# From metabarcoding to metaphylogeography: separating the wheat from the chaff

**DOI:** 10.1101/629535

**Authors:** Xavier Turon, Adrià Antich, Creu Palacín, Kim Præbel, Owen Simon Wangensteen

## Abstract

Metabarcoding is by now a well-established method for biodiversity assessment in terrestrial, freshwater and marine environments. Metabarcoding datasets are usually used for α- and β-diversity estimates, that is, interspecies (or inter-MOTU) patterns. However, the use of hypervariable metabarcoding markers may provide an enormous amount of intraspecies (intra-MOTU) information - mostly untapped so far. The use of cytochrome oxidase (COI) amplicons is gaining momentum in metabarcoding studies targeting eukaryote richness. COI has been for a long time the marker of choice in population genetics and phylogeographic studies. Therefore, COI metabarcoding datasets may be used to study intraspecies patterns and phylogeographic features for hundreds of species simultaneously, opening a new field which we suggest to name metaphylogeography. The main challenge for the implementation of this approach is the separation of erroneous sequences from true intra-MOTU variation. Here, we develop a cleaning protocol based on changes in entropy of the different codon positions of the COI sequence, together with co-occurrence patterns of sequences. Using a dataset of community DNA from several benthic littoral communities in the Mediterranean and Atlantic seas, we first tested by simulation on a subset of sequences a two-step cleaning approach consisting of a denoising step followed by a minimal abundance filtering. The procedure was then applied to the whole dataset. We obtained a total of 563 MOTUs that were usable for phylogeographic inference. We used semiquantitative rank data instead of read abundances to perform AMOVAs and haplotype networks. Genetic variability was mainly concentrated within samples, but with an important between-seas component as well. There were inter-group differences in the amount of variability between and within communities in each sea. For two species the results could be compared with traditional Sanger sequence data available for the same zones, giving similar patterns. Our study shows that metabarcoding data can be used to infer intra- and interpopulation genetic variability of many species at a time, providing a new method with great potential for basic biogeography, connectivity and dispersal studies, and for the more applied fields of conservation genetics, invasion genetics, and design of protected areas.

## INTRODUCTION

Metabarcoding, whereby information on species present in a variety of communities can be obtained from so-called environmental DNA (eDNA), or from bulk or community DNA (Creer et al 2016, Macher et al 2018), is by now established as a robust method for biodiversity assessment (Baird & Hajibabaei 2012, Deiner et al 2017, Taberlet et al 2018, Adamowicz et al 2019). Metabarcoding provides a fast and accurate method for measuring biodiversity, allowing identification of many more taxa (Molecular Operational Taxonomic Units or MOTUs) than morphological methods (Dafforn et al 2014, Cowart et al 2015, Elbrecht et al 2017). Further, metabarcoding is independent of taxonomic expertise, which is dwindling worldwide (Wheeler et al 2004), albeit it is highly dependent on the completeness of reference databases to reliably assign taxonomic names to MOTUs (Cowart et al 2015, Briski et al 2016). Biodiversity assessment, detection of invasive or endangered species, paleoecological reconstruction or diet analyses are among the main applications of metabarcoding to date (e.g., Ji et al 2013, Pochon et al 2013, Kelly et al 2014, Hajibabaei et al 2016, Ficetola et al 2018). All of them are highly relevant for basic biodiversity research and for establishing management policies. There is, however, more information in metabarcoding datasets than just α- and β-diversity related issues. Further exploitation requires a shift from interspecies genetic patterns, that constitute most of the metabarcoding applications so far, to intraspecies genetic patterns (Adams et al 2019), making use of the within-MOTU genetic variability uncovered by metabarcoding.

Being heirs to studies in prokaryotes, eukaryotic metabarcoding initially relied heavily on ribosomal RNA sequences for MOTU delimitation. These sequences lack variability for within-MOTU studies in many groups, particularly metazoans (Tang et al 2012, Leray & Knowlton 2016, Wangensteen et al 2018a). However, in recent years, intense efforts have been devoted to optimize the use of mitochondrial DNA (mostly COI) in metabarcoding (Andújar et al 2018). The use of mitochondrial sequences was hindered by the lack of universal primers (Deagle et al 2014), but new sets of COI primers for general purposes or for specific groups (Leray et al 2013, Elbrecht & Leese 2017, Vamos et al 2017, Gunther et al 2018) are overcoming this problem and COI sequences are now being increasingly used in general biodiversity studies (e.g. Leray & Knowlton 2015, Aylagas et al 2016, Macher et al 2018, Porter & Hajibabaei 2018a), where they typically uncover a much higher degree of α-diversity than rDNA (Stefanni et al 2018, Wangensteen et al 2018a,b). Furthermore, the use of COI opens the door to taxonomic assignment using the extensive database of the Barcode of Life Datasystems (BOLD), which is continuously increasing in depth and coverage (Ratnasingham & Hebert 2007, Porter & Hajibabaei 2018b).

COI sequences have been extensively used in studies of population genetics and phylogeography of terrestrial, freshwater, and marine organisms (Avise 2009, Emerson et al 2011). The shift to COI-based metabarcoding (Andújar et al 2018), therefore, implies the generation of databases containing an untapped reservoir of intraspecies variation that can allow characterizing intra- and inter-population genetic features of many species (MOTUs) simultaneously. This could constitute a gigantic leap from the current single-species studies, effectively opening a new field in population genetics for which we suggest the name of metaphylogeography.

The possibility of using metabarcoding for population genetics was hinted at by Bohmann et al (2017) and Adams et al (2019), but has been hardly developed. Current instances are in general preliminary, proof of concept, applications. For instance, within population genetic structure using bulk DNA has been assessed for ichthyosporean parasites of the cladoceran *Daphnia* (González-Tortuero et al 2015), or for a *Xyleborus* beetle collected at two locations with differing management practices (Pedro et al 2017). In the marine realm, Sigsgaard et al (2016), reliably obtained haplotype data in a whale shark aggregation from seawater eDNA. In invasion biology, eDNA was proven useful to assess native vs non-native strains of common carp in Japan (Uchii et al 2016).

An integrated phylogeography encompassing a range of species would be a powerful tool to investigate landscape-level processes (either natural or anthropogenic), over and above the signal given by each species. Studies that combine population genetics data on adamomultiple species by traditional methods are costly and usually involve just a handful of species (e.g., Haye et al 2014). The alternative is to use meta-analyses to collate the information scattered in different works (e.g., Zink 2002, Pascual et al 2017), or to use the information contained in georeferenced genetic databases (Gratton et al 2017). However, the pace at which climate change affect our ecosystems and the projected increased exploration of our resources in the coming decades urge for increased knowledge of population structure and phylogeography at the global biome level. The potential of metaphylogeography ranges from basic questions about biogeography, connectivity and dispersal patterns to more applied fields of conservation genetics, invasion genetics, and protected areas design. Nowadays, the consideration of multispecies genetic conservation objectives is seen as crucial to preserve community-wide genetic and evolutionary patterns (Vellend et al 2014, Nielsen et al 2017).

The main problem for the application of eDNA or community DNA to analyse intraspecies patterns lies in the fact that metabarcoding generates a high number of reads containing sequencing errors, which can occur at different steps in the procedure. Reads obtained by amplification and sequencing can be thought of as a “cloud” of erroneous sequences surrounding the correct one (Edgar & Flyvbjerg 2015). Sequencing errors will typically occur as low abundance reads with one or few base changes, while errors during amplification (PCR point errors, chimeras) have the potential of generating “daughter clouds” as they can reach higher read abundances (Edgar & Flyvbjerg 2015). As erroneous sequences in general diverge very little from the true sequences, they are often incorporated into the right MOTU during the clustering step, thus reducing potential impacts on the results of “standard” metabarcoding approaches. However, they can severely bias intraspecies genetic patterns by artificially inflating the true haplotype diversity. Thus, separating the “wheat” (true sequences) from the “chaff” (false sequences) is the main challenge for the application of metabarcoding data to metaphylogeography.

To our knowledge, the problem of the correct assessment of intraspecific genetic diversity from community DNA in complex samples has been explicitly addressed only in a recent work by Elbrecht et al (2018a). Using a single-species mock sample with known Sanger-sequenced haplotypes, they assayed a combination of denoising procedures to reduce the number of spurious haplotypes obtained using a metabarcoding pipeline. They then applied the best performing strategy to natural samples of freshwater invertebrates, deriving population genetic patterns for some of the species present.

We sought here to develop a practical strategy to make metabarcoding datasets amenable to phylogeographic studies. There is an ever-increasing number of such datasets publicly available in repositories. Mining COI-metabarcoding data has been suggested for species discovery (Porter & Hajibabaei 2018b), and these databases can be a resource for phylogeography as well. These data comprise different information, from raw sequences to filtered and paired sequences to simply MOTU tables. In many cases, no ground truth data or mock community analyses exist for them. We therefore need a strategy for cleaning noisy databases in the absence of ground truth information. We contend that the properties of coding sequences such as COI can provide such a strategy. Indeed, coding DNA sequences have naturally a high amount of variation concentrated in the third position of the codons, while errors at any step of the metabarcoding pipeline would be randomly distributed across codon positions. Examination of the change of diversity values (measured here as the entropy of each position, Schmidt & Herzel 1997) as we eliminate noisy sequences can therefore guide the choice of the best cleaning parameters in the presence of an unknown amount of noisy data.

A parallel inspection of the distribution of sequences across samples is also necessary. Error-containing sequences will typically co-occur in the same sample with the correct sequence, albeit with less abundance, and co-occurrence patterns can be incorporated to detect these sequences in cleaning steps. At the same time, while error sequences are likely to appear randomly in the samples, true sequences should feature a given ecological distribution, meaning that a sequence appearing in all replicates of a community, for instance, is unlikely to be an error. Distribution patterns of sequences have been suggested to guide MOTU calling or MOTU curating procedures (Olesen et al 2017, Froslev et al 2017), but have not been applied, to our knowledge, for within-MOTU sequence curation.

Combining patterns of variation in entropy and sequence distribution patterns can lead to meaningful ways to reduce noisy datasets to operational datasets. This approach can be used to generate customized procedures for each different study system that take into consideration its particulars (replication level, pre-filtering applied, clustering procedure). It only requires that, for a given study, the information about which sequences have been pooled in each MOTU in the clustering step - with their sample distribution – is provided.

We want to point out that the “metaphylogeography” concept is not equivalent to “conventional phylogeography of many species” and we therefore need to adapt some definitions. In particular, relative frequencies of reads of the different haplotypes are available instead of the relative frequencies of individuals bearing these. Both are unlikely to be equivalent. The high difference in number of reads that can be obtained in metabarcoding can easily reach orders of magnitude and is hardly representative of conventional frequencies based on the number of individuals bearing a particular haplotype. Further, the quantitative value of metabarcoding data is debatable (Elbrecht & Leese 2015, Wares & Pappalardo 2016, Piñol et al 2018). Once we have a curated dataset, we suggest to perform phylogeographic inference using a semiquantitative abundance ranking applied within each MOTU, as a compromise between a strictly quantitative interpretation of the data on one hand, and losing all the information contained in the number of reads on the other. For comparative inference, the traditional analytical framework including haplotype networks, AMOVA, and the like, is perfectly valid if one keeps in mind these differences in the interpretation of results.

In the present study, we developed cleaning strategies to make community data derived from COI amplicon sequencing amenable to the analysis of intraspecific variation. As a case study we used a COI-based metabarcoding survey of biodiversity of sublittoral marine benthic communities. We then extracted phylogeographic trends from the MOTUs obtained with the best pruning parameters selected. We finally compared results with those of traditional phylogeographic studies for two species for which information exists for the same (or nearby) sampling areas. Our general goal was to show the feasibility of the metaphylogeographic approach using a “standard” metabarcarcoding dataset obtained from natural samples.

## MATERIAL AND METHODS

### Dataset

The dataset consisted of COI-based biodiversity data obtained from benthic marine communities in two Spanish National Parks, one in the Atlantic and one in the Mediterranean (Appendix S1: Fig. S1). The dataset has different replication levels: over time (two years), within communities, and within samples (size fractions). Sample collection and processing followed Wangensteen & Turon (2017) and Wangensteen et al (2018a). Appendix S1 provides further information about the communities, collection techniques, and laboratory and bioinformatics procedures to generate the dataset.

For the metaphylogeographic approach, we selected MOTUs that had at least two sequences. We also required that the MOTU appeared in the two Parks with 20 or more reads in each one, and appeared at least once in each of the two study years. Note that this MOTU selection does not imply that discarded MOTUs are artefacts, but simply that they are not useful for population genetics inference (e.g., one MOTU appearing only in a given community, even if abundant).

Using the list of retained MOTUs, the original sequence file, and the information of which sequence belongs to each MOTU (contained in the output of the clustering program used to generate MOTUs), we obtained separate MOTU files containing, for each MOTU, all sequences included with their abundances in the different samples. We then aligned sequences within each MOTU with the msa R package (Bodenhofer et al 2015), and misaligned sequences, likely due to slippage of degenerate primers (Elbrecht et al 2018b), were detected and eliminated.

### Simulation analysis

All data manipulation and analyses were conducted using R software (R_Development_Core_Team 2008). To avoid confusion between different terms, sometimes used interchangeably, we will use the name denoising to refer to any procedure that tries to infer which sequences contain errors and merges their reads with those of the correct “mother” sequence. We will call filtering any method that actually deletes sequences from the dataset, based on abundance thresholds or otherwise. Clustering will refer to any procedure for combining sequences - without regard to whether they are correct or not - into meaningful MOTUs.

We ran a simulation study to infer the best cleaning strategy and the best parameters for our data. The rationale was to start with a known dataset, introduce sequencing errors, and clean it again to recover the original dataset. Following Wang et al (2012), we considered that the 1,000 sequences with highest frequency in our dataset were error-free, and used them for parameter estimation on a dataset representative of our actual sequences. For this simulation we didn’t keep the ecological information, and used just the total number of reads of each of these 1,000 top sequences.

We simulated that these allegedly correct amplicons were sequenced with error rates between 0.001 and 0.01 per base, bracketing values published for HTS sequencers and, in particular, for the MiSeq platform (Schirmer et al 2016, Pfeiffer et al 2018). For simplicity, we assumed a constant error rate for all bases in a sequence.

For the highest error rate (0.01) we then denoised the resulting sequences using a procedure adapted from the algorithm of Edgar (2016). We merged the reads of presumably incorrect daughter sequences with those of the correct mother sequences if the number of sequence differences (d) is small and the abundance of the incorrect sequence with respect to the correct one (abundance ratio) is low. The higher the number of differences, the lower the ratio should be for the sequences to be merged. This was formalized by the expression (Edgar 2016):

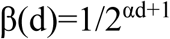

Where β(d) is the maximum abundance ratio allowed between two sequences separated by d changes so that the less abundant was merged with the more abundant. The α parameter is user-settable to seek a compromise between accepting as correct erroneous sequences (high α values) or merging true sequences (low α values). The denoising was done for values of α from 10 to 1.

We analysed changes in diversity of the different codon positions as we introduced increasing levels of noise (erroneous reads) and as we denoised the dataset with increased stringency (lower α values). As a measure of diversity we used the Shannon entropy value computed with the R package *entropy* (Hausser & Strimmer 2009). We expected that random error will increase more the entropy of the less variable position (second position of the codons) and less the entropy of the third, more variable, position. Thus, the entropy ratio (hereafter E_r_):

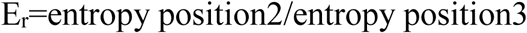

was expected to increase as simulated error rates increased and to decrease when denoising. After each round of denoising we noted the number of original sequences remaining, the number of noisy sequences remaining, and the entropy ratio of the sequences. We expected that at some value of α the E_r_ will reach the original value and remain more or less constant afterwards. As at this point many erroneous sequences remained in the dataset (see Results), we completed the simulation with a filtering procedure in which low frequency sequences were eliminated.

We assayed a range of minimal number of reads to keep a sequence and looked at the number of original and noisy sequences remaining, as well as their entropy ratio. As before, we expected the Er to decrease markedly and stabilize after some threshold is reached. The best α parameter and the best minimal number of reads should allow us to recover most of the original sequences with as few erroneous sequences as possible.

### Dataset cleaning

The cleaning procedure followed the findings of the simulation and was therefore based on two steps: denoising (without loss of reads) and filtering by minimal abundance (with loss of reads). We applied denoising within defined MOTUs, under the assumption that most erroneous sequences would have been included in the same MOTU as the correct sequence, and thus sequence distances and abundances, a key part of the denoising algorithm, are more meaningful if compared within MOTUs. Once denoising was complete and, thus, all “salvageable” sequences had been merged with the correct sequence, the second step consisted of an abundance filtering, in which low-abundance sequences, likely erroneous, “surviving” the denoising step were eliminated.

During the previous steps, co-occurrence patterns were used to avoid merging or eliminating sequences whose sample distribution and co-occurrence patterns suggested they were not artifacts (f.i., sequences that do not co-occur with similar sequences will not be merged with them, and sequences found in all replicates of a community will not be filtered out). The use of distribution data can reduce the risk of eliminating true sequences, particularly when they are present at low abundances (e.g., reflecting a low biomass of the organism).

To allow a daughter sequence presumed to be a sequencing error to be merged with a more abundant mother sequence, we required that the former co-occurs with the latter. This is formalized by a co-occurrence (C_occ_) ratio in the form:

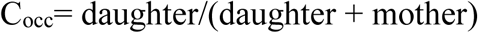

were daughter= n. of samples with only the daughter sequence, and daughter+ mother= number of samples with the daughter and the mother sequence.

We set this parameter to a value of 1 (i.e., whenever a daughter sequence was present, the mother sequence should be present in the same sample). Any “daughter” sequence with co-occurrence ratio <1 was considered a genuine sequence and was not merged. This stringent value was used considering that we enforce the presence at the sample level, and not at the fraction level, which means that the sequence needs to be present in just one of the three fractions of the sample. In preliminary assays, changing C_occ_ influenced the number of sequences retained, but represented little change in the entropy ratios obtained. In addition, in the filtering step sequences appearing in all replicates of a given community were considered correct and not filtered out, even if present at low abundance.

Taking these distribution patterns into consideration we applied the denoising and filtering steps. A diagrammatic representation of the pipeline used is presented in Fig. 1. Denoising was performed at α values between 10 and 1, and for the best-performing α, filtering was done for increasing minimal numbers of reads from 2 to 100. After each round of sequence denoising or filtering, the MOTUs were examined and retained only if they still met the requirements of having at least two sequences, appearing in the two Parks with 20 or more reads in each one, and appearing at least once in the two study years. The changes in E_r_ of the retained MOTUs were examined over the range of α and minimal abundance values. In both cases the entropy ratio should decrease and, following the simulation results, the points where it became stabilized (i.e., where the slope fell below 0.005) were used as optimal parameter cut-offs.

**Figure 1.**
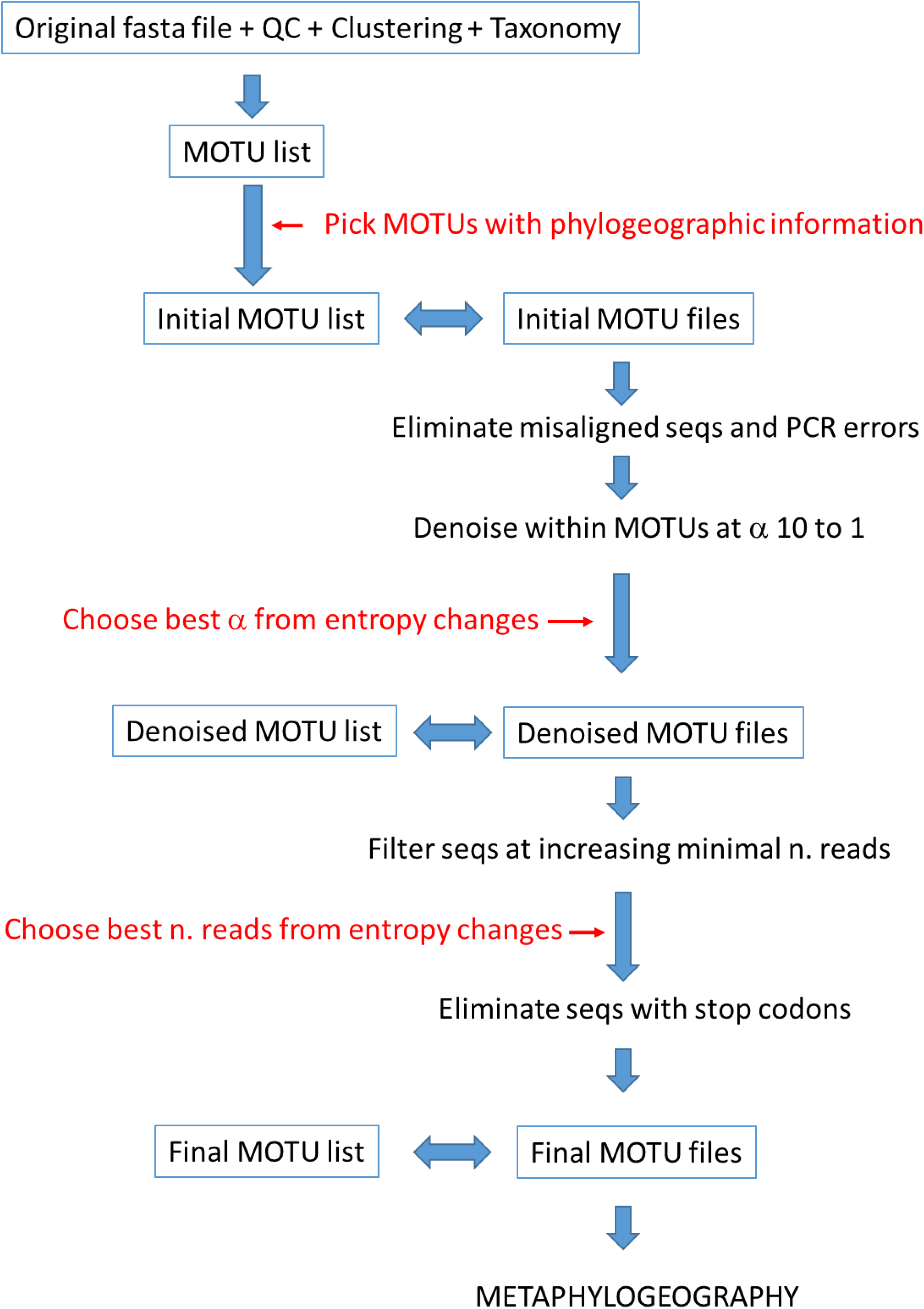
Schematic representation of the pipeline followed in this study. See Methods for details.

Finally, even if sequences retained were mostly correct, they can still include a number of true variants due to heteroplasmy or numts (Elbrecht et al 2018a). However, numts tend to accumulate mutations resulting in stop codons and can therefore be detected by inspecting the sequences retained. As the dataset included many different eukaryotic groups with different genetic codes, we adopted a conservative approach. For each MOTU, we tried the 18 genetic code variants stored in the Biostrings R package (Pagès et al 2018) and used the *translate* function to obtain the corresponding aminoacid sequence. We then chose, for each MOTU, the genetic code giving the lower number of stop codons (often several code variants resulted in no stop codons). The MOTUs denoised with the optimal α-value (first step), once filtered with the optimal abundance cut-off (second step) were checked for the presence of stop codons and the sequences presenting them were removed from the dataset. The remaining MOTUs and sequences constituted the curated dataset for further analyses (Fig. 1).

### Metaphylogeographic analyses

We performed network analyses with function *HaploNet* of the R package pegas (Paradis 2010). We used function *amova* of the R package ade4 (Dray & Dufour 2007) to compute analyses of molecular variance (AMOVA) in order to ascertain the percent variation associated with the hierarchical organization of the samples. For AMOVA, we used the proportion of the different sequences present (option distances=NULL). Preliminary assays considering also sequence distances (not just sequence frequencies) gave highly similar results and were computationally slower.

In these analyses we needed to capture the quantitative information regarding frequencies of the different sequence variants. As mentioned above, using number of reads as a proxy for individual-based abundances can be misleading. We adopted a semiquantitative index based on Wangensteen et al (2018b) applied within each MOTU. To obtain this semiquantitative ranking we ordered the sequences of each sample in each MOTU by increasing number of reads and ranked them from 0 to 4, indicating that the sequence is either absent in that sample (rank 0) or falls in the following percentiles of the distribution of ordered sequences: rank 1, ≤50%, rank 2,> 50≤ 75%; rank 3,> 75 ≤90%, rank 4,> 90%. These semiquantitative ranks were used as proxies for haplotype abundances in the analyses.

### Comparison with previous studies

After examination of the curated MOTU dataset, we found only two species for which conventional phylogeographic analyses had been performed using COI information in the same geographic area: the sea urchin *Paracentrotus lividus* and the brittle star *Ophiothrix fragilis*.

For *Paracentrotus lividus*, we collated haplotype information from studies spanning the atlanto-mediterranean transition (Duran et al 2004), trimmed the sequences to the same fragment amplified herein, and compared the haplotypes with the ones encountered in our metabarcoding dataset. Duran et al (2004) included two populations close to our localities: Eivissa Island in the Balearic Archipelago, and Ferrol in the Galician coast. Networks were generated with the haplotypes found in these localities and compared with our results.

For *Ophiothrix fragilis*, our MOTU corresponded to Lineage II of Pérez-Portela et al (2013). This brittle star is in fact a complex of species, and Lineage II is likely a cryptic species (Taboada & Pérez-Portela 2016), but it remains unnamed so far. As before, we extracted haplotype information from all localities in Pérez-Portela et al (2013), spanning the atlanto-mediterranean area, and compared with our results. We also obtained haplotype networks for the two closest populations studied in that work: Alcudia in the Balearic Archipelago, and Ferrol in the Galician coast.

## RESULTS

### The dataset

The original dataset, once quality and length filtered, contained 25,772,264 sequences of 8,900,080 unique sequences. Without singletons, the numbers were reduced to 17,808,524 reads and 936,340 unique sequences. Following the pipeline of Wangensteen et al (2018a) with some modifications including clustering with SWARM2 (Mahé et al 2015) (see Appendix S1), we obtained a MOTU list of 26,561 eukaryote MOTUs. Of these, 13,410 MOTUs were present only in the Mediterranean site, 8,247 only in the Atlantic locality, and 4,904 were shared by both basins. Of the latter, only 722 MOTUs (with a total of 362,177 unique sequences and 9,430,236 reads) fulfilled the conditions of having at least two sequences, being present in the two Parks with at least 20 reads in each one, and having appeared in the two years of study. After checking the alignment, only 158 sequences, comprising 689 reads, appeared as misaligned, mostly as a result of one bp slippage, and were removed. The singleton-free fasta sequence file (paired, demultiplexed, and quality-filtered), the original MOTU list, and the output of the SWARM analyses have been uploaded as a Mendeley dataset (DOI: http://dx.doi.org/10.17632/xpmtvn2k7m.1). The 722 MOTUs selected for the study are listed in Data S1, together with their taxonomic assignment and abundance (number of reads) per sample. The actual sequences of each MOTU, with their abundances per sample, are available at the Mendeley dataset.

### Simulation study

In our case, the top 1,000 sequences in the 722 MOTUs dataset contained 5,948,135 reads. The entropy values of the codon positions of these sequences were: first position, 0.4298±0.037 bits (mean±SE); second position, 0.1833±0.028 bits; third position, 0.9256±0.023 bits. The simulation of increasing sequencing error rates clearly increased the entropy of the three positions (Fig. 2A), but more so for the less variable second position, which increased its value ca. 30% at the highest error rate. On the other hand, the third position increased entropy only about 1.8%. As a result, the entropy ratio (E_r_, entropy2/entropy3) increased linearly with error rate, from 0.198 to 0.252 (Fig. 2B).

**Figure 2.**
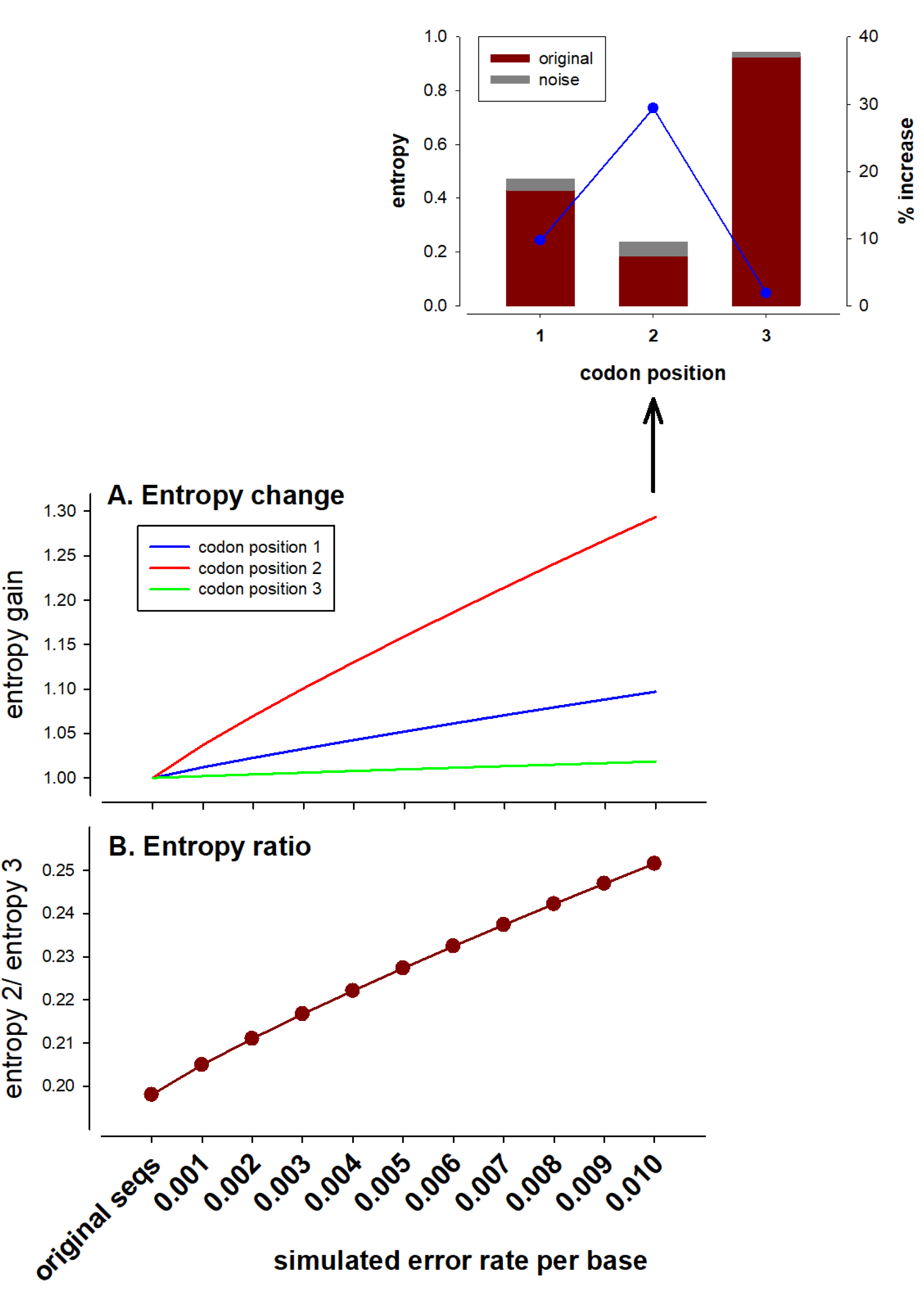
Simulation analysis. (A) relative increase (initial value=1) of the entropy values of each position at increased error rates. Barplot shows the original and added entropy of each position at the highest (0.01) error rate. (B) change in the entropy ratio.

We then used the “noisy-most” dataset, the one simulated at the highest (0.01%) error rate. It had the same original number of reads, but 5,141,683 erroneous sequences (besides the 1,000 correct ones) were generated. For coherence with the global dataset used, singletons were removed, leaving 144,791 sequences. This dataset was then denoised at α values between 10 (least stringent) and 1 (most stringent). The E_r_ decreased drastically at the initial steps, concomitantly with a decrease in the number of erroneous sequences (Fig. 3A). The E_r_ value of the simulated dataset reached the original value at α between 6 and 5.

**Figure 3.**
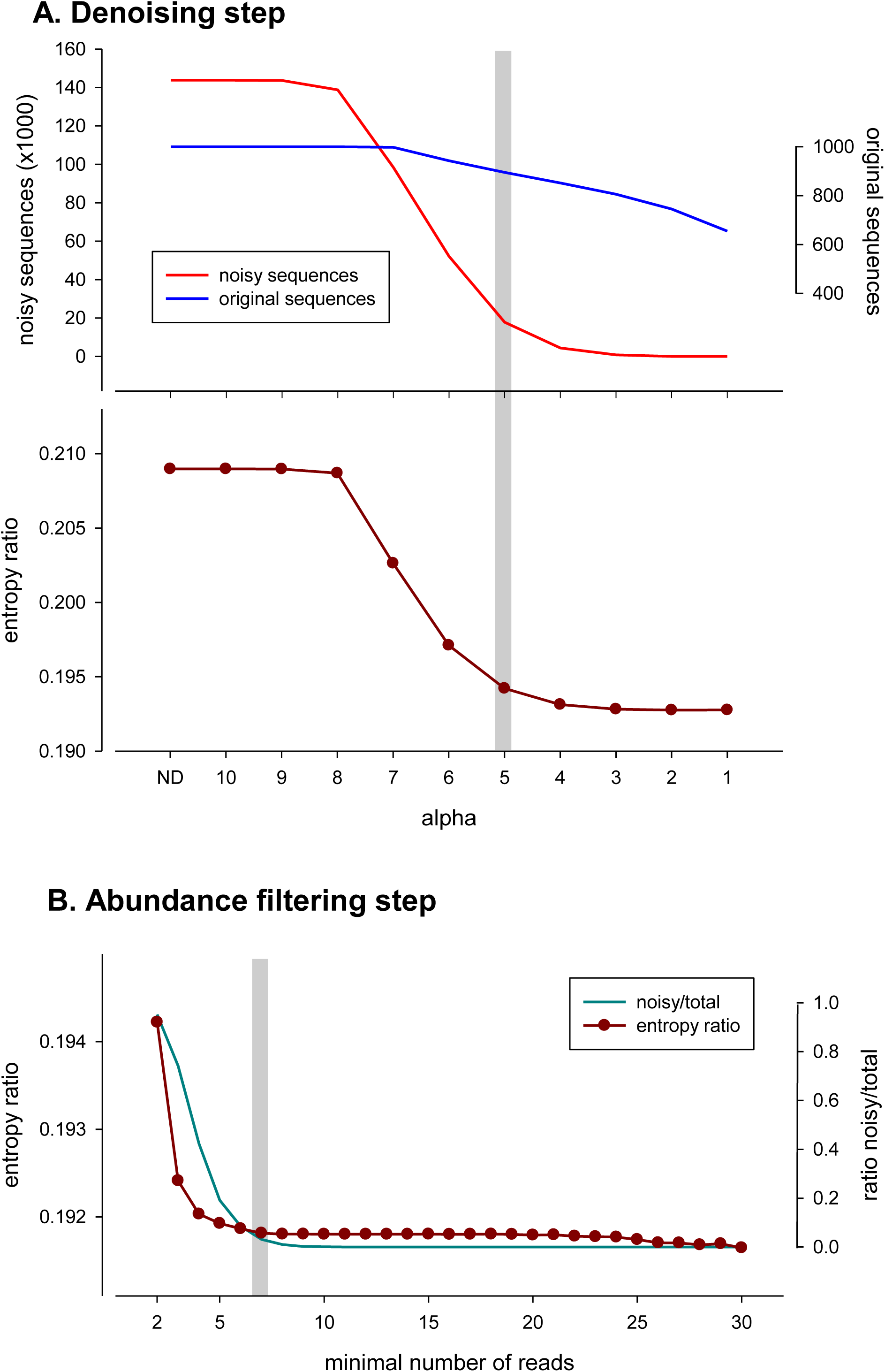
Simulation analysis. (A) variation in the number of original and erroneous (“noisy”) sequences and entropy ratio at decreasing values of the alpha parameter of the denoising algorithm (ND: no denoising). (B) change in the entropy ratio and in proportion of noisy vs original sequences after filtering the dataset by minimal abundance. The gray bars indicate the selected values of alpha (5) and minimal number of reads (7).

Taking the more conservative α=5, which is also the point where the curve leveled off (slope <0.005), we found that the dataset contained 895 of the original sequences and 17,799 erroneous sequences. In other words, while ca. 10% of the original sequences have been incorrectly merged, there remained still a high number of errors in the dataset. Using only the denoise procedure, we got completely rid of erroneous sequences only at α=1. But at this value only 66% of the correct sequences were retained.

We therefore applied a round of filtering by minimal number of reads to the dataset denoised at α=5. Again, the E_r_ decreased sharply at increasing thresholds of minimal reads, following the elimination of erroneous sequences (Fig. 3B), and stabilized clearly at 7 reads (Fig. 3B). The combination of denoising (α=5) and filtering (minimal abundance=7) allowed us to recover 924 sequences, of which 895 (97%) were among the 1,000 original sequences and only 3% were erroneous sequences. The frequency distribution of the number of reads in both the original (1000) and the recovered (924) sequences was almost identical (not shown). Importantly, the shape of the E_r_ curve, concretely the stabilization points, proved informative to select the cut-points for the two variables.

### Dataset cleaning

As a first step, we tried to identify PCR errors during amplification, as they can result in abundant sequences and be more difficult to spot. We assumed that PCR errors will affect one nucleotide at most, will be rare, will occur in the same sample as the original sequence, and will be abundant. Therefore, we looked within the 722 MOTUs for sequences differing by one nucleotide from a more abundant one, co-occurring always with it, being present in at most 3 samples, and having an abundance of >200 reads. Only 14 such sequences were identified and merged with the more abundant ones, so PCR errors did not seem to play an important role in our study.

After applying to the whole dataset of 722 MOTUs the denoising step for α values from 10 to 1 and a co-occurrence index of 1, we examined the change in number of retained MOTUs and entropy ratio (Fig. 4A). The number of MOTUs remained constant but started decreasing at α=6. As expected, the E_r_ decreased fast at first and more slowly at lower α-values (i.e., with higher merging power) (Fig. 4A). The curve leveled off (slope below 0.005) at α=5, with only a slight loss of MOTUs (6 out of 722). We thus retained α=5 as the optimal denoising parameter.

**Figure 4.**
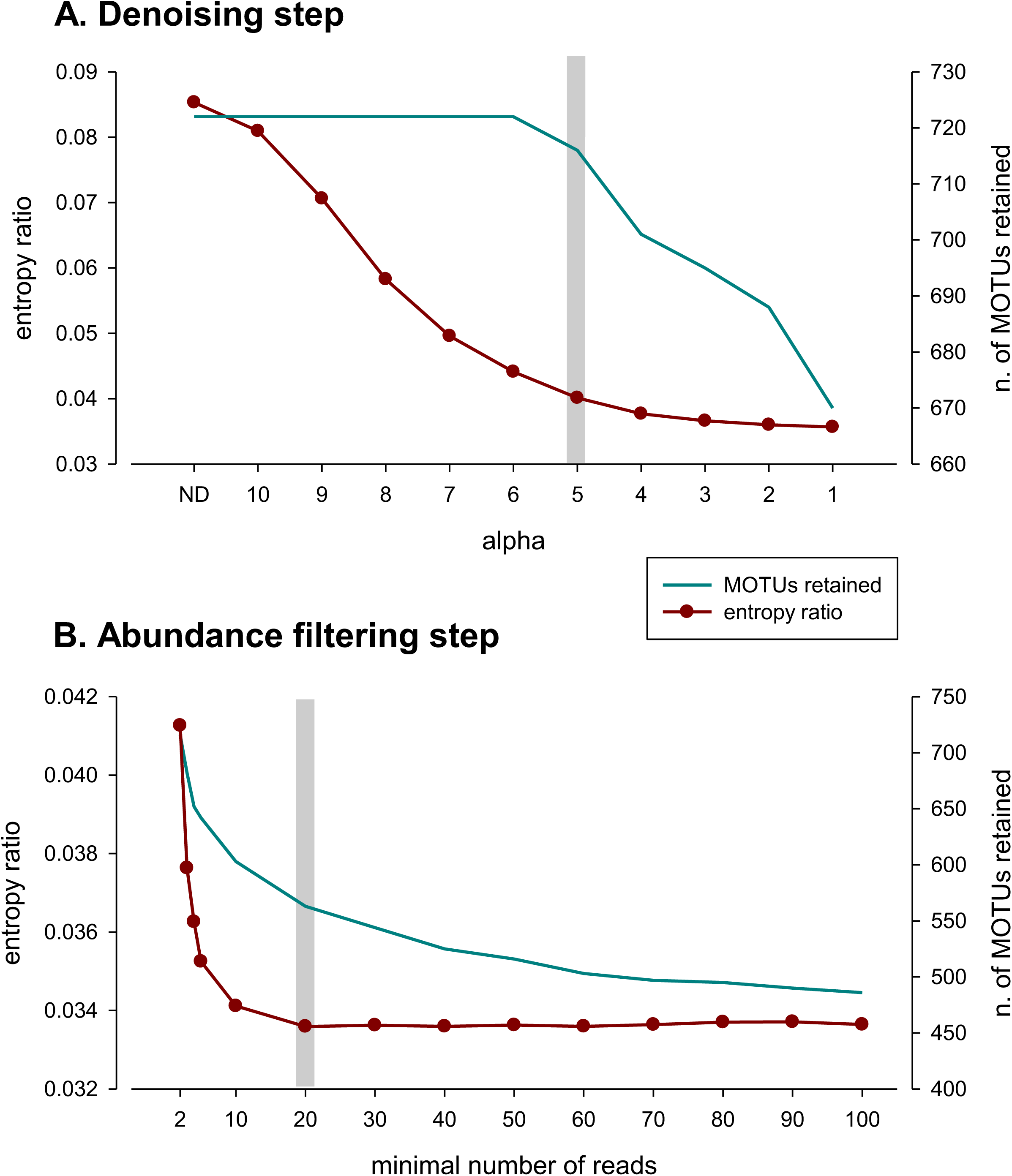
(A) variation in the number of sequences and number of MOTUs remaining at decreasing values of the alpha parameter (ND: no denoising) of the denoising algorithm. (B) change in the entropy ratio and, (C) change in residual (within-sample) variance of the amova model. The gray bars indicate the selected alpha value (5) and abundance threshold (20).

The MOTU list corresponding to the denoised dataset had 716 MOTUs, with 49,995 sequences (86% of the original sequences had been merged) and 9,426,339 reads (Data S1). The corresponding MOTU files (available at the Mendeley dataset http://dx.doi.org/10.17632/xpmtvn2k7m.1) were submitted to an abundance filter, with a threshold from 2 to 100 reads. There was a decrease the number of MOTUs retained at increasing minimal number of reads, particularly in the interval 2-50 (Fig. 4B). The entropy ratio fell markedly and became stabilized at a value of 20 reads, after which it remained more or less constant (Fig. 4B). Thus, 20 reads was used as a minimal abundance threshold.

The sequences of the resulting MOTU files were translated and checked. Only 8 sequences had stop codons and were eliminated. The final MOTU list thus consisted of 563 MOTUs, with 7,206 sequences and 8,912,772 reads (Data S2). The final MOTU files were uploaded to the Mendeley dataset http://dx.doi.org/10.17632/xpmtvn2k7m.1

As for the taxonomy assigned, the most diverse groups of Eukarya in the final dataset were Rhodophyta (91 MOTUs), Stramenopiles (90 MOTUs, mostly diatoms and brown algae), and Metazoa (273 MOTUs) (Data S2). A total of 99 eukaryotic MOTUs remained unassigned taxonomically (identified as Eukarya). Among metazoans, 112 MOTUs were assigned a species-level taxon, while 225 MOTUs were assigned at least at the phylum level and 48 MOTUs remained unassigned (Data S2). The phyla of metazoans identified in the final MOTU list were: Annelida (34 MOTUs), Arthropoda (56 MOTUs), Bryozoa (17 MOTUs), Chordata (8 MOTUs), Echinodermata (7 MOTUs), Mollusca (22 MOTUs), Nemertea (6 MOTUs), Porifera (30 MOTUs), and Xenacoelomorpha (1 MOTU).

Further analyses concentrated in the major groups detected, which accounted for 437 of the 464 MOTUs that could be assigned: red algae (Rhodophyta), diatoms (Bacillariophyta), brown algae (Phaeophyceae) and metazoans (Metazoa). In the latter, phylum-level analyses were performed.

### Phylogeography

Network graphs of the MOTUs (Appendix S2) showed different patterns, albeit in most cases one or a few haplotypes appeared as the most abundant, linked to a varying number of low abundance haplotypes. Some selected instances are presented in Fig. 5, showing also the change in network shape along the process of cleaning. It can be seen that the major pruning effect was due to the initial denoising step.

**Figure 5.**
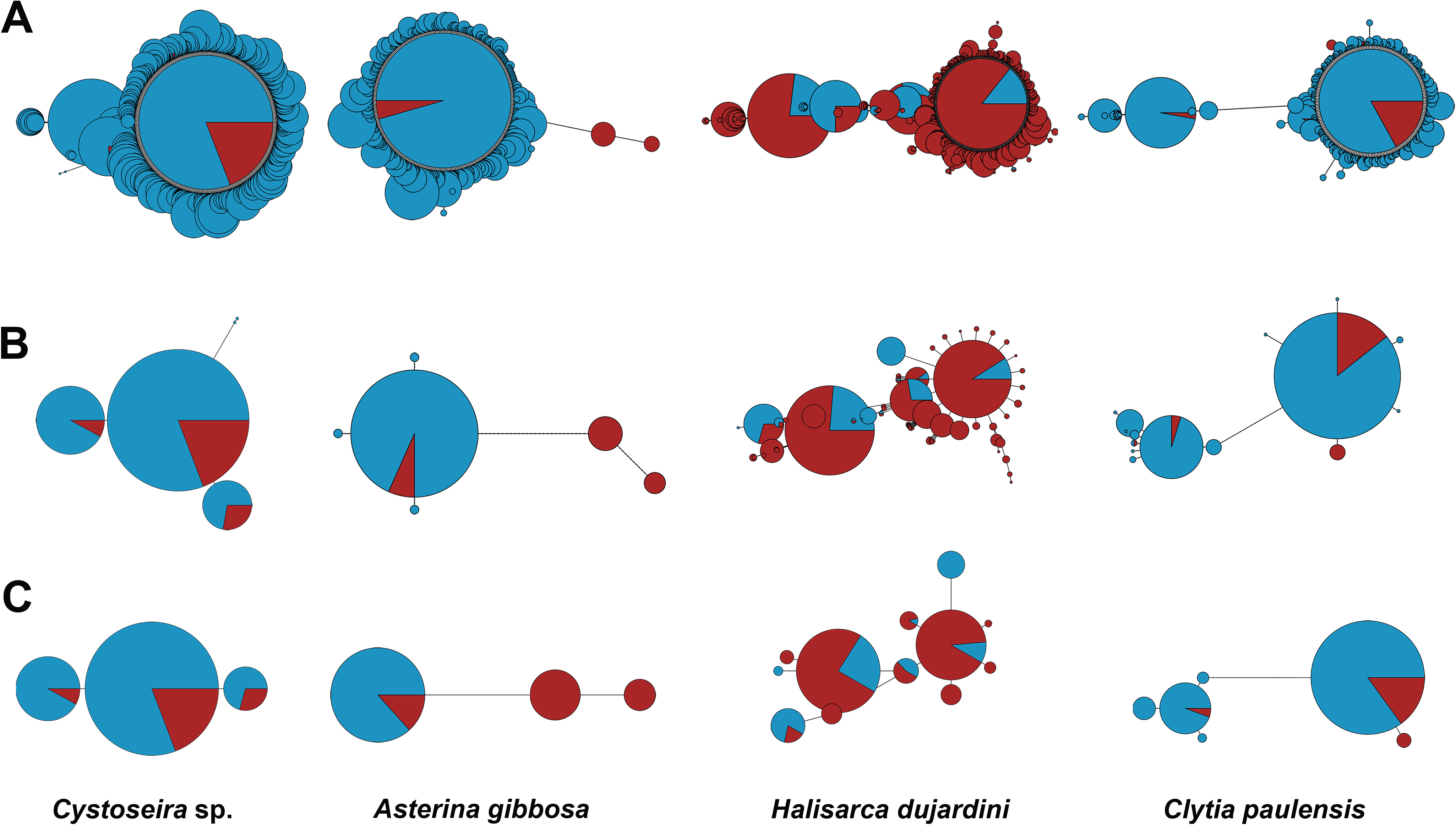
Selected instances of networks obtained at different stages of the pipeline. (A) without filters; (B) after denoising at alpha=5; (C) after denoising at alpha=5 plus minimal abundance filtering (threshold 20 reads). Circles represent haplotypes, and their diameters are proportional to their abundance (in semiquantitative ranks) in the samples. Blue color represent abundance in Mediterranean samples, red color in Atlantic samples. Length of links is proportional to the number of changes between haplotypes. Note that Figs in A, B, and C are not drawn to the same scale. The names correspond to the taxonomical identification of the MOTUs with ecotag. The MOTU ids (as per Data S1) are, from left to right: 143, 1740, 2500, 25366.

AMOVAs were used to partition the genetic variance hierarchically into components due to the differences between seas, between communities within seas, between samples (replicates) within communities, and within samples. The average values of these variance components for the major groups detected, and for metazoan phyla separately, displayed a clear overall trend: genetic variance was concentrated within samples (60-75%) in all major groups (Fig. 6A). The other components of variance followed a decreasing trend, with a remarkable variance associated to differentiation between the two seas (14-25% of variance), and smaller variance between communities within each sea, and even lower between replicate samples of a given community. The latter component was almost negligible (<1.2%) in the non-metazoan groups considered, but reached 5.4% in metazoans. The different components were compared across groups with ANOVA (followed by Student-Newmann-Keuls *post-hoc* tests if significant). The between sample component was significantly higher (all p<0.001) in metazoans than in the other groups. For the other components the values were in general comparable, the only significant differences being a higher between seas differentiation in diatoms than in metazoans, and a higher within sample variance in red algae than in diatoms.

**Figure 6.**
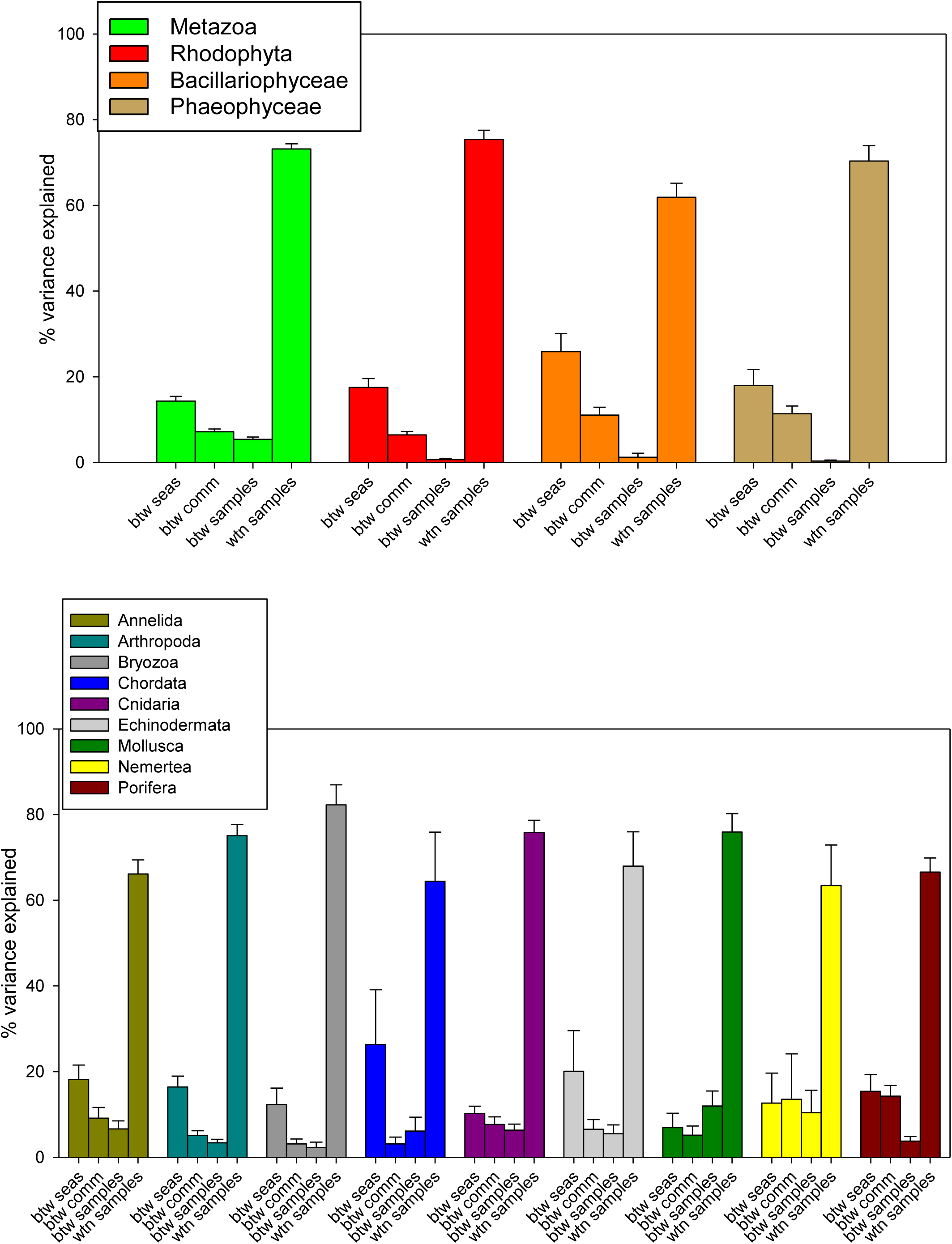
Summary of the mean % of variance explained by the hierarchical structure of the AMOVA. (A) as per eukaryote groups; (B) per metazoan phyla. Bars are standard errors. (btw seas: between seas, btw comm: between communities within seas; btw samples: between samples within communities, wtn samples: within samples)

Metazoans therefore showed a higher heterogeneity between replicate samples of a given community than the other groups. When examined across phyla (Fig. 6B), albeit the overall trend was in general maintained - a dominant within sample component and a variance between seas > between communities > between samples - there were exceptions. In particular, molluscs had a high between sample variability, and other groups presented important small-scale (between communities and/or between samples) variability as compared to the between seas differentiation (Cnidaria, Nemertea, Porifera). ANOVA showed few significant differences between phyla, the only significant comparisons involving the between samples component in molluscs, which was significantly higher than in bryozoans or sponges.

As for the comparison with previous studies, MOTU 697 was identified as the sea urchin *Paracentrotus lividus* with 100% sequence identity. This MOTU had 15 sequences. This species has an atlanto-mediterranean distribution, and Duran et al (2004) analysed populations spanning the W Mediterranean and NE Atlantic with COI. In that work, 65 different haplotypes (of a longer fragment of COI) were detected. Once trimmed to our sequence length and collapsed, there were 32 remaining haplotypes. 9 out of the 15 sequences detected herein had already been found by Duran and co-workers, while the remaining 6 were new.

We then selected the haplotypes found in the previous work in the two localities closest to our sampling points (Eivissa in Balearic Islands and Ferrol in Galicia). There were 11 haplotypes (4 of which were also present in our MOTU). We performed a network with the 2004 information and compared it with the one obtained for MOTU 697 with our semiquantitative abundance rank (Fig. 7A, B). The two networks had a similar shape, with a highest abundance of haplotype 2 (named after the order of sequences obtained for this MOTU), followed by haplotypes 1, 3, and 6. For the shared haplotypes, the between seas distribution was the same in the two studies (1, 2, and 3 shared between seas, 6 present only in the Atlantic). An AMOVA with a randomisation test (n=1,000) of our MOTU 697, revealed a significant differentiation between seas, between and within samples (p<0.001), but not between communities (p=0.812).

**Figure 7.**
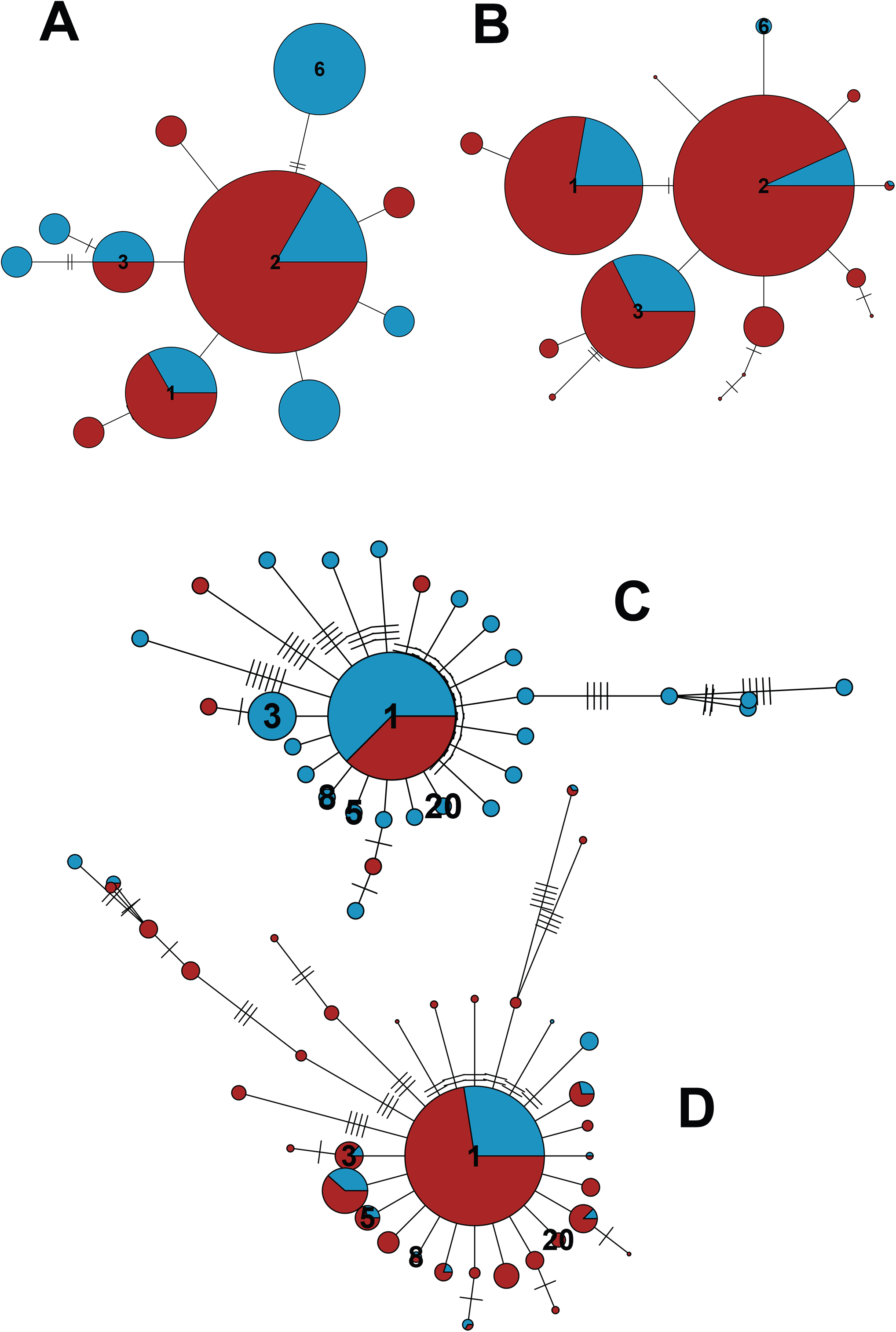
(A) network constructed with the 11 haplotypes of the sea urchin *Paracentrotus lividus* found by Duran et al (2004) in localities close to our sampling points and (B) network constructed with the 13 haplotypes comprising the MOTU corresponding to this species (id: 697). Haplotypes common to both studies are numbered. (C) network with the 29 haplotypes of the brittle star *Ophiothrix fragilis* identified by Perez-Portela et al (2013) in localities close to our sampling points, (D) network of the 34 haplotypes found in the present study in the MOTU corresponding to this species (id: 15396). Haplotypes common to both studies are numbered. The short slashes in the links between haplotypes represent mutational steps. Colors as in Figure 5.

The MOTU 15396, comprising 37 sequences, was identified (100% identity) with *Ophiothrix* sp. in Pérez-Portela et al (2013). In that work the authors studied a controversial species complex of the genus *Ophiothrix* in the European waters using 16S and COI. Our sequences corresponded to the Lineage II of *Ophiothrix fragilis* in that work, that spanned from Britanny to Turkey. Pérez-Portela et al (2013) reported 125 haplotypes of Lineage II which, once trimmed to our 313 bp length, resulted in 90 different haplotypes. When merged with our dataset, 9 out of 37 sequences in MOTU 15396 had already been found in the previous study, while another 28 were new.

As before, we selected in Pérez-Portela et al (2013), the two localities closest to our sampling points (Alcudia in Balearic Islands, and Ferrol in Galicia). There were 29 haplotypes in these localities, of which 5 were shared with our study. The corresponding networks (Fig. 7C, D) showed a star-shaped structure with a dominant haplotype 1 found in the two studies, with many low abundance sequences separated by one or a few mutations from the central haplotype and some longer branches. It is noteworthy that, in this case, the shared haplotypes do not have always the same inter-basin distribution, thus, haplotype 1 was present in both oceans, but haplotypes 3, 8, and 5 present only in the Mediterranean site in the previous work, appeared now in the two seas (it should be noted that haplotype 3 did appear in other Atlantic sites in Pérez-Portela et al 2013). Finally, haplotype 20 was present only in the Mediterranean site in Pérez-Portela et al (2013) and only in the Atlantic locality in the present work. An AMOVA with a randomisation test (n=1,000) of our MOTU 15396 showed a significant component of variation related to between and within samples genetic variability (p<0.001), but not between seas (p=0.729) or between communities within seas (p=0.212).

## DISCUSSION

In this study we have developed a method to apply metabarcoding datasets to the study of intraspecies patterns of many species at a time using a highly variable coding fragment (COI). An initial denoising step, aimed at merging erroneus sequences with the correct ones, was followed by an abundance filtering step aimed at removing the remaining erroneous sequences. We used information from the variability of the different codon positions – following a simulation study - to select the best parameter-values in the denoising and filtering steps. In addition, sample distribution information was used in the different steps to minimize loss of low abundance true sequences.

All cleaning procedures are a compromise between eliminating spurious sequences and losing true signal. In the benchmarking approach of Elbrecht et al (2018a), 943 erroneous haplotypes appeared in a sample known to have only 15 before any processing. After a denoising process 15 haplotypes remained, but of these 6 (40%) were still sequences not present in the original sample, while 6 of the 15 original variants were discarded during the process. Clearly, separating wheat from chaff is a challenging problem.

Herein, we suggest an operational approach based on the stabilization of the entropy ratio to guide the cleaning procedures. Both the simulation approach and the analysis of the real dataset pointed to an α-value of 5 in the denoising step, which was also the optimal value selected in Elbrecht et al (2018a). Whether this value can be taken as a general rule of thumb or not will require analyses of more datasets. For the filtering step, our method indicated 20 reads as the optimal threshold.

Some authors proposed that denoising should be performed before clustering to identify genuine sequence variants, using different procedures, such as the UNOISE2 algorithm that we have adopted here (Edgar 2016), the MED (minimum entropy decomposition, Eren et al 2015) procedure, or the DADA2 algorithm (divisive amplicon denoising algorithm, Callahan et al 2016). It has also been suggested that sequence variants should replace MOTUs to capture relevant biological variation (Edgar 2016, Callahan et al 2017). This suggestion may be adequate in prokaryotes, where strains of the same species can have different characteristics (f.i., pathogenicity). However, for eukaryotes, and particularly metazoans, given the high amount of intraspecies information contained in the datasets, we think that it is more advisable to define meaningful MOTUs and perform denoising procedures within them, in order to obtain a “clean” dataset and be able to use the intra-MOTU sequence variability to make phylogeographic and population genetics inference. Clearly, our procedure is applicable only to coding sequences, which excludes much work done on protists based on ribosomal DNA. However, the growing number of metabarcoding studies using COI sequence data, together with the steady development of the BOLD database, makes us confident that many metabarcoding datasets of enormous potential for metaphylogeographic inference will become available in the near future.

We found a couple of instances of previous studies that have analysed COI structure in species recovered in our MOTU dataset and in nearby localities. For *Paracentrotus lividus* there were phylogeographic studies of the atlanto-mediterranean area using COI (Duran et al 2004), 16S (Calderón et al 2008), and the nuclear ANT intron (Calderón et al 2008). In all cases a low, but significant, signal corresponding to the separation between Atlantic and Mediterranean was found. Our COI results were in agreement with those of Duran et al (2004) for the localities that could be compared. We detected a somewhat higher number of haplotypes (11 in the previous work, 15 in our study) and the most common haplotypes were shared. The shape of the network was also similar. We want to emphasize that, as far as we could detect, not a single sea urchin of this species was present in our samples, so we obtained a similar level of haplotype diversity with community DNA than in a study specifically devoted to collect sea urchin specimens. For *Ophiothrix fragilis*, we also found a higher haplotype diversity (37 haplotypes) than in comparable localities in the work of Pérez-Portela et al (2013) (29 haplotypes). We identified five haplotypes that were shared in the two studies, including the commonest one in both datasets, and the networks again had similar structure. Of note here is that we could expand the distribution range of some of the haplotypes. Our AMOVA results for these two instances were equivalent to previous results for the only component that was analysed in both studies (the between seas differentiation). Thus, Duran et al (2004) found a significant (p<0.05) between-basin differentiation in *Paracentrotus lividus*, while Pérez-Portela et al (2013) didn’t find any significant genetic variability between Atlantic and Mediterranean for Lineage II of *Ophiothrix fragilis* (p=0.790). This is consistent with our metabarcoding-derived AMOVAs (p<0.001 and p=0.729, respectively).

We have used an already collected dataset, which can mimic the situation that many *a posteriori* studies can encounter. However, future metabarcoding studies can be planned taking into consideration the potential application for intraspecies analyses as well. For instance, PCR replicates for each sample can be of tremendous advantage to eliminate noise in the first steps. Increasing ecological replication can also be of great value for metaphylogeographic studies. We strongly advocate that published metabarcoding studies include in their datasets the information about which sequences are grouped into each MOTU with their sample distribution. This information is not commonly provided, and is necessary to make these studies amenable for intraspecies and metaphylogeographic analyses.

Metabarcoding now occupies a well-deserved prominent place among the methods for assessing community-level diversity (Kelly et al 2014, Adamowicz et al 2019). We have shown that it can be also an important source for species-level genetic diversity information. The mining of metabarcoding data for intraspecies information opens up a vast field with both basic and applied implications (Adams et al 2019). Among the latter, the possibility of effectively basing conservation efforts on multispecies and community-wide parameters (Nielsen et al 2017). It will also open the biogeography field, nowadays restricted almost exclusively to macro-organisms, to the myriad of meio- and micro-eukaryotes that make up most of the diversity present in natural communities.

Another related field is the assessment of connectivity between populations. This is important for endangered species, invasive species, protected areas design and management in general. For instance, in the marine environment, differences in larval dispersal have often been suggested as responsible for determining population genetic structure, but other factors such as variation in divergence times and changes in effective population sizes must be taken into account (Hart & Marko 2010). A powerful test for these contrasting assumptions is to compare phylogeographic patterns among species that concur or differ in larval type. Metaphylogeography can provide such comparative data. For instance, we have shown that metazoans in general have more between-replicate variability than other groups, and within metazoans the between community and between-replicate components of genetic variation can be significantly different between phyla.

In conclusion, our study shows the feasibility of mining metabarcoding datasets for the analysis of intraspecies genetic diversity using objective parameters for denoising and filtering spurious sequences. We cannot at present advice a set pipeline to do this, as procedures should be customized for the particulars (e.g., replication level, number of habitats, number of localities) of each study dataset. With this article, we hope to stir further discussion and developments in this field. The metaphylogeography application should be borne in mind to guide the planning and reporting of metabarcoding studies to ease the recovery of this - so far unexplored - vast amount of information.

## Acknowledgements

We are indebted to the staff of the Atlantic Islands of Galicia and Cabrera Archipelago National Parks for sampling permits and invaluable logistic help. Thanks also to Xavier Roijals for his skilful bioinformatic assistance in the CEAB computing cluster facilities. This work has been funded by projects Metabarpark (Spanish National Parks Autonomous Agency, OAPN 1036/2013) and PopCOmics ((CTM2017-88080, MCIU/AEI/FEDER/UE) of the Spanish Government.

