## Appendix S1 for "From metabarcoding to metaphylogeography: separating the wheat from the chaff"

Appendix S1. The dataset.

The dataset consisted of COI-based biodiversity data obtained from benthic marine communities at two Spanish National Parks, one in the Atlantic and one in the Mediterranean (Fig. S1). Several communities were sampled in 2014 and 2015 by completely scraping off standardized 25 x 25 cm quadrats. Three replicate samples were collected per community, and each sample was then separated through sieving into three size fractions (>10 mm, 1-10 mm, 63 µm-1mm, roughly corresponding to mega-, macro- and meiobenthos, Rex & Ettter, 2010).

Wangensteen et al (2018a) reported α- and β-diversity results of the sampling performed in 2014, including four communities in the Mediterranean Park (Cabrera Archipelago, Balearic Islands) and four in the Atlantic Park (Atlantic Islands of Galicia). These communities were, in each Park, two well-lit communities, one deeper, invertebrate-dominated, community, and a detritic bottom with coralline algae (Table S1). In 2015, the sampling was repeated on the same localities and communities, except for a new community sampled in Cabrera (*Caulerpa cylindracea* community) and the change of one of the two well-lit communities in the Atlantic (*Asparagopsis armata* community instead of *Cystoseira tamariscifolia* community, Table S1). Some of the communities sampled in 2015 were used in a study of the effect of invasive seaweeds (Wangensteen et al 2018b). A total of 51 samples separated in 153 fractions were included in the present study (Table S1).

Samples were extracted and sequenced using a modification of the Leray et al. (2013) primers for a 313 bp fragment of COI, with the adequate blanks and negatives, following procedures detailed in Wangensteen et al (2018a). Separate libraries were built with samples from 2014 and 2015 and sequenced in two runs on an Illumina MiSeq platform (2 x 300 bp paired-end) at Fasteris SA (Plan-les-Ouates, Switzerland).

For the present study, we pooled the reads of the two years and reanalysed the joint dataset with the same pipeline as in Wangensteen et al (2018a), based mostly on the OBITools suite (Boyer et al. 2016), and including quality and chimera controls. We nevertheless made three changes in the published pipeline: first, we applied a strict length filter keeping only sequences of the expected length (313 bp). Second, instead of applying the CROP algorithm for clustering (Hao et al. 2011), we used the SWARM2 method (Mahé et al. 2015), with a d-parameter of 13. Previous comparisons on restricted datasets showed that the two procedures yielded highly similar values of α-diversity, and SWARM was much faster than CROP. Third, we didn't perform the filtering by minimal relative abundance, whereby all MOTUs that did not have a relative abundance greater than 0.01% in at least one sample were removed. This was done because we wanted to keep all potentially informative sequence data for the curation procedure. At this step, we only discarded sequences with just one read in all the dataset, as is common practice in metabarcoding studies. We also pooled the sequences of the three fractions of each sample for downstream analyses. Following the pipeline, we generated a MOTU list and assigned a taxonomical rank to each MOTU. Non-eukaryotic MOTUs were removed.

Table S1. Sample characteristics, with indication of locality, type of community, dominant species, depth, coordinates, and number of replicate samples collected in each study year

| National Park | Community | Dominant species | Depth (m) | Coordinates | samples 2014 | samples 2015 |
|---|---|---|---|---|---|---|
| Cabrera Archipelago | Photophilic algae | *Lophocladia lallemandii* | 7-10 | 39.1250,2.9603 | 3 | 3 |
| Cabrera Archipelago | Photophilic algae | *Padina pavonica* | 7-10 | 39.1250,2.9603 | 3 | 3 |
| Cabrera Archipelago | Sciaphilic algae | Sponges and invertebrates | 30 | 39.1250,2.9603 | 3 | 3 |
| Cabrera Archipelago | Sciaphilic algae | *Caulerpa cylindracea* | 30 | 39.1250,2.9603 | -- | 3 |
| Cabrera Archipelago | Detritic bottoms | Coralline algae | 50 | 39.1249,2.9604 | 3 | 3 |
| Atlantic Islands | Photophilic algae | *Cystoseira nodicaulis* | 3-5 | 42.2259,-8.8969 | 3 | 3 |
| Atlantic Islands | Photophilic algae | *Cystoseira tamariscifolia* | 3-5 | 42.2260,-8.8970 | 3 | -- |
| Atlantic Islands | Photophilic algae | *Asparagopsis armata* | 4-6 | 42,2146,-8.8973 | -- | 3 |
| Atlantic Islands | Sciaphilic algae | *Saccorhiza polyschides* | 16 | 42.1917,-8.8885 | 3 | 3 |
| Atlantic Islands | Detritic bottoms | Coralline algae | 20 | 42.2123,-8.8972 | 3 | 3 |

Figure legends

Figure S1. Map of the Iberian Peninsula showing the two sampling sites in the Cabrera Archipelago (Mediterranean) and Cies Islands (Atlantic). The sampling zone in each site is indicated in yellow.

Figure S1

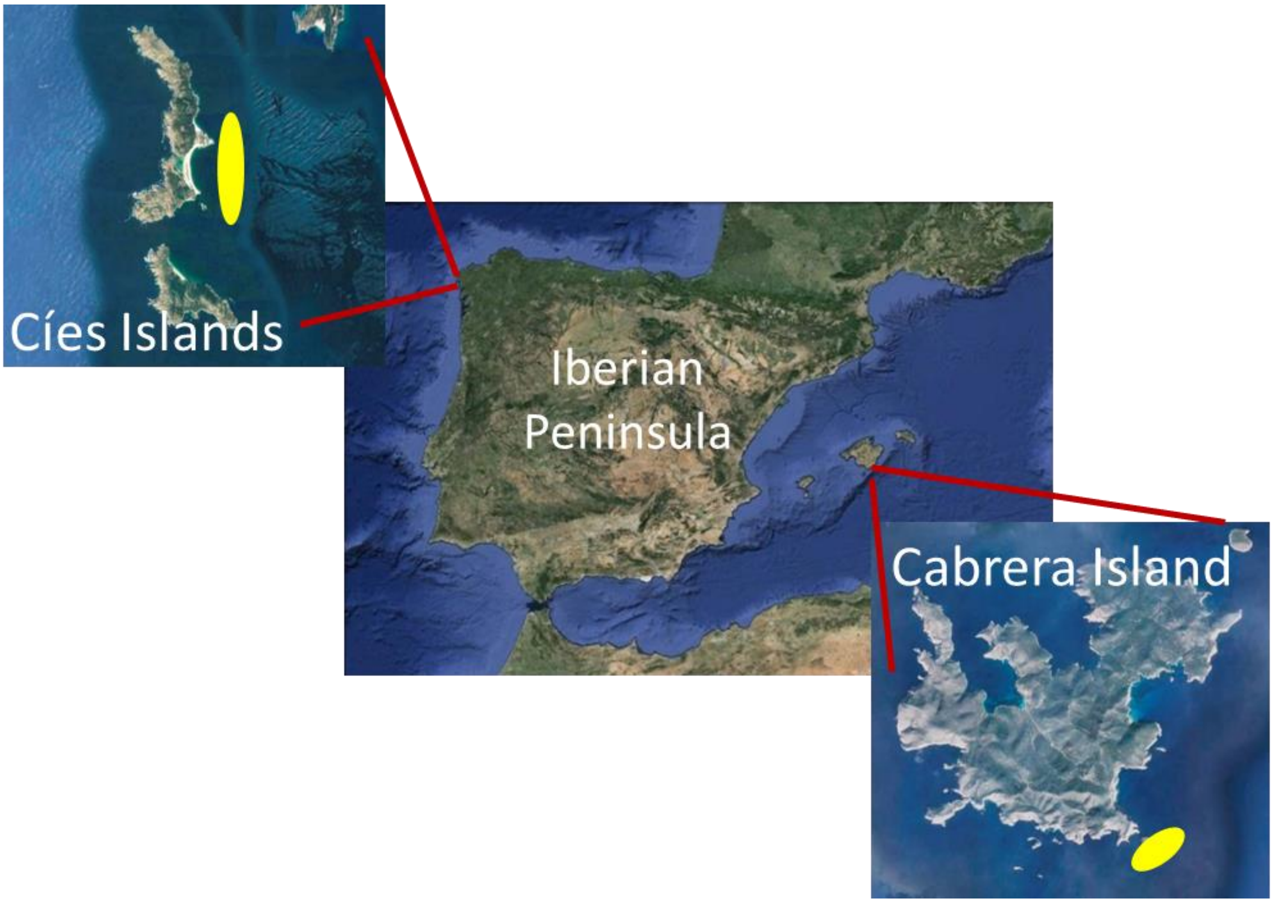
