## Appendix S2 for "From metabarcoding to metaphylogeography: separating the wheat from the chaff"

Network analyses of the MOTUs retained after denoising and filtering. The size of the pies is proportional to the semiquantitative rank abundances used. Blue color represent abundance in Mediterranean samples, red color in Atlantic samples. The code of the MOTU and the main group where they belong are indicated. For details on each MOTU, refer to Data S2. Only graphs for MOTUs with >2 and < 230 sequences are represented.

MBPA_000000099 Rhodophyta

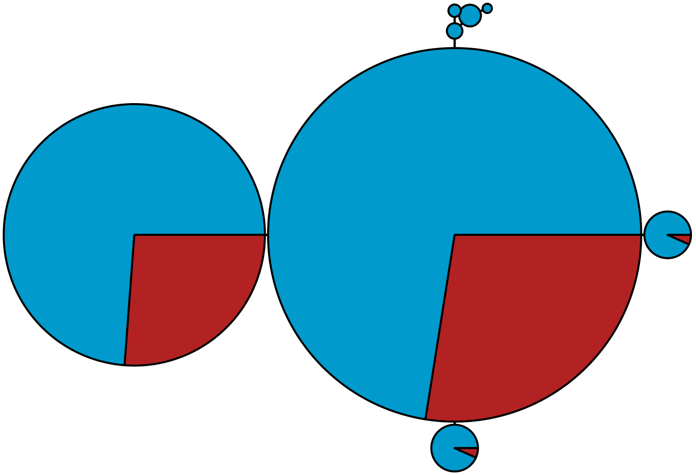

MBPA_000000127 Metazoa

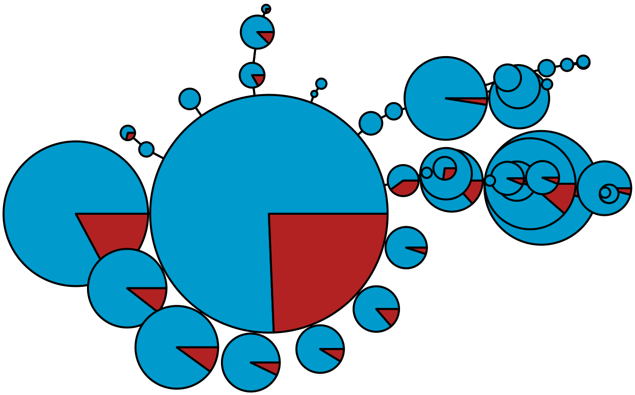

MBPA_000000141

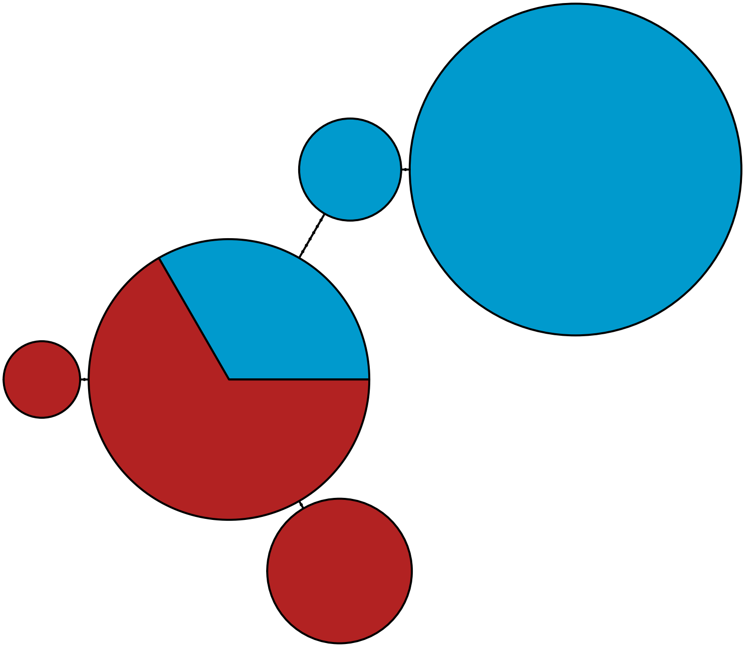

MBPA_000000143 Stramenopiles

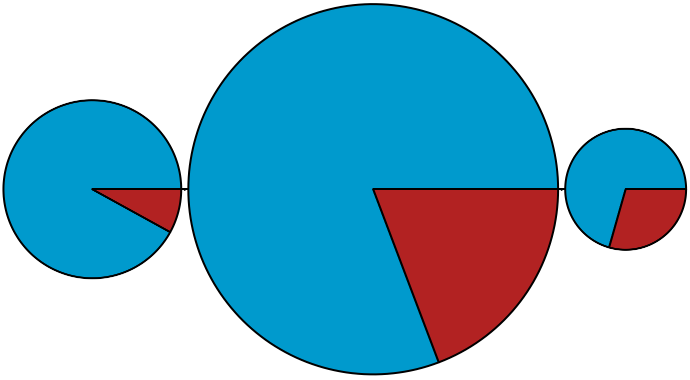

MBPA_000000159 Rhodophyta

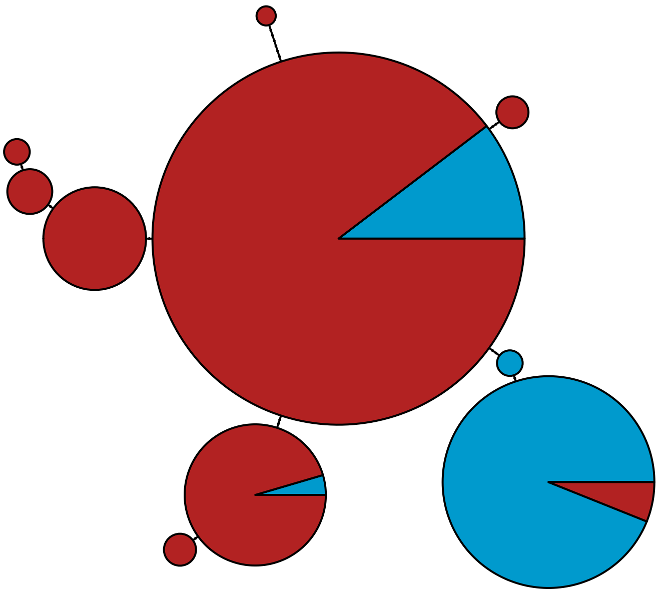

MBPA_000000178 Metazoa

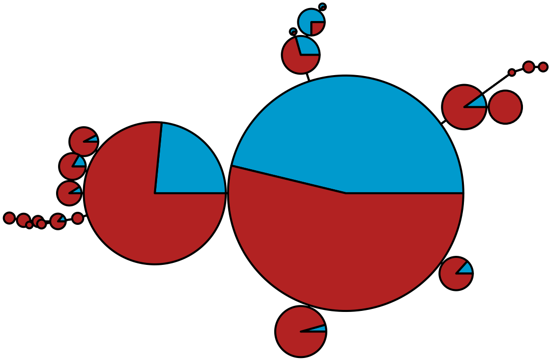

MBPA_000000180 Metazoa

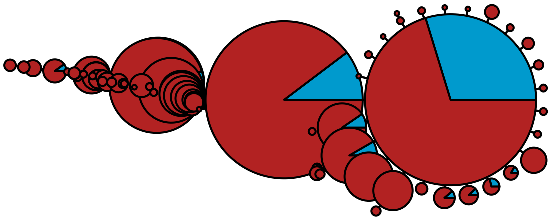

MBPA_000000189 Rhodophyta

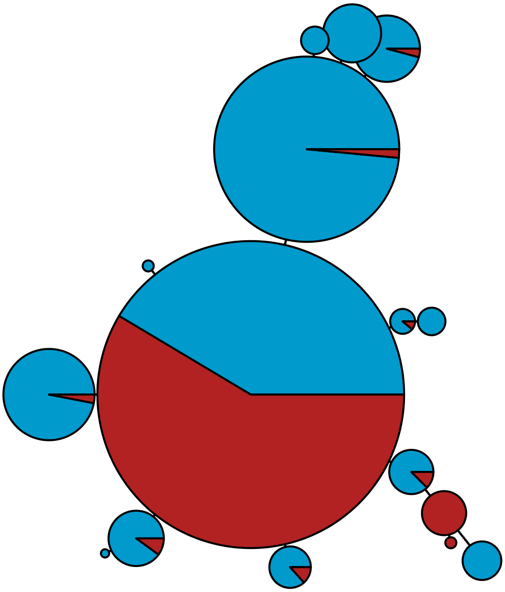

MBPA_000000191 Stramenopiles

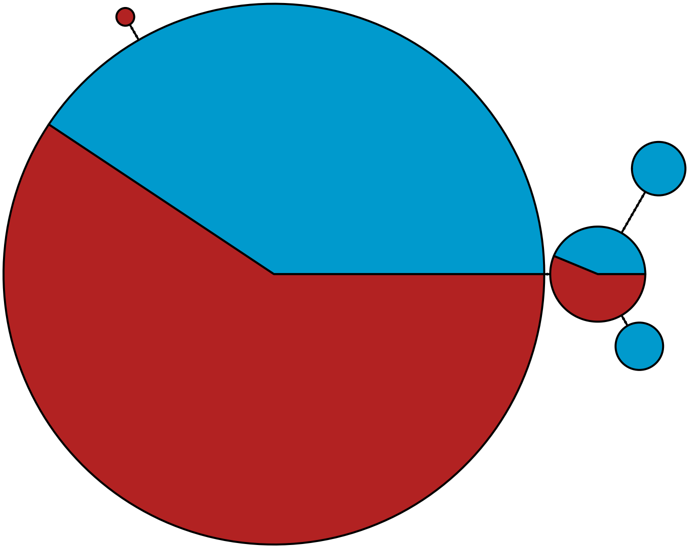

MBPA_000000206 Rhodophyta

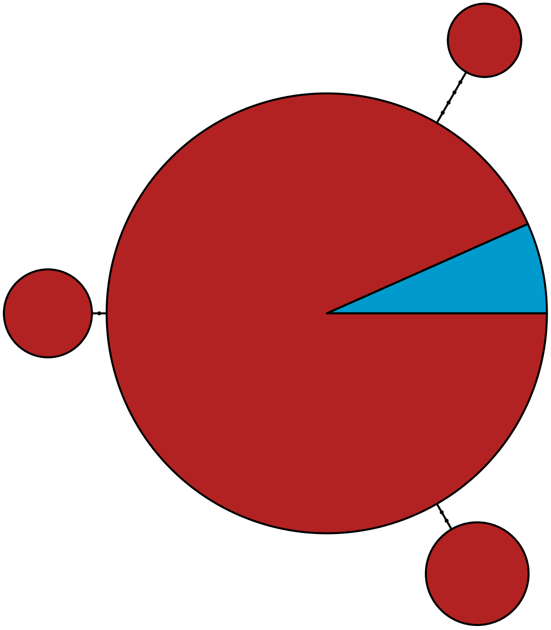

MBPA_000000229 Metazoa

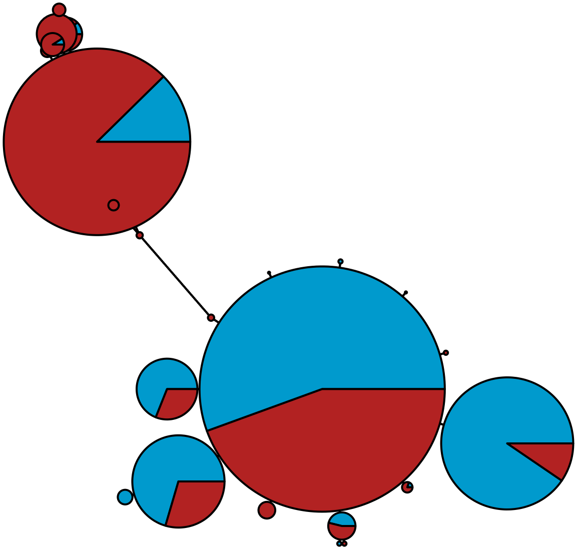

MBPA_000000240 Metazoa

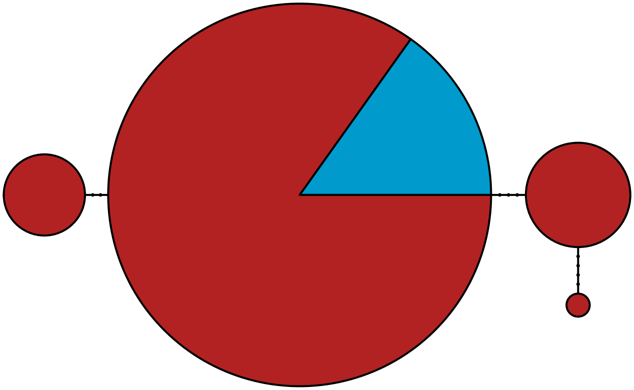

MBPA_000000245 Rhodophyta

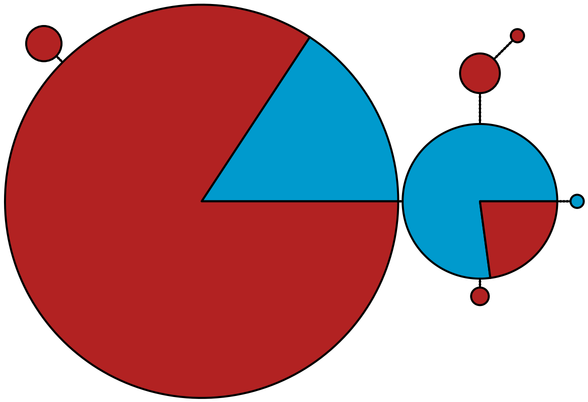

MBPA_000000247 Rhodophyta

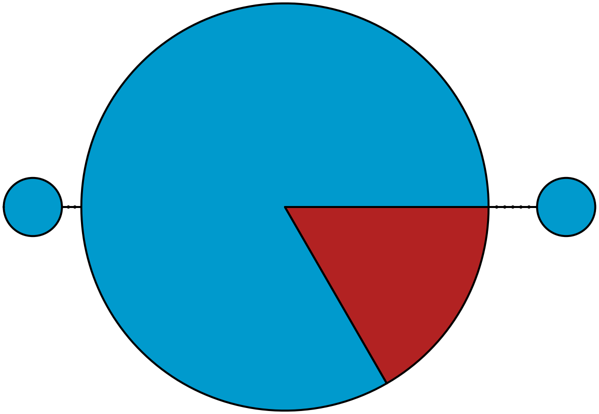

MBPA_000000254 Rhodophyta

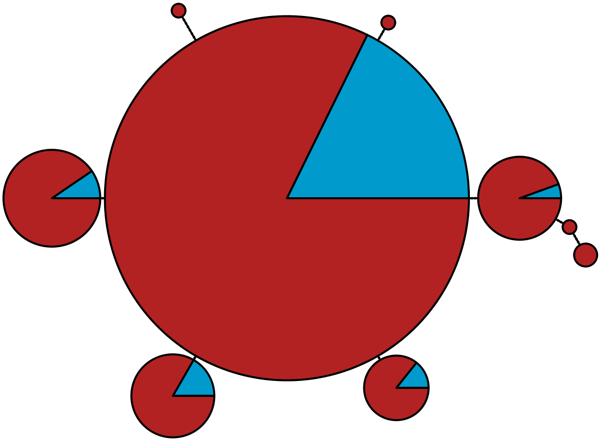

MBPA_000000269 Rhodophyta

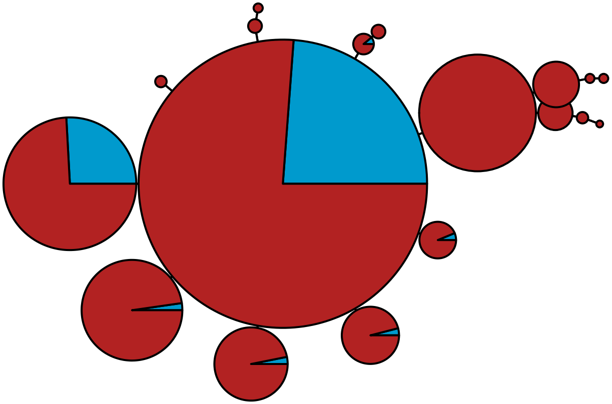

MBPA_000000279 Metazoa

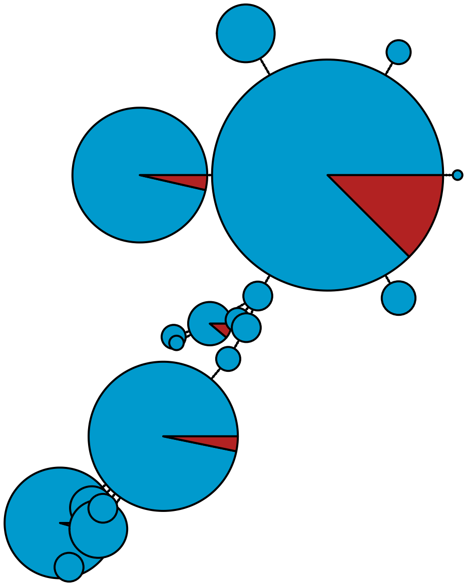

MBPA_000000284 Metazoa

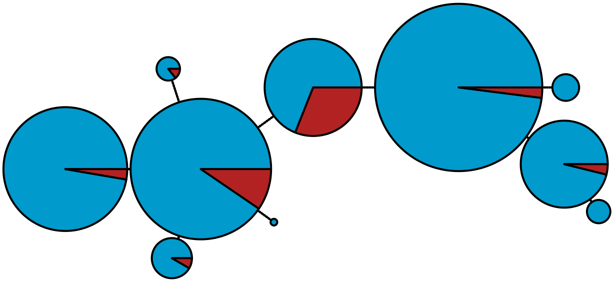

MBPA_000000300 Metazoa

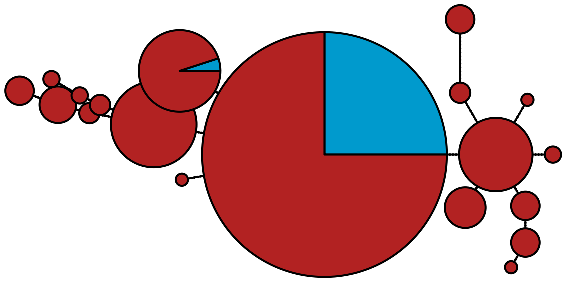

MBPA_000000301 Metazoa

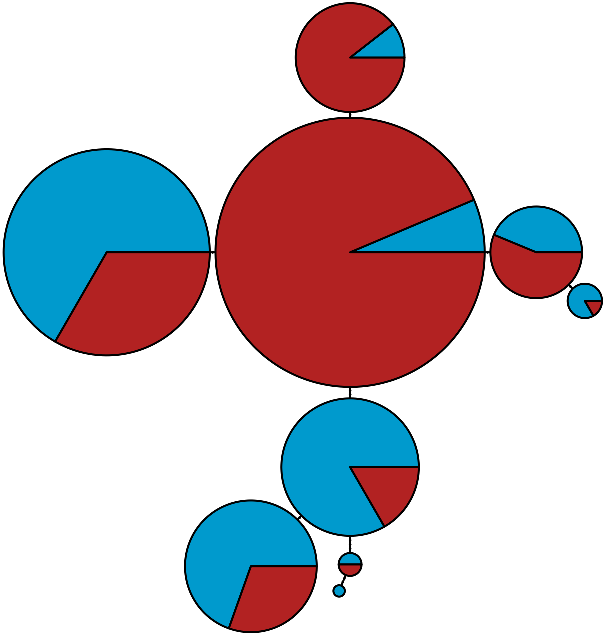

MBPA_000000310 Rhodophyta

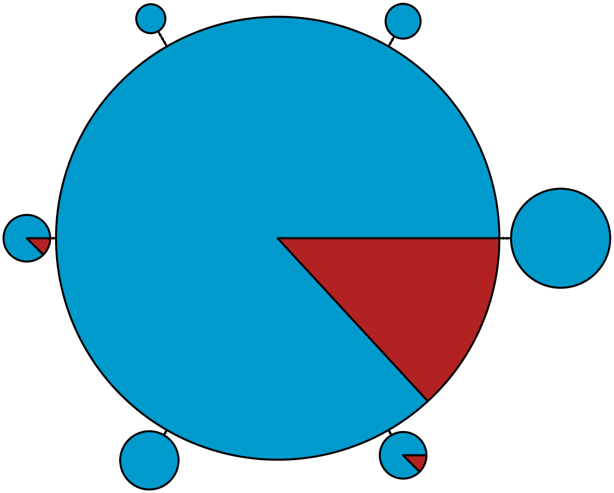

MBPA_000000314 Metazoa

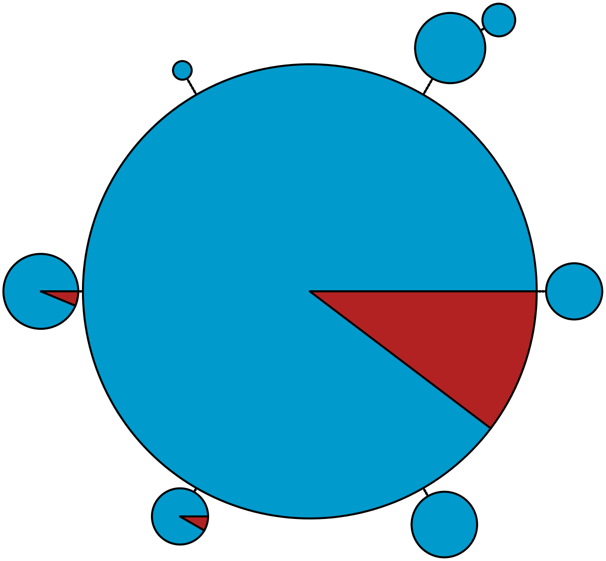

MBPA_000000350 Metazoa

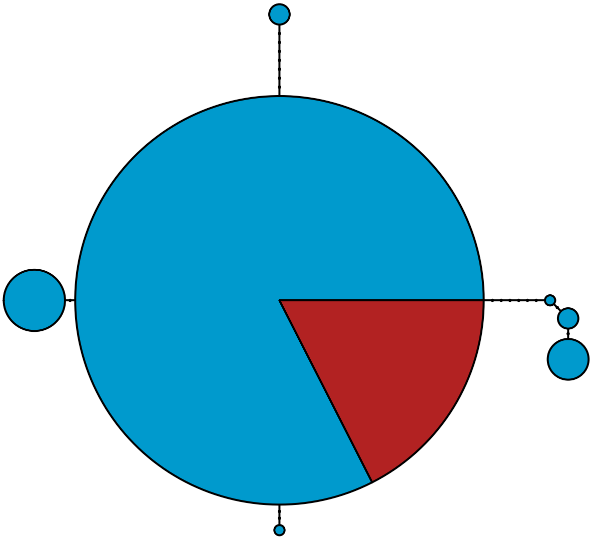

MBPA_000000355 Metazoa

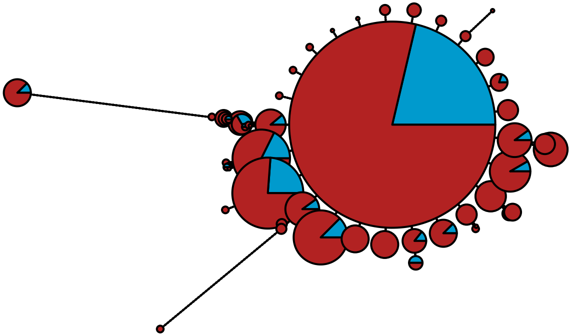

MBPA_000000362 Stramenopiles

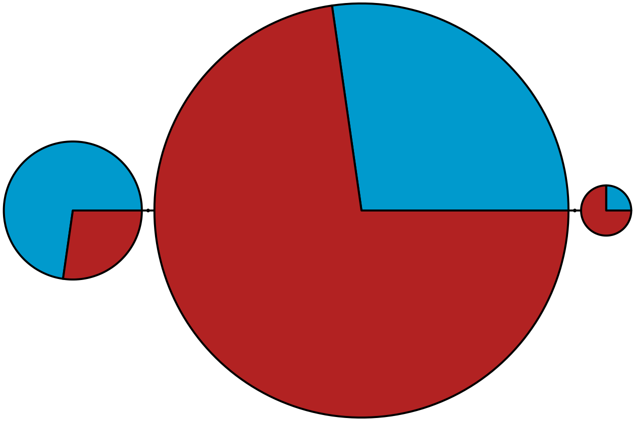

MBPA_000000364 Metazoa

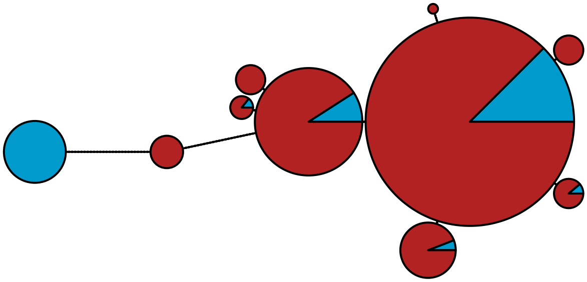

MBPA_000000386 Metazoa

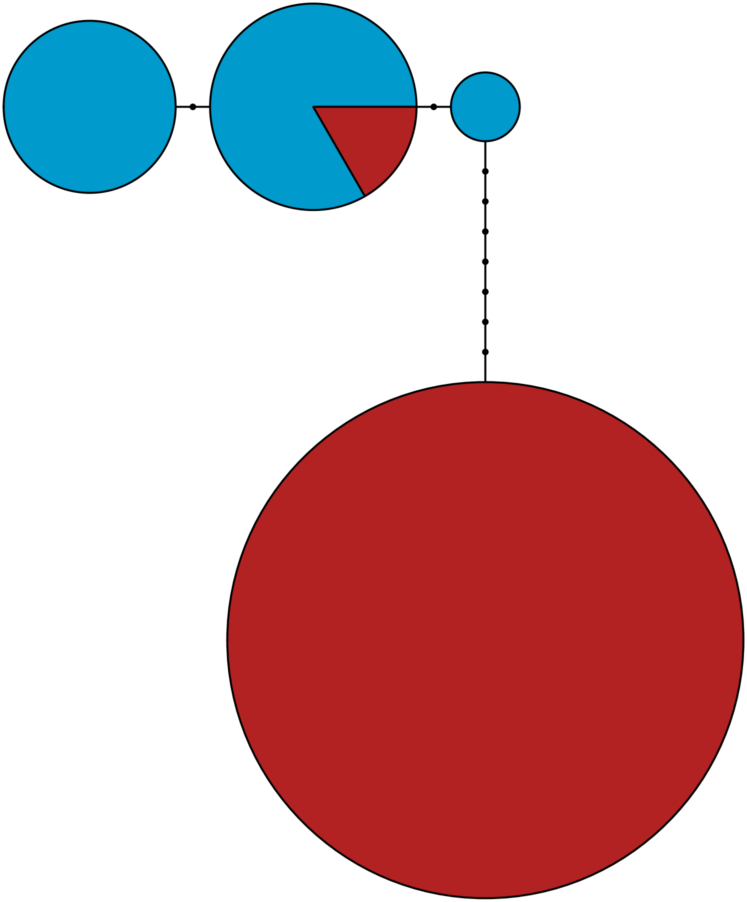

MBPA_000000398

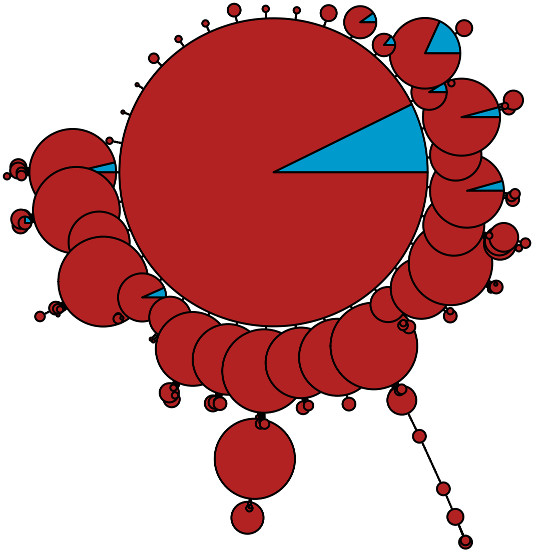

MBPA_000000400 Rhodophyta

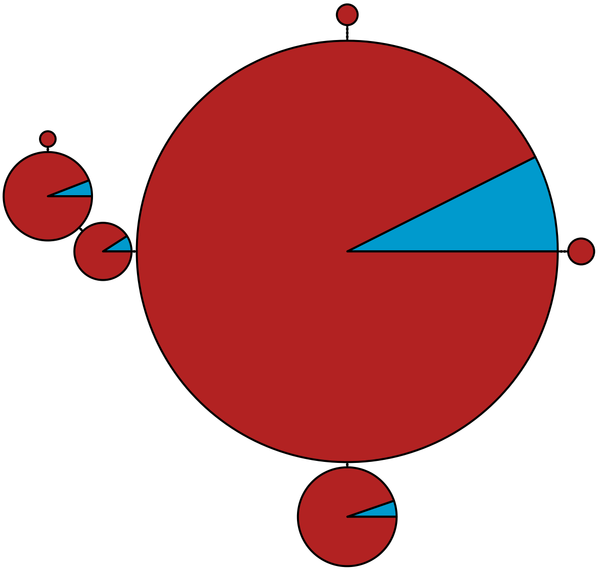

MBPA_000000450 Metazoa

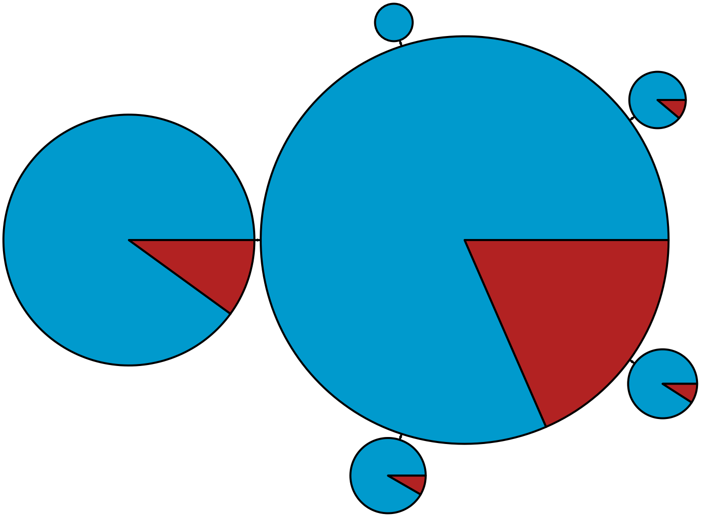

MBPA_000000479 Metazoa

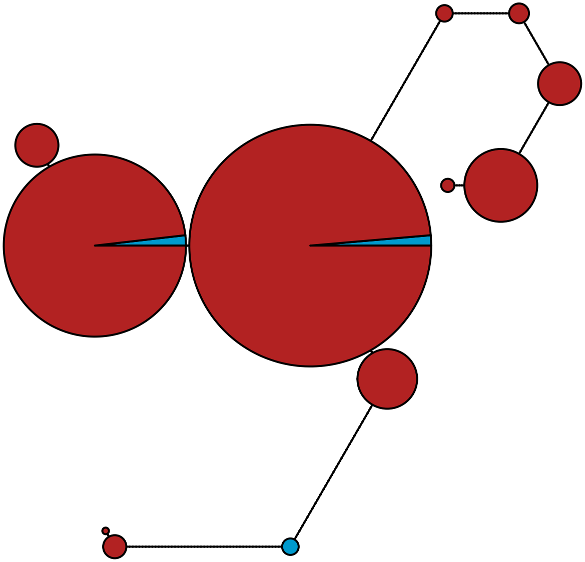

MBPA_000000507 Metazoa

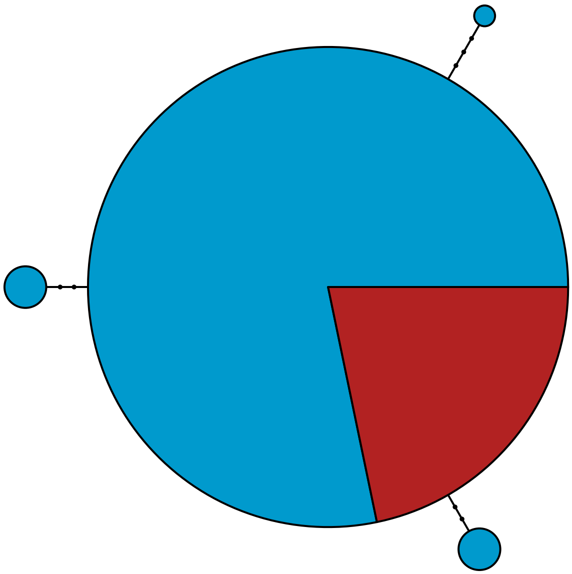

MBPA_000000542 Rhodophyta

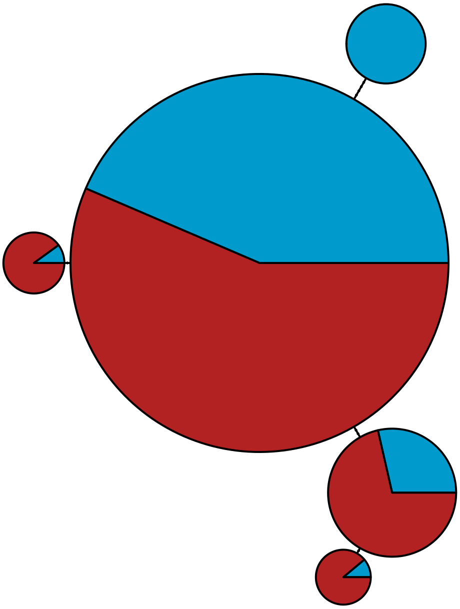

MBPA_000000550

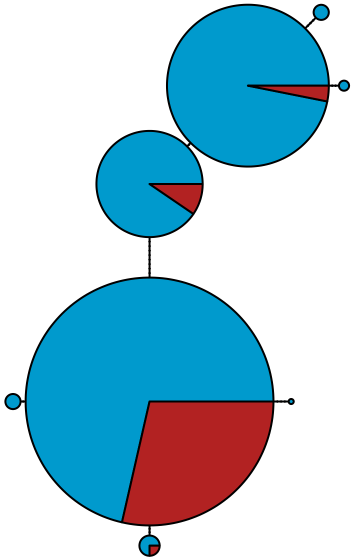

MBPA_000000581 Stramenopiles

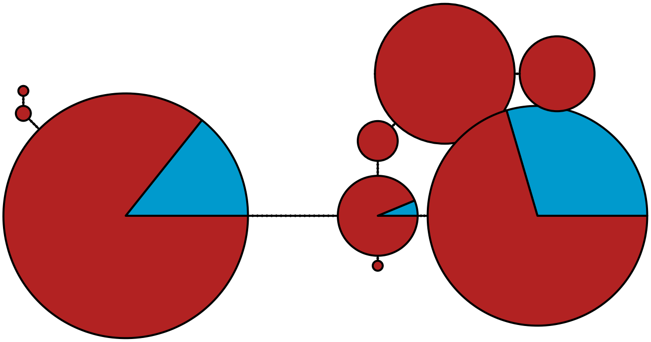

MBPA_000000587 Metazoa

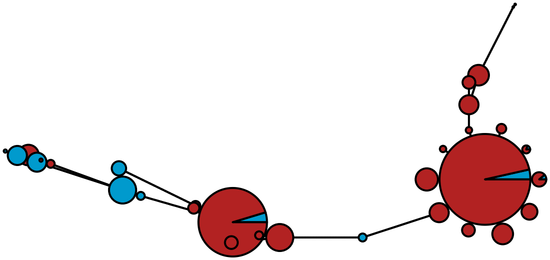

MBPA_000000595 Metazoa

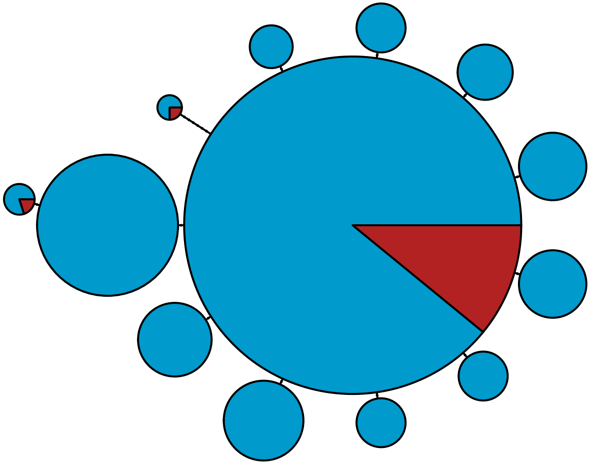

MBPA_000000636 Rhodophyta

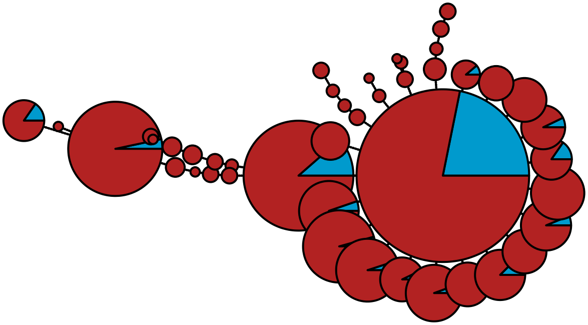

MBPA_000000697 Metazoa

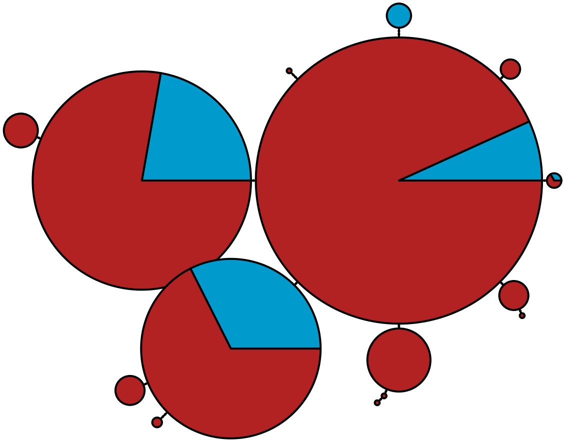

#### MBPA_000000715 Metazoa

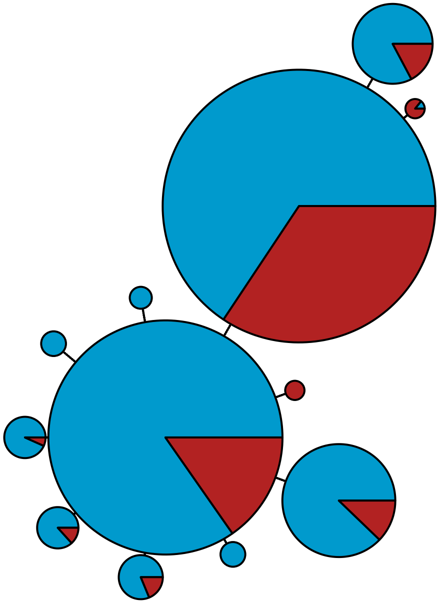

MBPA_000000729 Metazoa

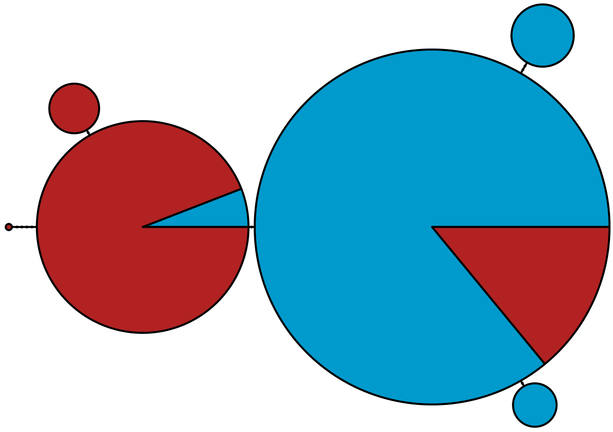

MBPA_000000740 Rhodophyta

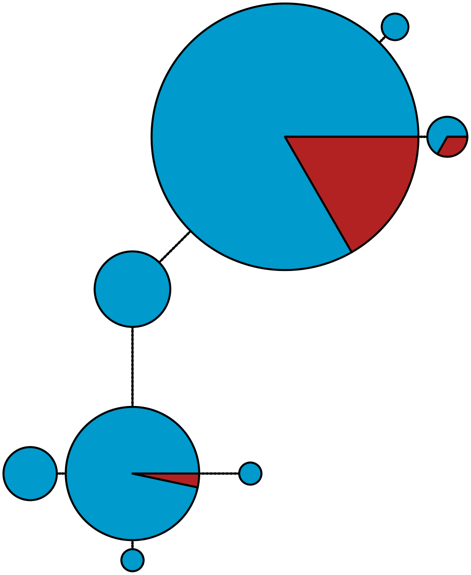

MBPA_000000750 Metazoa

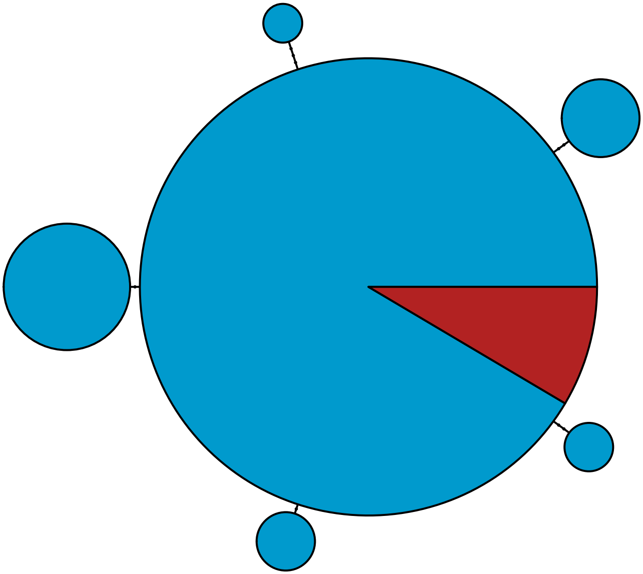

MBPA_000000752 Metazoa

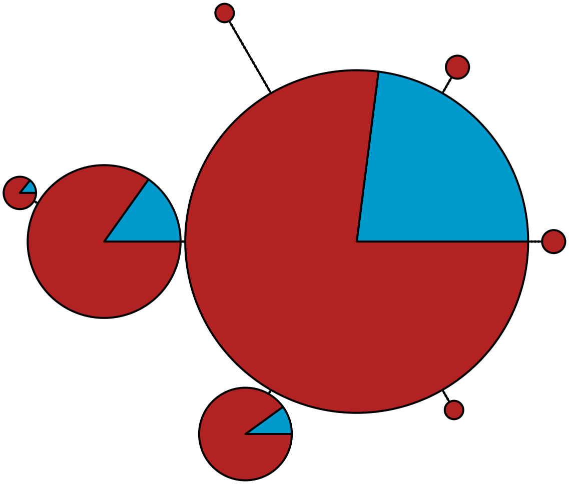

MBPA_000000754 Rhodophyta

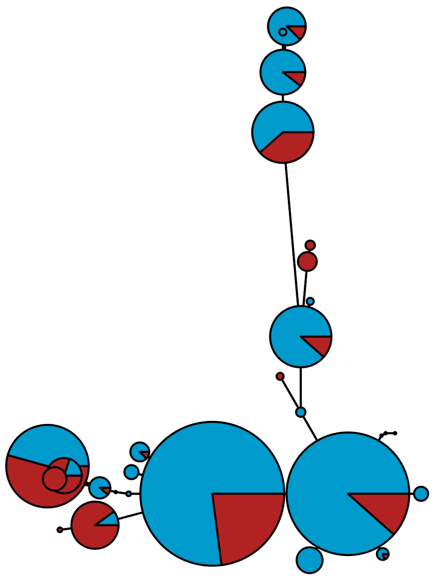

MBPA_000000758 Metazoa

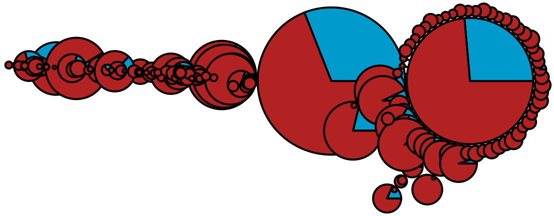

MBPA_000000762 Metazoa

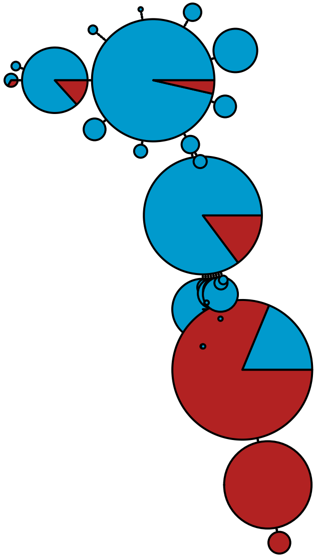

MBPA_000000774 Metazoa

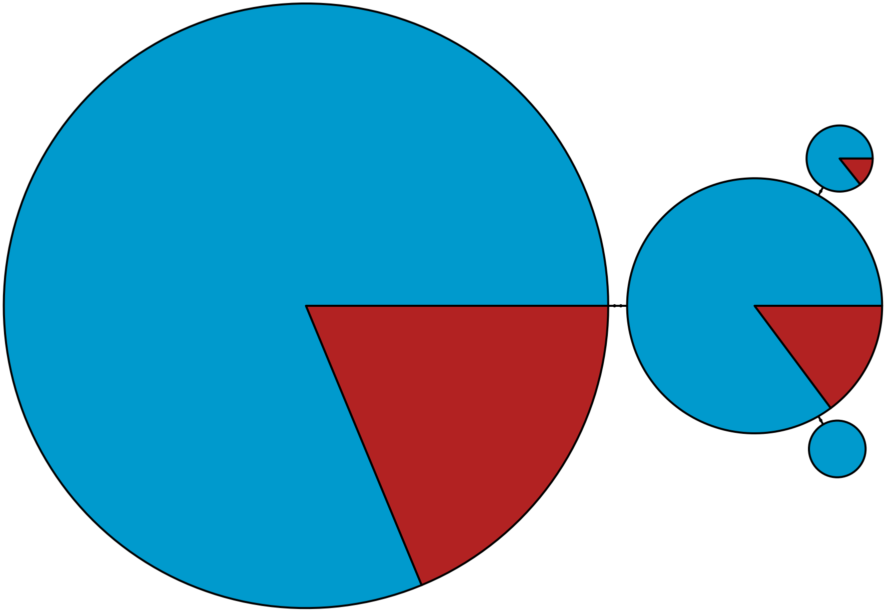

MBPA_000000790 Metazoa

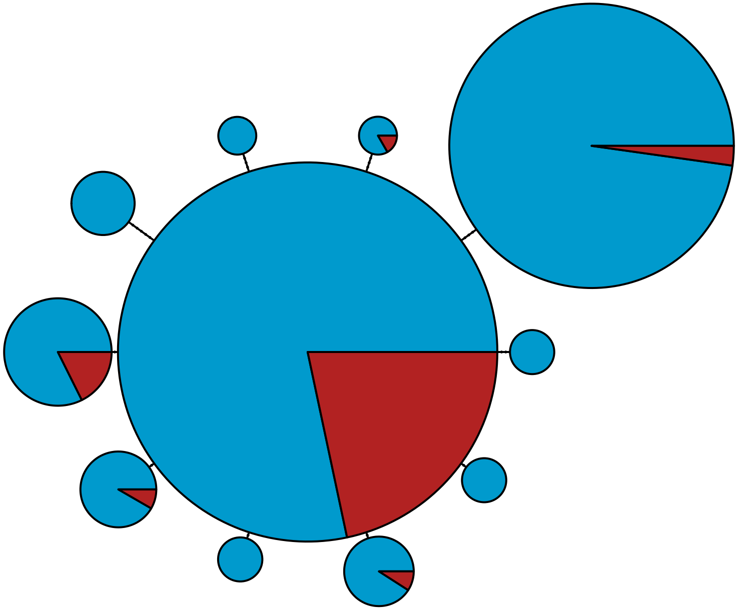

MBPA_000000816 Metazoa

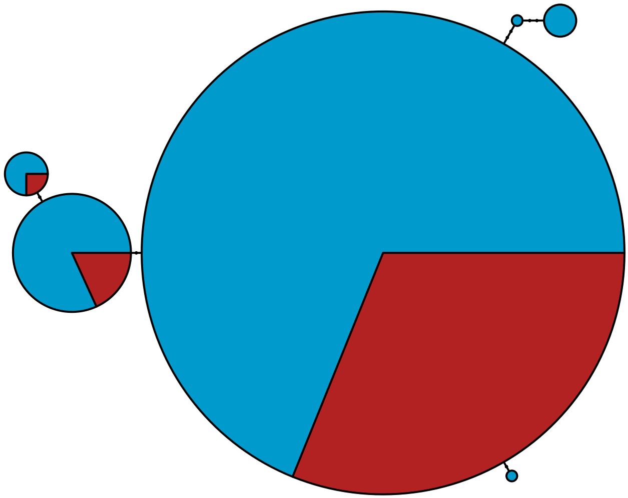

MBPA_000000846 Metazoa

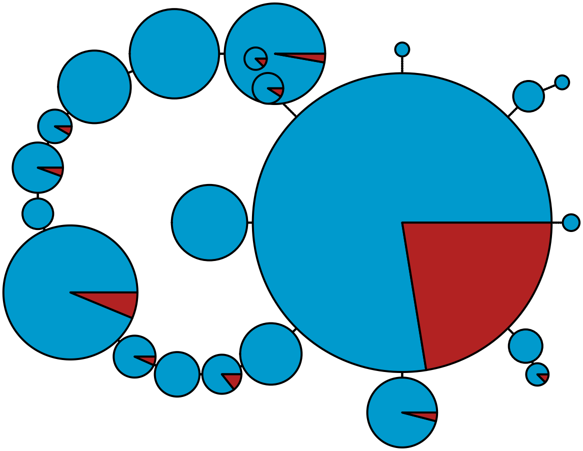

MBPA_000000863 Metazoa

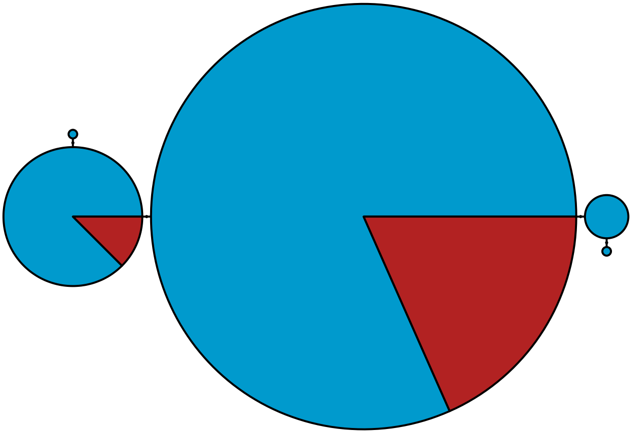

MBPA_000000887 Metazoa

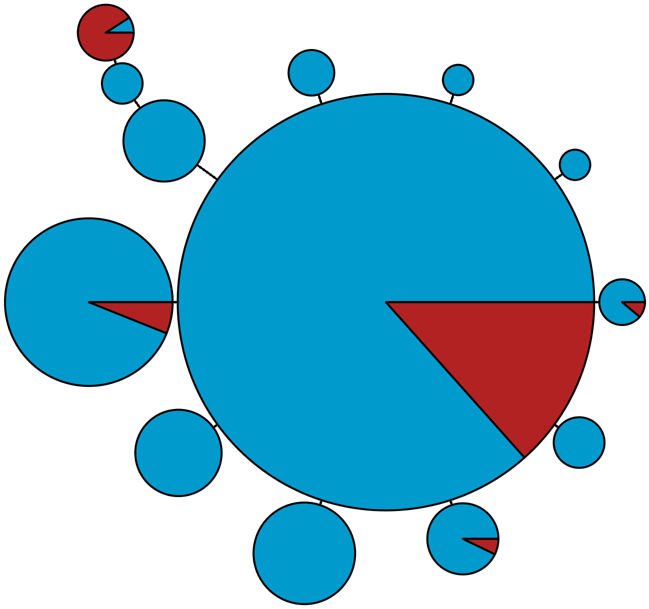

MBPA_000000935 Metazoa

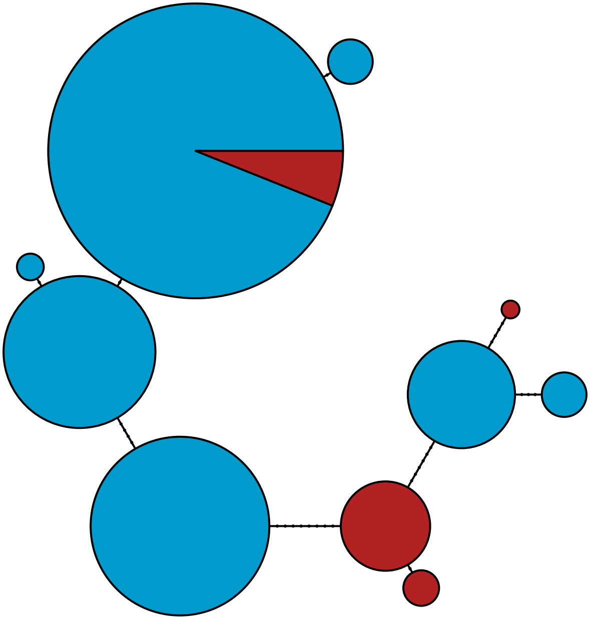

MBPA_000000965 Metazoa

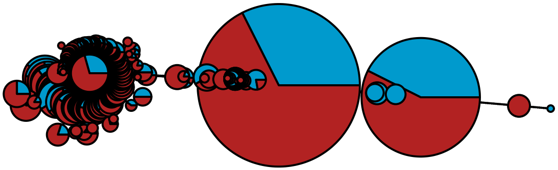

MBPA_000000975 Metazoa

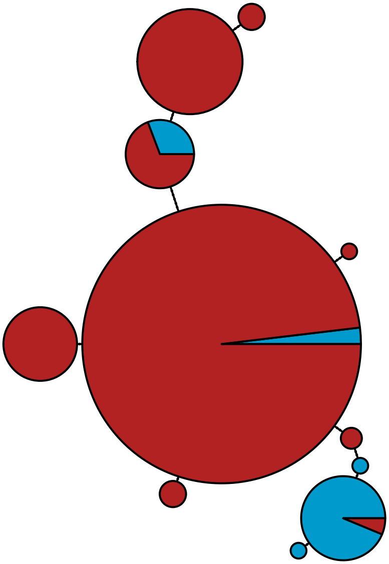

MBPA_000000976 Metazoa

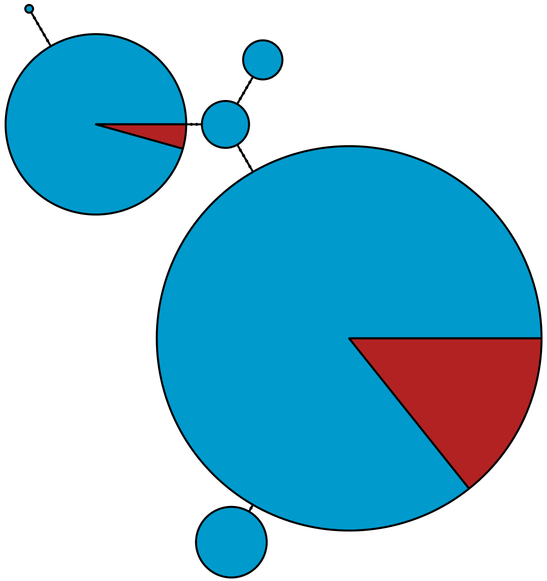

MBPA_000000995 Metazoa

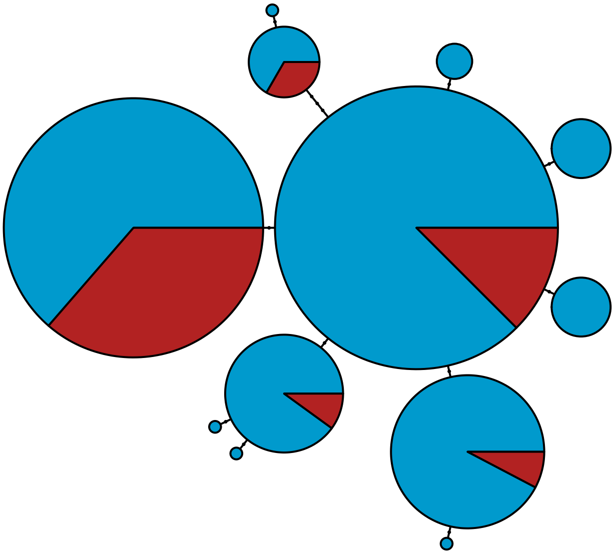

### MBPA_000001009 Metazoa

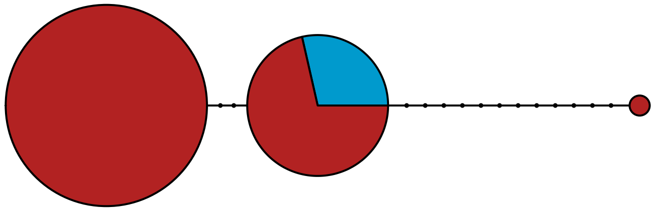

MBPA_000001011 Metazoa

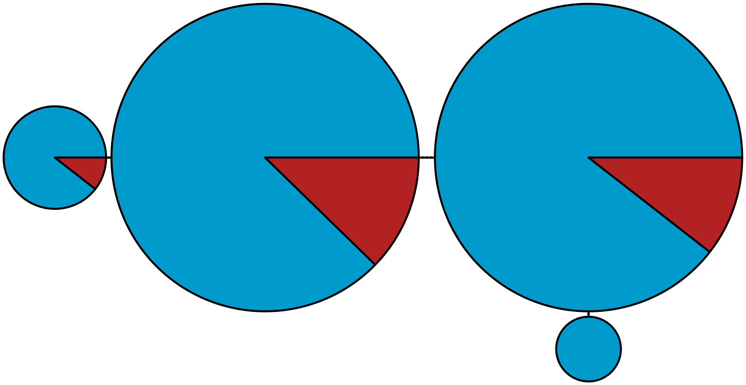

MBPA_000001014 Metazoa

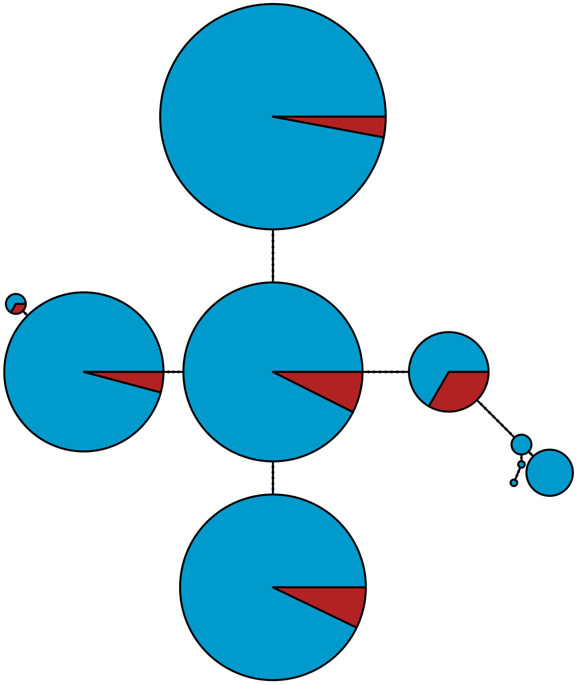

MBPA_000001015 Metazoa

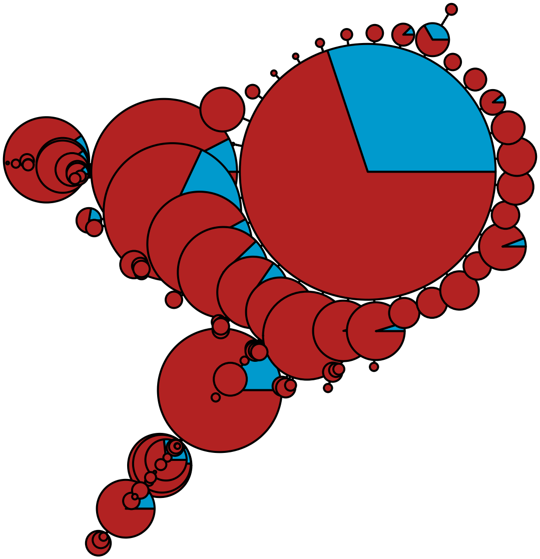

MBPA_000001019 Metazoa

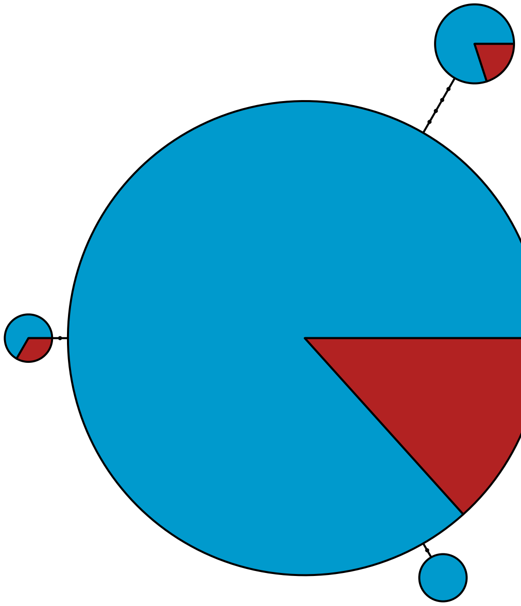

MBPA_000001038 Rhodophyta

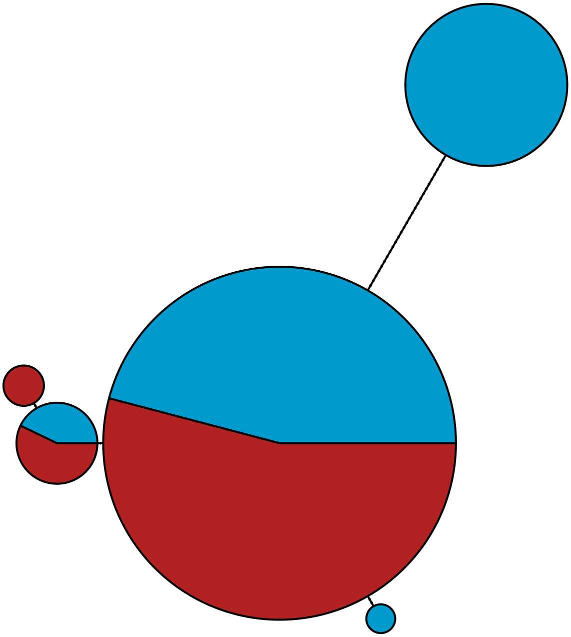

MBPA_000001075 Metazoa

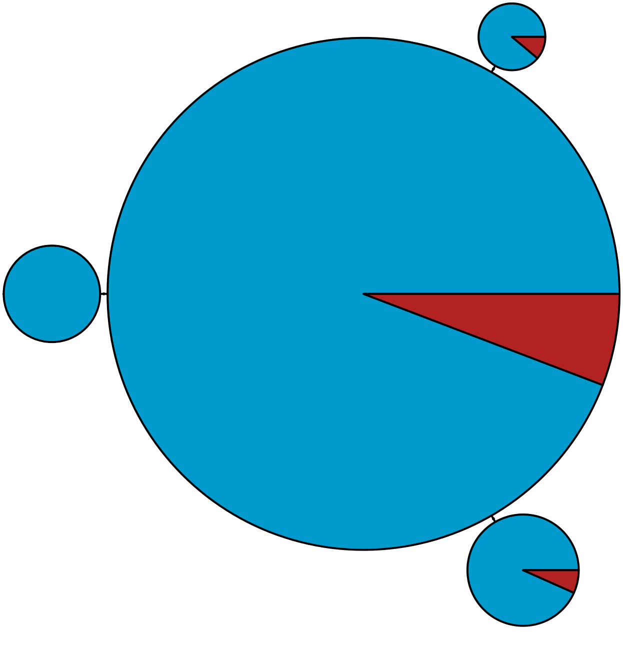

MBPA_000001081 Metazoa

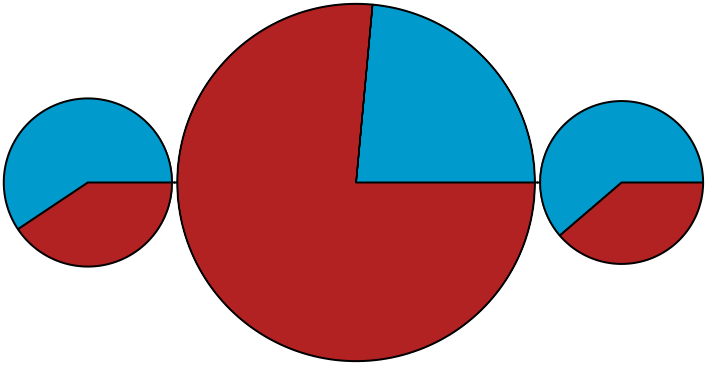

MBPA_000001116 Metazoa

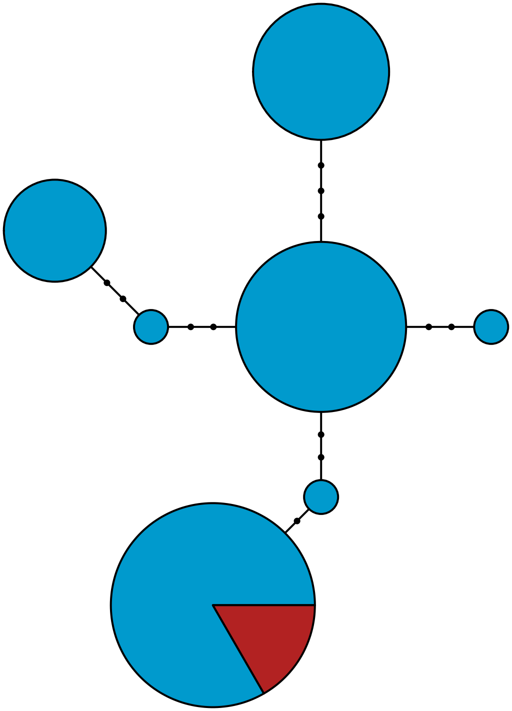

MBPA_000001158 Metazoa

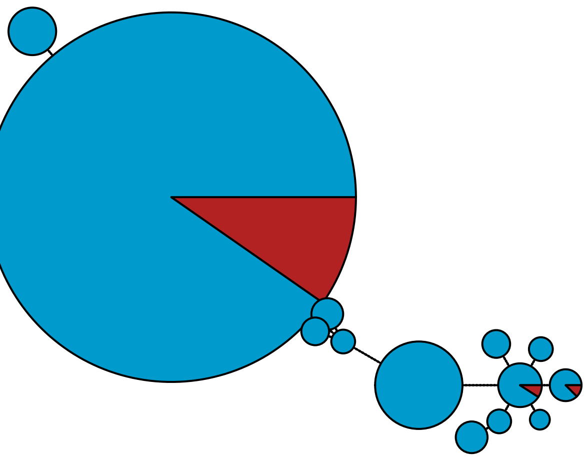

MBPA_000001172 Metazoa

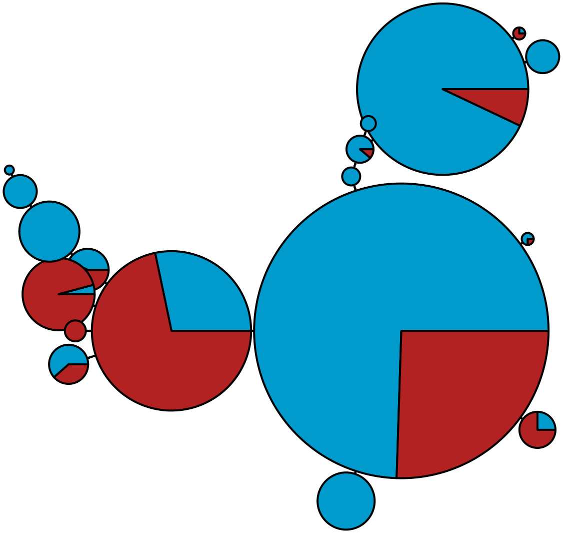

MBPA_000001190 Rhodophyta

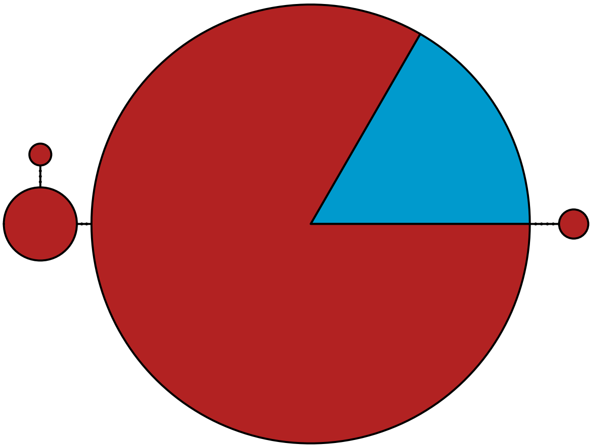

MBPA_000001194 Stramenopiles

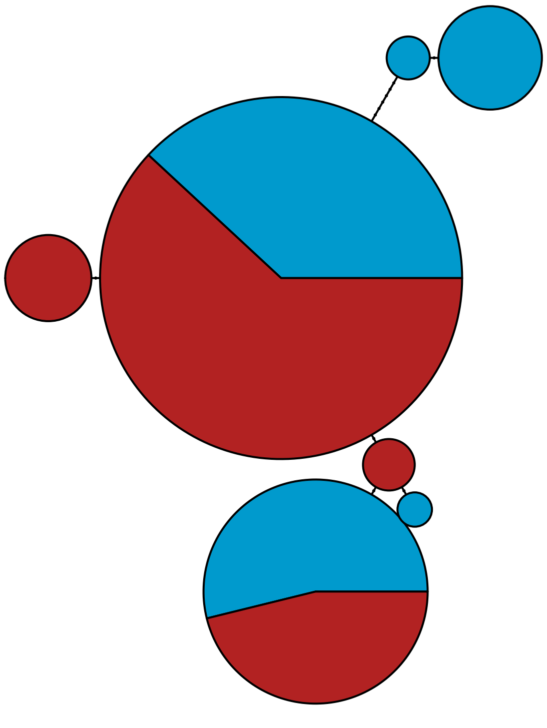

MBPA_000001196 Metazoa

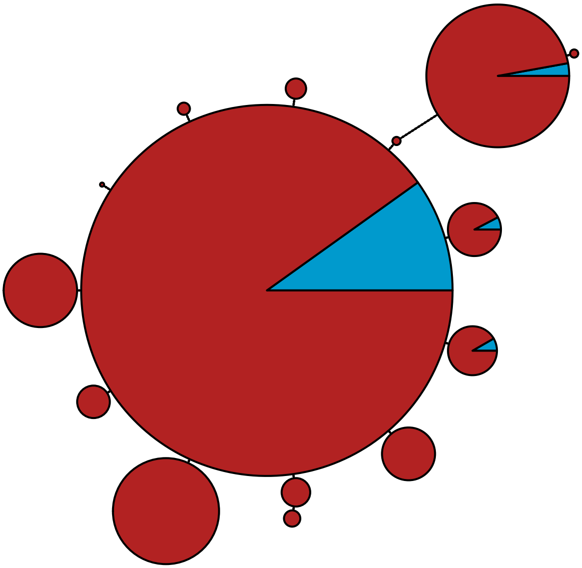

MBPA_000001205 Stramenopiles

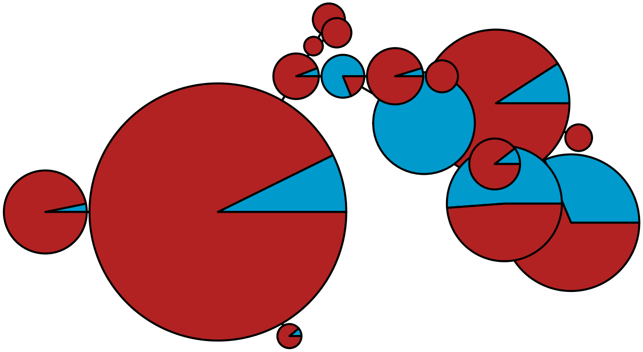

MBPA_000001214 Metazoa

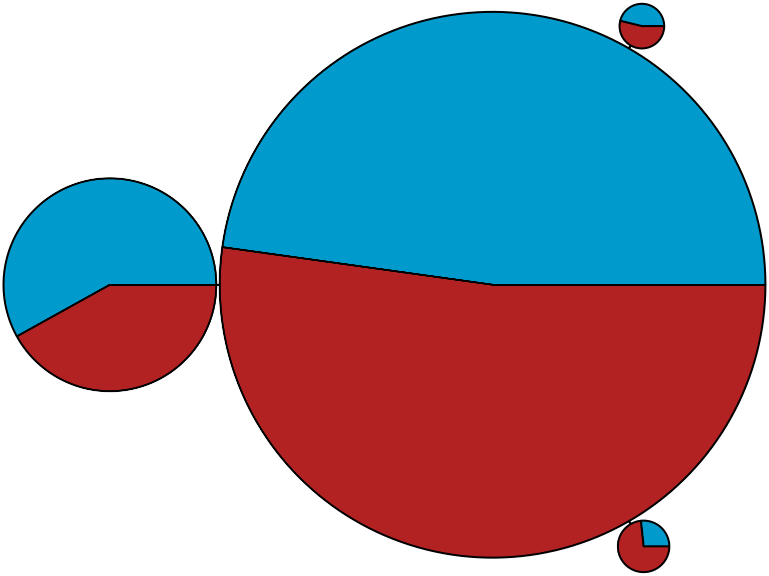

MBPA_000001240 Metazoa

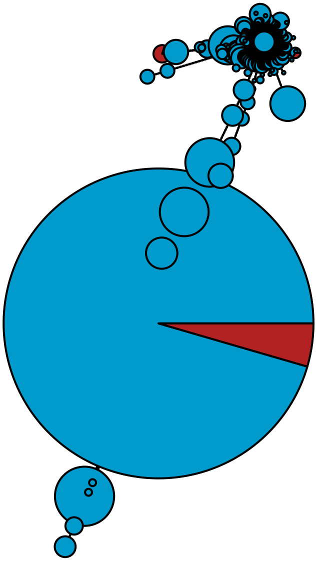

MBPA_000001378 Metazoa

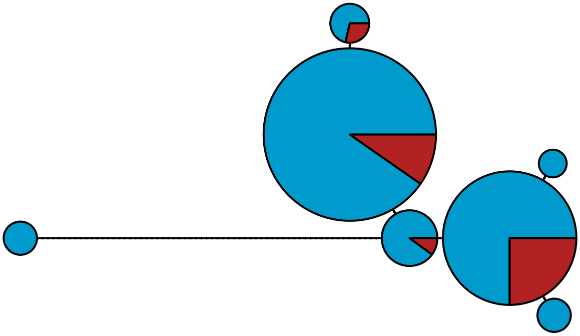

MBPA_000001396 Rhodophyta

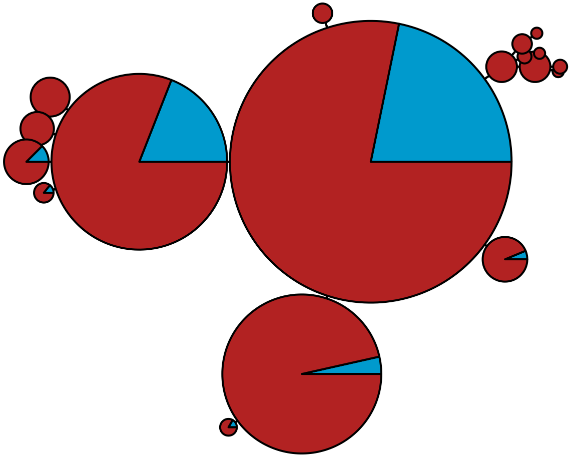

MBPA_000001417 Metazoa

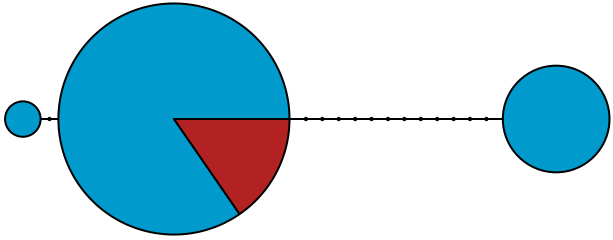

MBPA_000001432 Metazoa

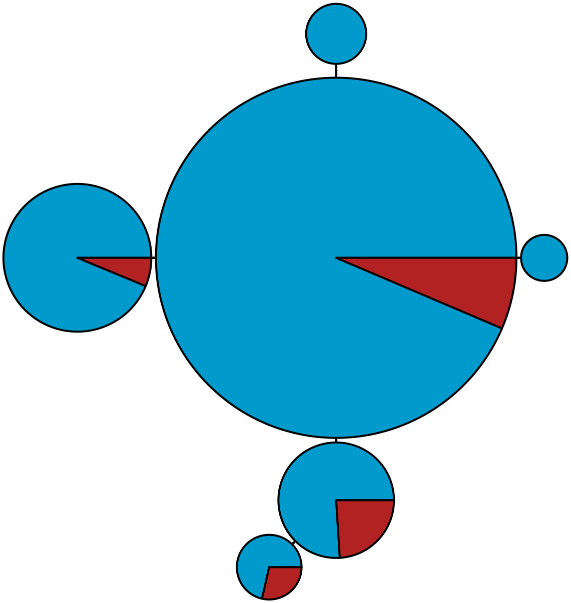

MBPA_000001444 Metazoa

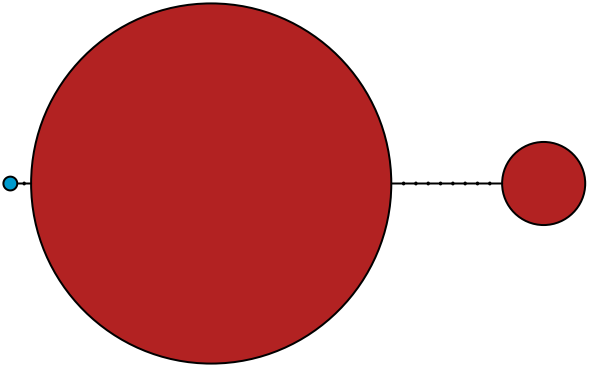

MBPA_000001446 Metazoa

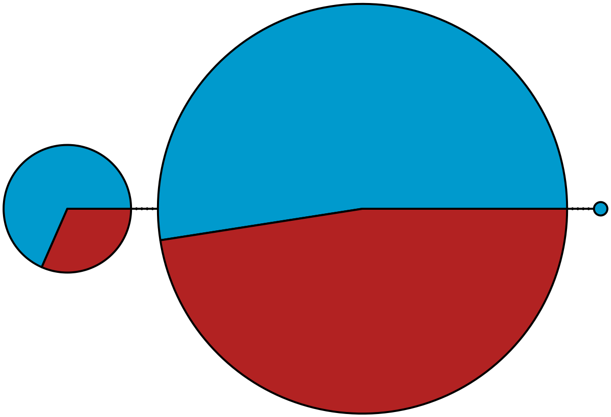

MBPA_000001457 Metazoa

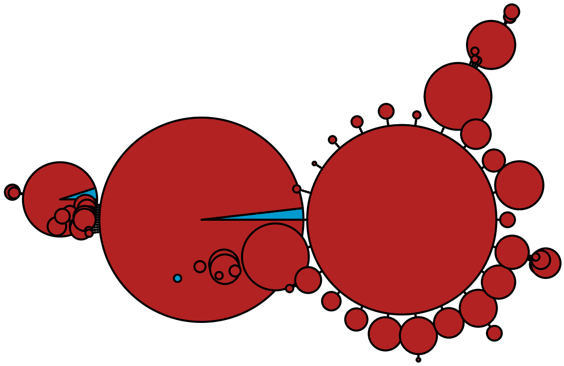

MBPA_000001473 Metazoa

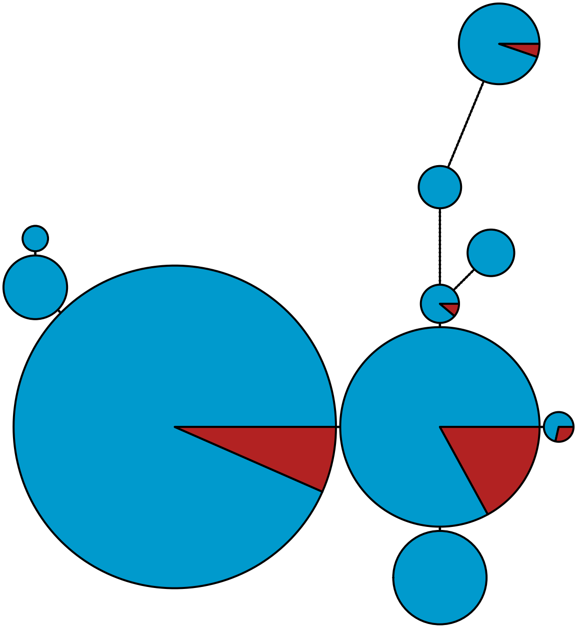

MBPA_000001520 Metazoa

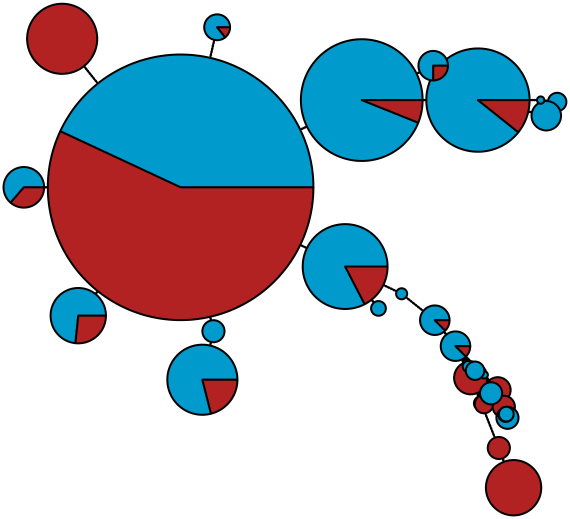

MBPA_000001531 Metazoa

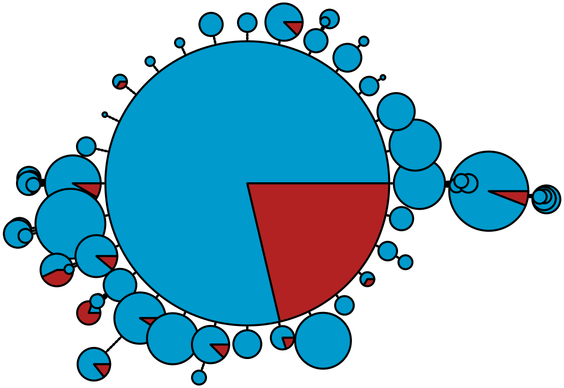

MBPA_000001532 Metazoa

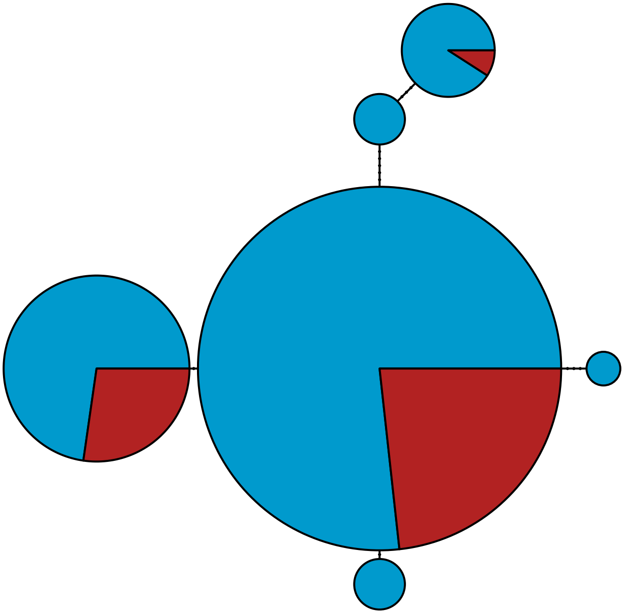

MBPA_000001534 Metazoa

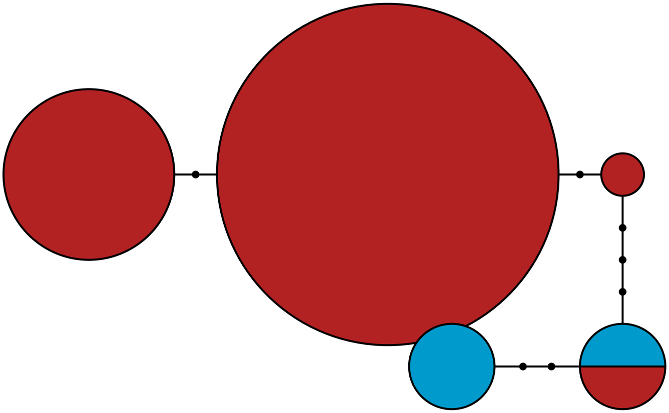

MBPA_000001541 Stramenopiles

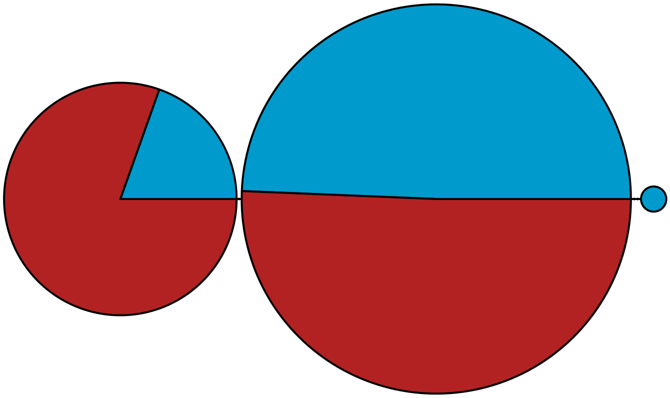

MBPA_000001606 Metazoa

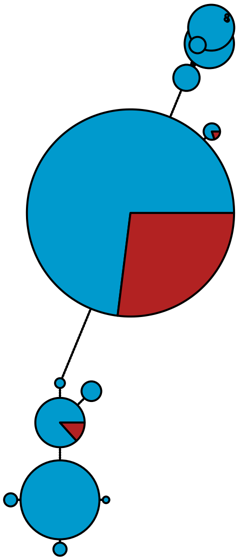

MBPA_000001625

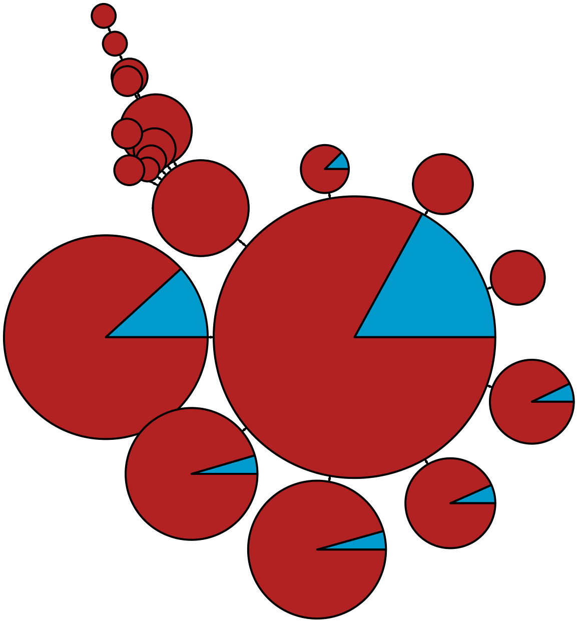

MBPA_000001680 Rhodophyta

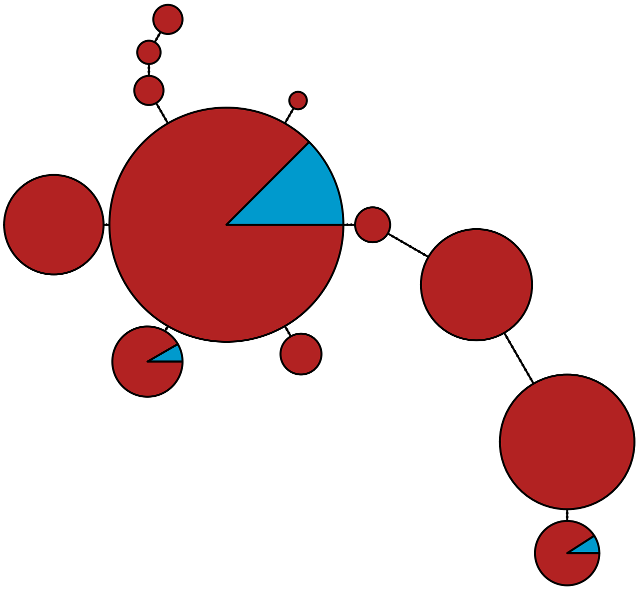

MBPA_000001687 Metazoa

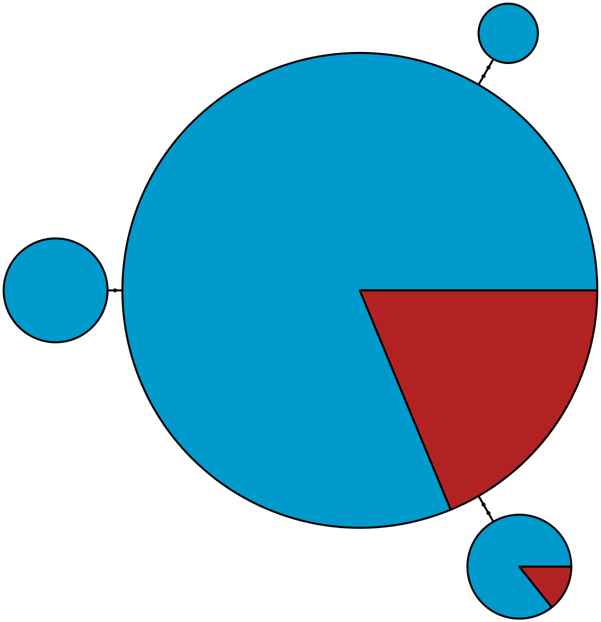

MBPA_000001715 Rhodophyta

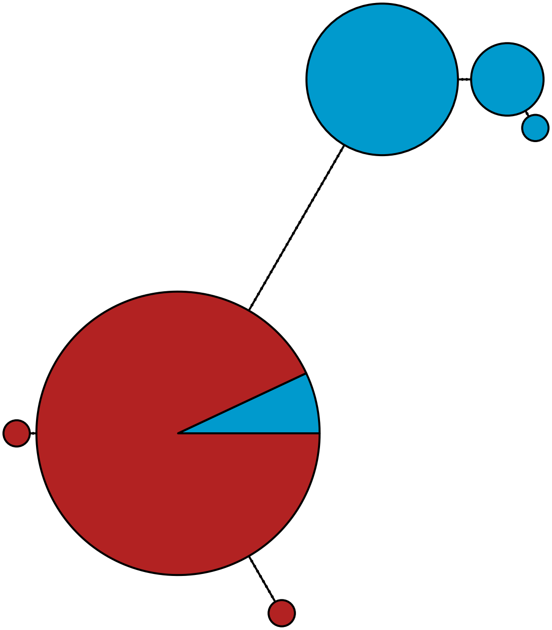

MBPA_000001740 Metazoa

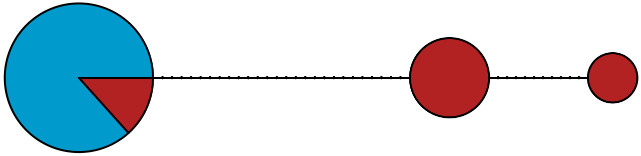

MBPA_000001748 Metazoa

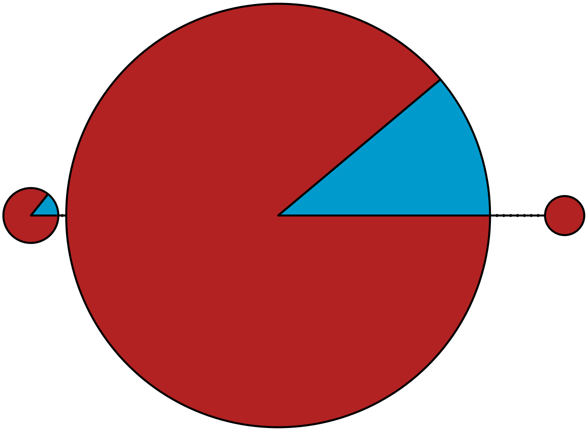

MBPA_000001908 Metazoa

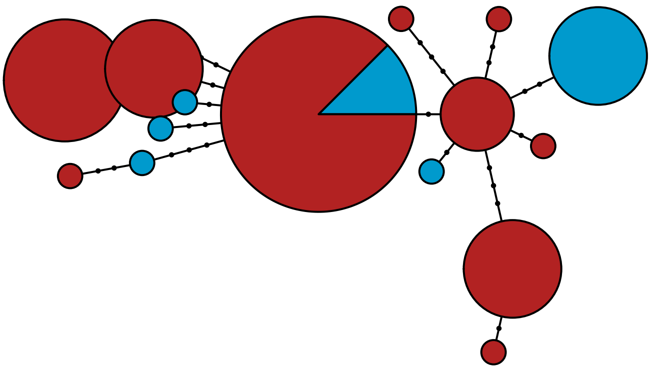

MBPA_000001958 Stramenopiles

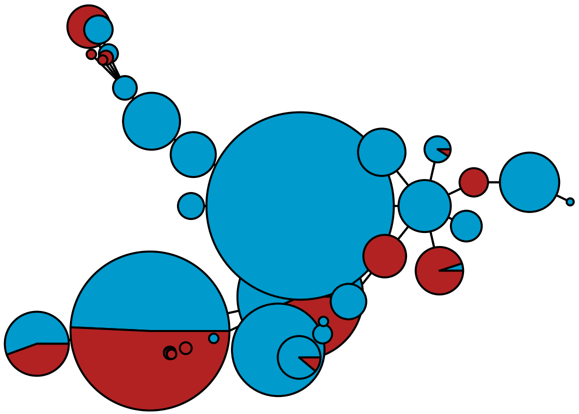

MBPA_000002017 Rhodophyta

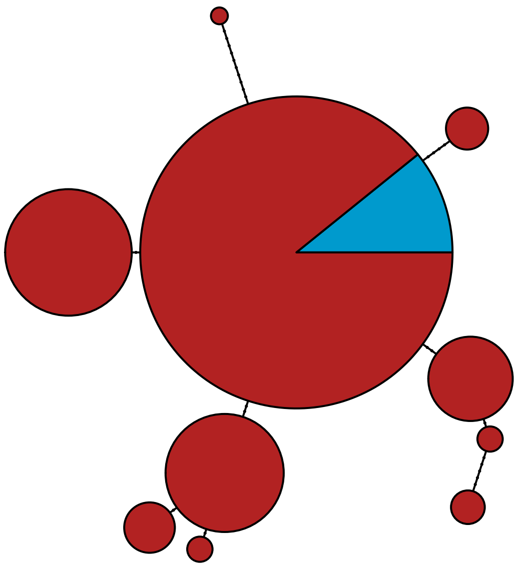

MBPA_000002140 Rhodophyta

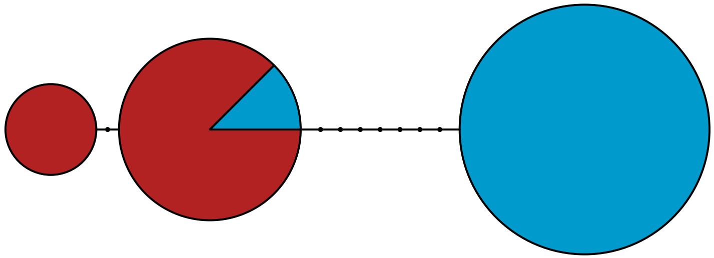

#### MBPA_000002155 Metazoa

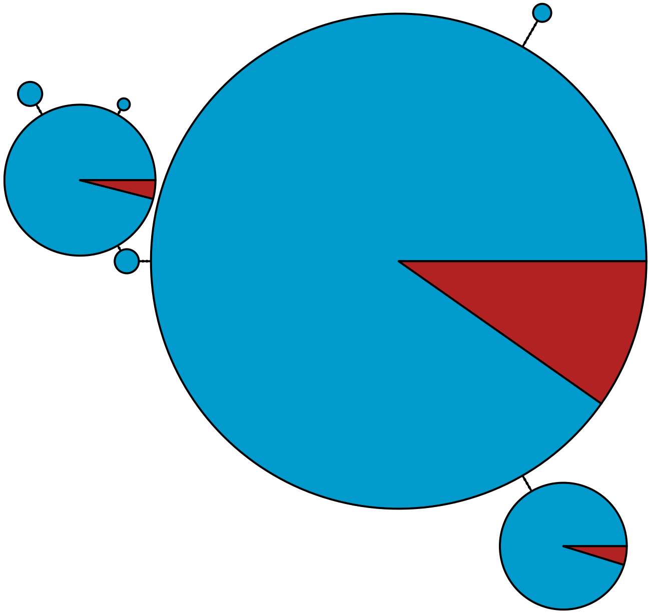

MBPA_000002164 Metazoa

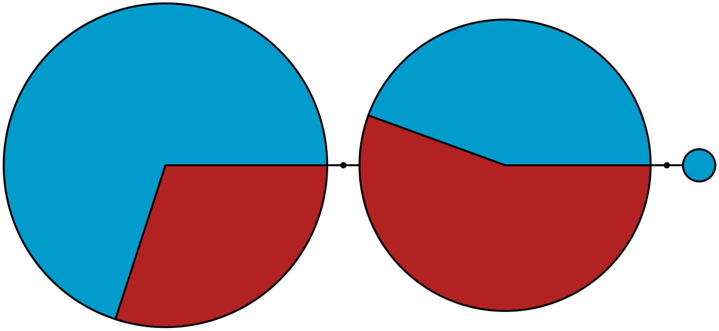

MBPA_000002175 Metazoa

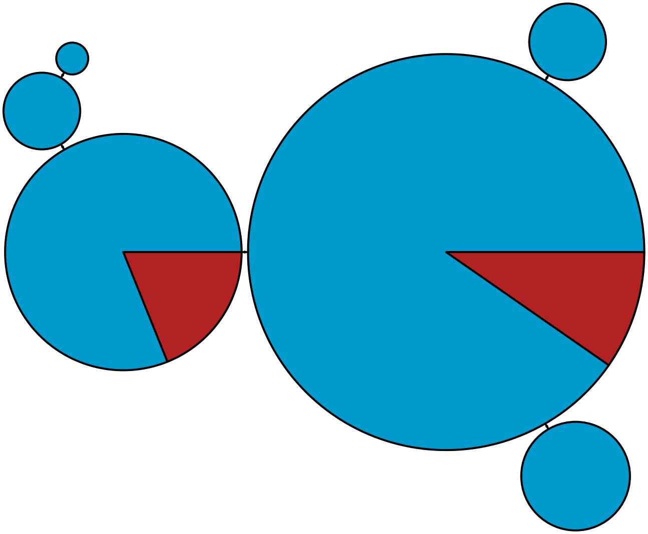

MBPA_000002183 Stramenopiles

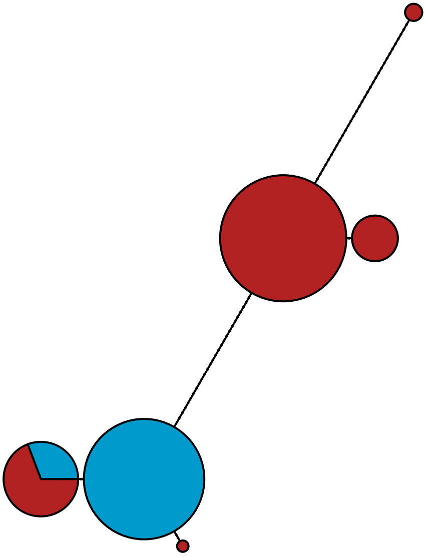

MBPA_000002241 Rhodophyta

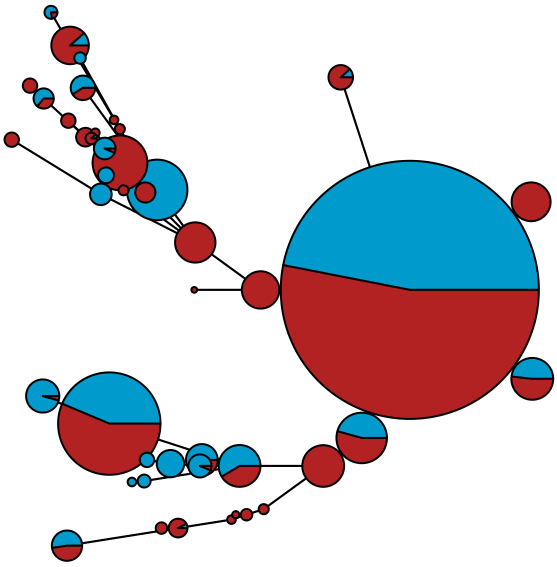

MBPA_000002268 Metazoa

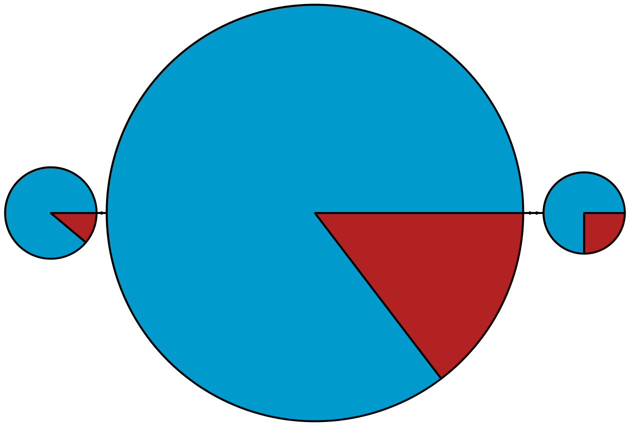

MBPA_000002274 Metazoa

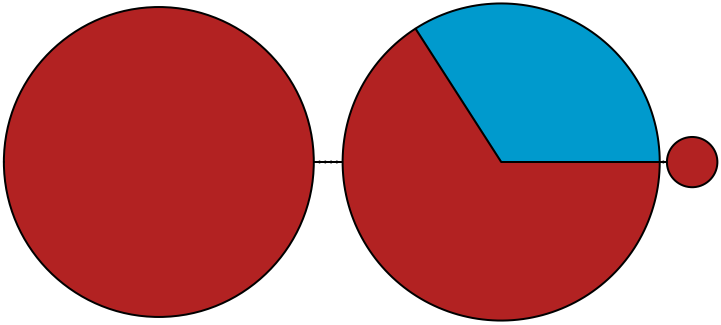

MBPA_000002284 Stramenopiles

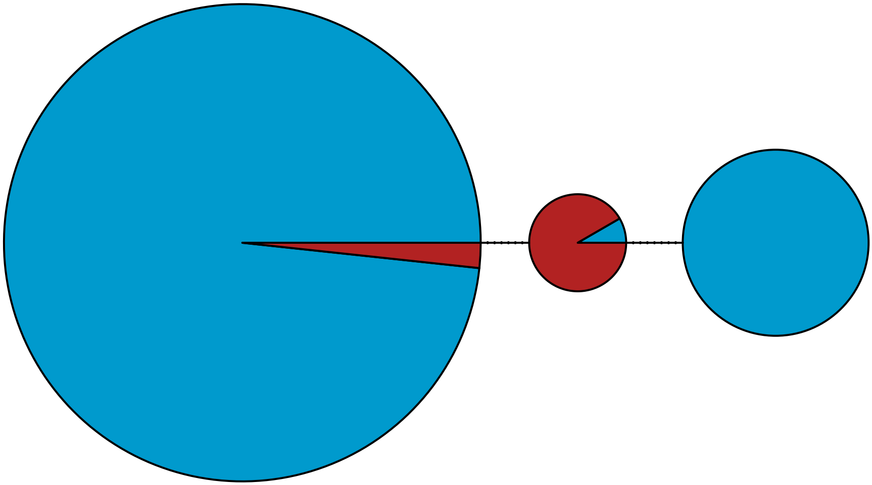

MBPA_000002320 Rhodophyta

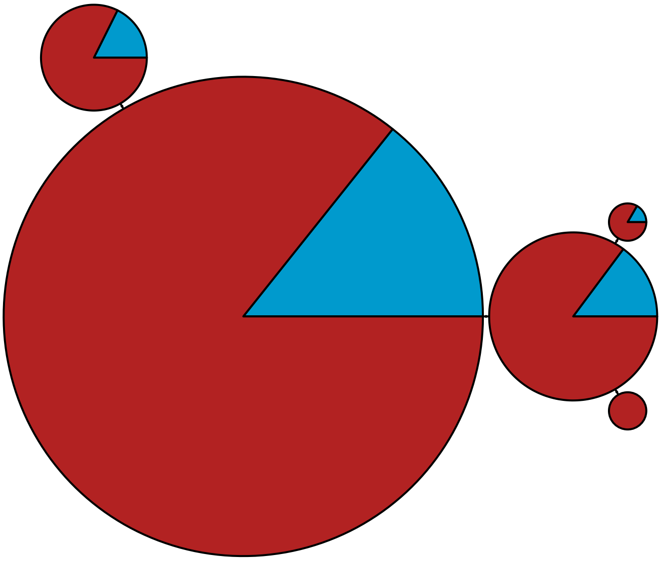

MBPA_000002335 Metazoa

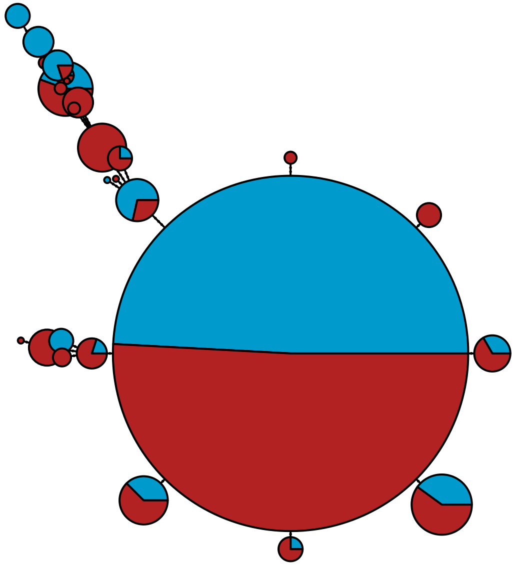

MBPA_000002368 Metazoa

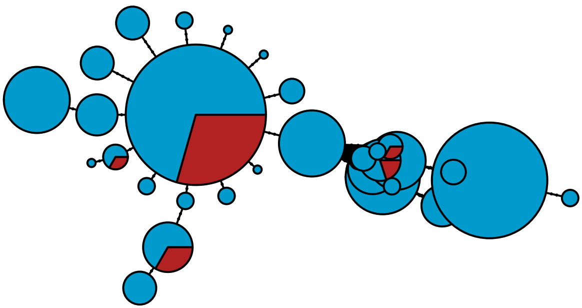

MBPA_000002373 Rhodophyta

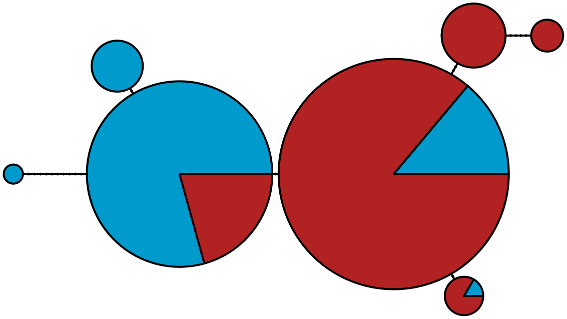

MBPA_000002384 Metazoa

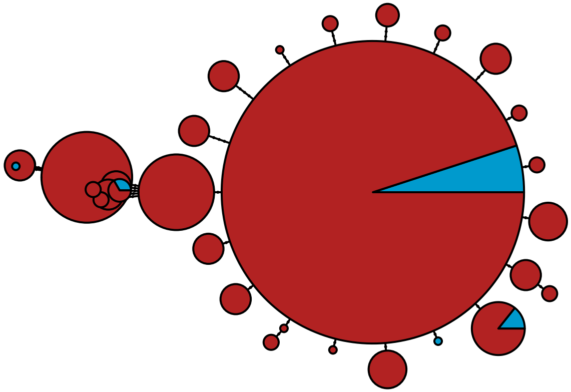

MBPA_000002404 Rhodophyta

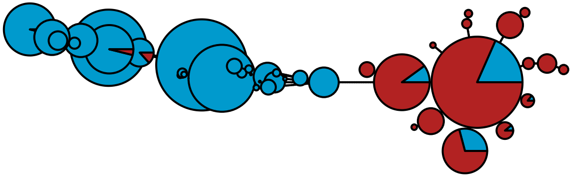

### MBPA_000002500 Metazoa

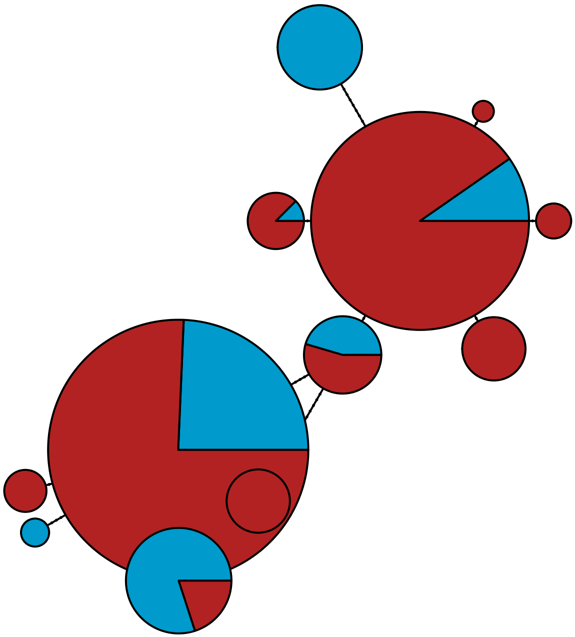

MBPA_000002581 Metazoa

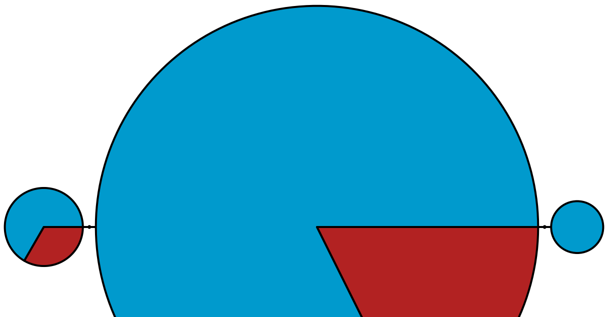

MBPA_000002597 Rhodophyta

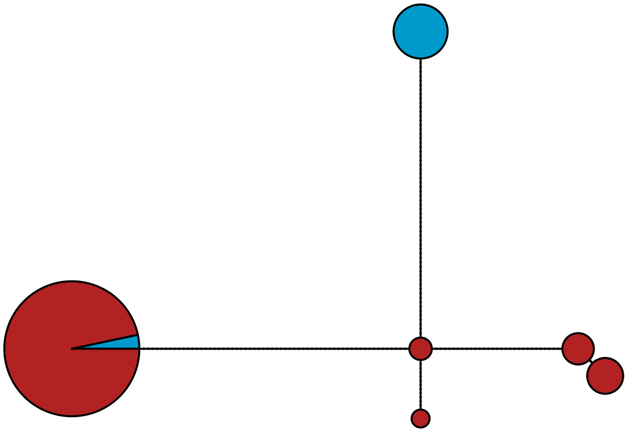

MBPA_000002708

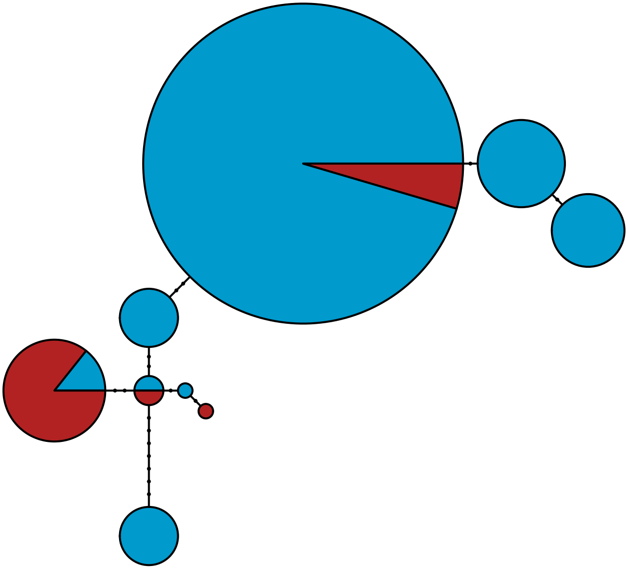

MBPA_000002792 Rhodophyta

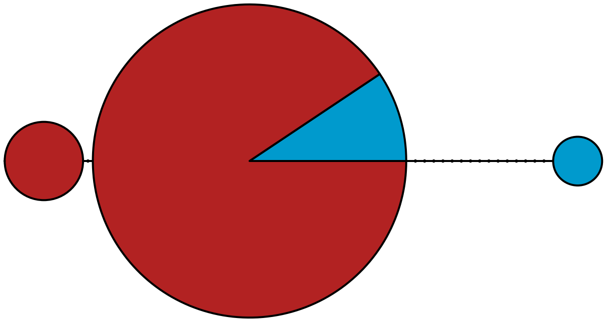

MBPA_000002796 Metazoa

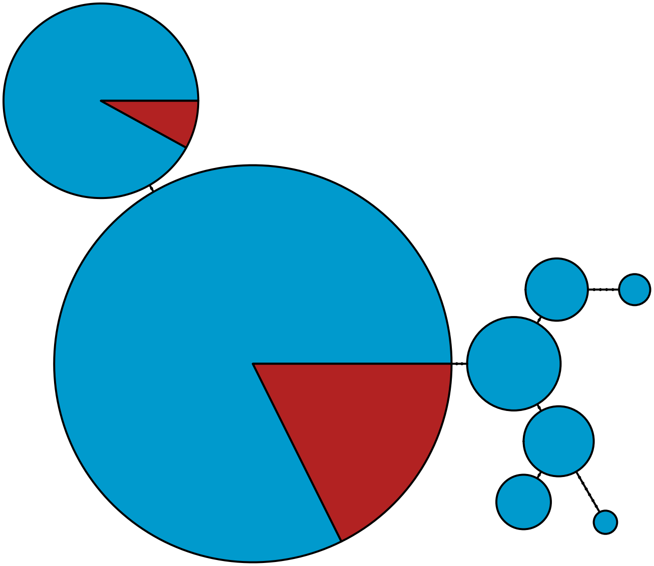

MBPA_000002801 Stramenopiles

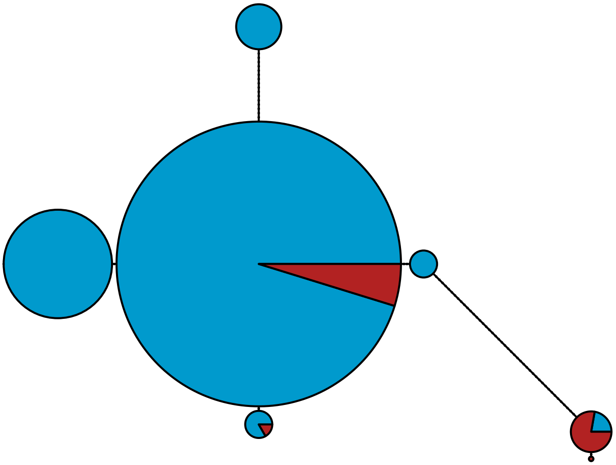

MBPA_000002808 Metazoa

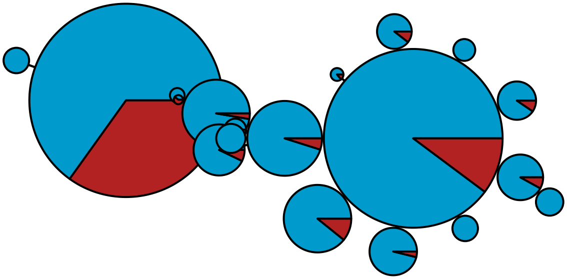

MBPA_000002905 Metazoa

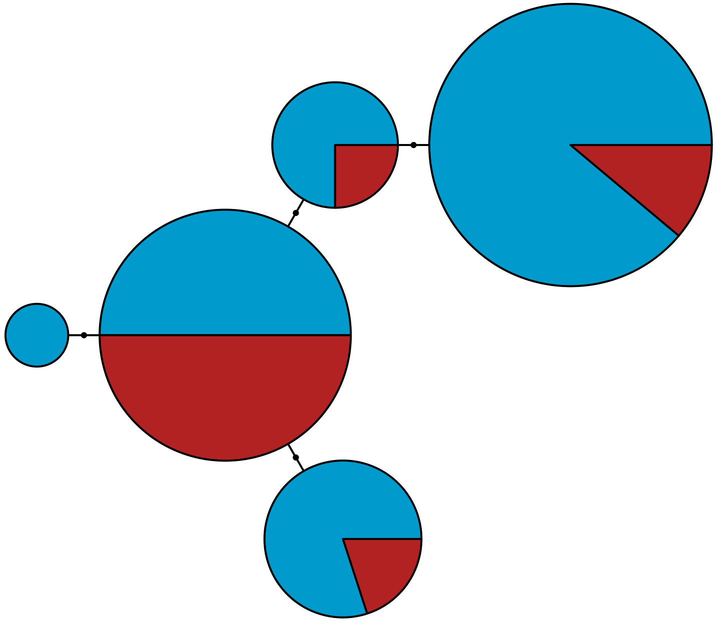

MBPA_000002906 Metazoa

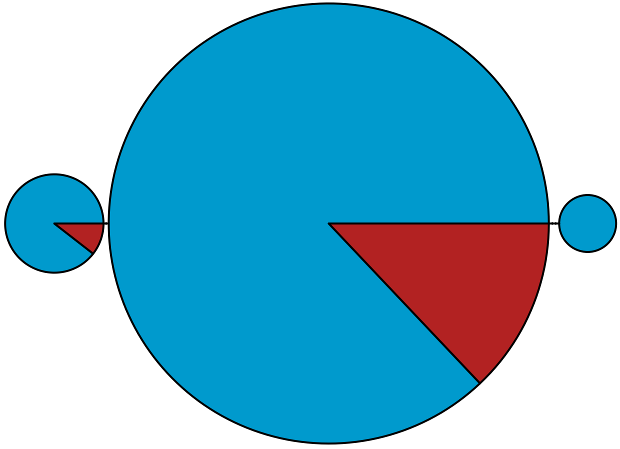

MBPA_000002932 Metazoa

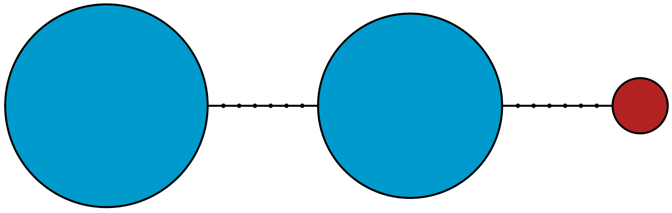

MBPA_000002984 Rhodophyta

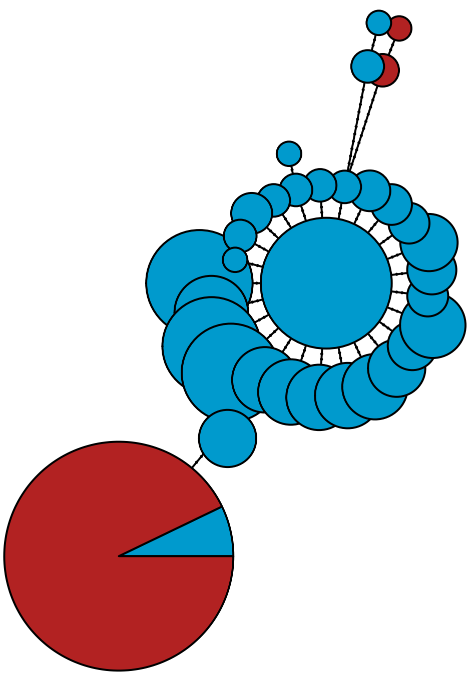

MBPA_000003116 Metazoa

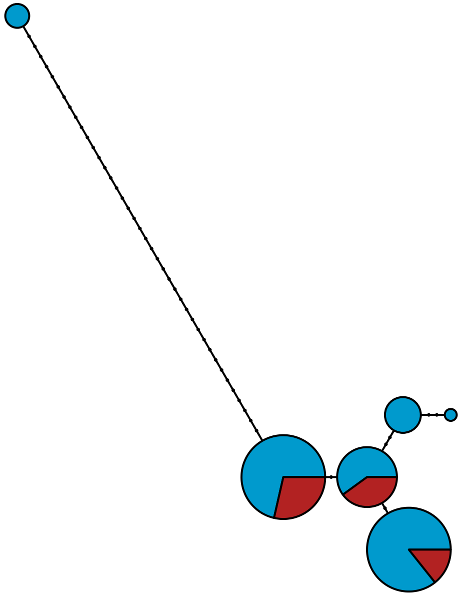

MBPA_000003133 Metazoa

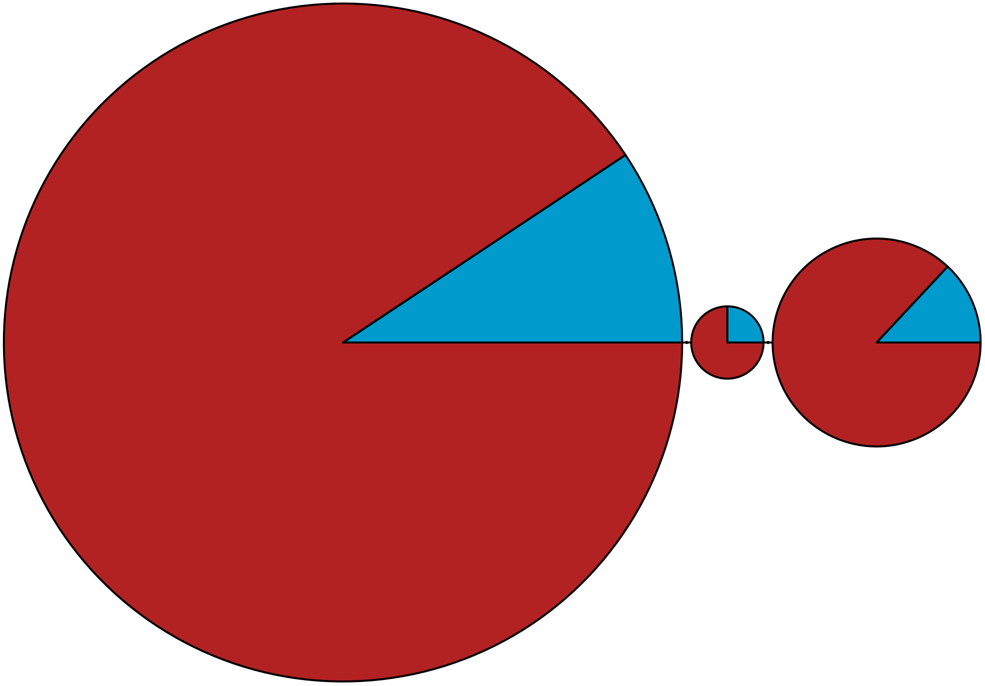

MBPA_000003178 Metazoa

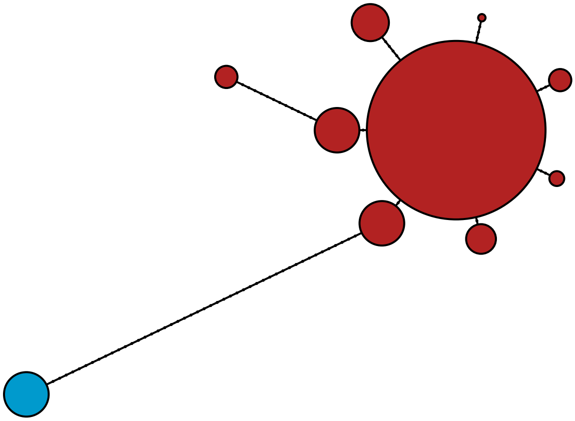

MBPA_000003207 Metazoa

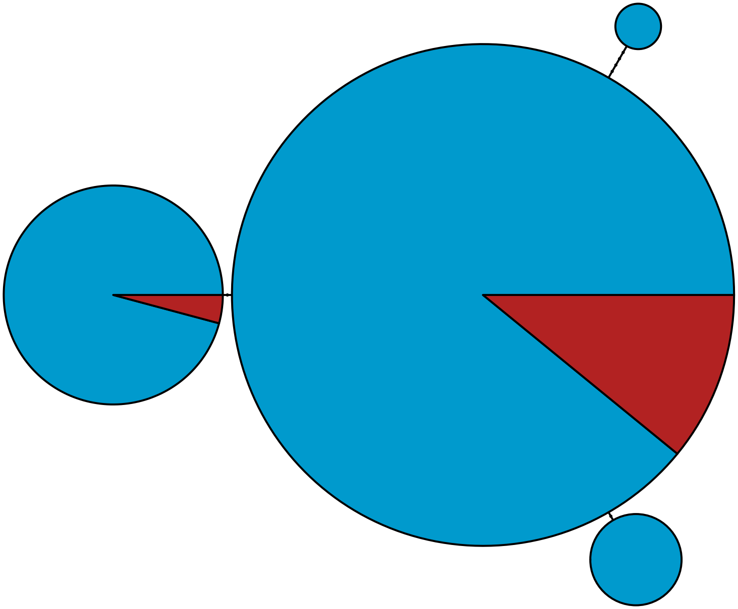

MBPA_000003261 Metazoa

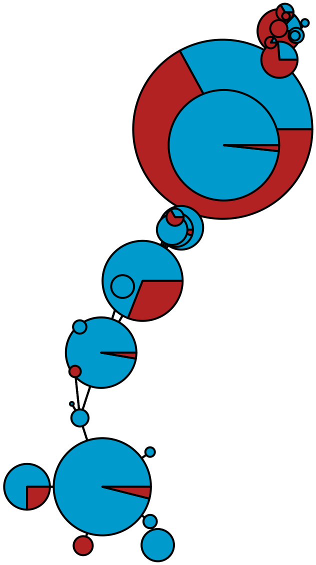

MBPA_000003289 Stramenopiles

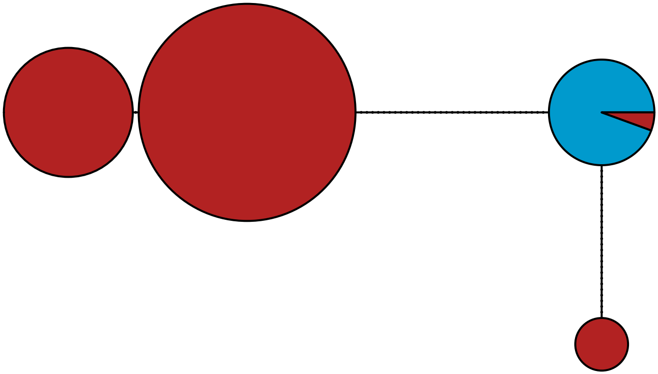

MBPA_000003295 Stramenopiles

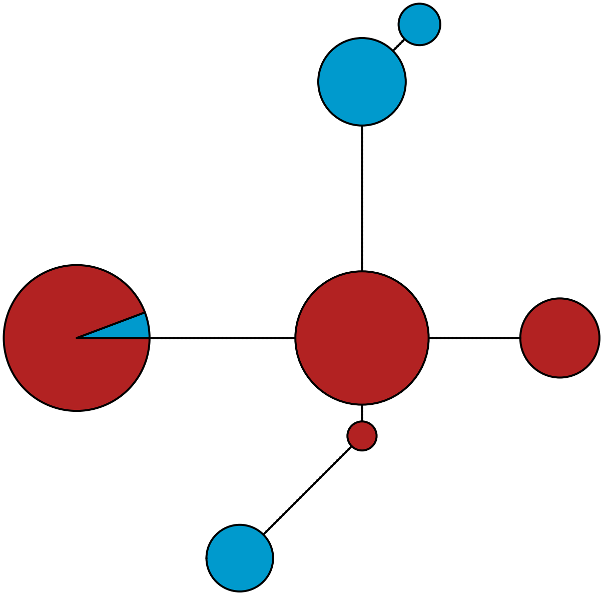

MBPA_000003304 Metazoa

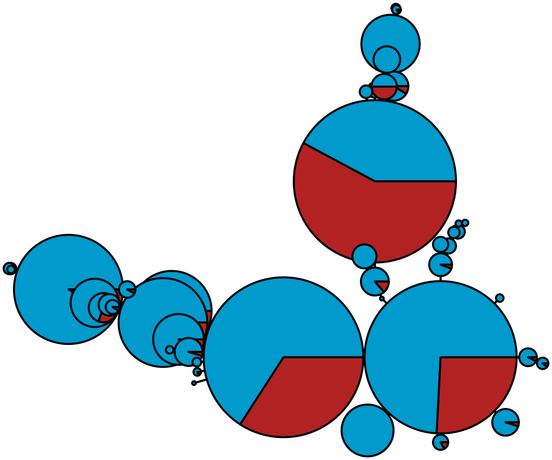

MBPA_000003418 Metazoa

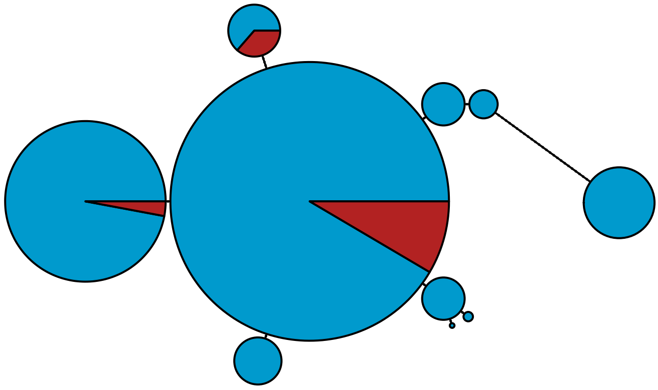

MBPA_000003528 Metazoa

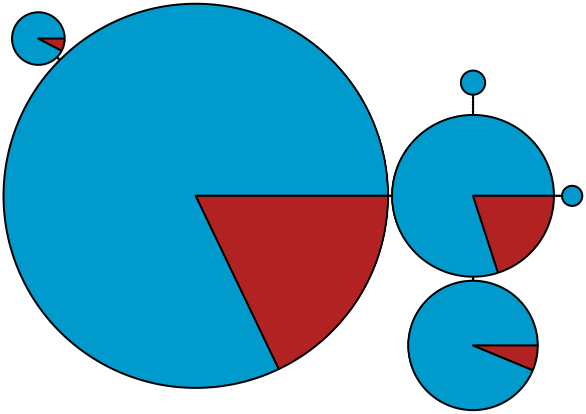

MBPA_000003530 Rhodophyta

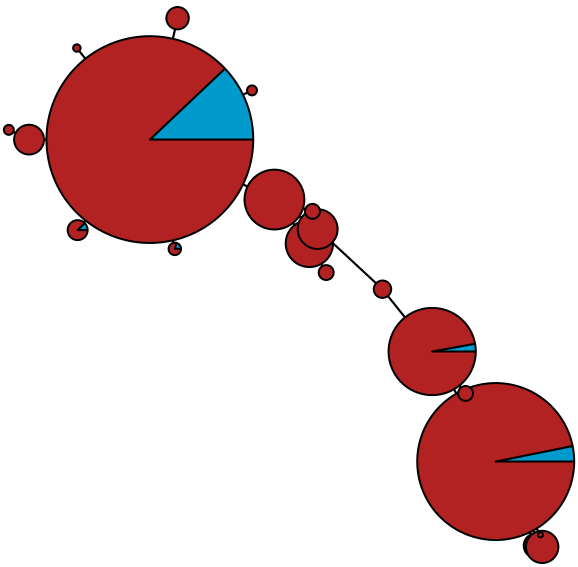

MBPA_000003535 Metazoa

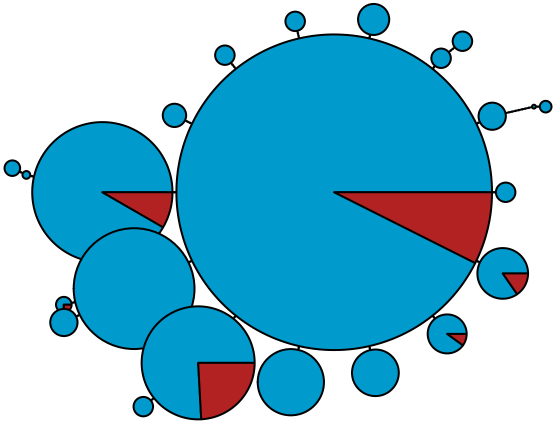

MBPA_000003542 Rhodophyta

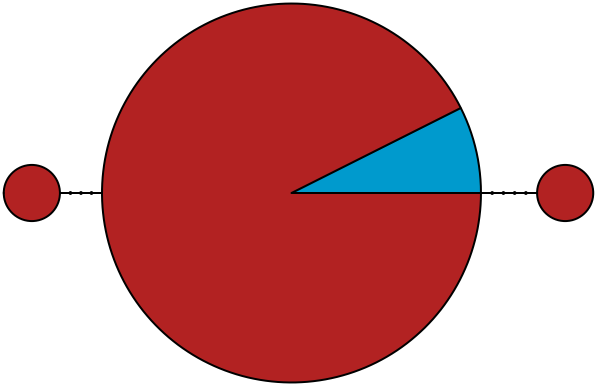

MBPA_000003576 Metazoa

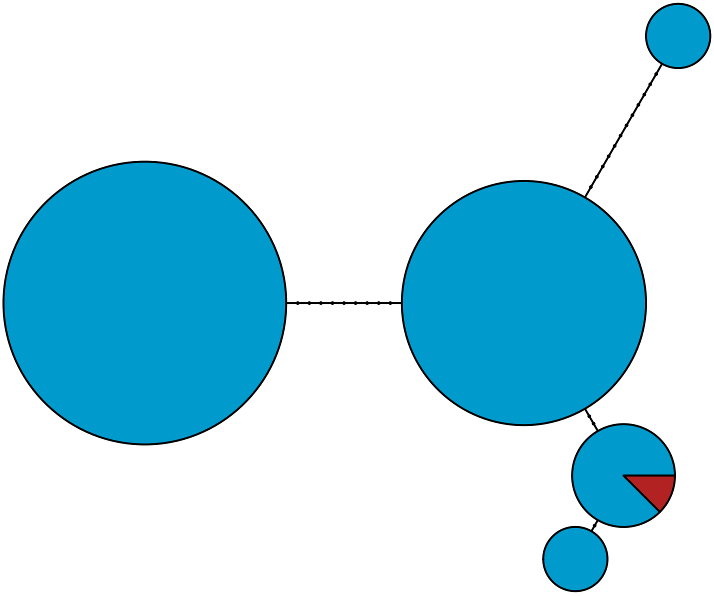

MBPA_000003744 Metazoa

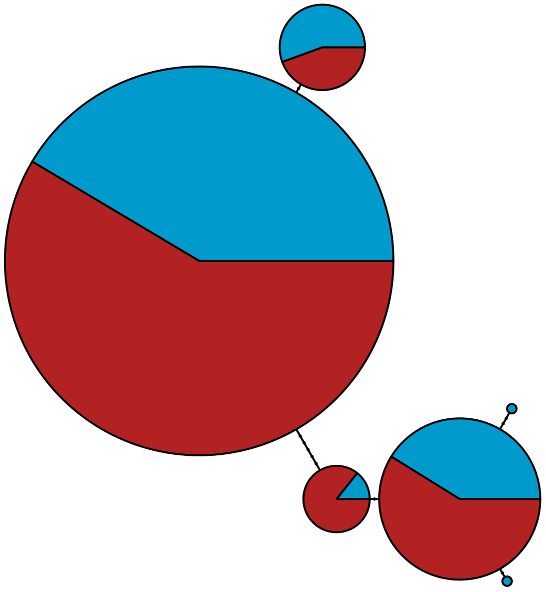

MBPA_000003840 Metazoa

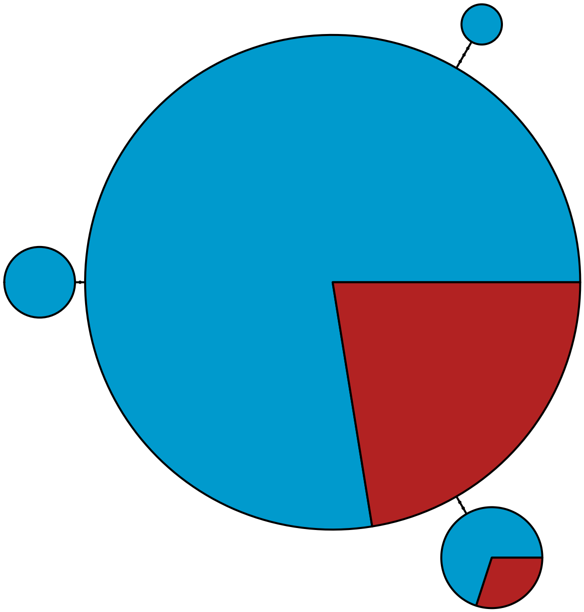

MBPA_000003904 Metazoa

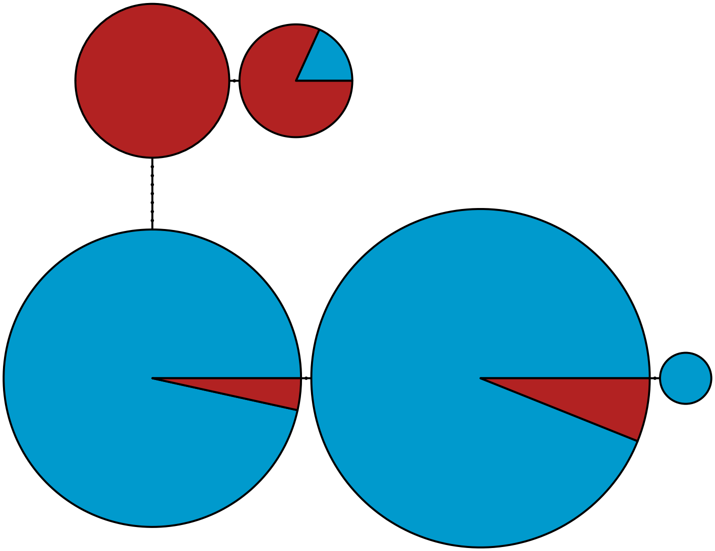

MBPA_000003913 Metazoa

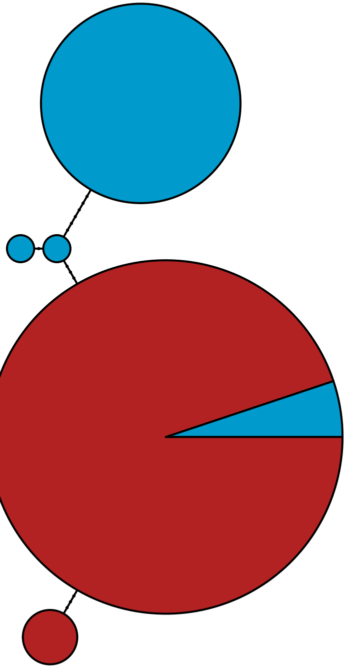

MBPA_000004142 Stramenopiles

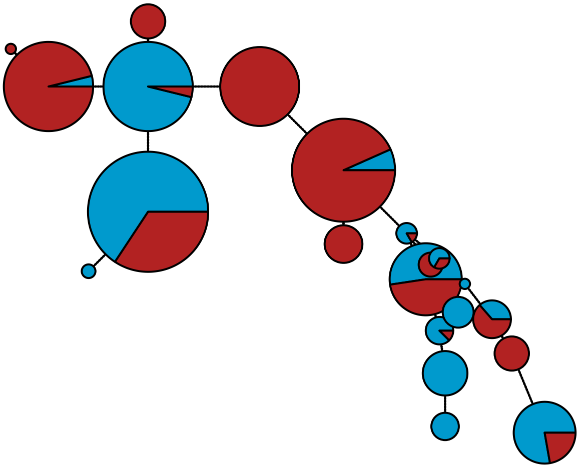

MBPA_000004294 Metazoa

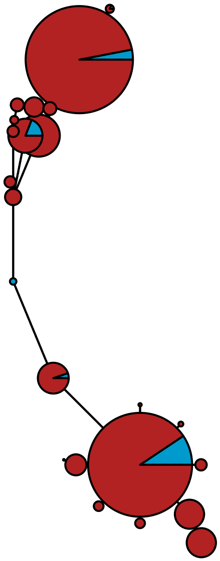

MBPA_000004478 Metazoa

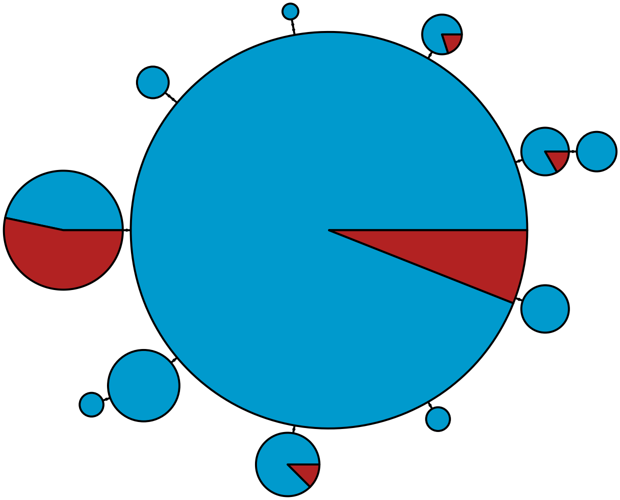

MBPA_000004483 Metazoa

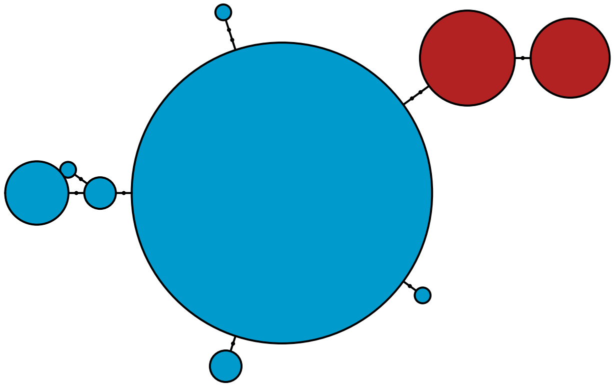

MBPA_000004503

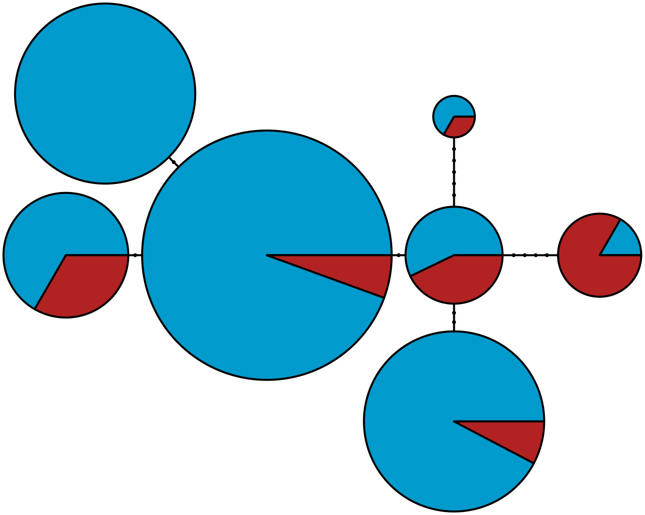

MBPA_000004561 Stramenopiles

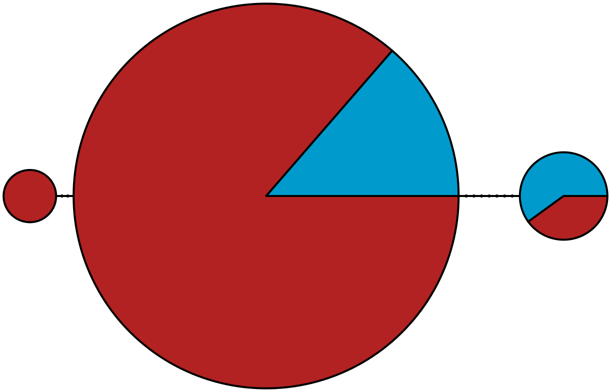

MBPA_000004645 Metazoa

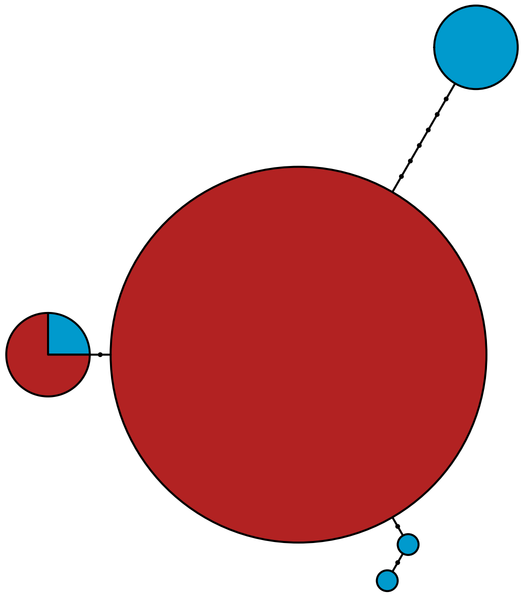

MBPA_000004675 Rhodophyta

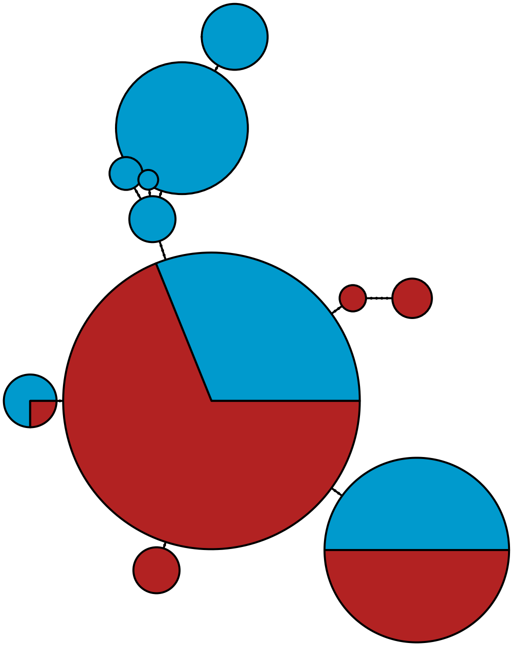

MBPA_000004681

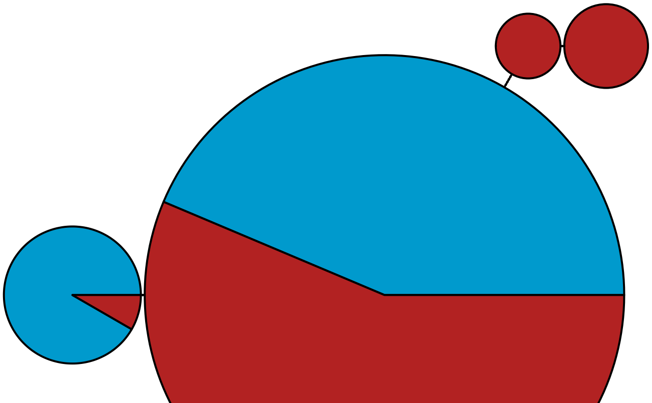

MBPA_000004688 Stramenopiles

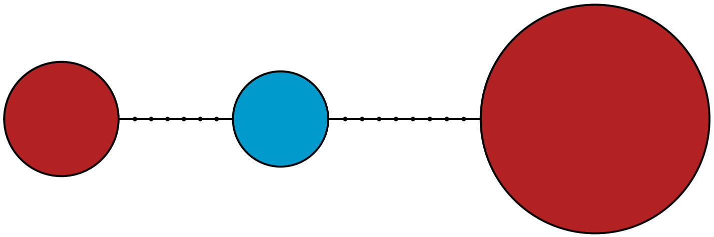

MBPA_000004696

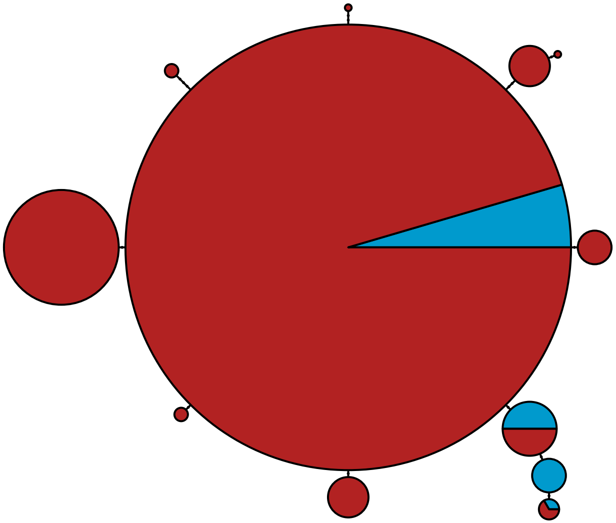

MBPA_000004765 Metazoa

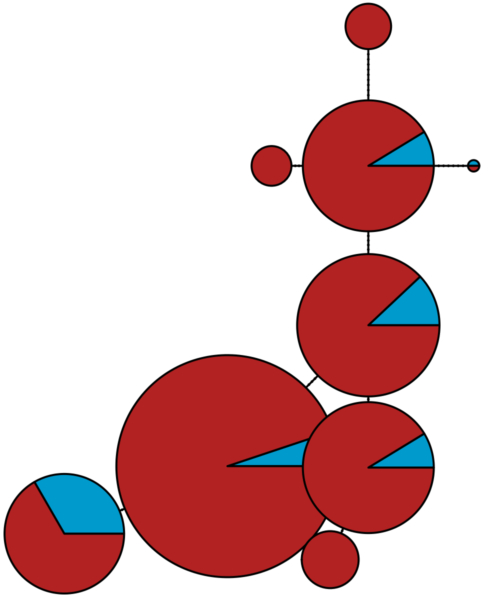

MBPA_000004906 Metazoa

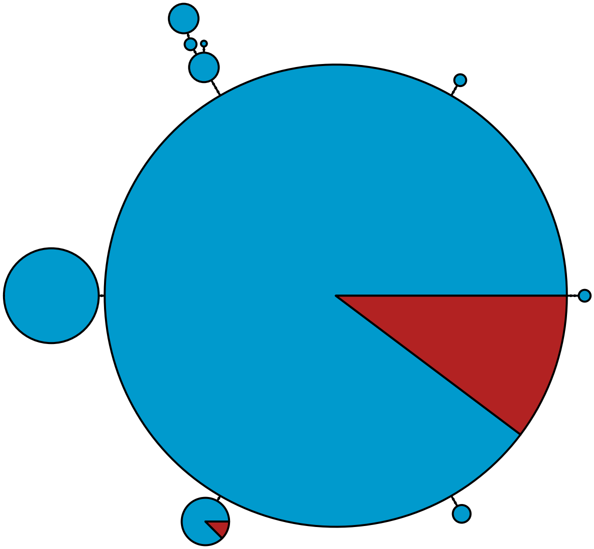

### MBPA_000005170 Metazoa

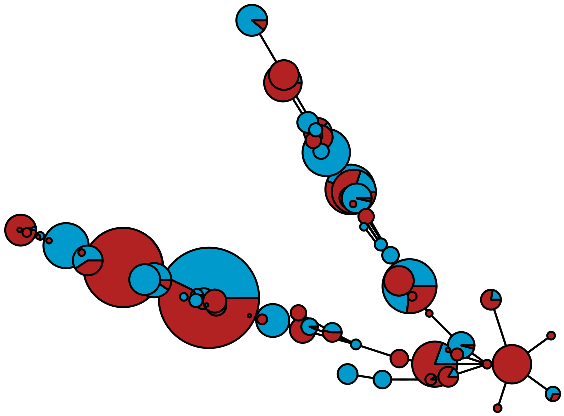

MBPA_000005205 Rhodophyta

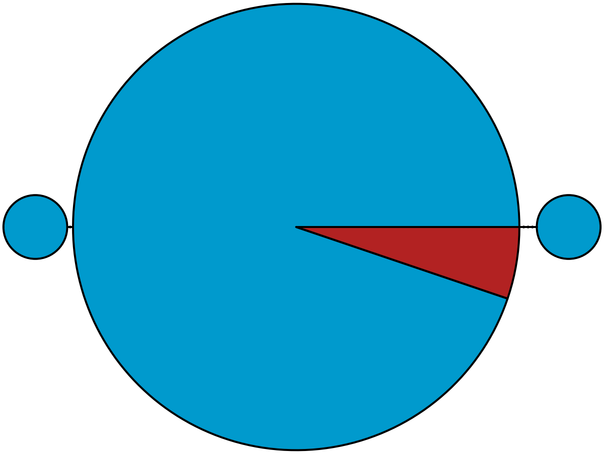

MBPA_000005225 Metazoa

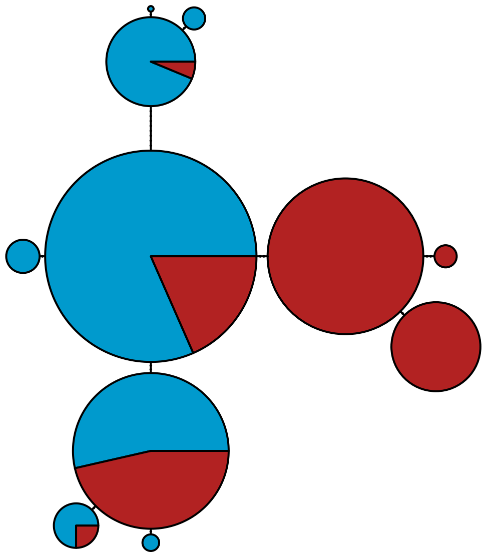

MBPA_000005265 Metazoa

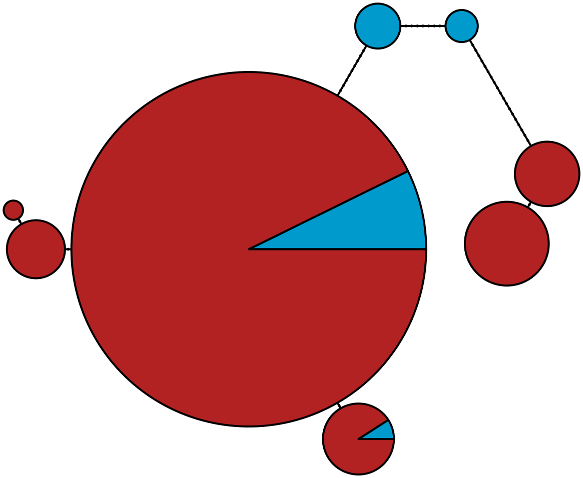

MBPA_000005365 Metazoa

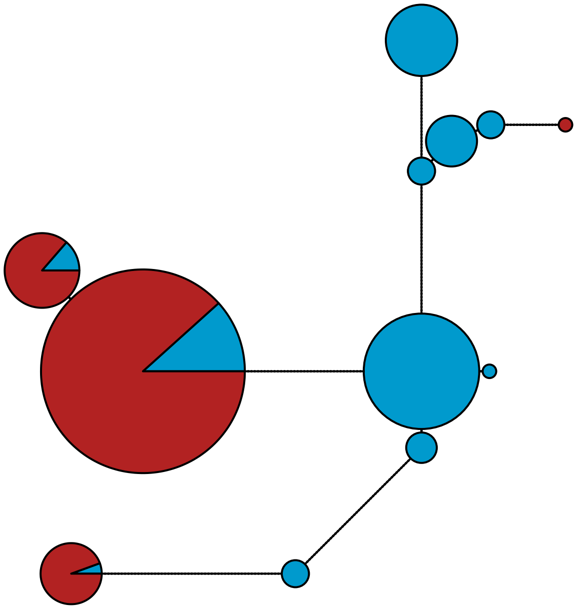

MBPA_000005379 Metazoa

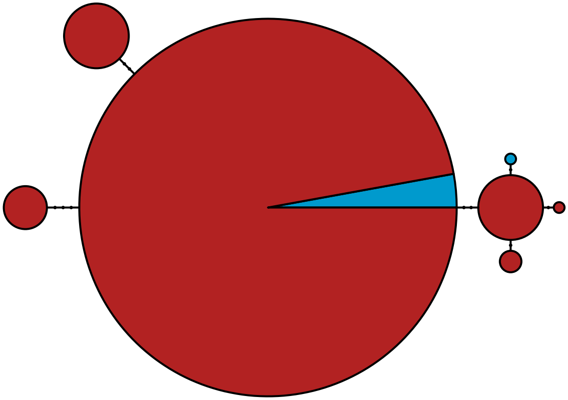

MBPA_000005497 Metazoa

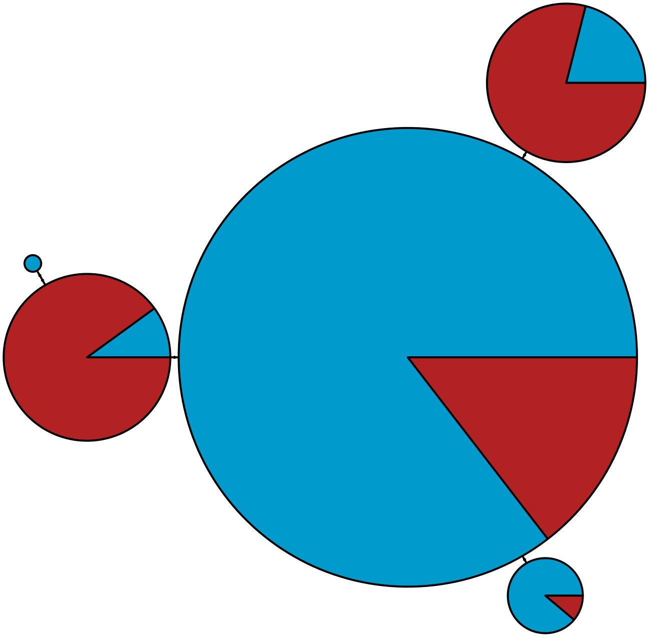

MBPA_000005561 Metazoa

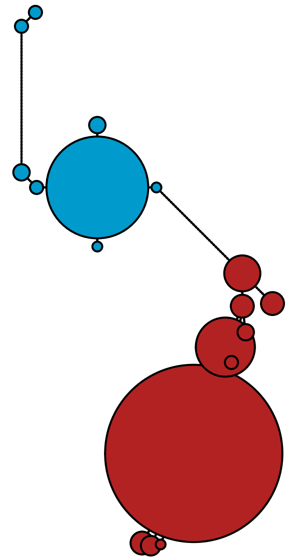

MBPA_000005961 Metazoa

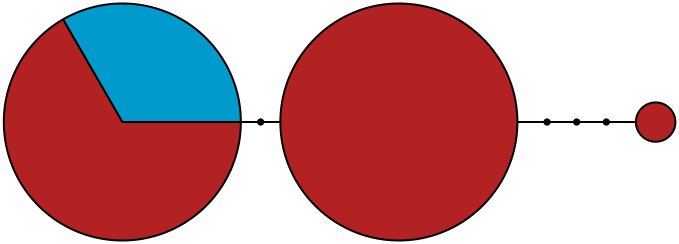

MBPA_000006003 Rhodophyta

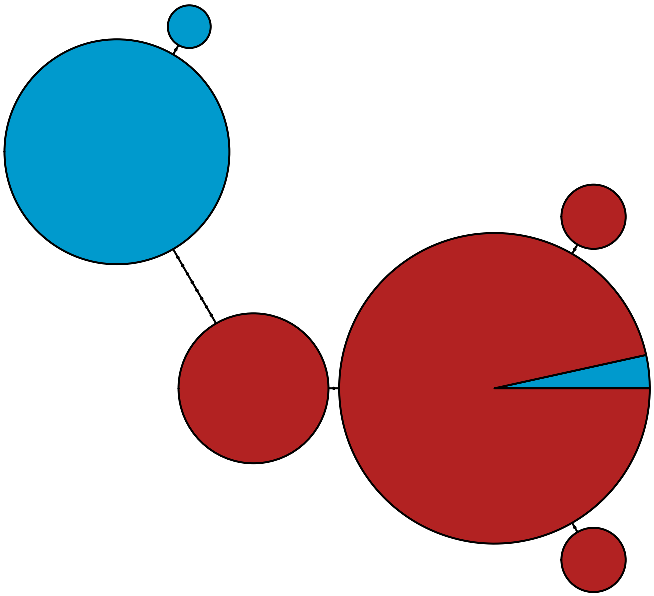

MBPA_000006026 Metazoa

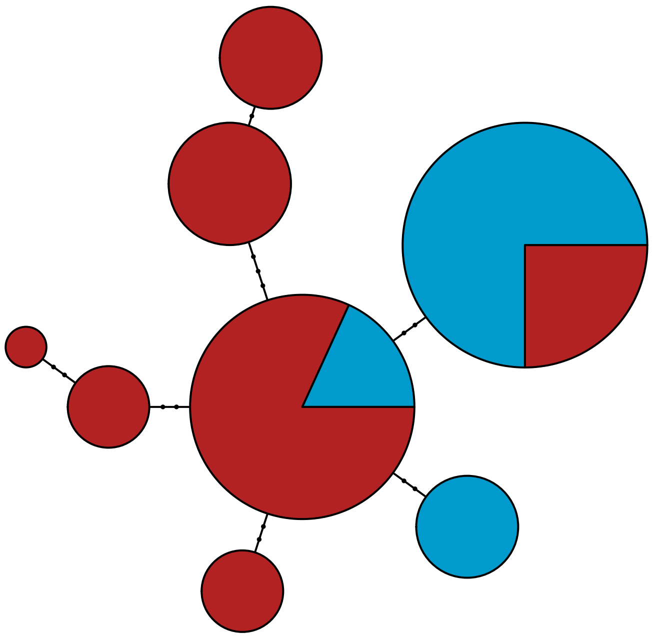

MBPA_000006050 Rhodophyta

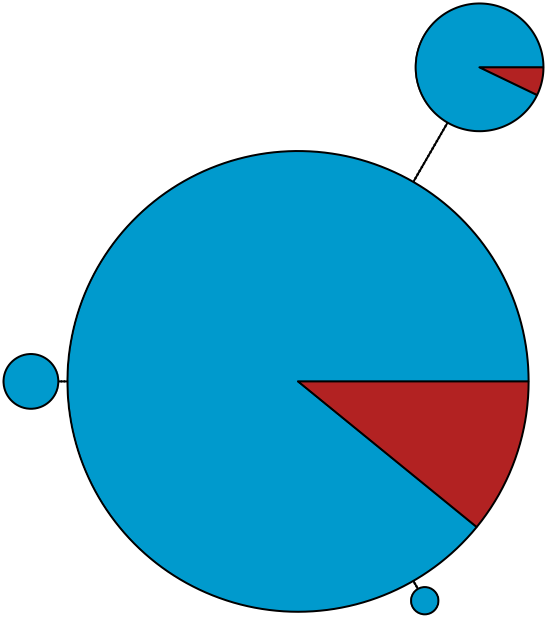

MBPA_000006086 Rhodophyta

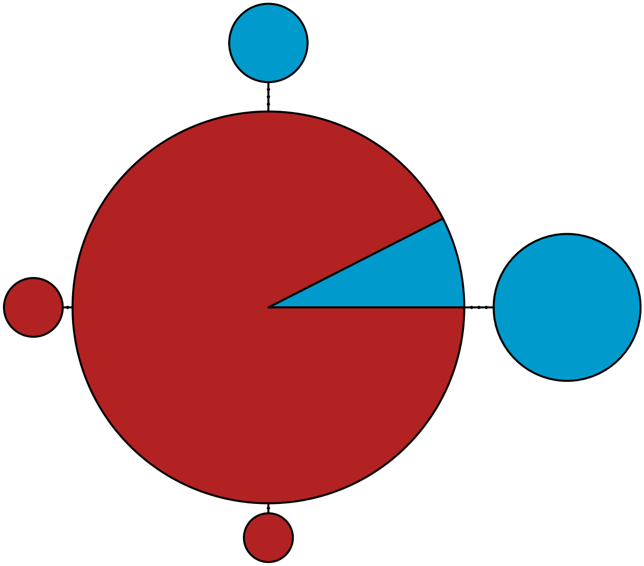

MBPA_000006103 Metazoa

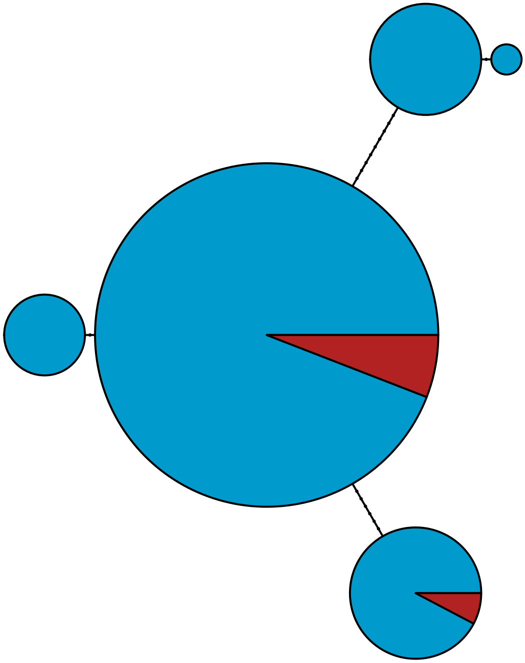

MBPA_000006224

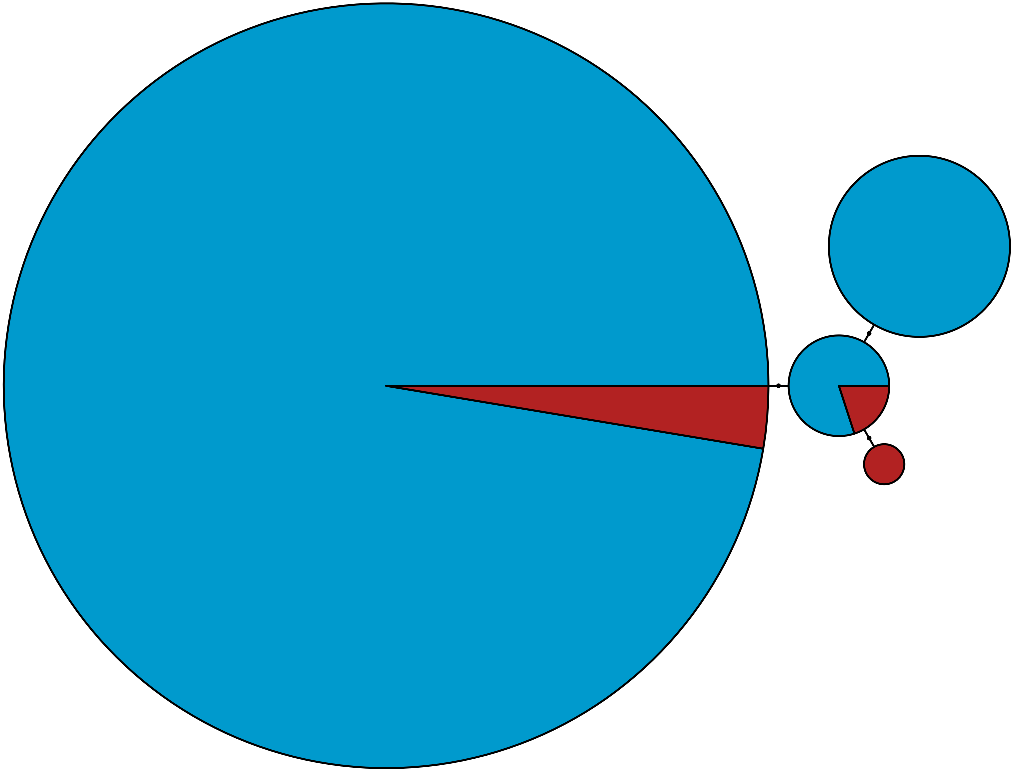

MBPA_000006572 Rhodophyta

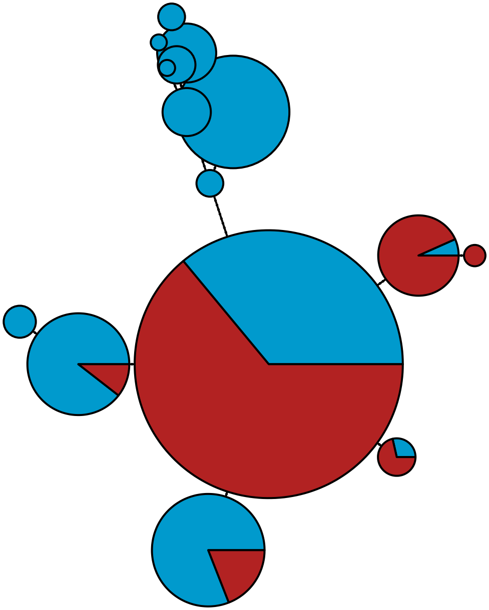

MBPA_000006708 Rhodophyta

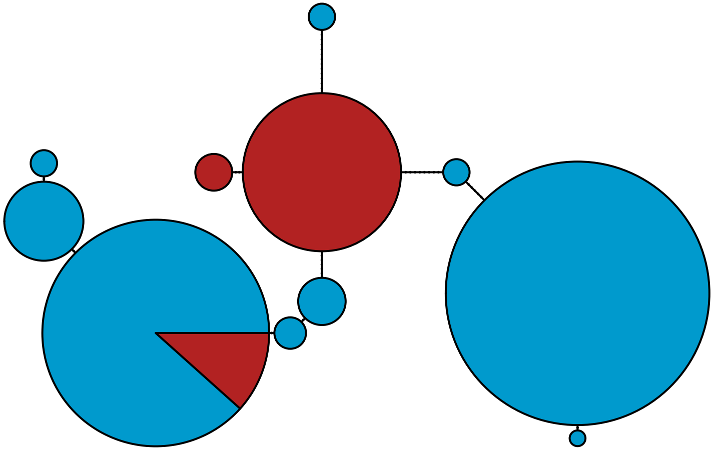

MBPA_000006823 Metazoa

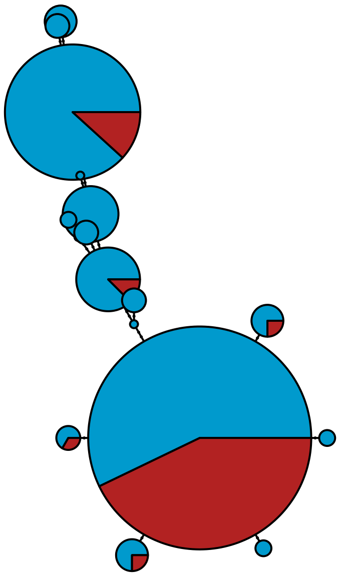

MBPA_000006837

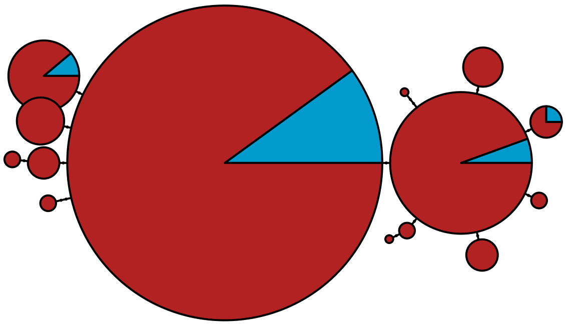

MBPA_000007078 Metazoa

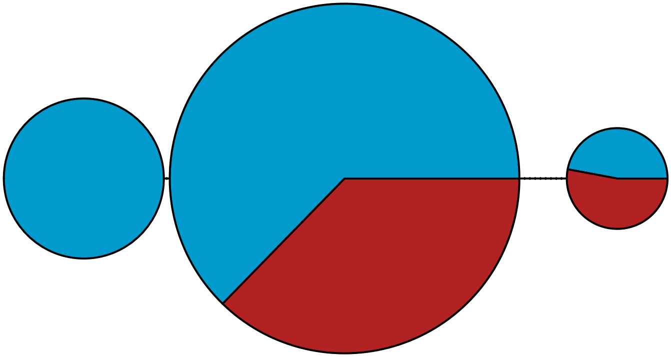

MBPA_000007262

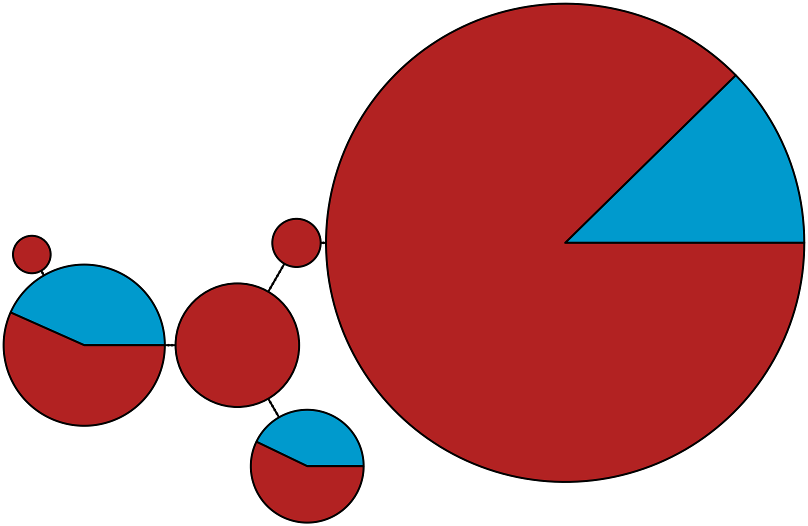

MBPA_000007294 Rhodophyta

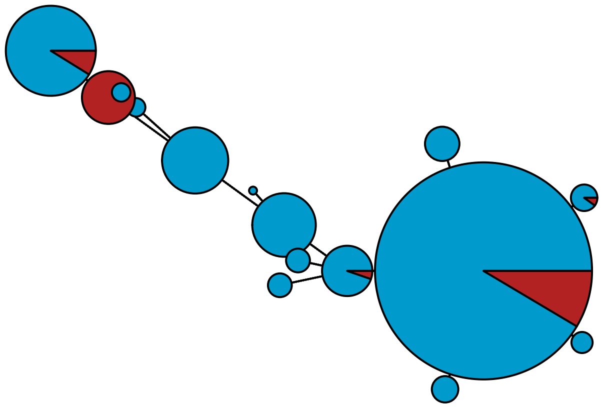

MBPA_000007328

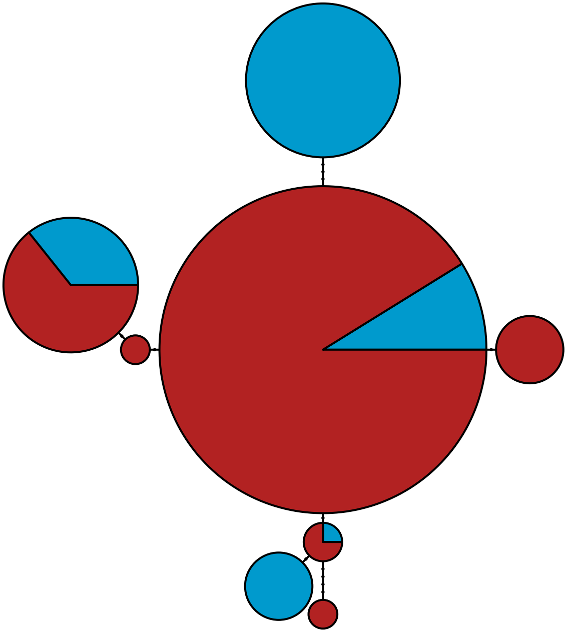

MBPA_000007354

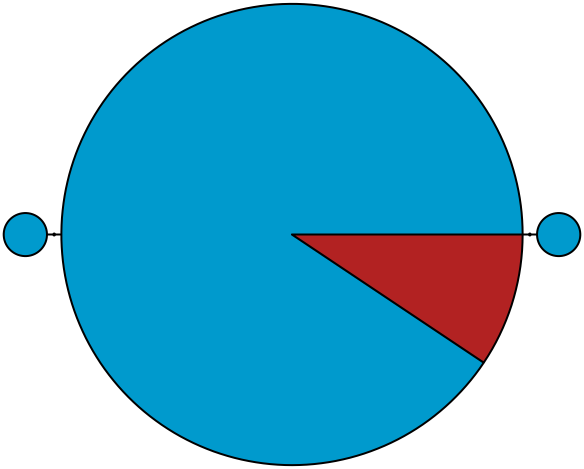

MBPA_000007400 Rhodophyta

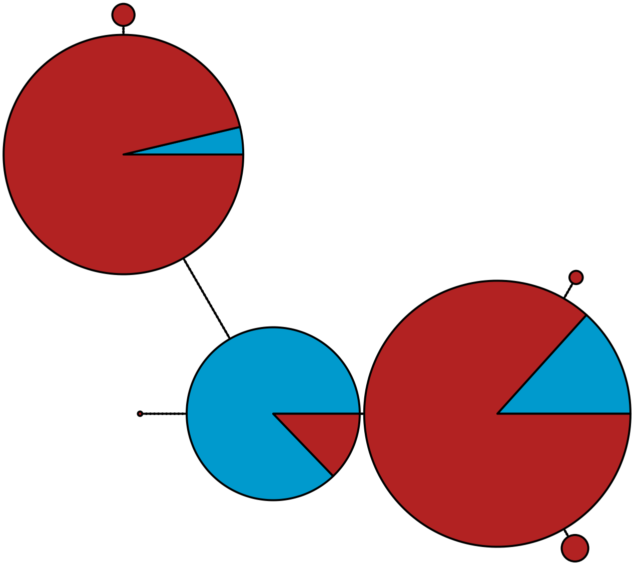

MBPA_000007428 Lobosa

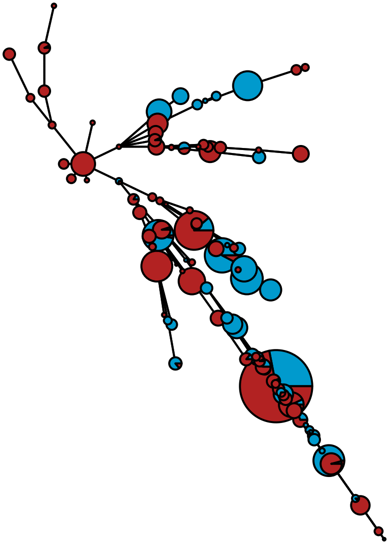

MBPA_000007461 Rhodophyta

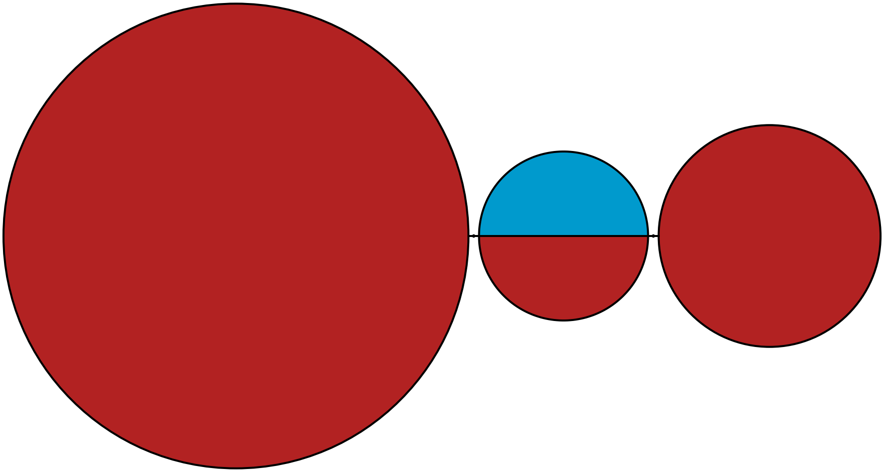

MBPA_000007565 Stramenopiles

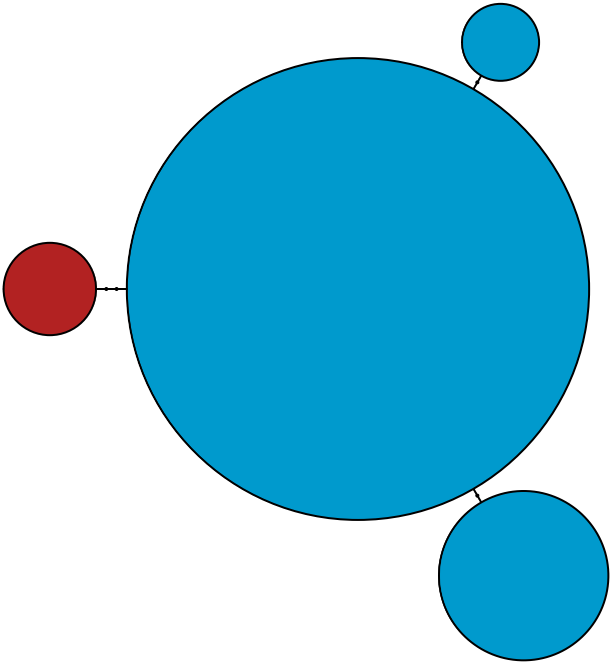

MBPA_000007571 Metazoa

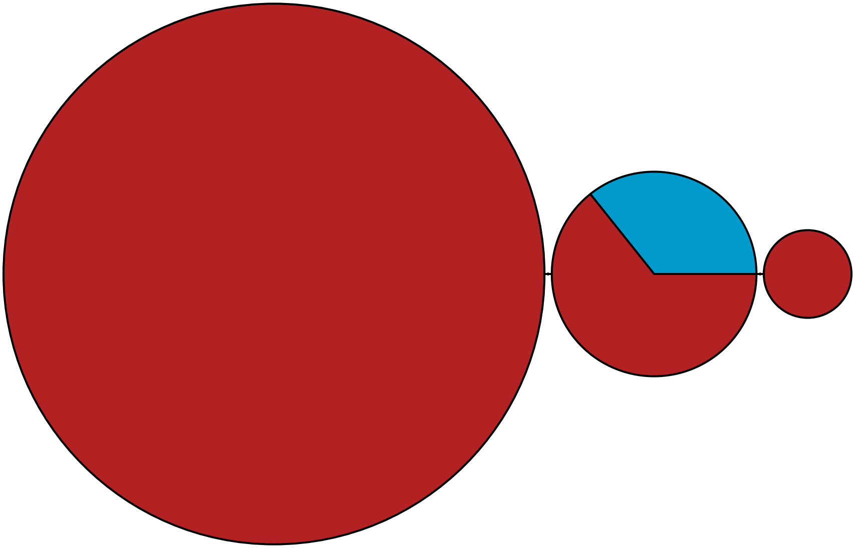

MBPA_000007824 Metazoa

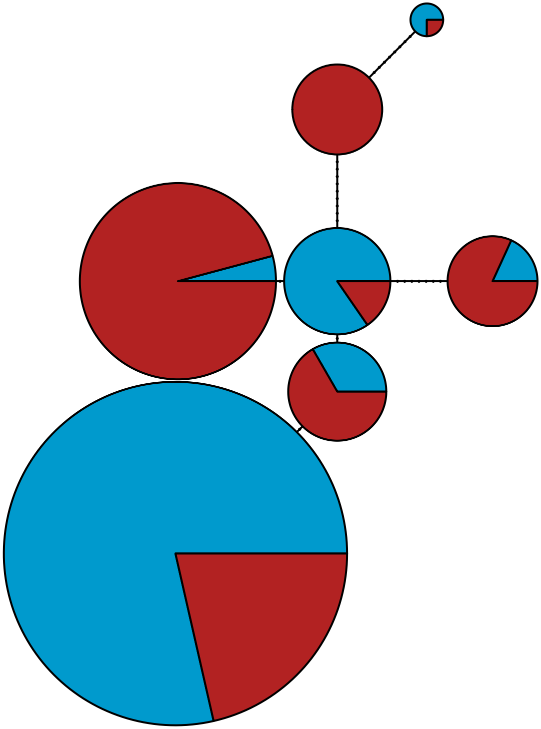

MBPA_000007825 Stramenopiles

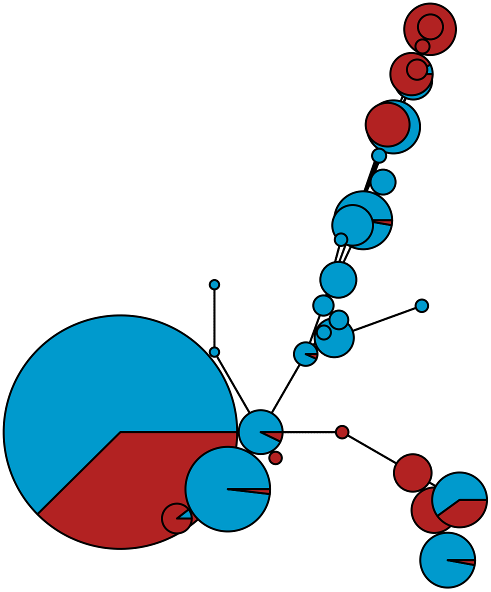

MBPA_000007879 Rhodophyta

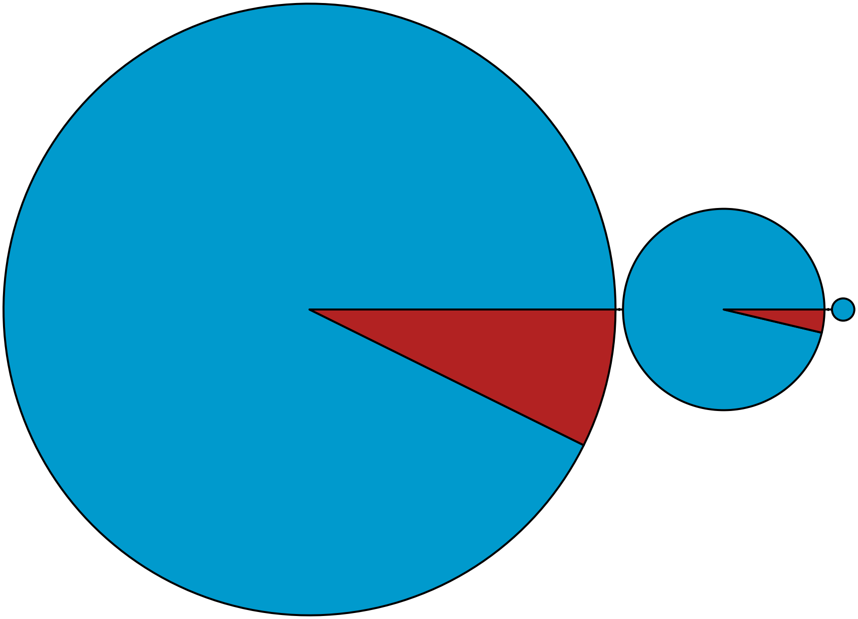

MBPA_000007980 Stramenopiles

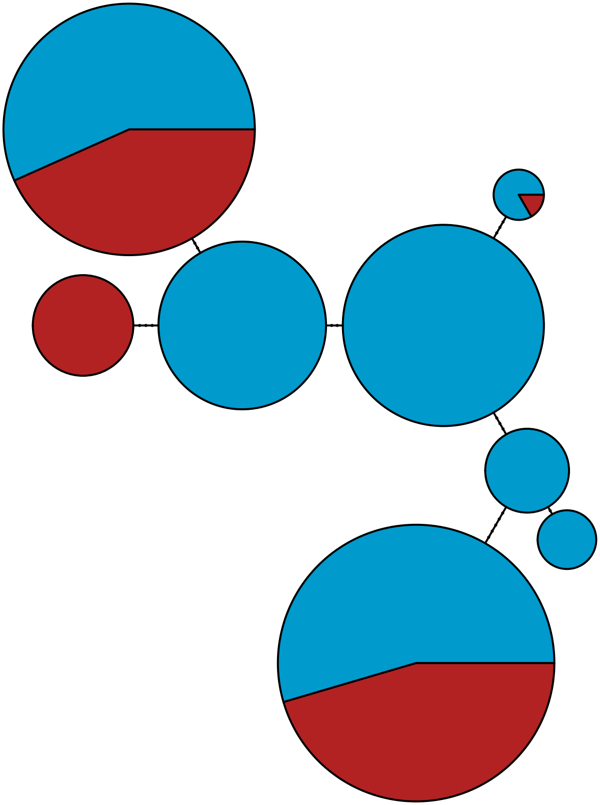

MBPA_000007992 Metazoa

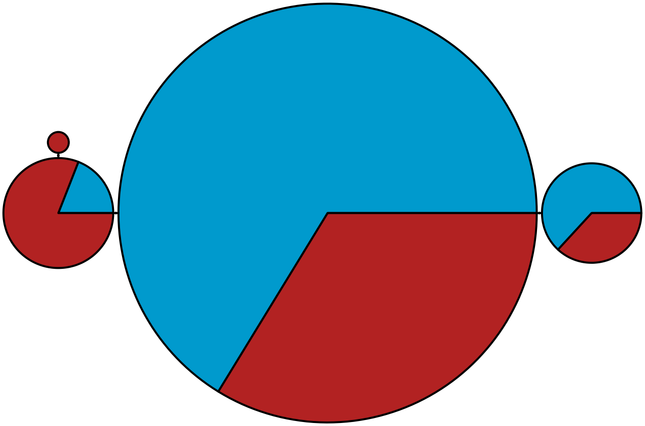

MBPA_000008054 Metazoa

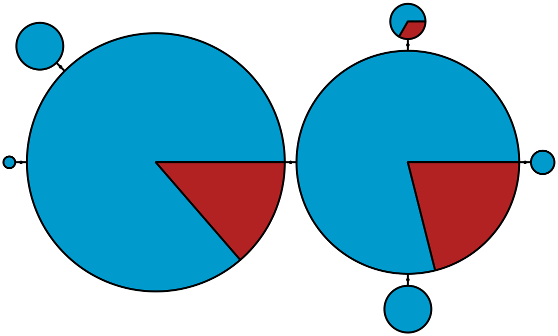

MBPA_000008072

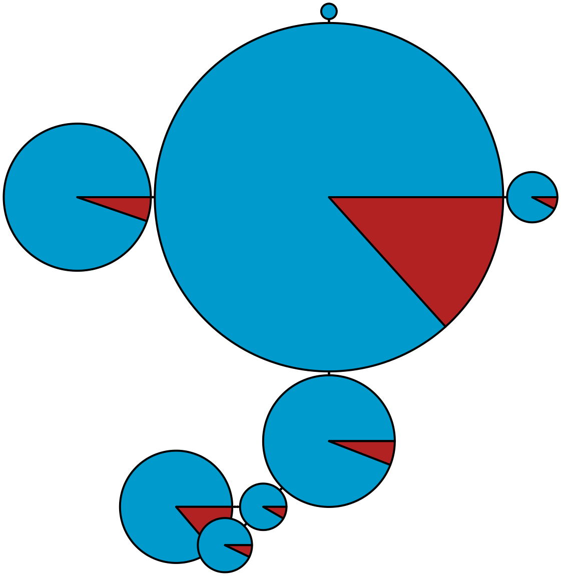

MBPA_000008077

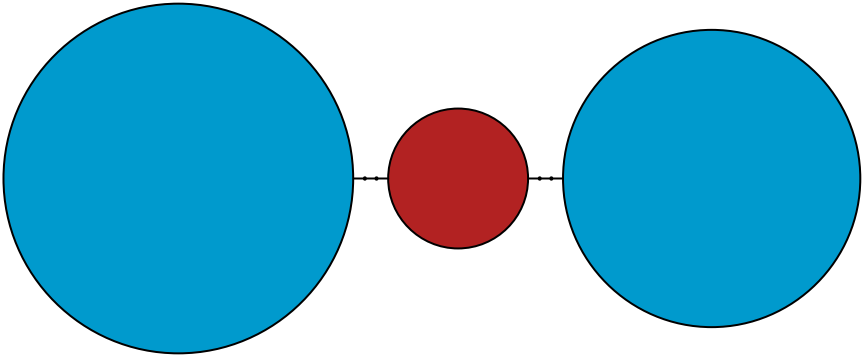

MBPA_000008180 Metazoa

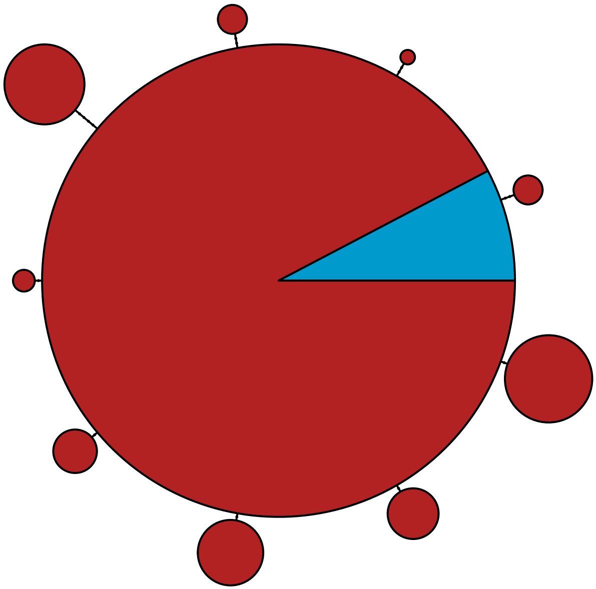

MBPA_000008254 Viridiplantae

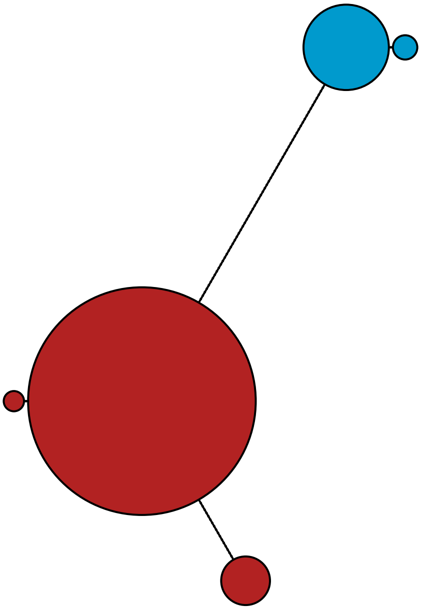

MBPA_000008348 Rhodophyta

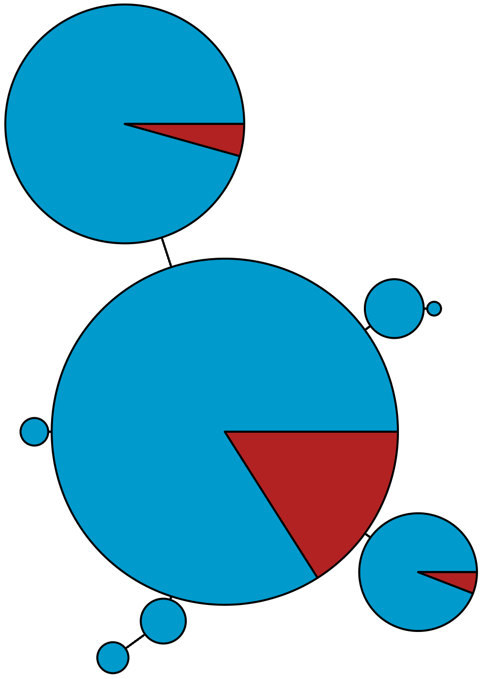

MBPA_000008447 Rhodophyta

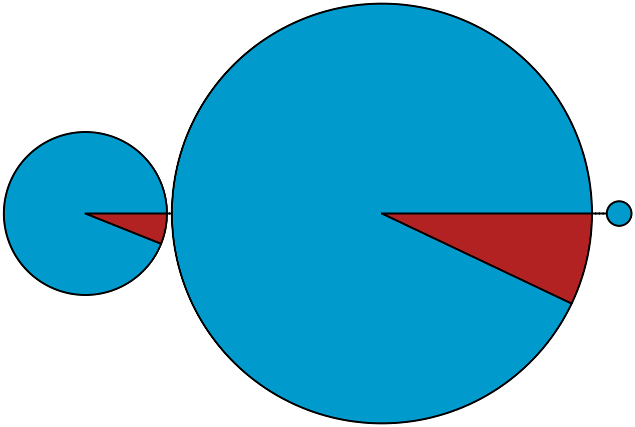

MBPA_000008678 Rhodophyta

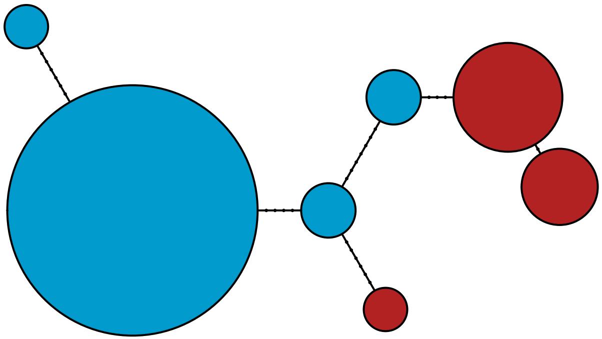

MBPA_000008816 Rhodophyta

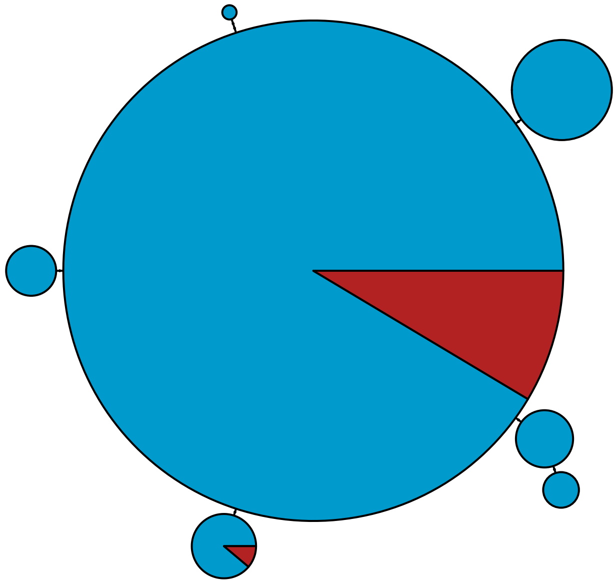

MBPA_000009011 Metazoa

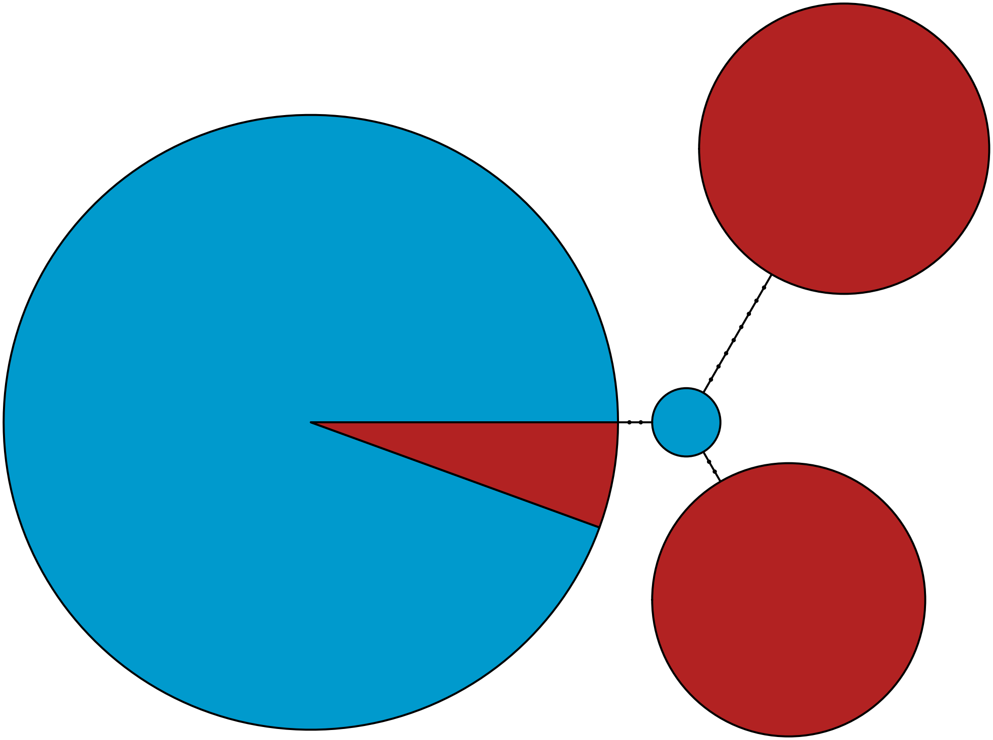

MBPA_000009042 Metazoa

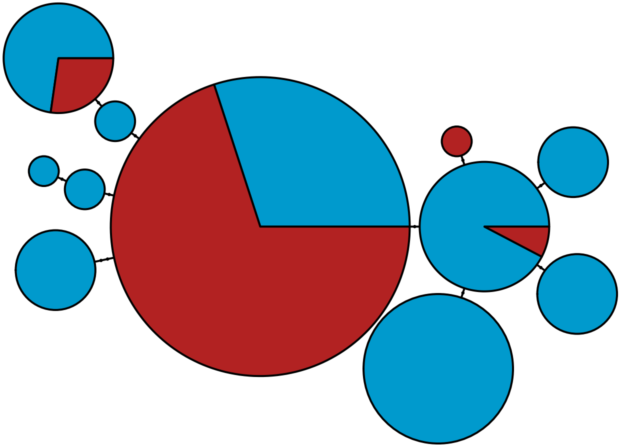

MBPA_000009116 Metazoa

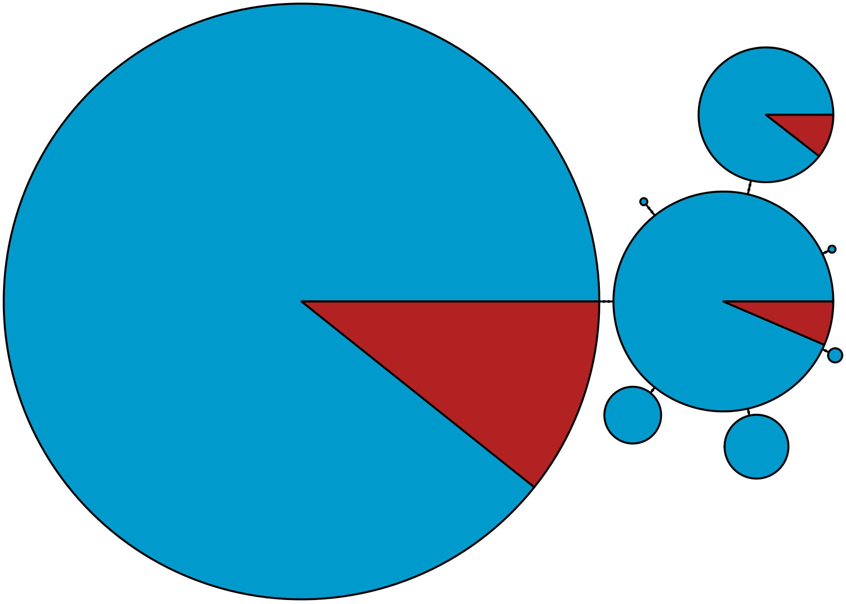

MBPA_000009268

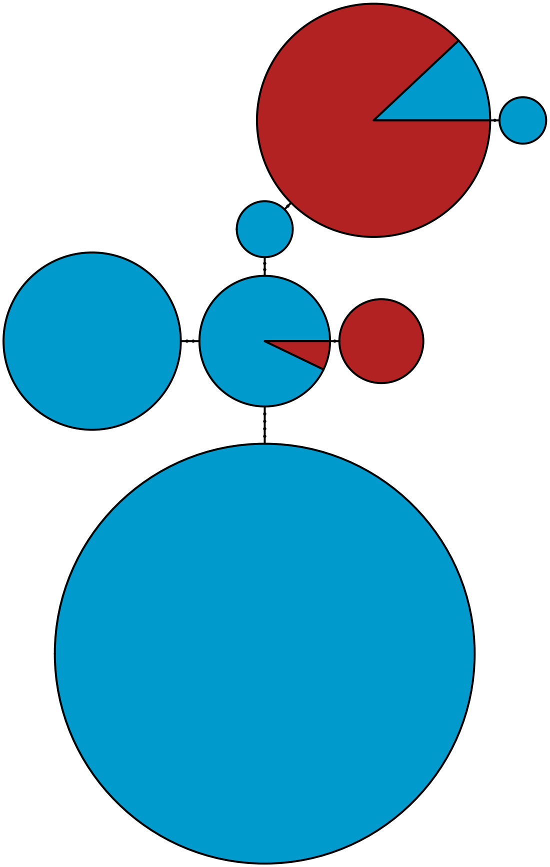

MBPA_000009366 Metazoa

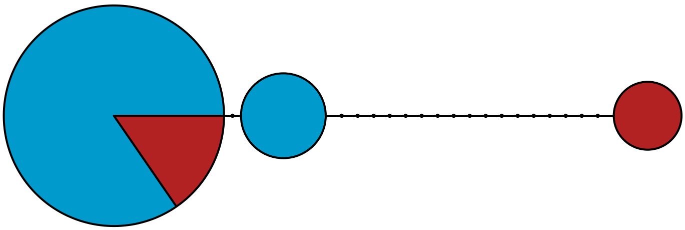

MBPA_000009385 Metazoa

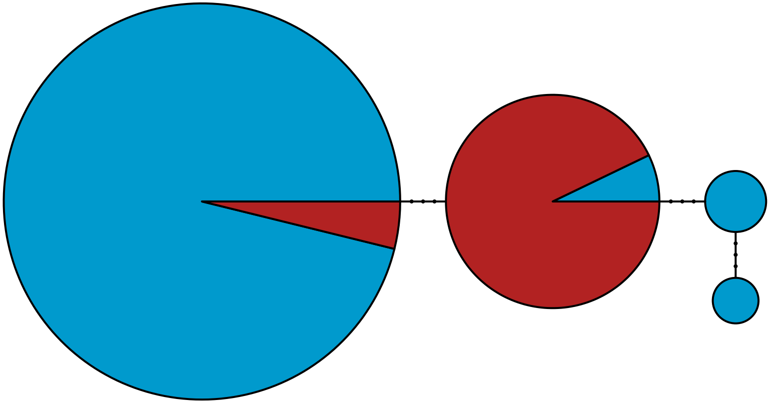

MBPA_000009450 Metazoa

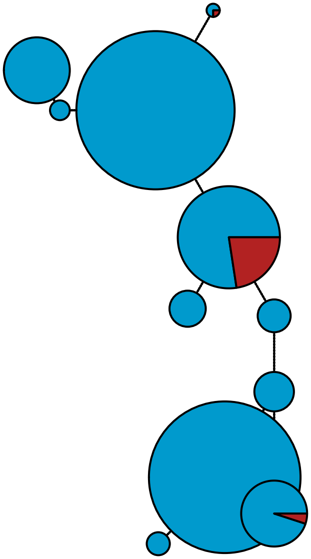

MBPA_000009497 Metazoa

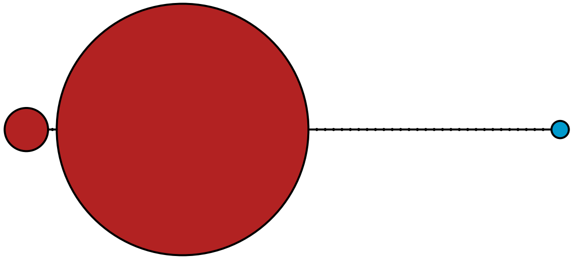

MBPA_000009522 Metazoa

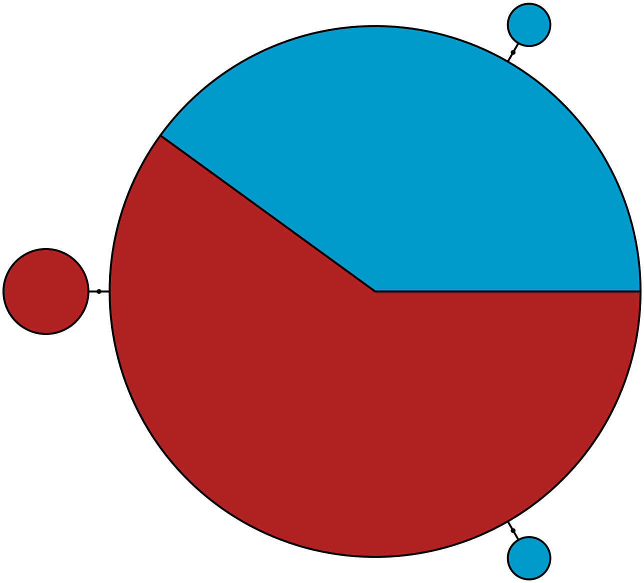

MBPA_000009591 Metazoa

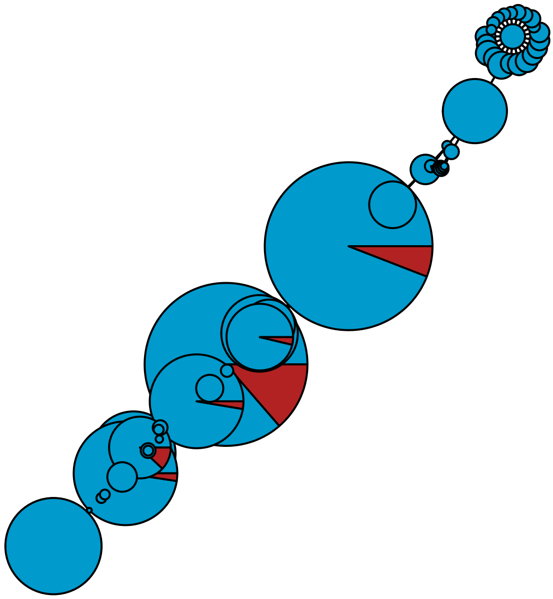

MBPA_000009603 Stramenopiles

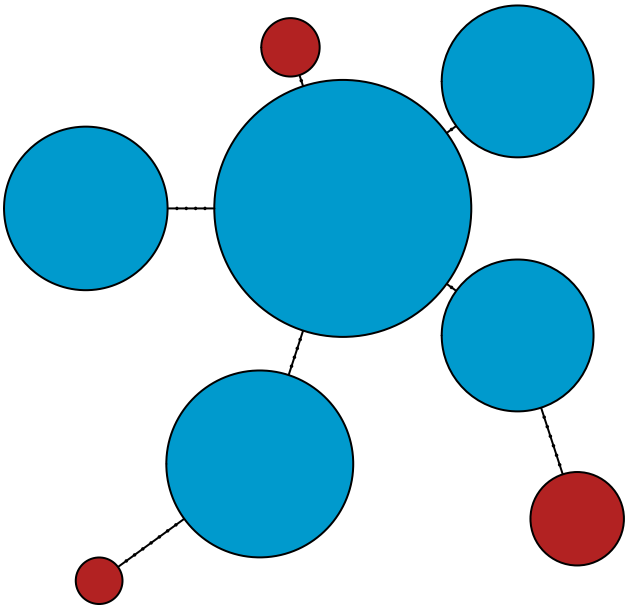

MBPA_000009968 Metazoa

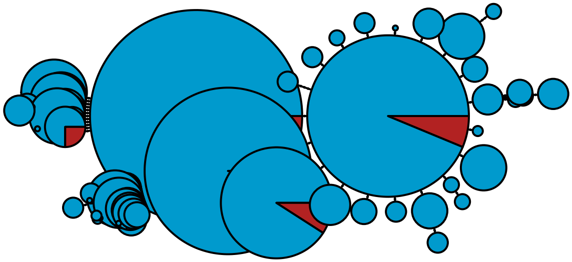

MBPA_000010095 Metazoa

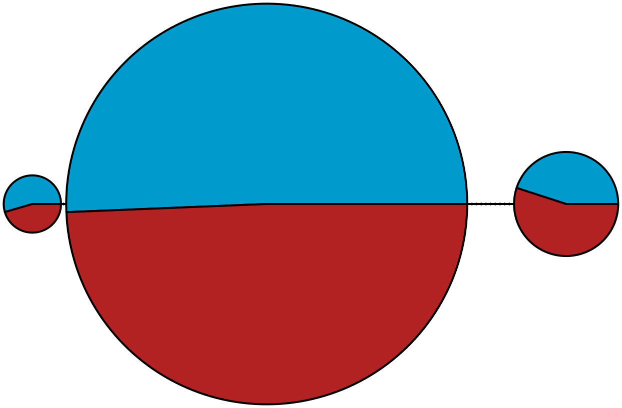

MBPA_000010185 Metazoa

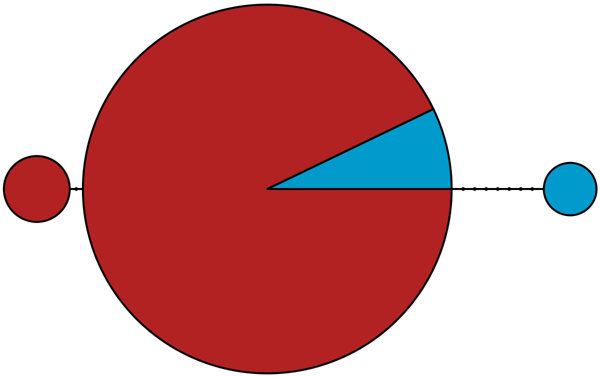

MBPA_000010283 Metazoa

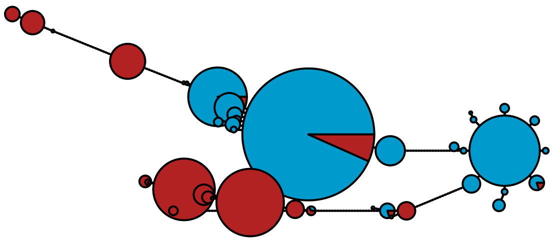

MBPA_000010863 Metazoa

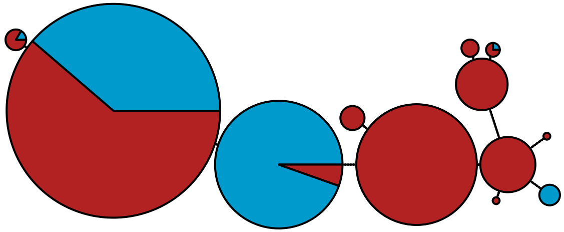

MBPA_000011152 Rhodophyta

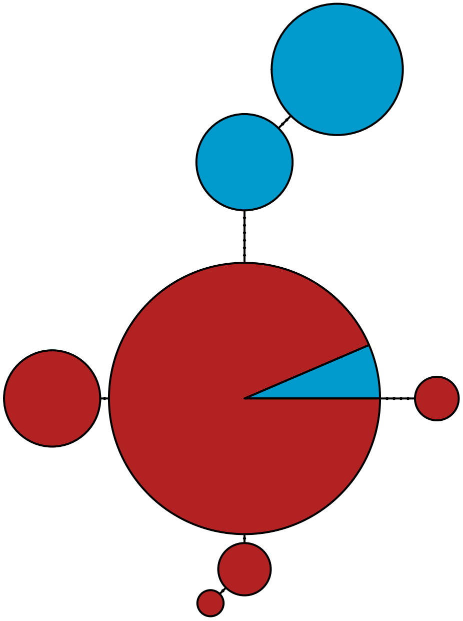

MBPA_000011187 Rhodophyta

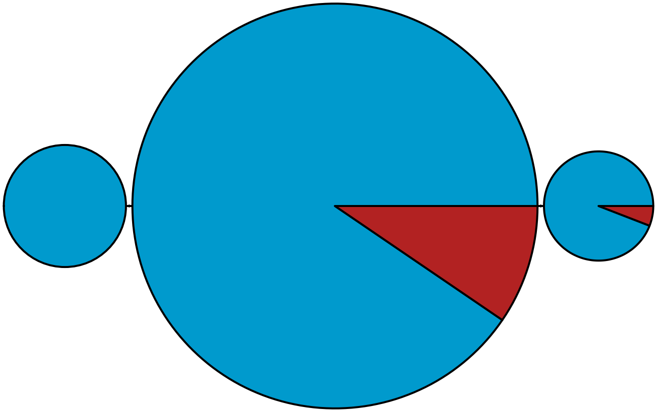

MBPA_000011342 Stramenopiles

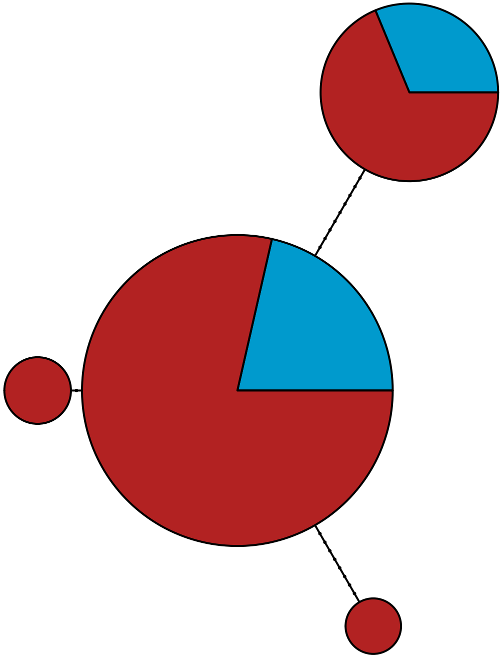

MBPA_000011408 Rhodophyta

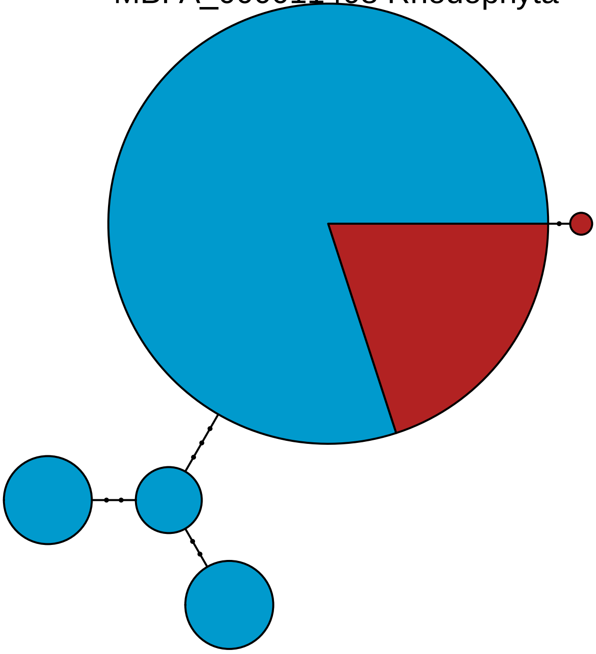

MBPA_000011511 Stramenopiles

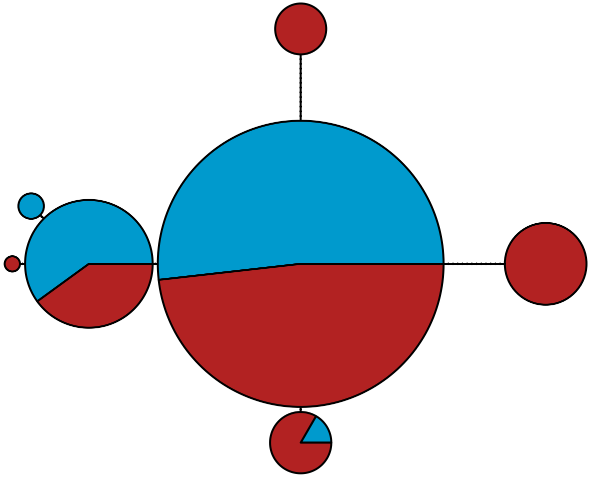

MBPA_000011533 Metazoa

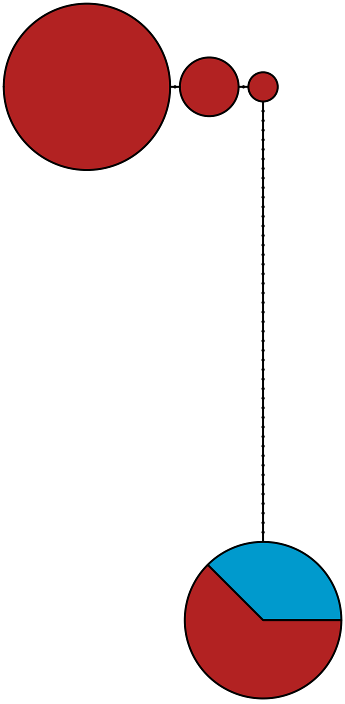

MBPA_000011558 Metazoa

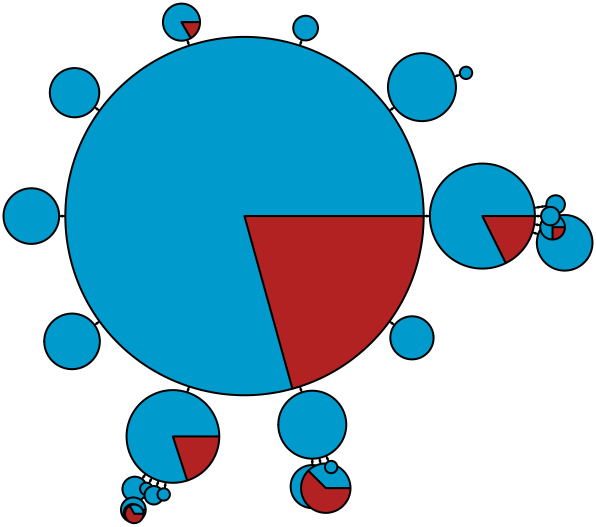

MBPA_000011575 Metazoa

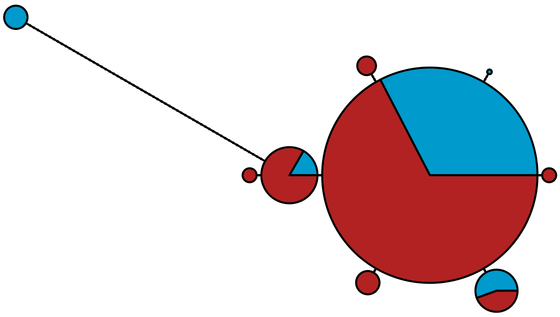

MBPA_000011795

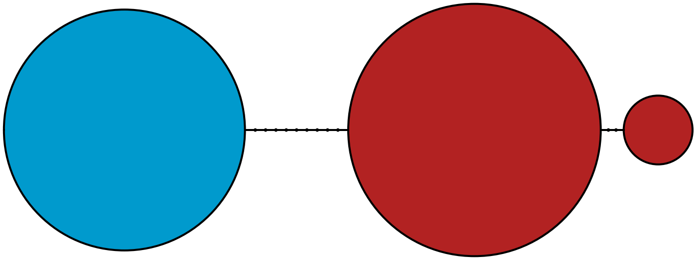

MBPA_000011942 Metazoa

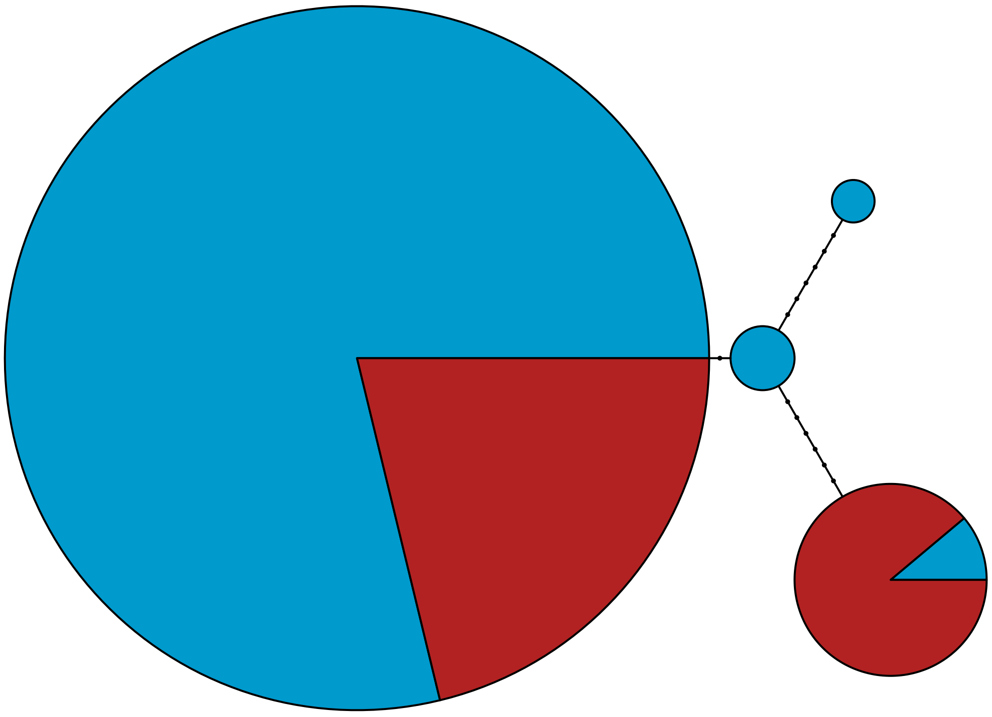

MBPA_000012149 Rhodophyta

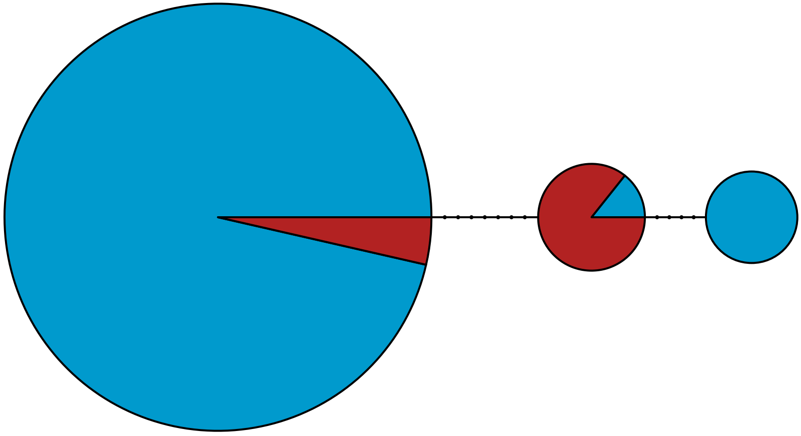

MBPA_000012398 Rhodophyta

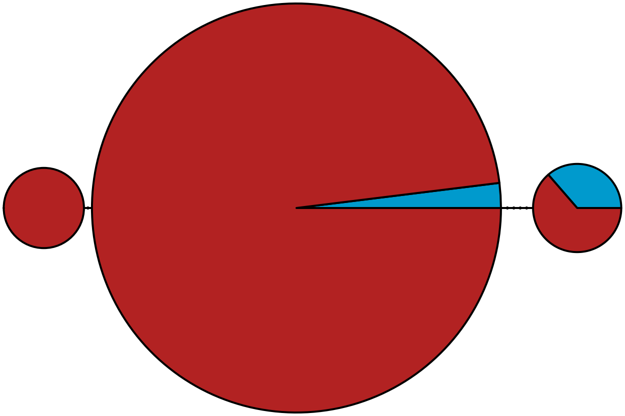

MBPA_000013383 Rhodophyta

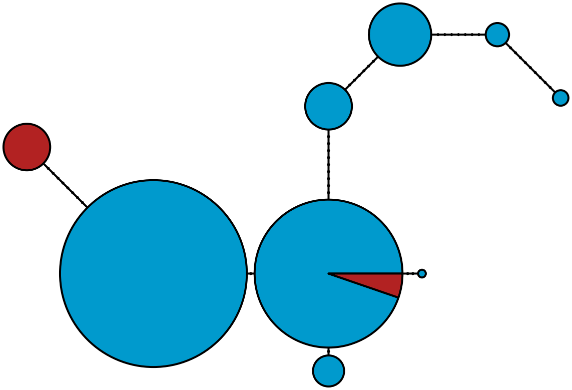

MBPA_000013488 Metazoa

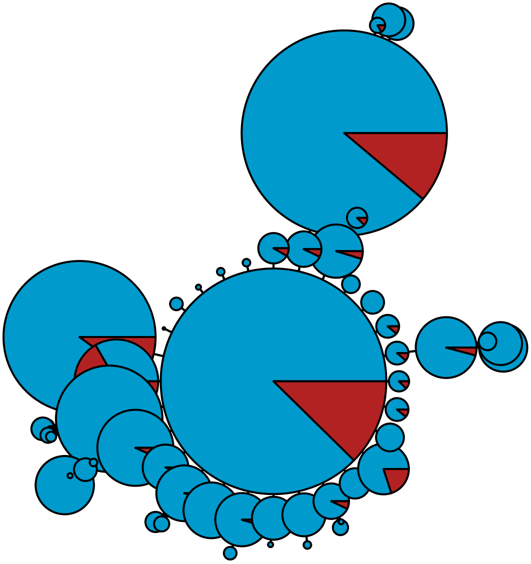

MBPA_000013614 Stramenopiles

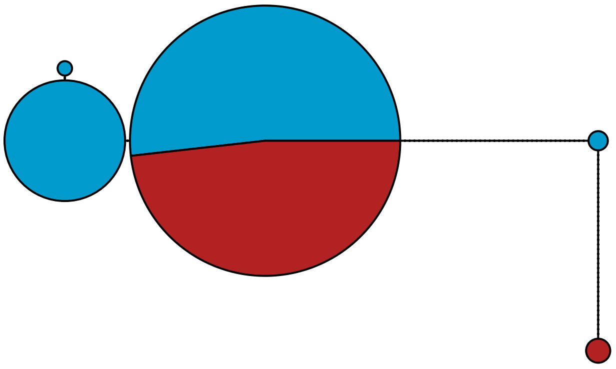

MBPA_000013638 Rhodophyta

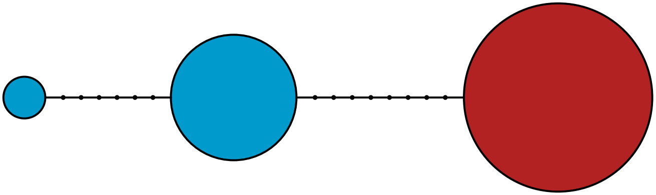

MBPA_000013680 Metazoa

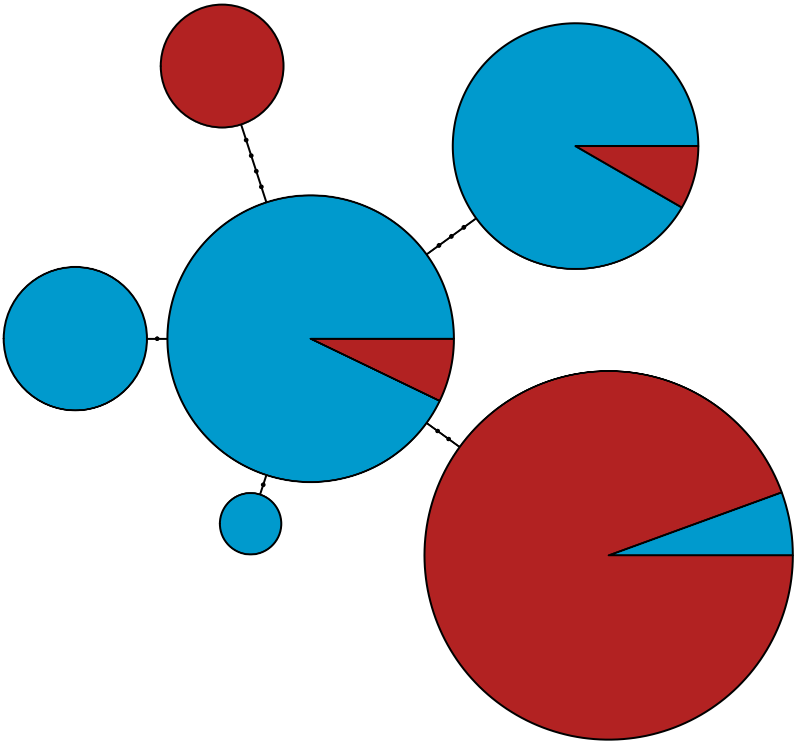

MBPA_000014041 Rhodophyta

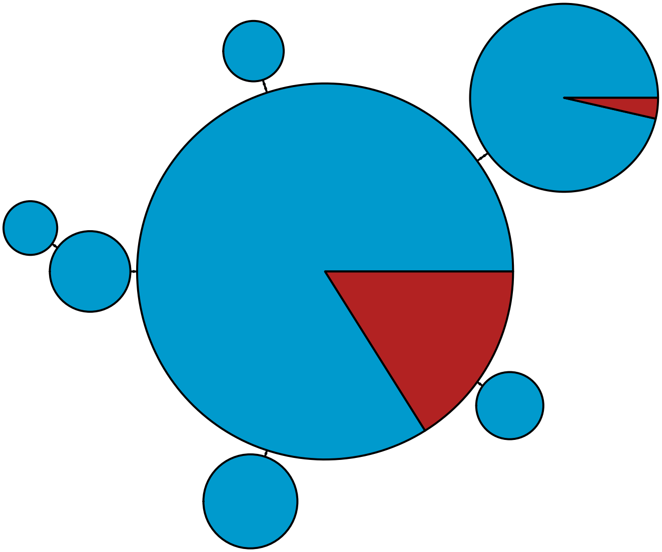

MBPA_000014171 Metazoa

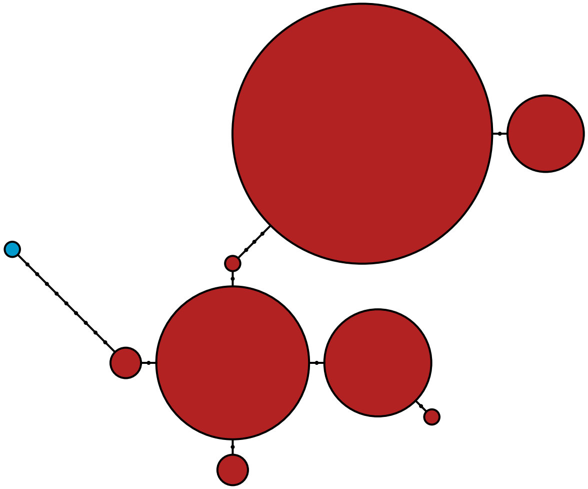

MBPA_000014317 Rhodophyta

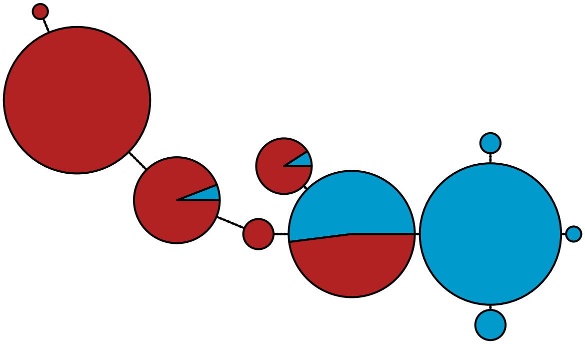

MBPA_000014353 Metazoa

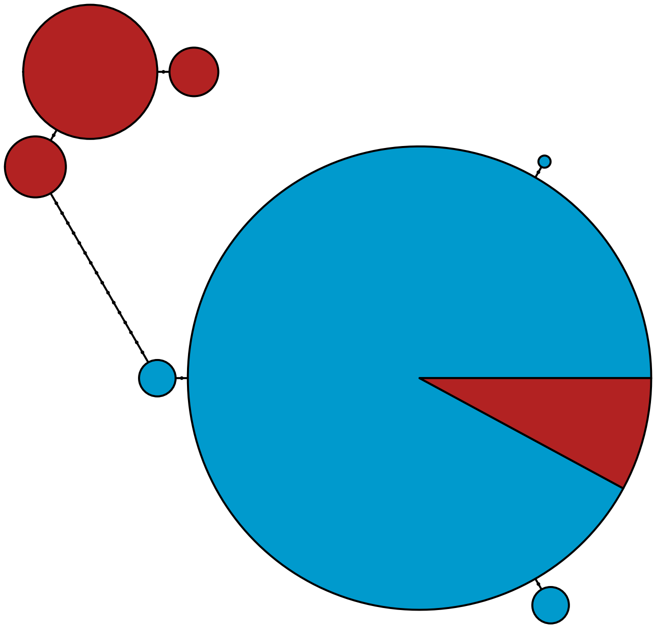

MBPA_000014365 Metazoa

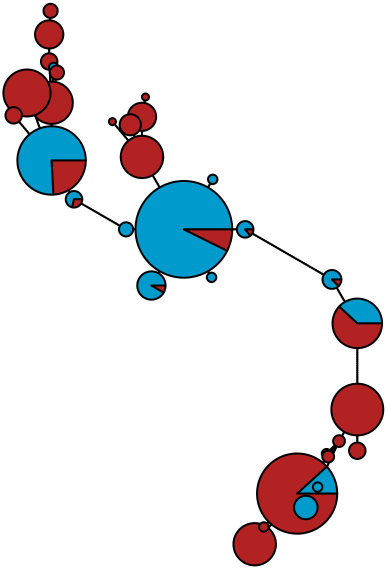

MBPA_000014569

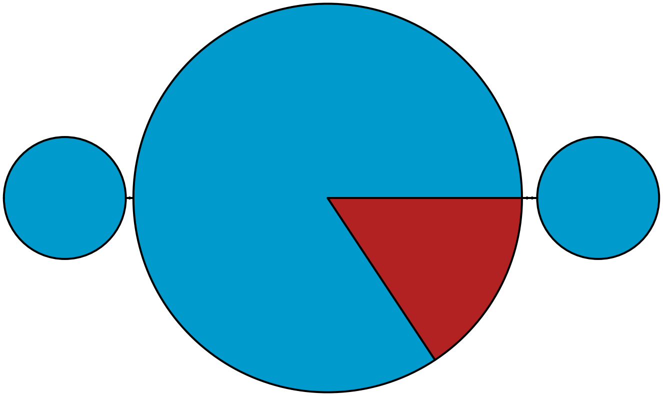

MBPA_000014641

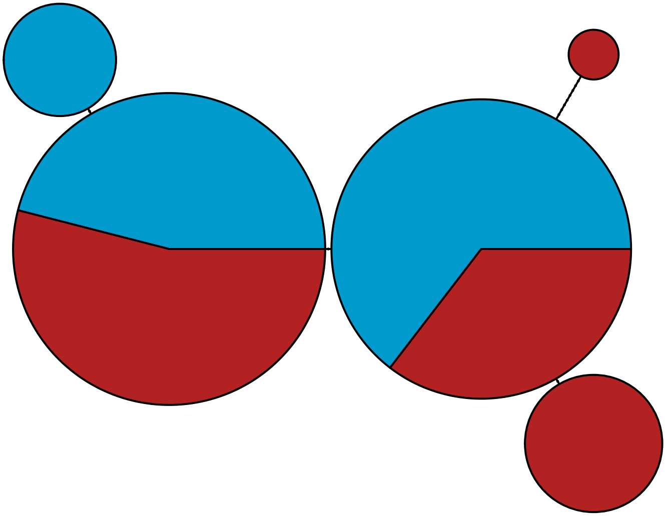

MBPA_000015316

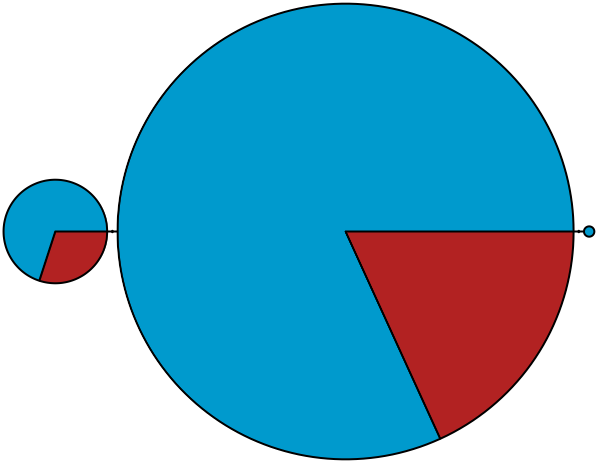

MBPA_000015396 Metazoa

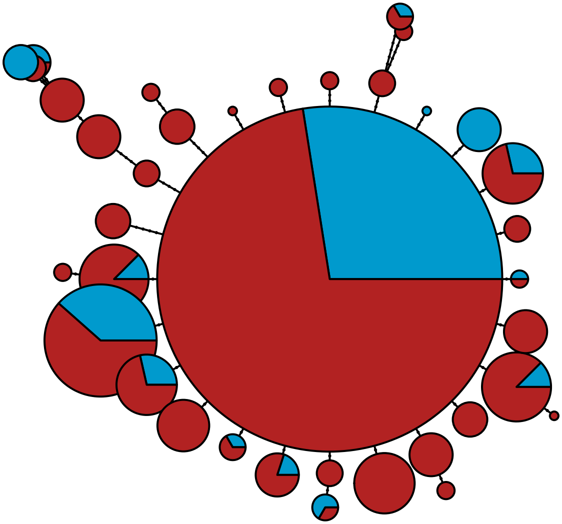

MBPA_000015741 Metazoa

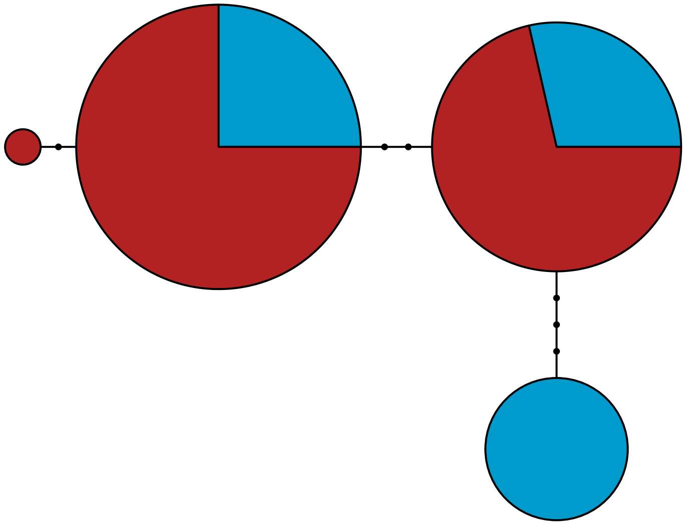

MBPA_000015751 Metazoa

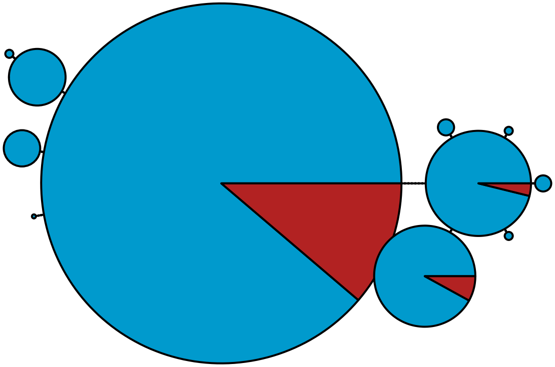

MBPA_000015964 Rhodophyta

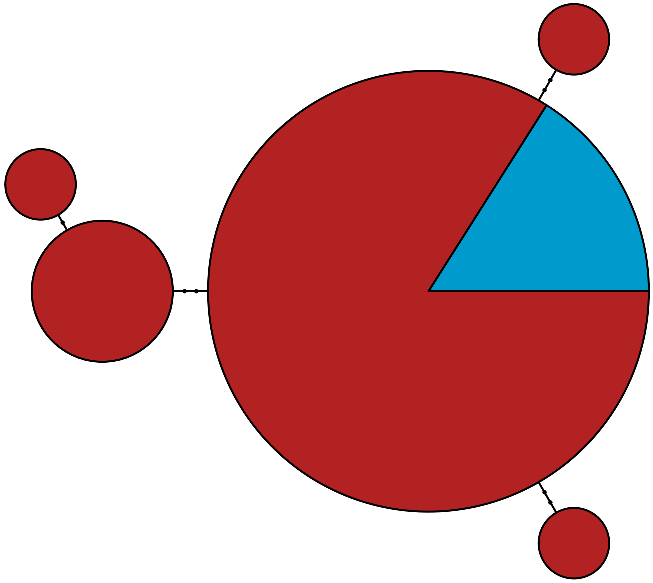

MBPA_000016543 Rhodophyta

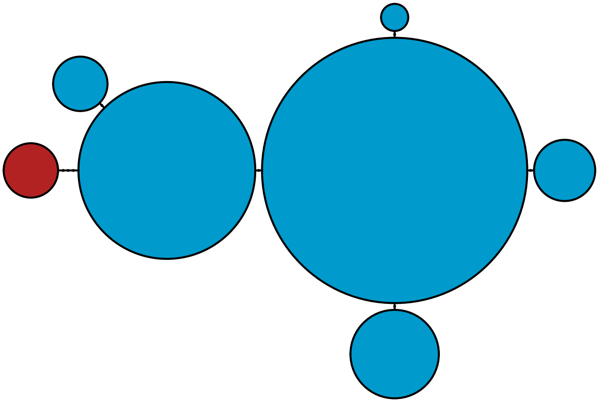

MBPA_000017127 Metazoa

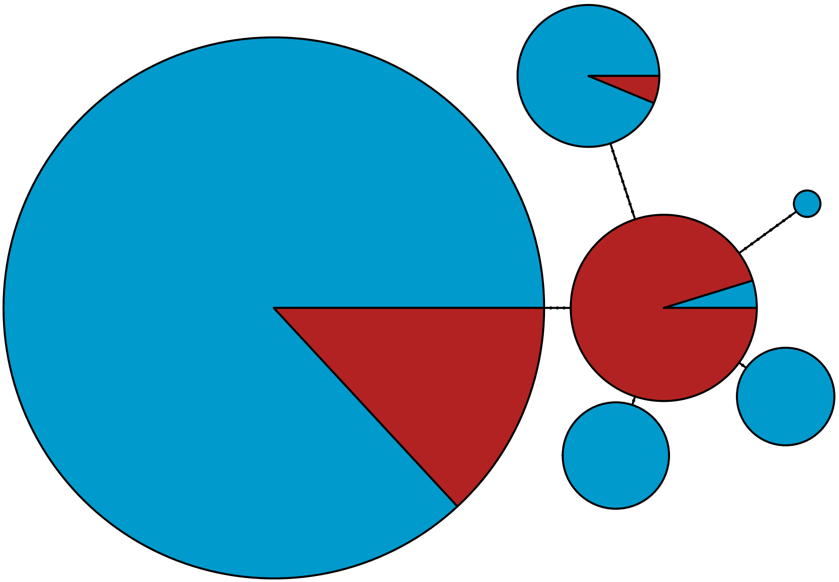

### MBPA_000017339 Rhodophyta

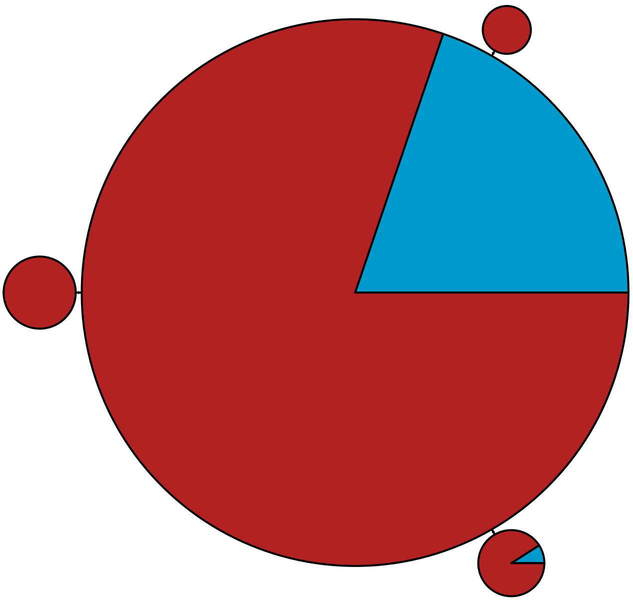

MBPA_000017880

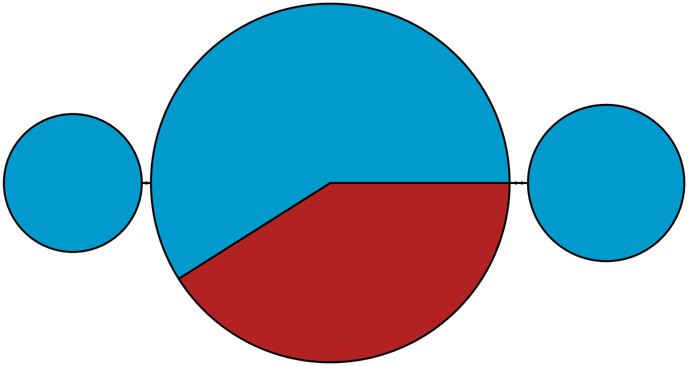

MBPA_000018047 Metazoa

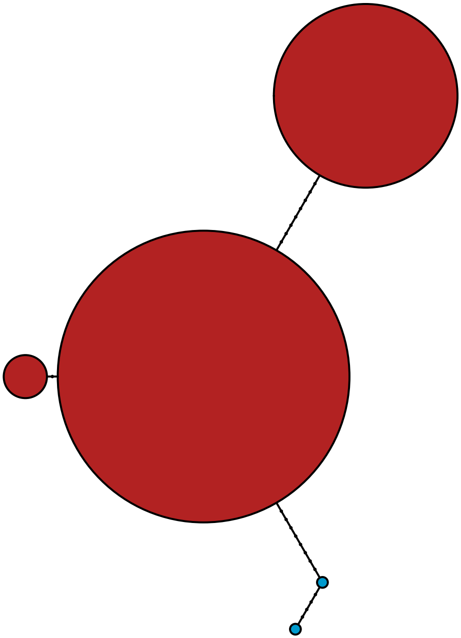

MBPA_000018125 Metazoa

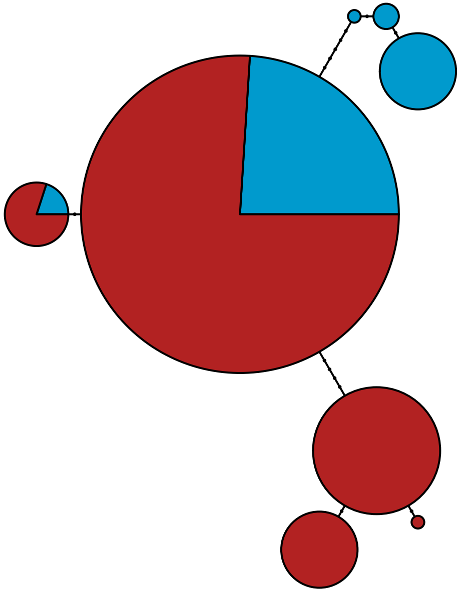

MBPA_000018321 Rhodophyta

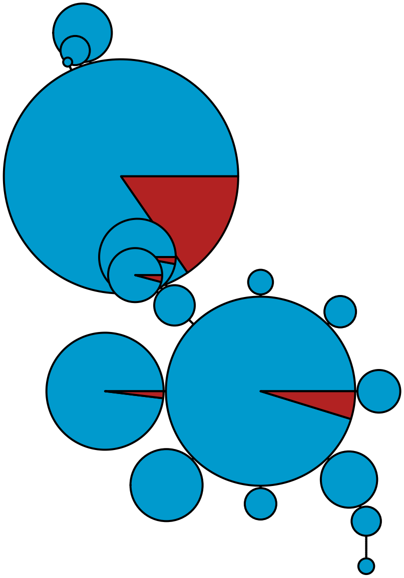

MBPA_000019194 Metazoa

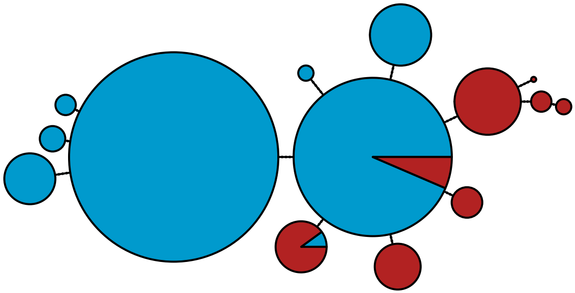

MBPA_000019499 Stramenopiles

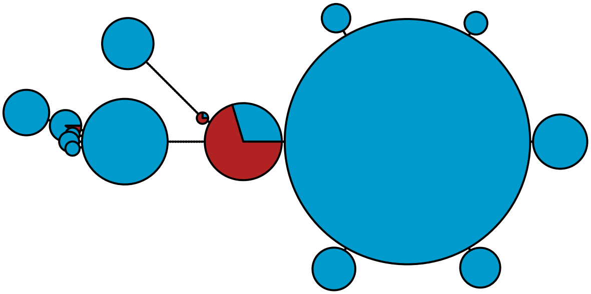

MBPA_000019575 Stramenopiles

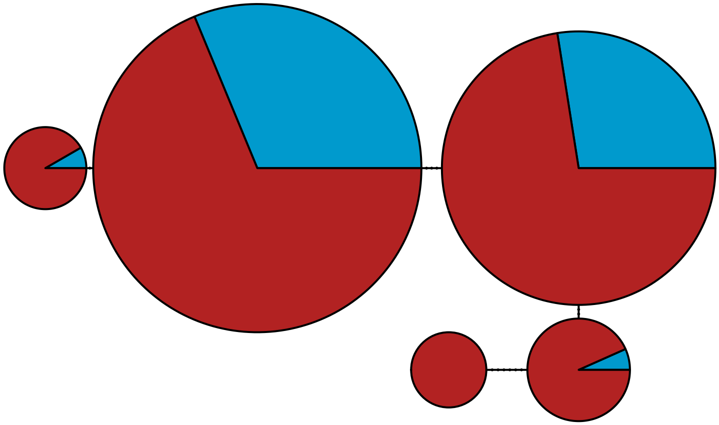

MBPA_000019614 Metazoa

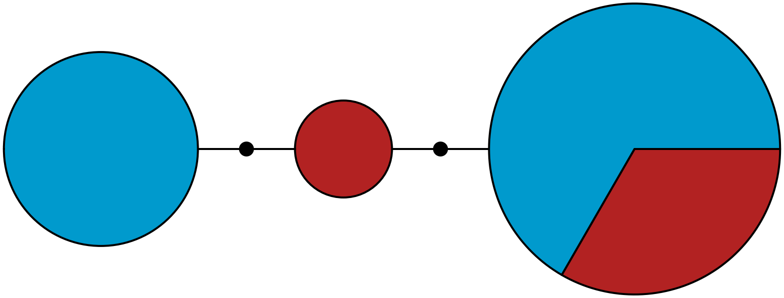

MBPA_000019805 Metazoa

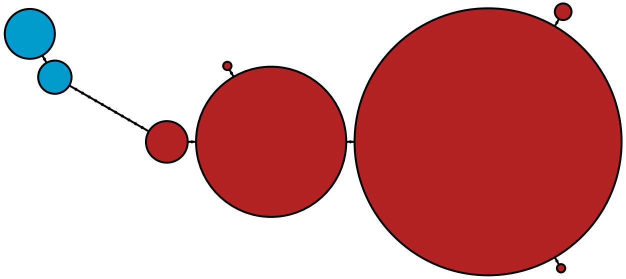

MBPA_000019843 Rhodophyta

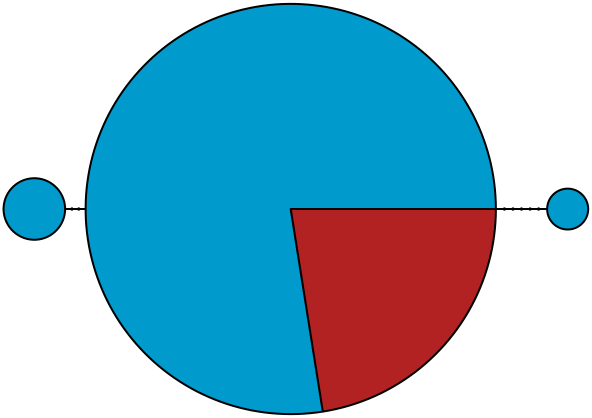

MBPA_000019963 Metazoa

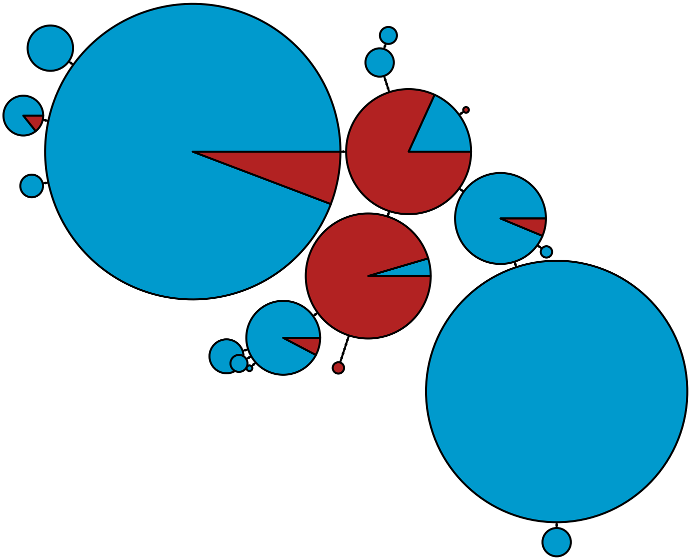

MBPA_000020026 Stramenopiles

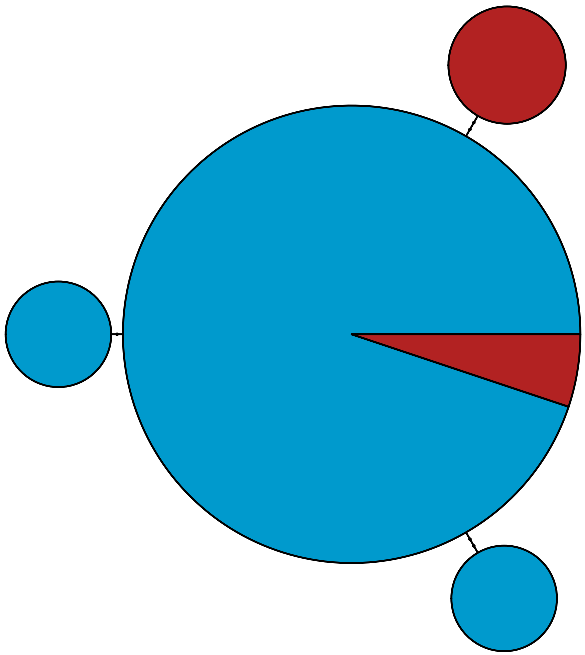

MBPA_000020370

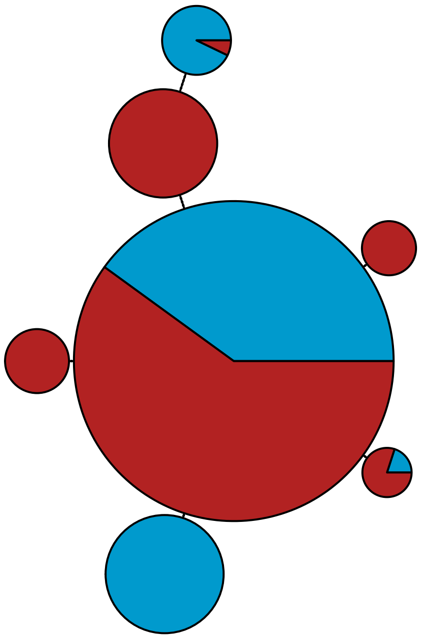

MBPA_000020384 Stramenopiles

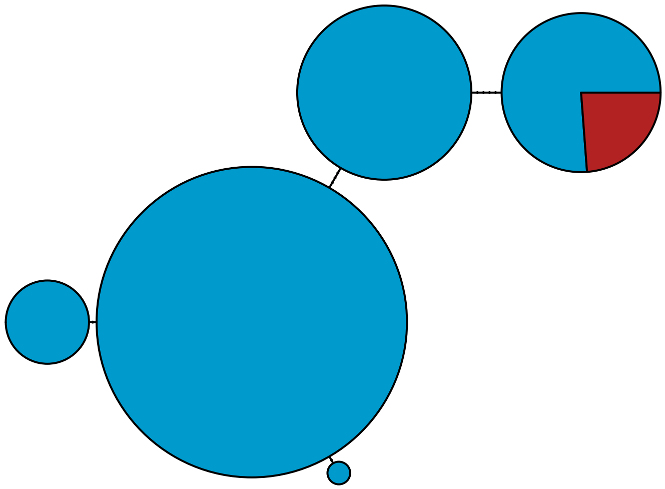

MBPA_000020965 Stramenopiles

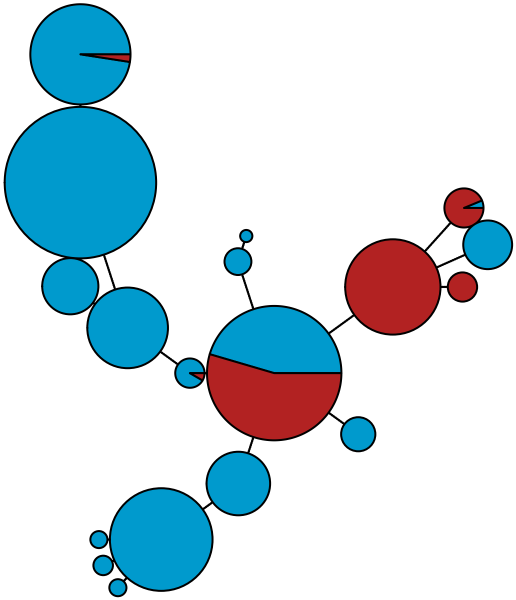

MBPA_000021047 Metazoa

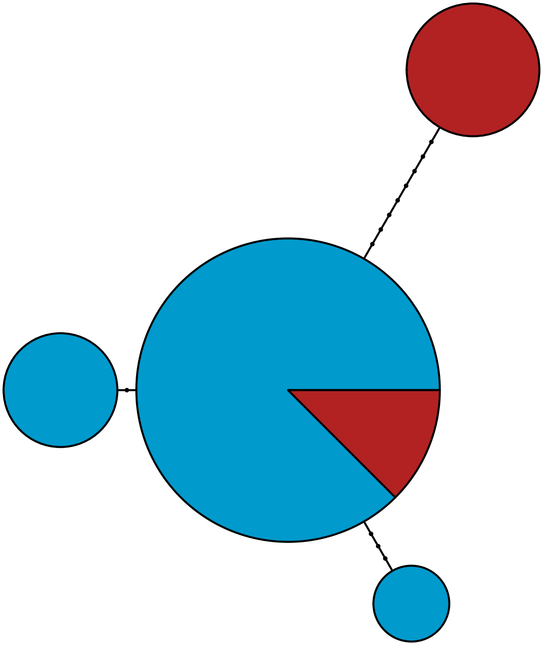

MBPA_000021173 Metazoa

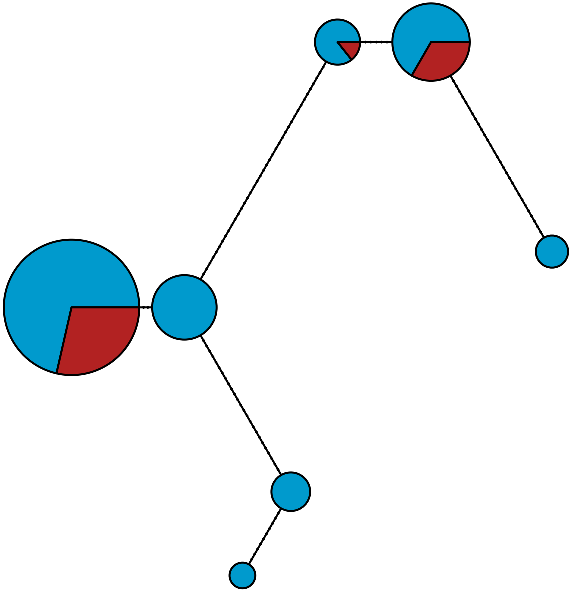

MBPA_000021660 Metazoa

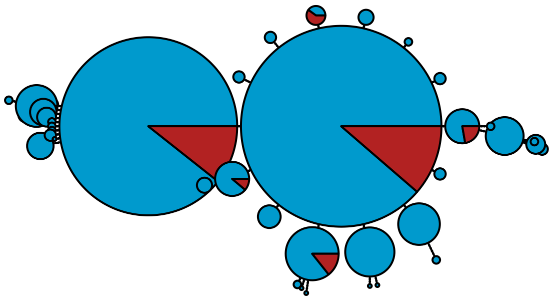

MBPA_000022983 Rhodophyta

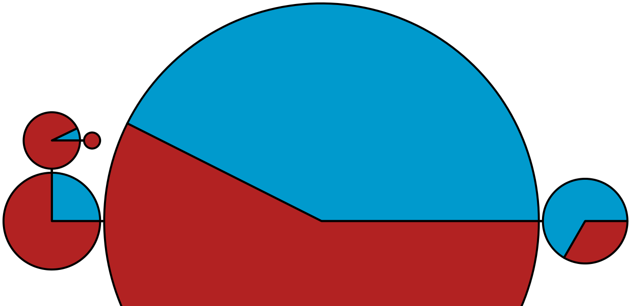

MBPA_000023160 Metazoa

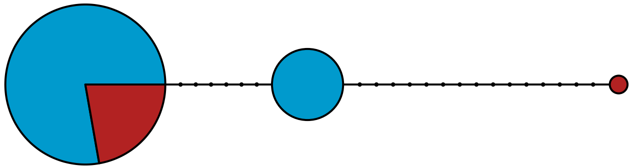

MBPA_000023304 Stramenopiles

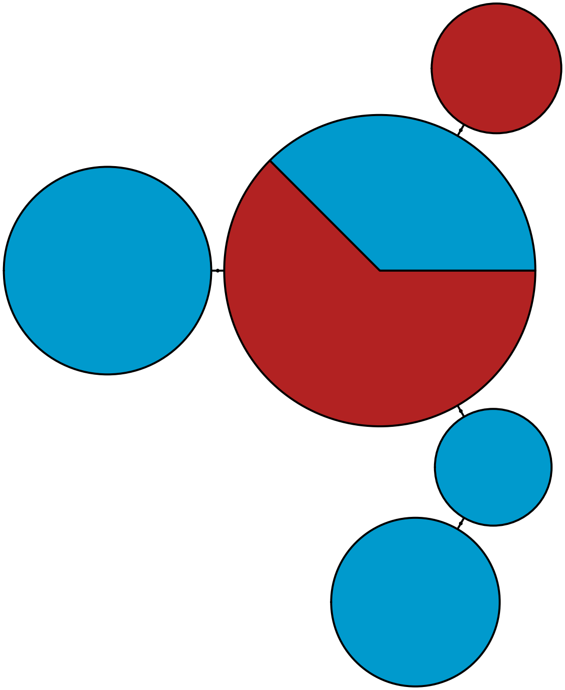

MBPA_000023526 Metazoa

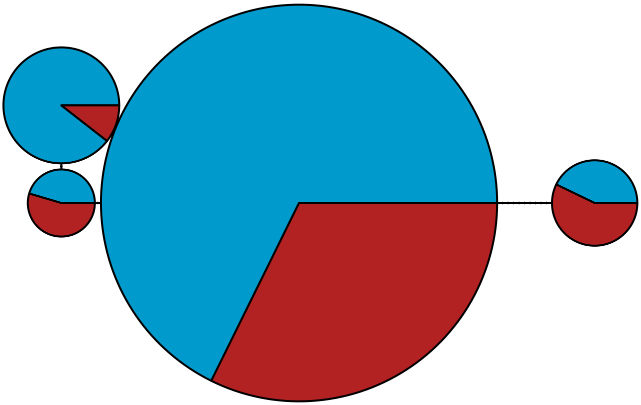

MBPA_000023558 Metazoa

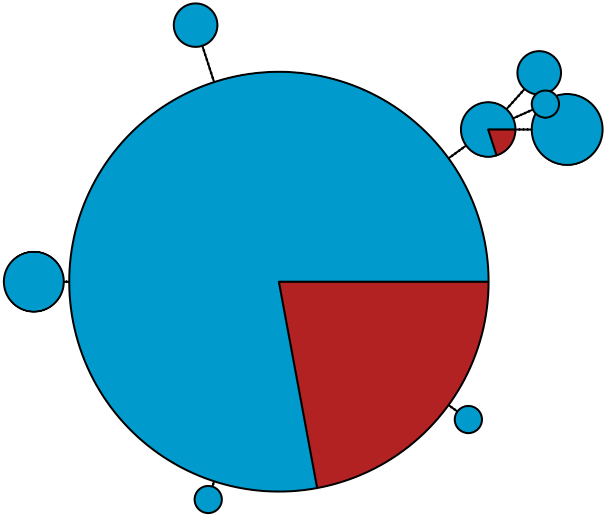

MBPA_000023808 Rhodophyta

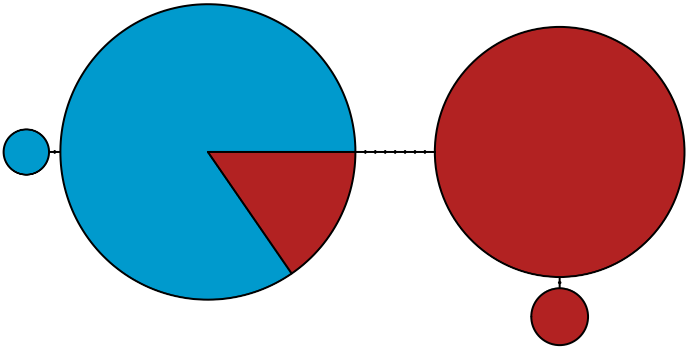

MBPA_000024011 Metazoa

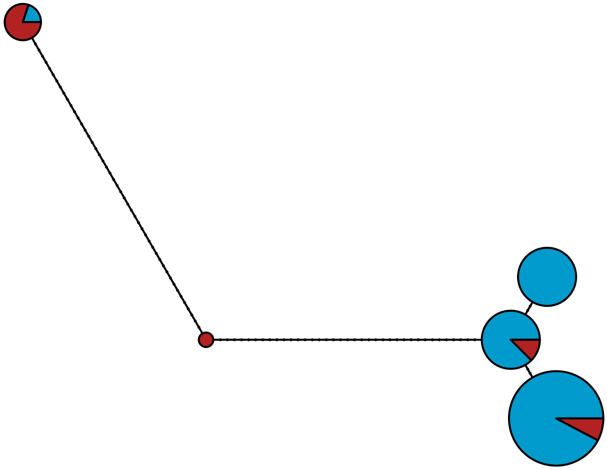

MBPA_000024329 Lobosa

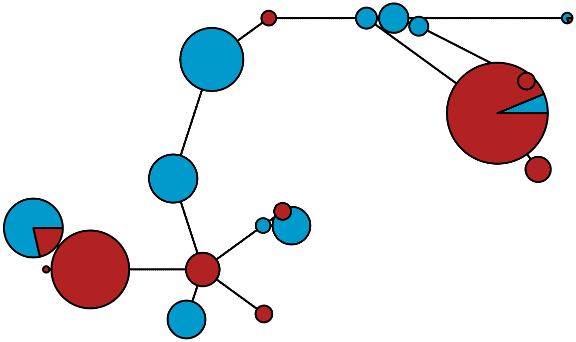

MBPA_000024401 Metazoa

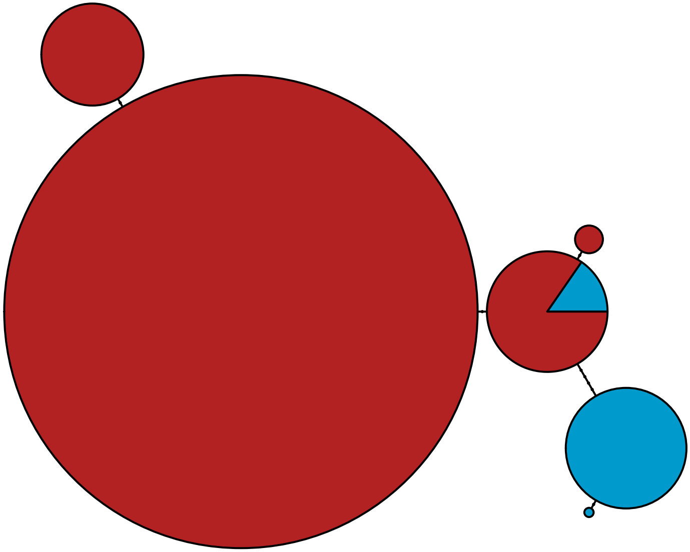

MBPA_000024738 Stramenopiles

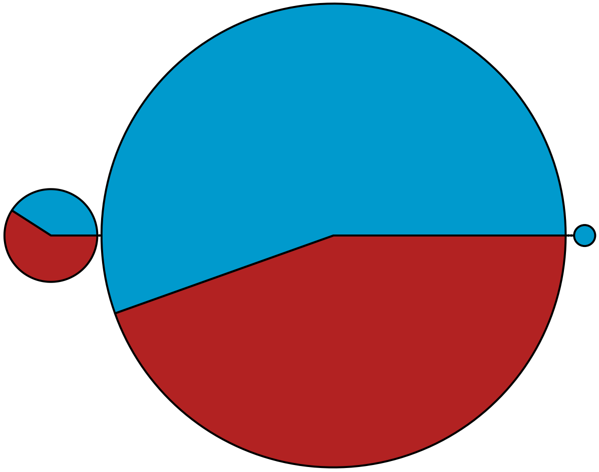

MBPA_000025120 Metazoa

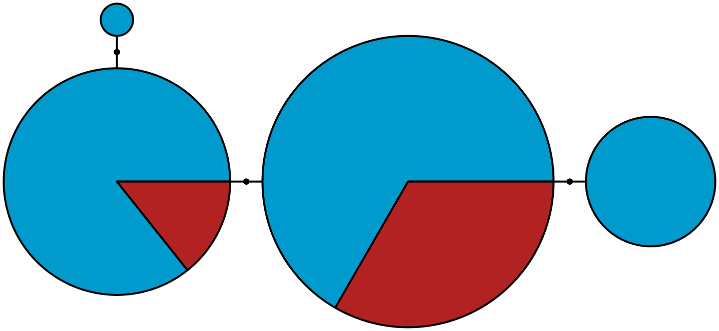

MBPA_000025150 Metazoa

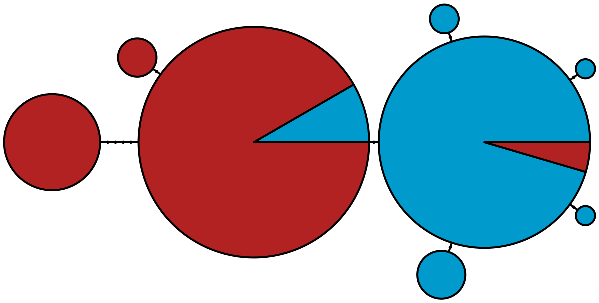

MBPA_000025366 Metazoa

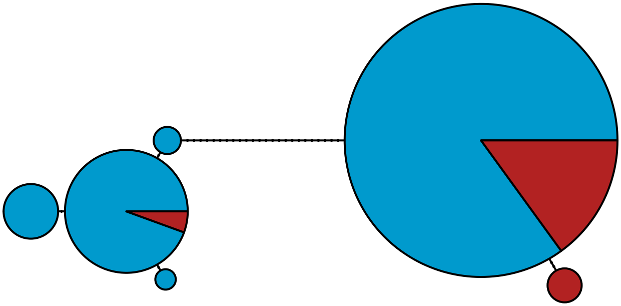

MBPA_000026133

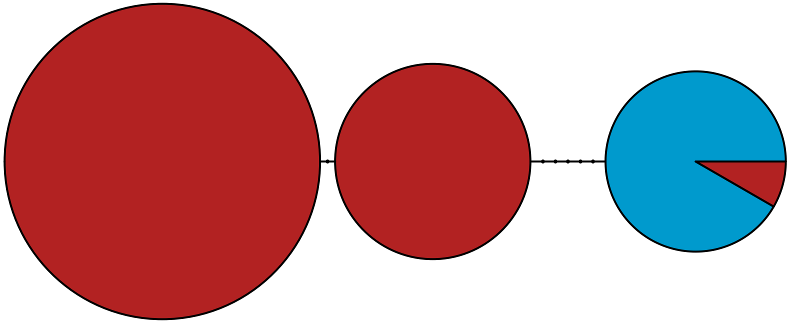

MBPA_000026711

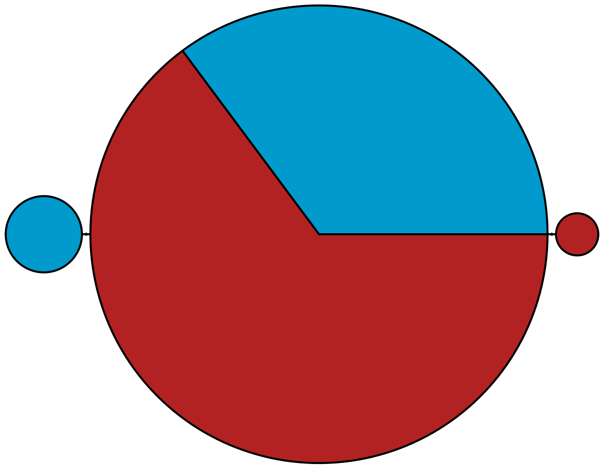

MBPA_000026716 Stramenopiles

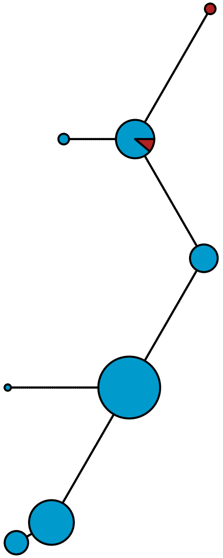

### MBPA_000027799 Stramenopiles

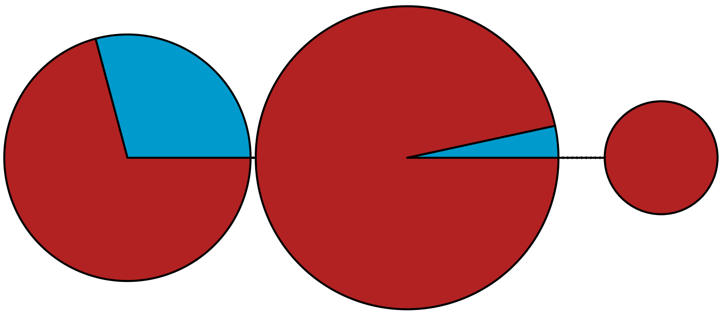

MBPA_000028231 Stramenopiles

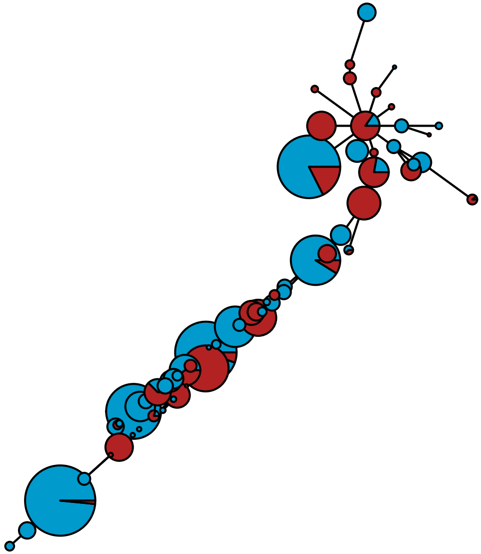

MBPA_000028274 Metazoa

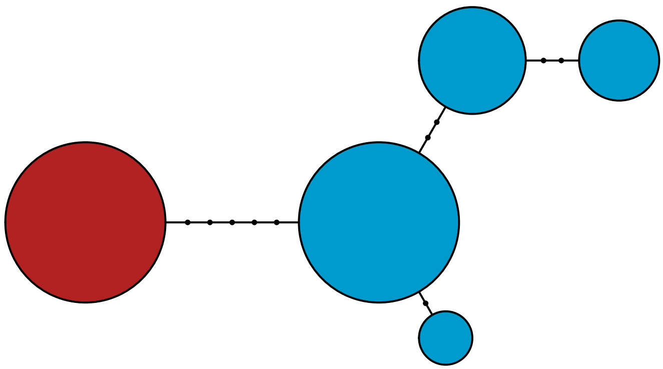

MBPA_000028539 Metazoa

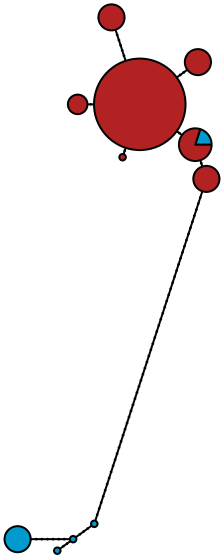

MBPA_000029048 Metazoa

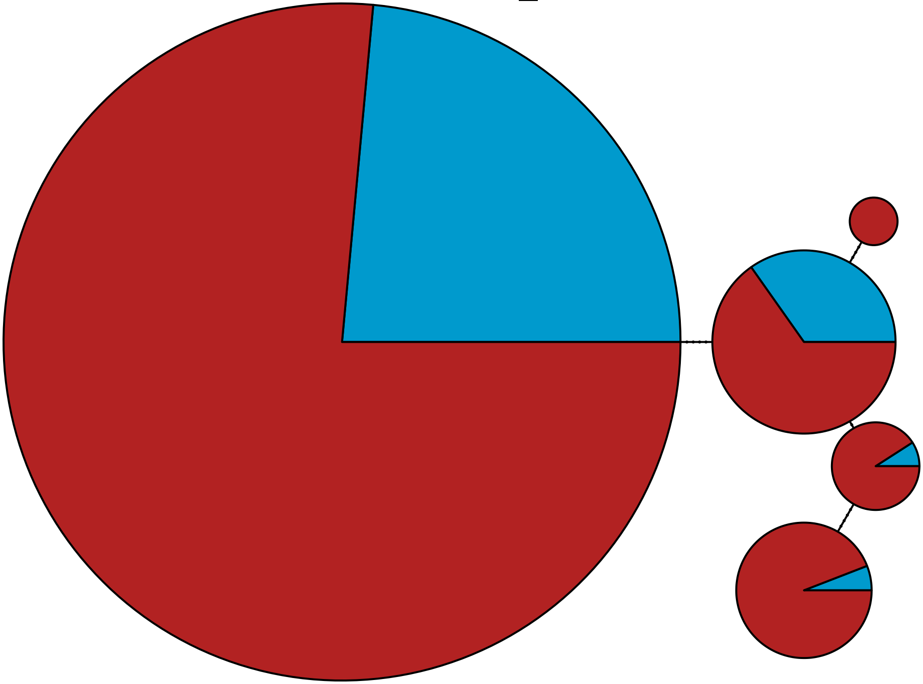

MBPA_000029195 Stramenopiles

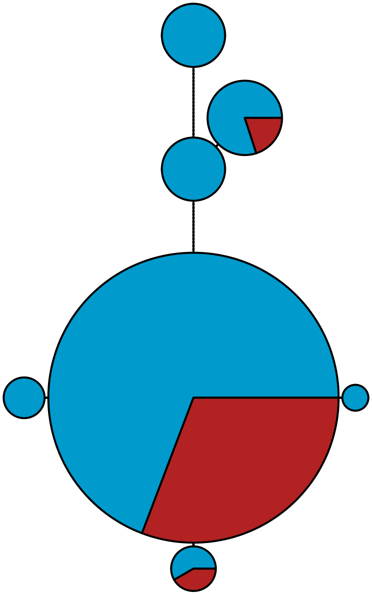

MBPA_000029552 Metazoa

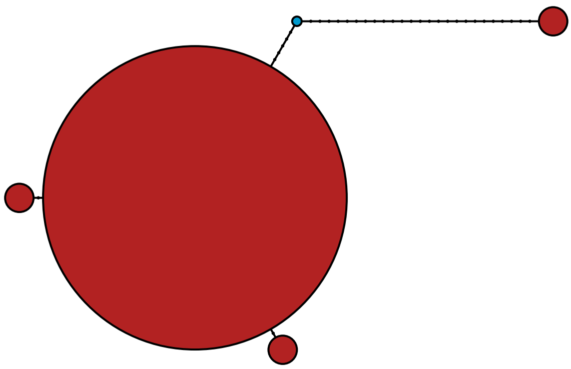

MBPA_000031064

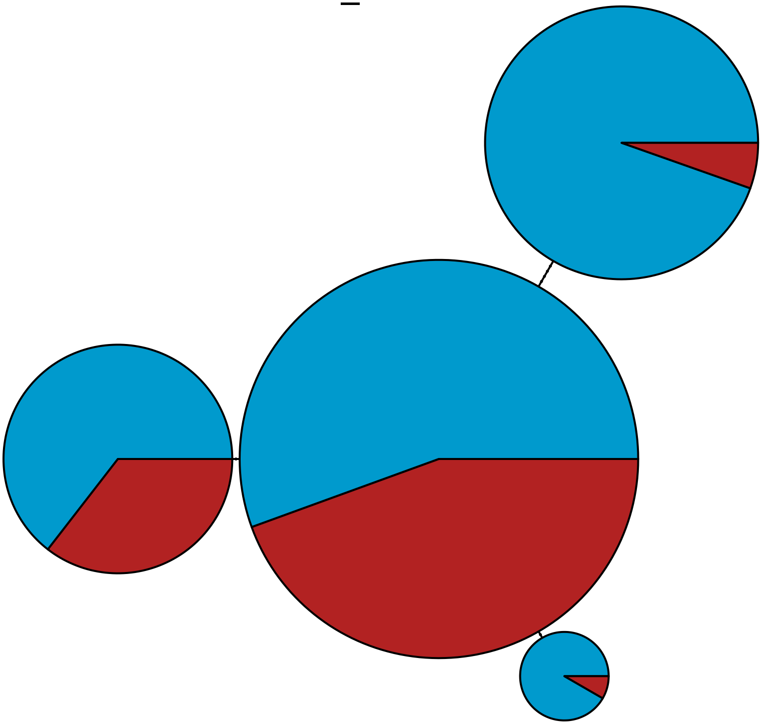

MBPA_000031322 Stramenopiles

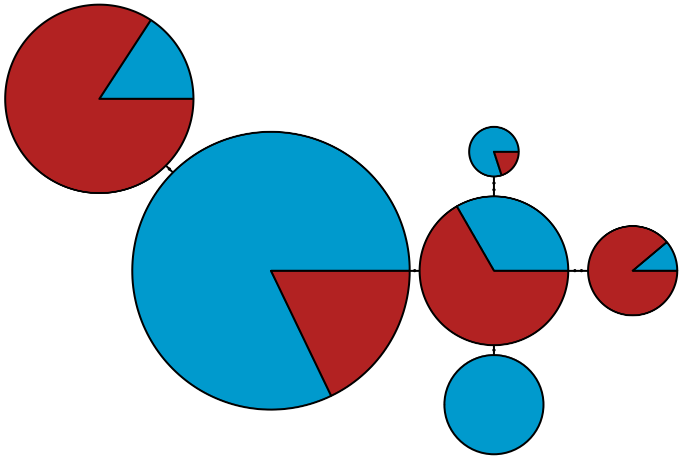

MBPA_000031574 Stramenopiles

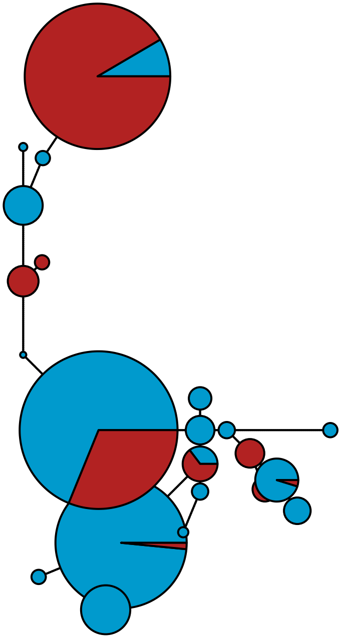

MBPA_000032954 Stramenopiles

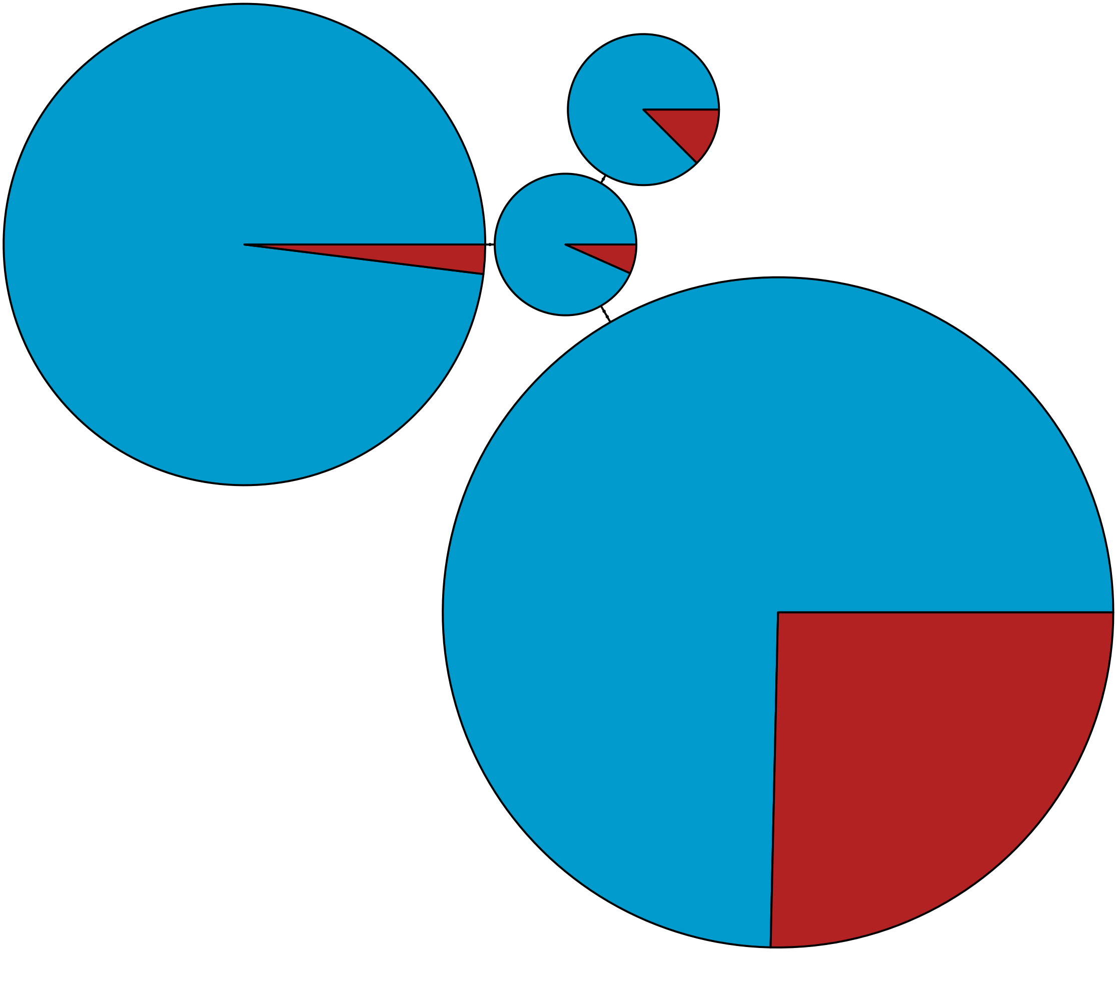

MBPA_000033226 Metazoa

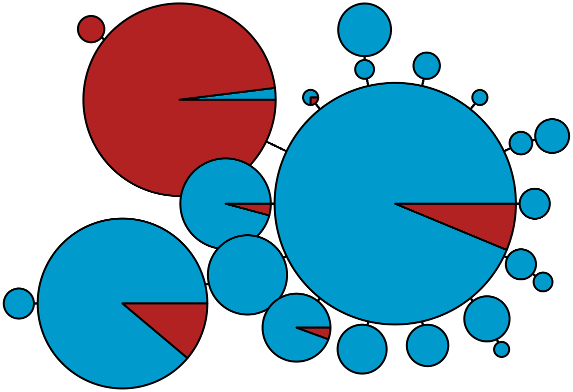

MBPA_000033962 Apusozoa

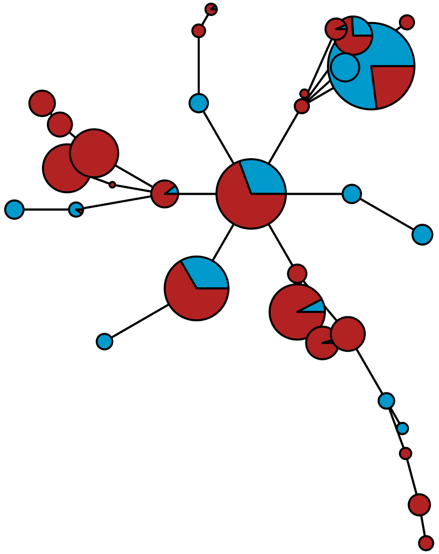

MBPA_000035627 Metazoa

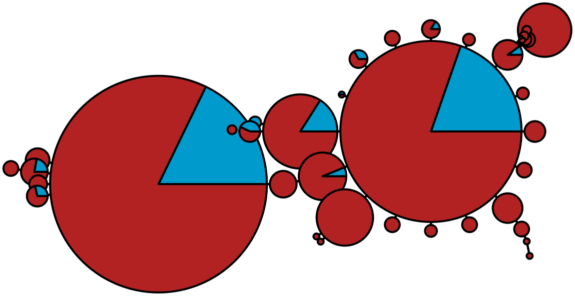

MBPA_000036272 Metazoa

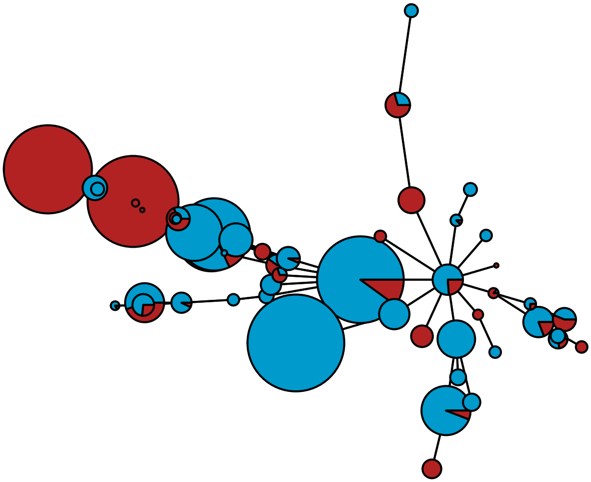

MBPA_000036534 Rhodophyta

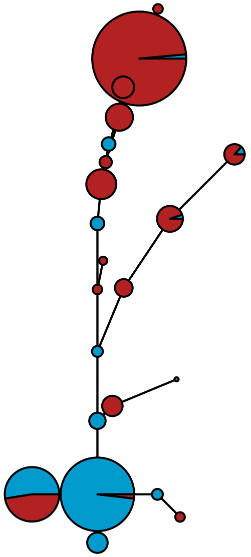

MBPA_000036793

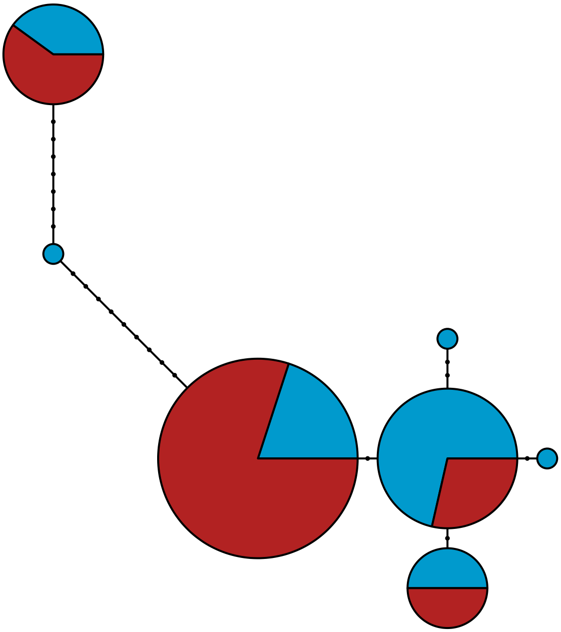

MBPA_000037107 Rhodophyta

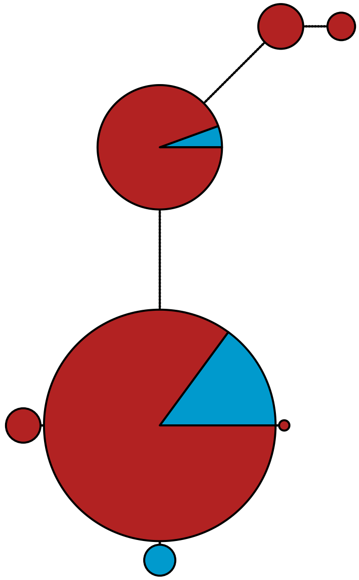

MBPA_000037644 Metazoa

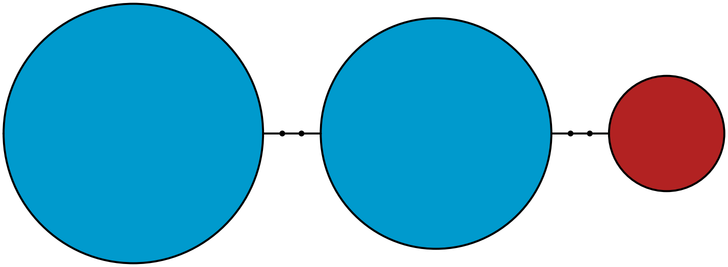

MBPA_000037724 Metazoa

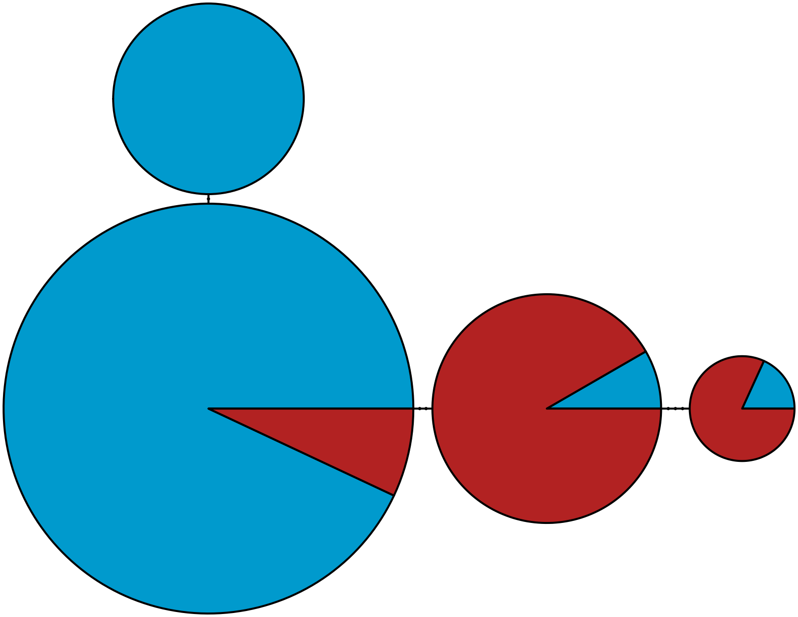

MBPA_000037738 Rhodophyta

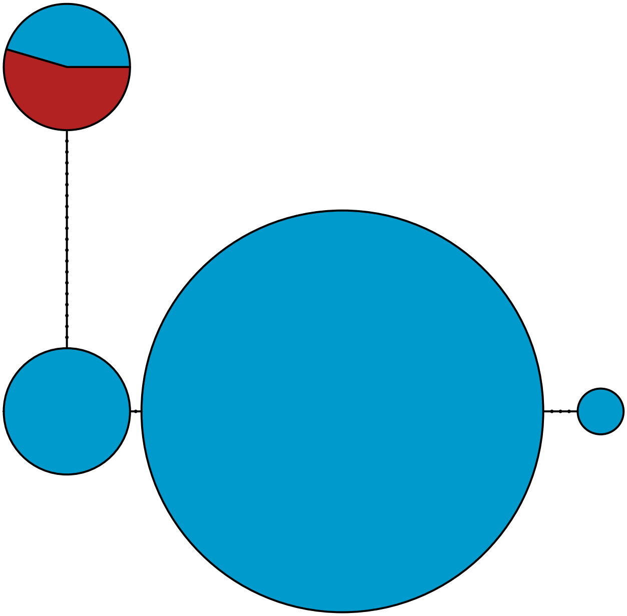

MBPA_000040545 Metazoa

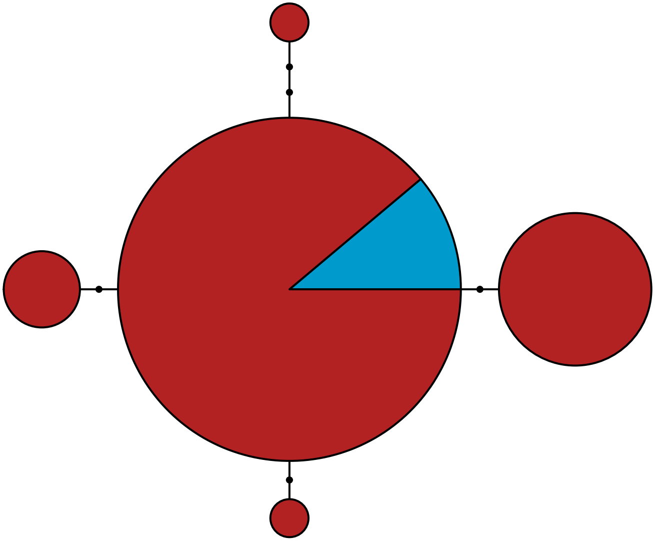

MBPA_000042387

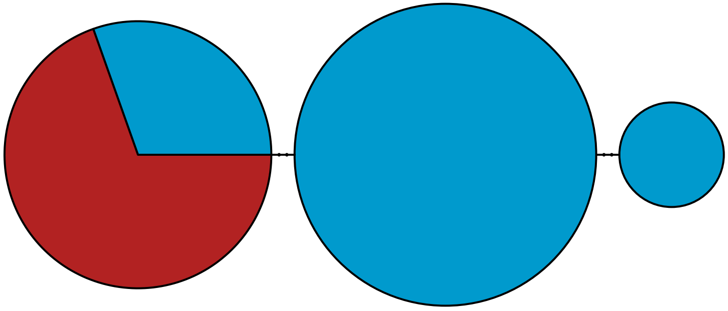

MBPA_000042908

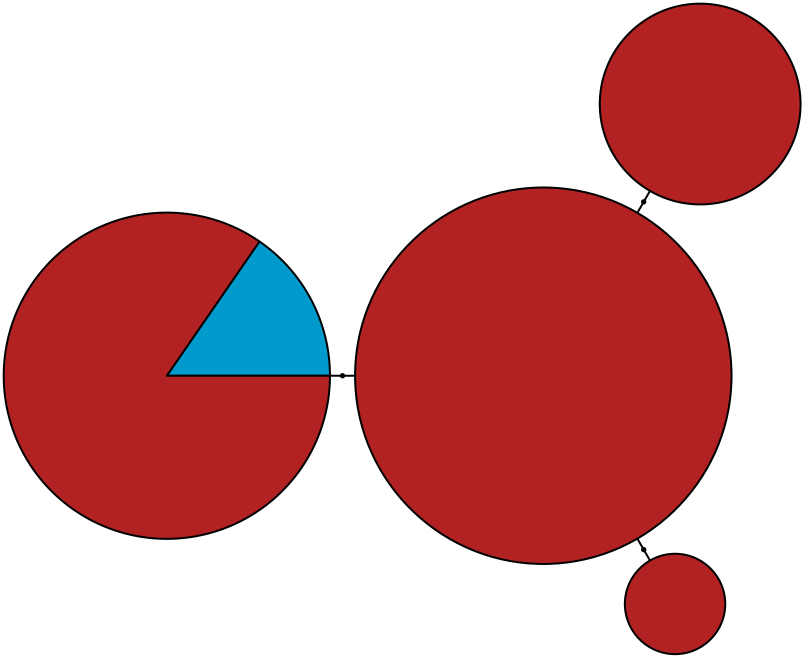

MBPA_000043002 Metazoa

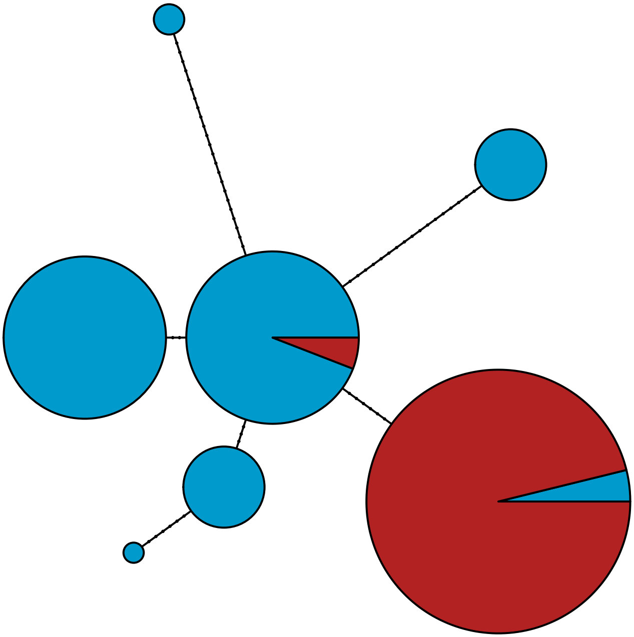

MBPA_000043371

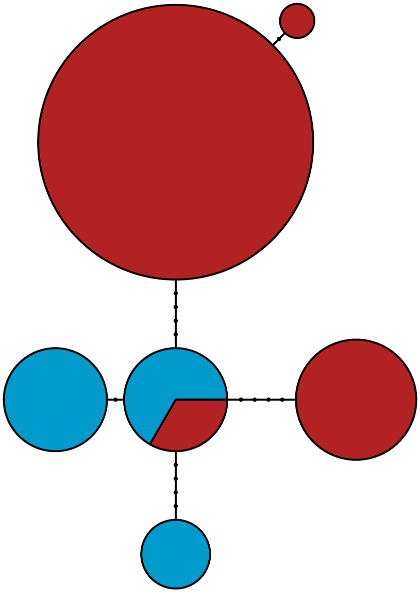

MBPA_000044058 Metazoa

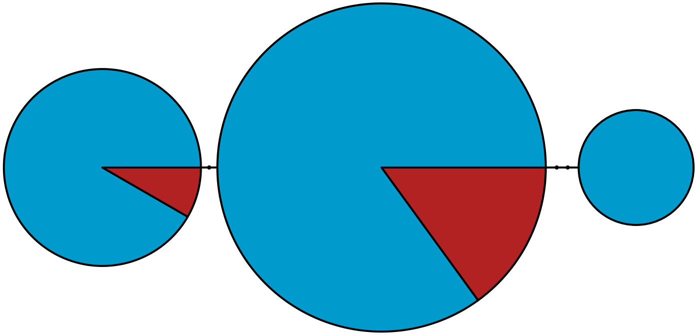

MBPA_000044934 Stramenopiles

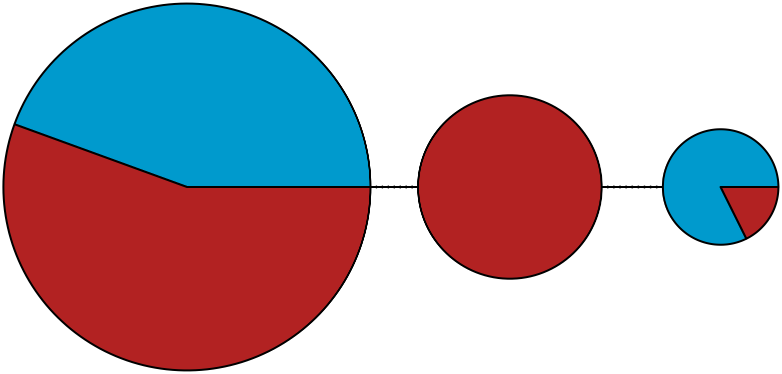

MBPA_000045592

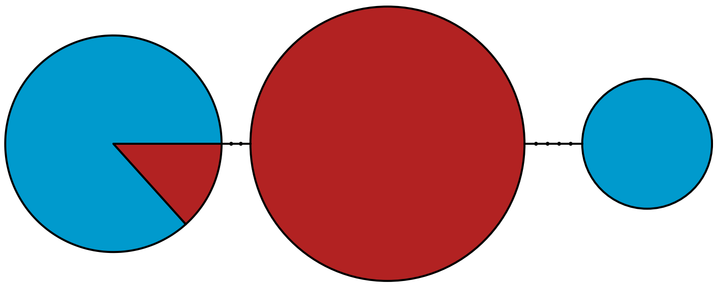

MBPA_000045838 Metazoa

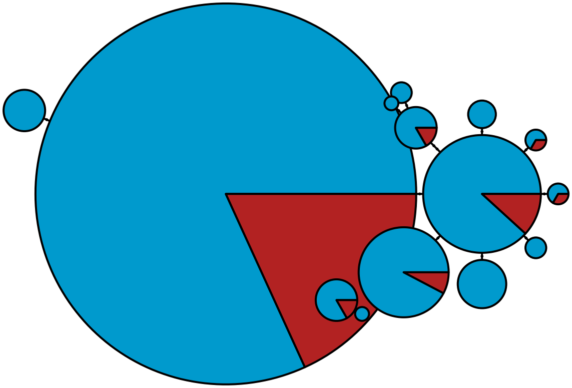

MBPA_000046034 Metazoa

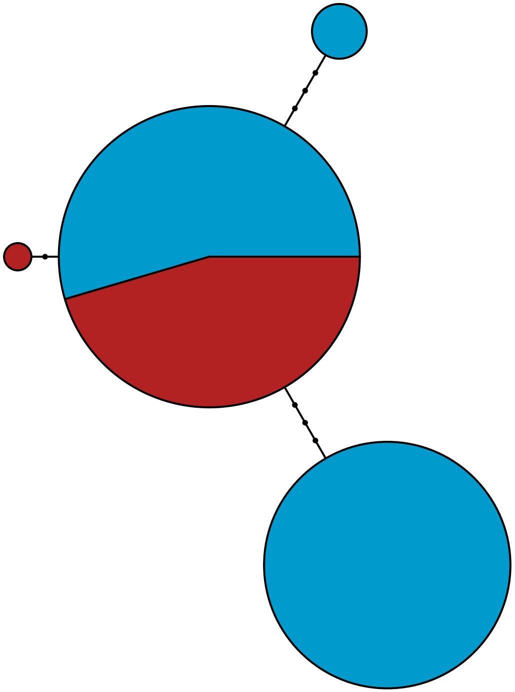

MBPA_000046040 Rhodophyta

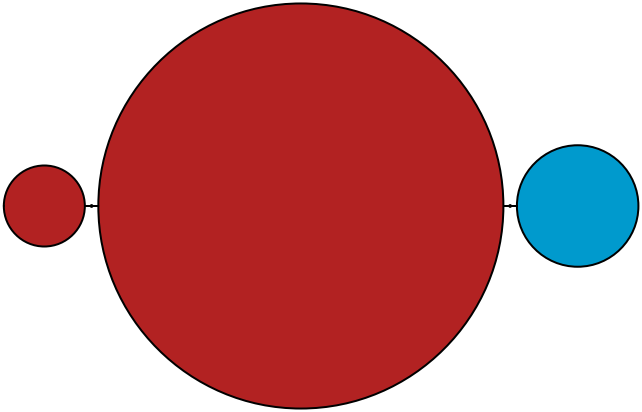

MBPA_000047017 Metazoa

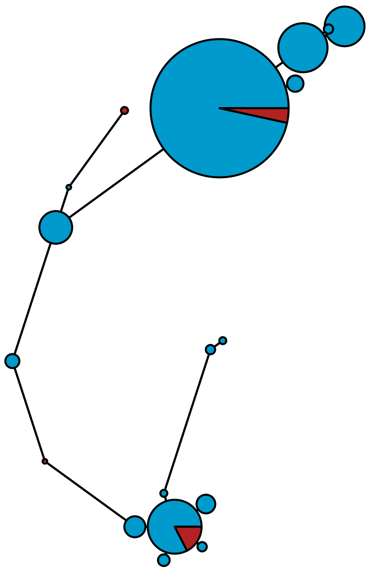

MBPA_000047601 Metazoa

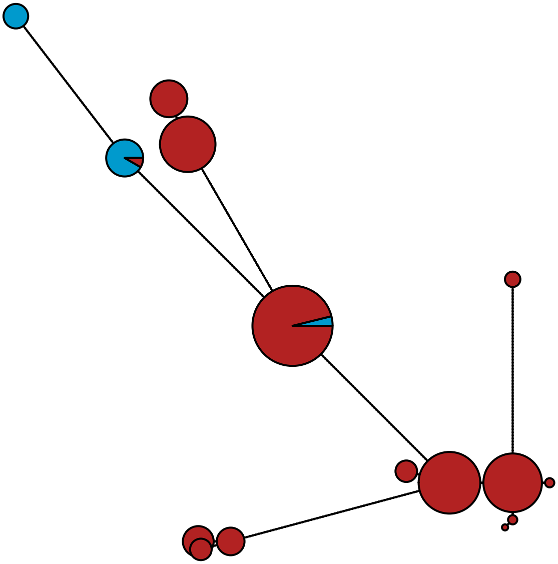

### MBPA_000048533 Metazoa

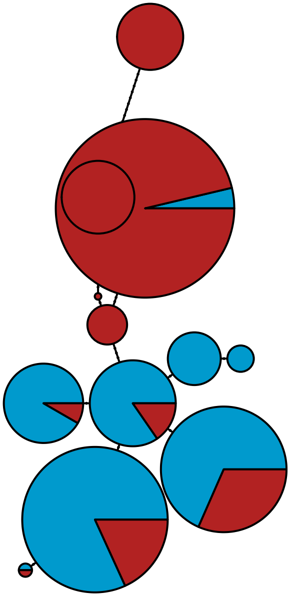

MBPA_000048986 Metazoa

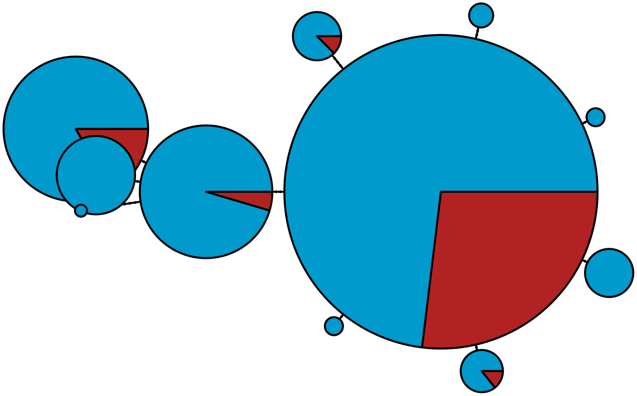

### MBPA_000049367 Metazoa

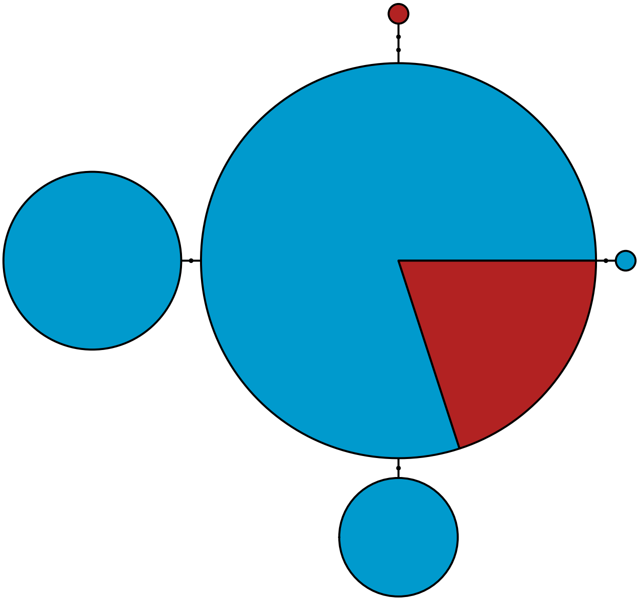

MBPA_000050844 Metazoa

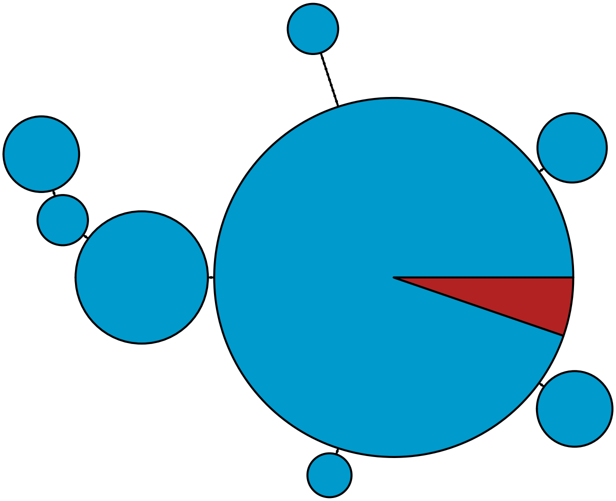

MBPA_000051093 Metazoa

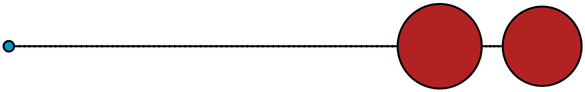

MBPA_000051659 Metazoa

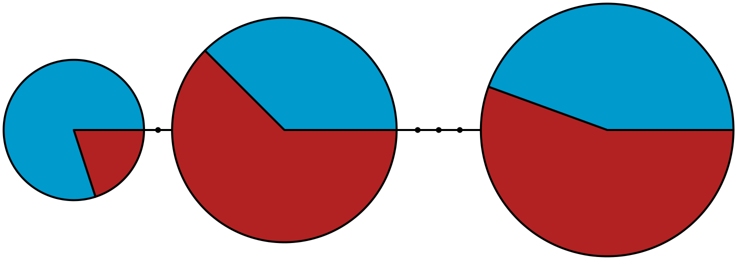

MBPA_000052016 Metazoa

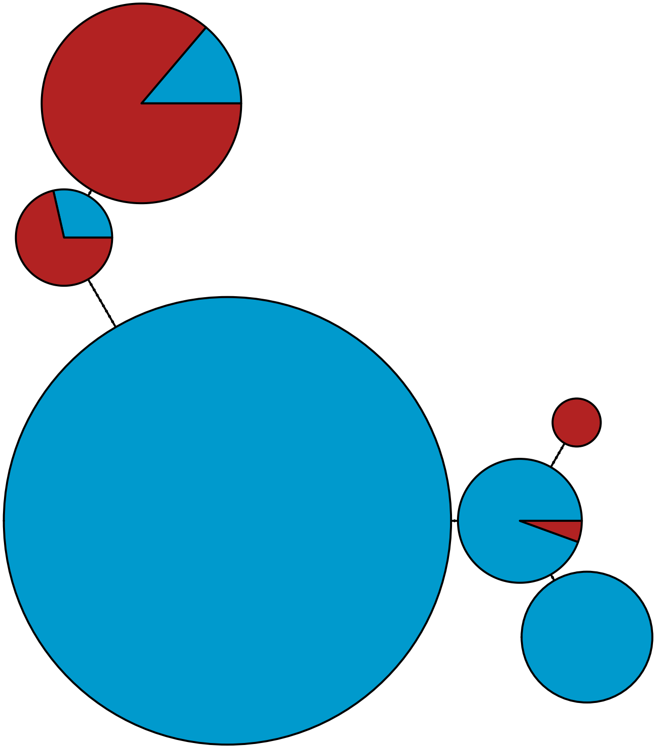

MBPA_000052152 Stramenopiles

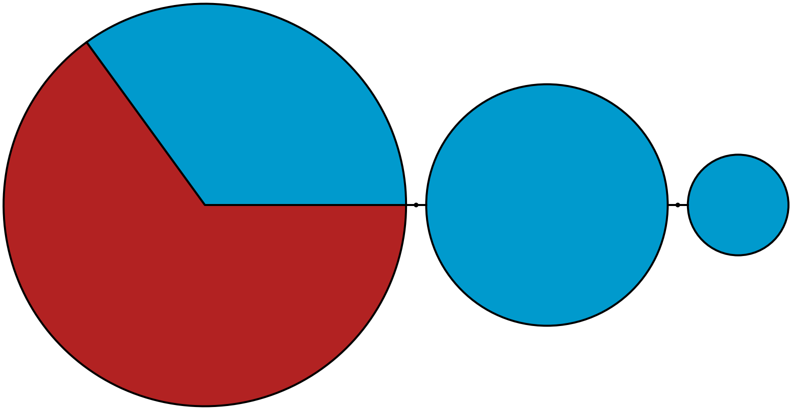

MBPA_000052777 Stramenopiles

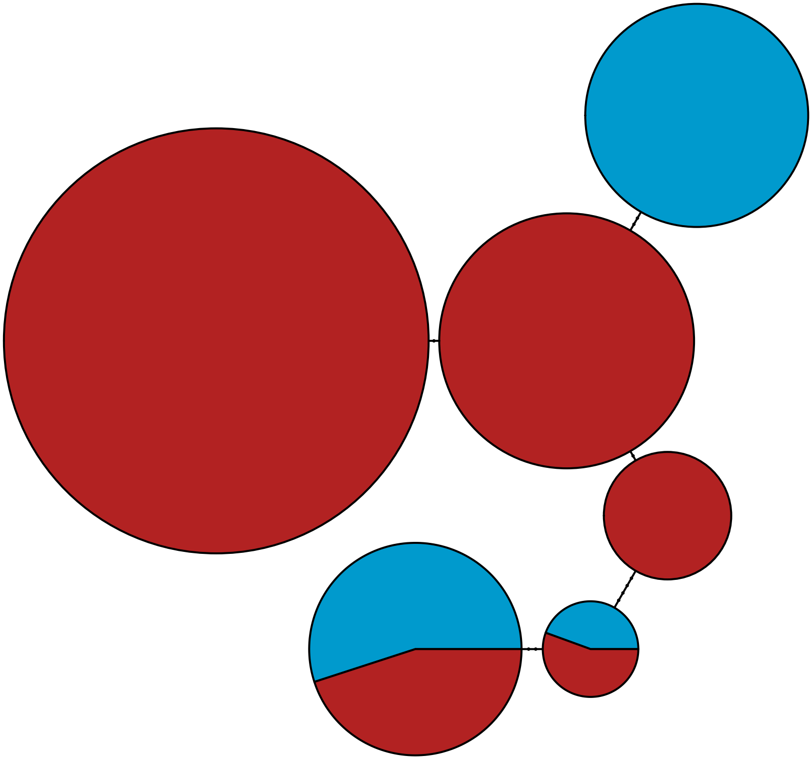

MBPA_000053184

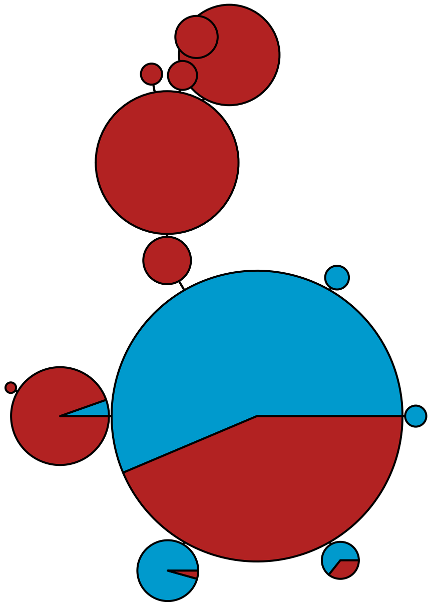

MBPA_000054110

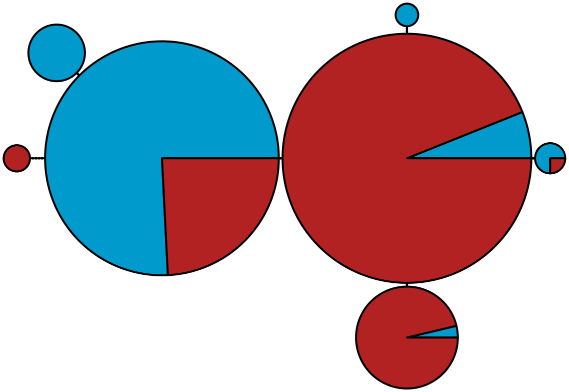

MBPA_000054249

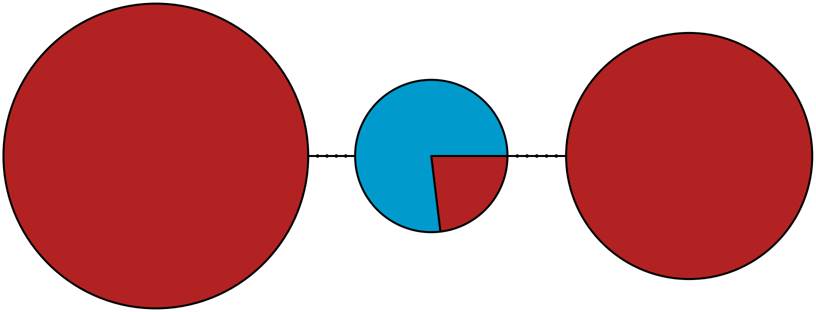

MBPA_000054334 Metazoa

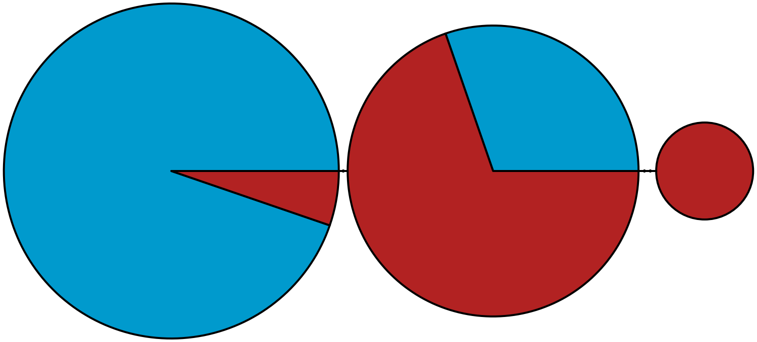

MBPA_000054777 Metazoa

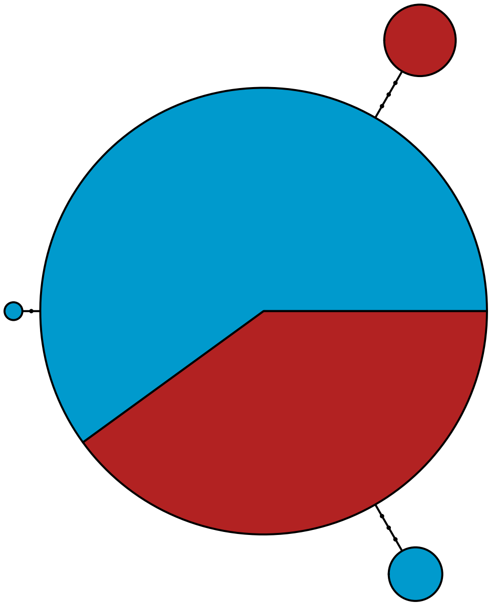

MBPA_000055452 Metazoa

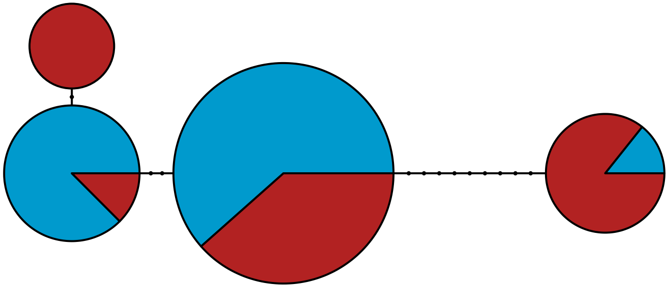

MBPA_000055600

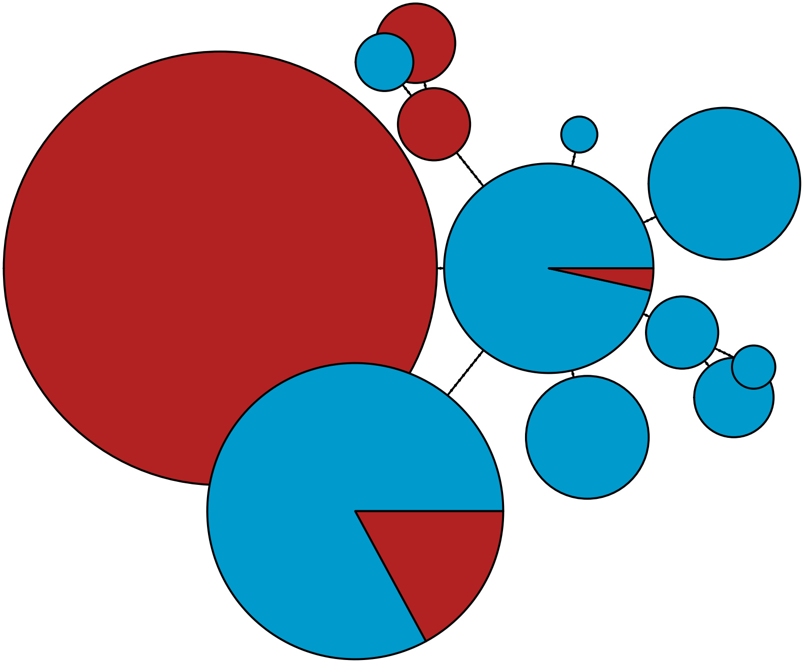

MBPA_000056965 Metazoa

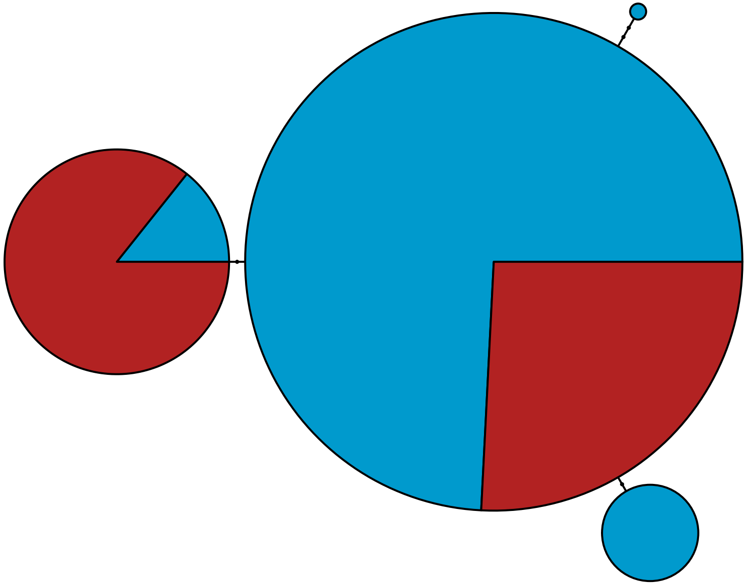

MBPA_000057470 Rhodophyta

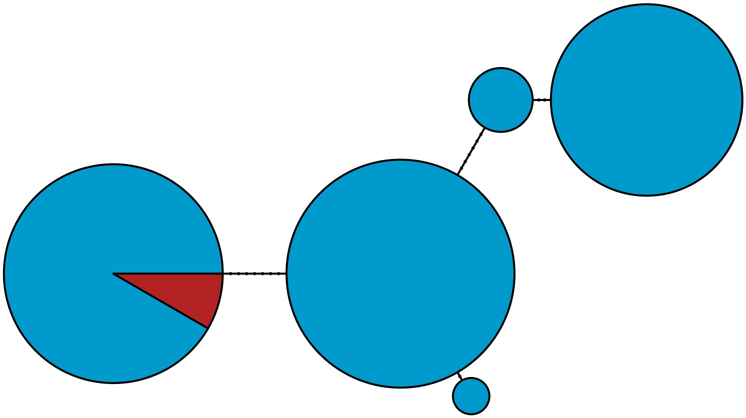

MBPA_000058489 Stramenopiles

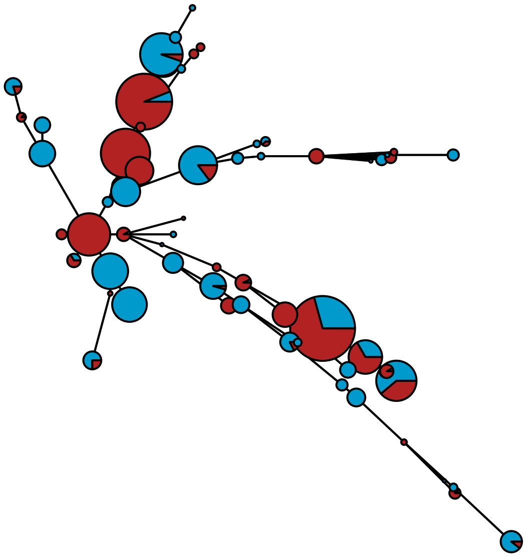

MBPA_000060150 Rhodophyta

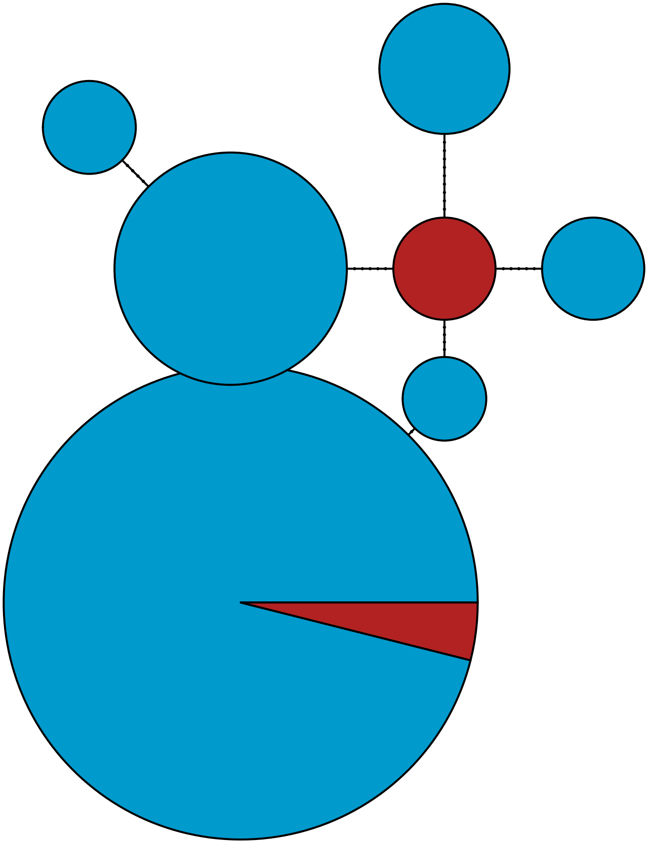

MBPA_000060921 Metazoa

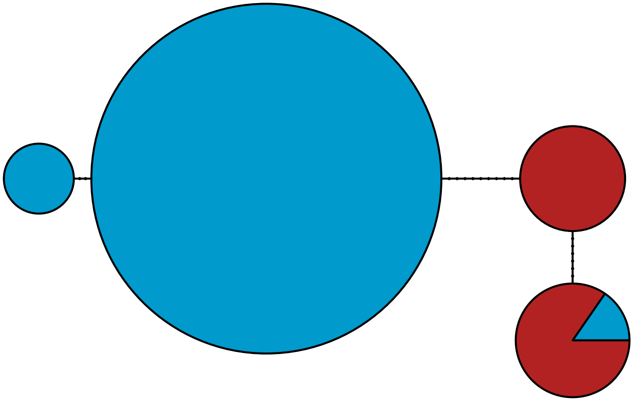

MBPA_000061051 Stramenopiles

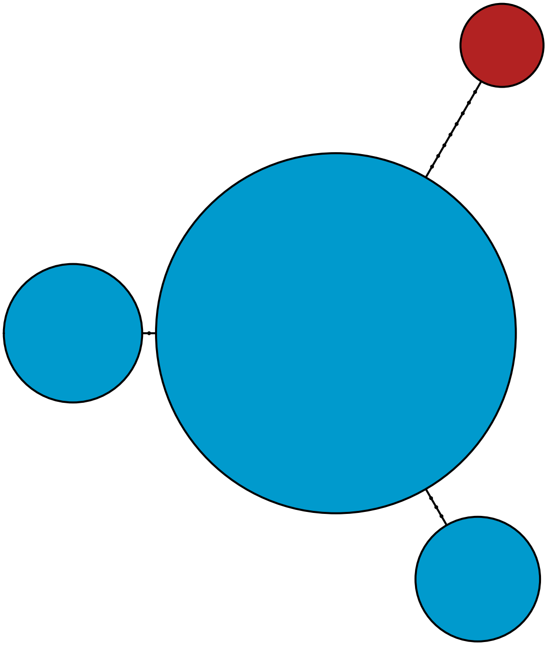

MBPA_000061226 Metazoa

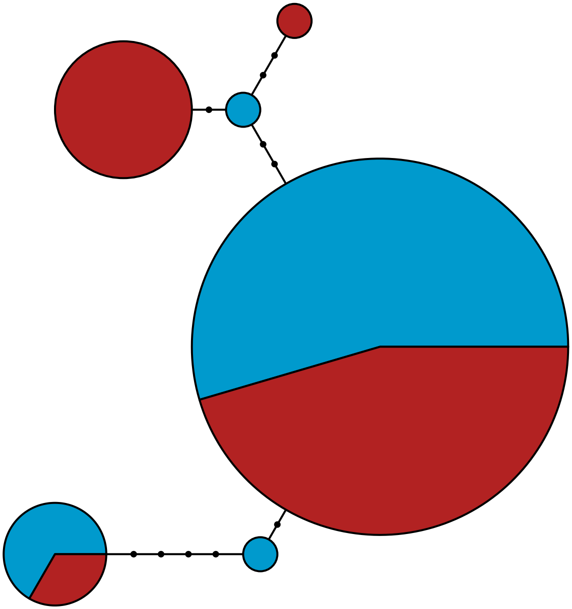

MBPA_000061821 Rhodophyta

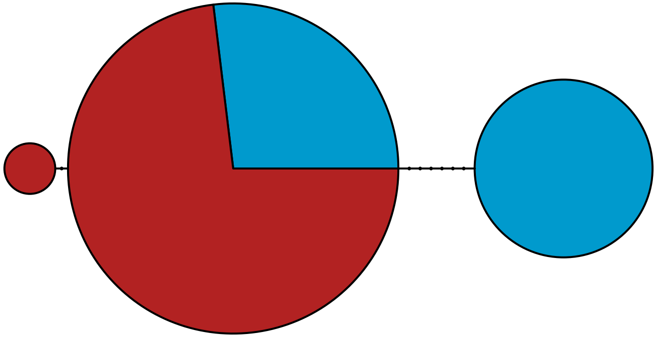

MBPA_000062823 Rhodophyta

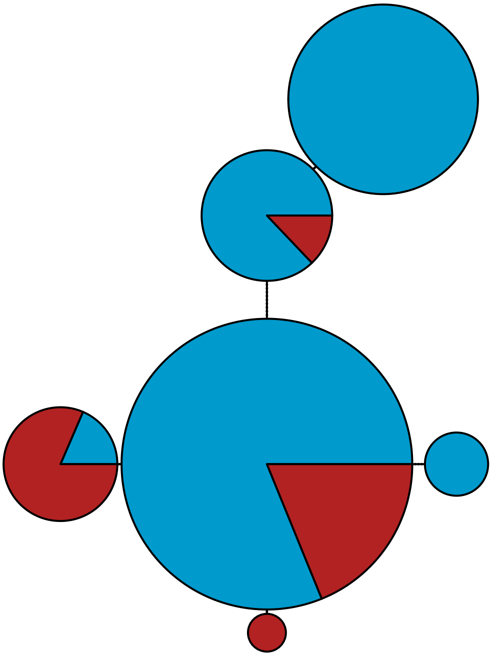

MBPA_000063504 Metazoa

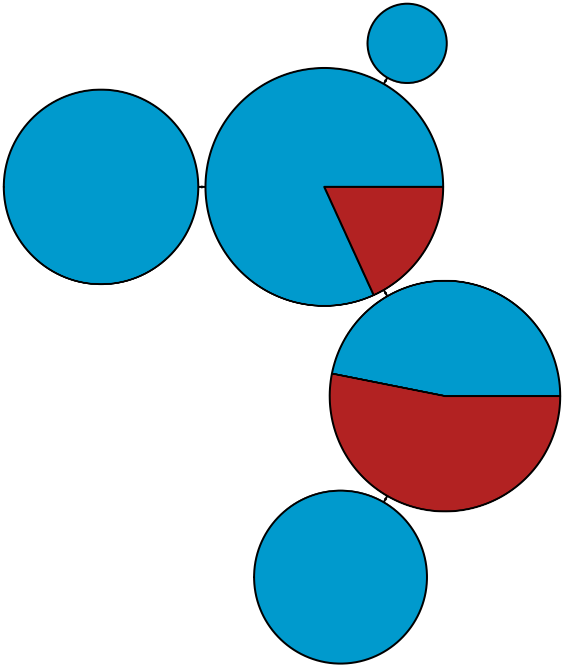

MBPA_000064145 Metazoa

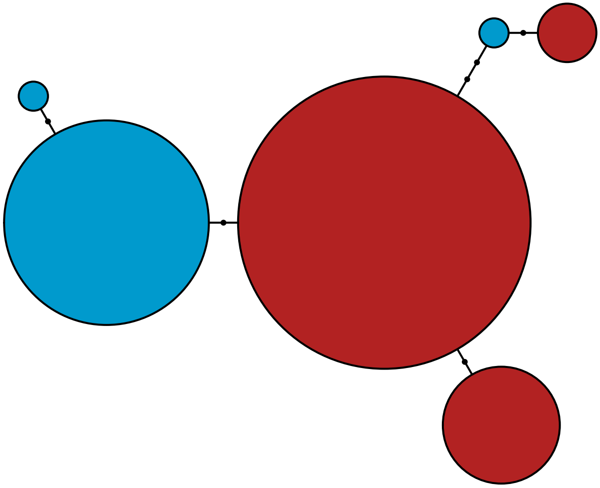

MBPA_000065282 Metazoa

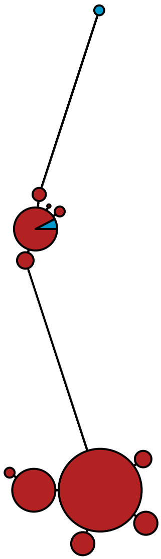

MBPA_000065793 Metazoa

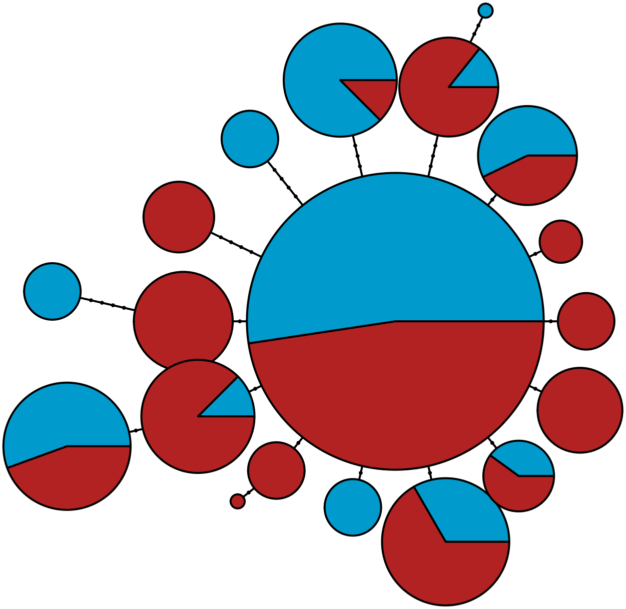

MBPA_000068277 Metazoa

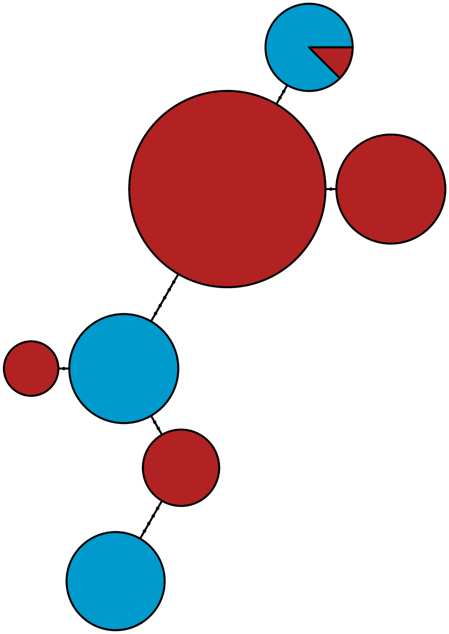

MBPA_000069234 Metazoa

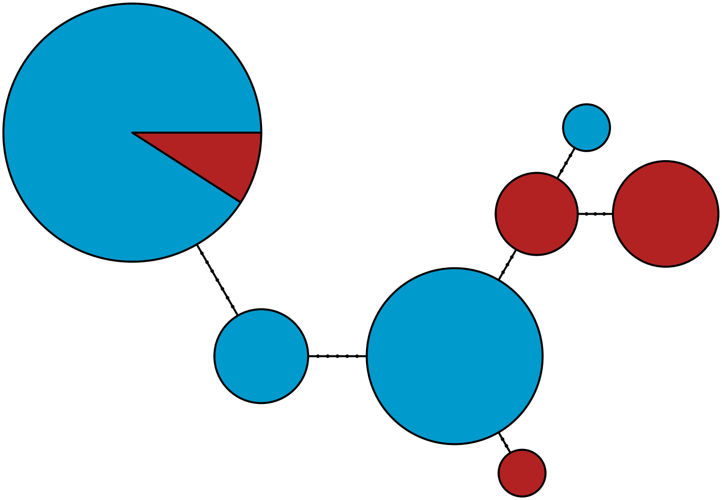

MBPA_000069671 Stramenopiles

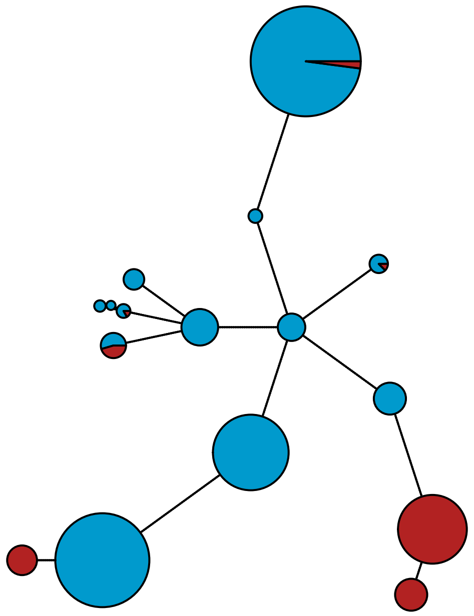

MBPA_000071470 Stramenopiles

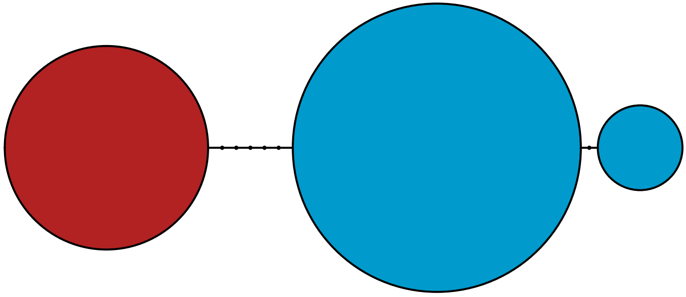

MBPA_000071968 Metazoa

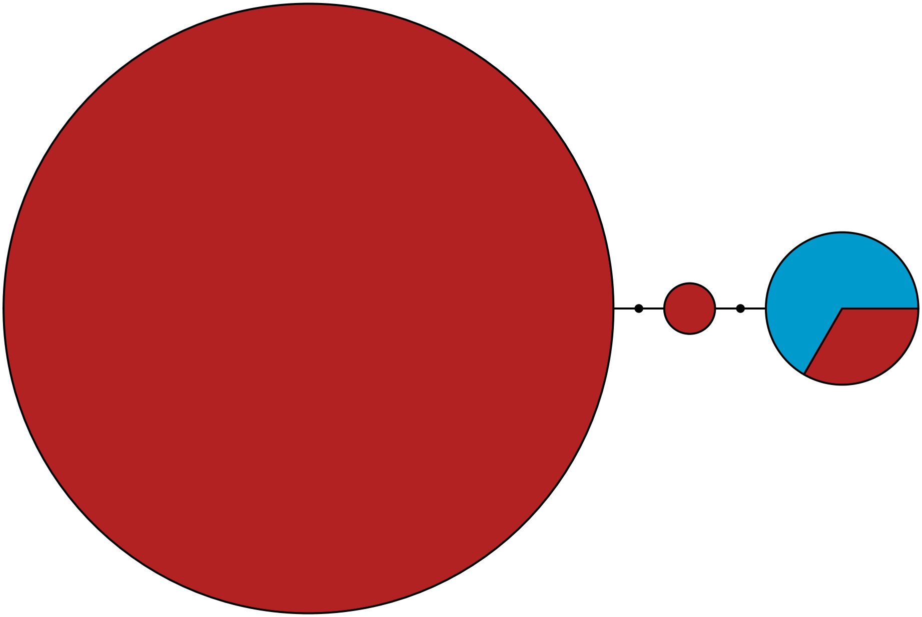

MBPA_000072398 Stramenopiles

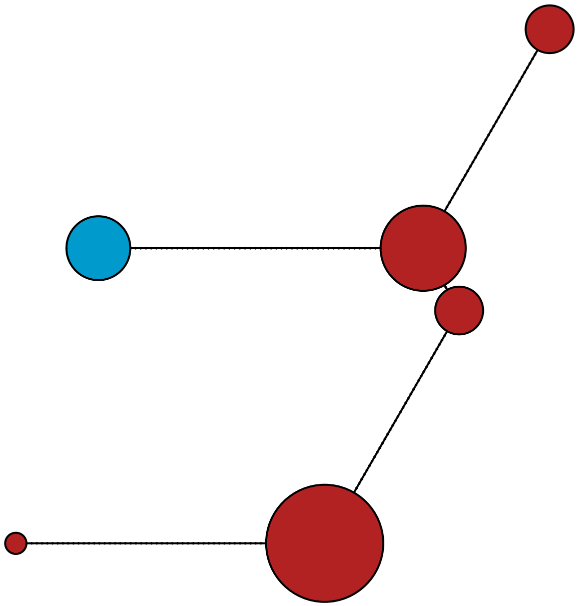

MBPA_000072895

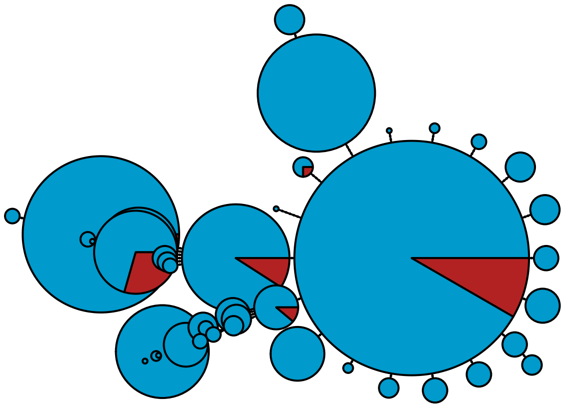

MBPA_000074040 Rhodophyta

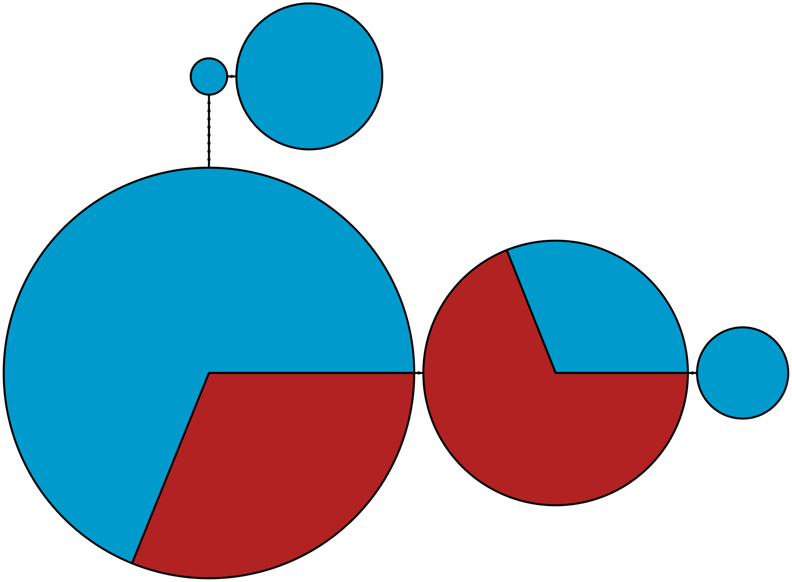

MBPA_000076199

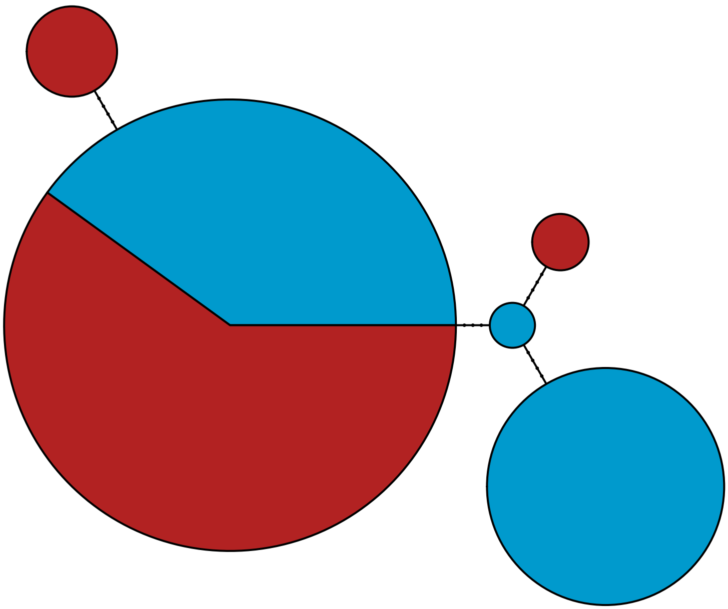

MBPA_000080298 Stramenopiles

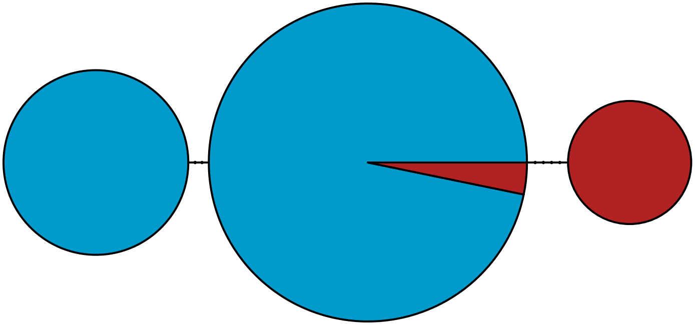

MBPA_000080538 Metazoa

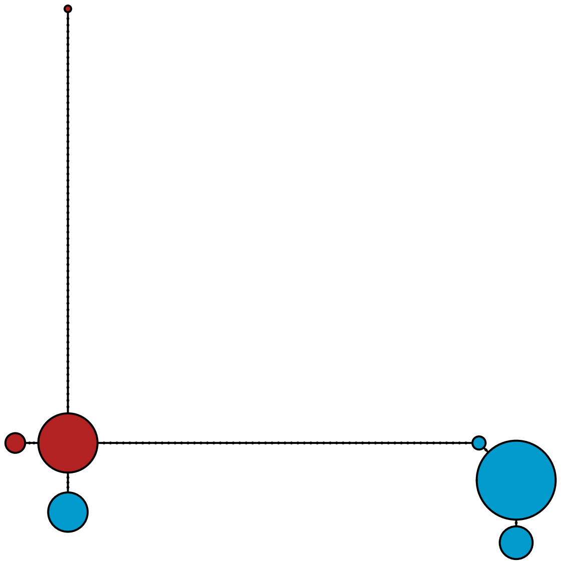

MBPA_000083133 Metazoa

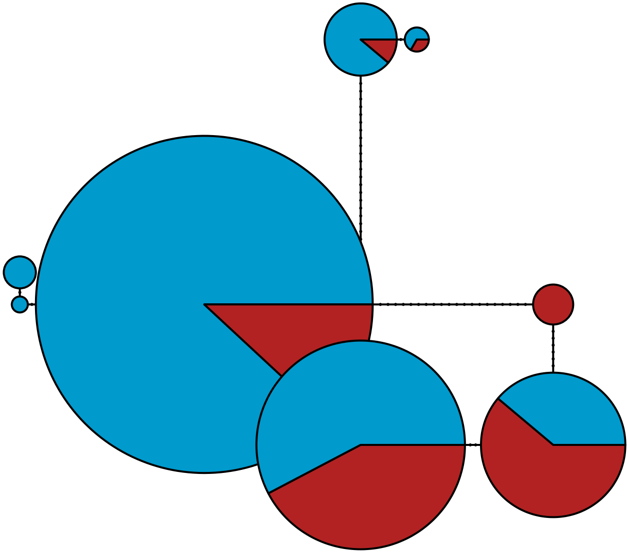

MBPA_000083685 Stramenopiles

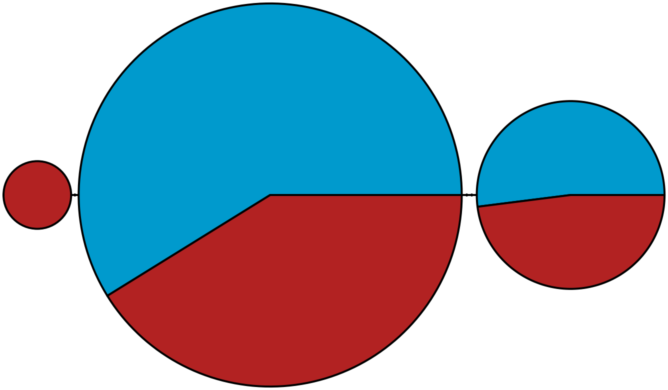

MBPA_000084918 Stramenopiles

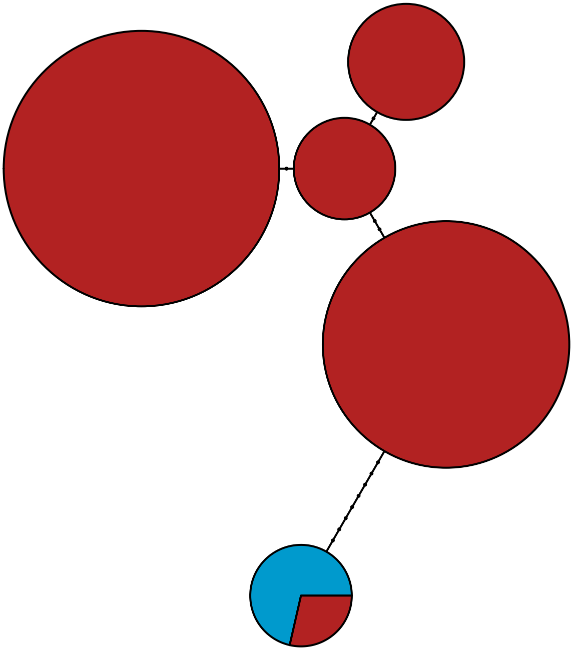

MBPA_000087707 Metazoa

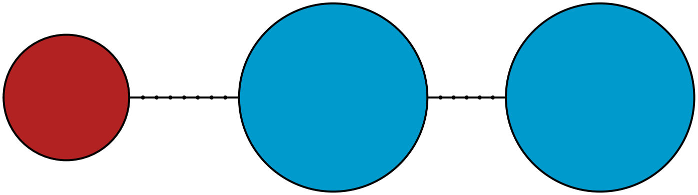

MBPA_000089403

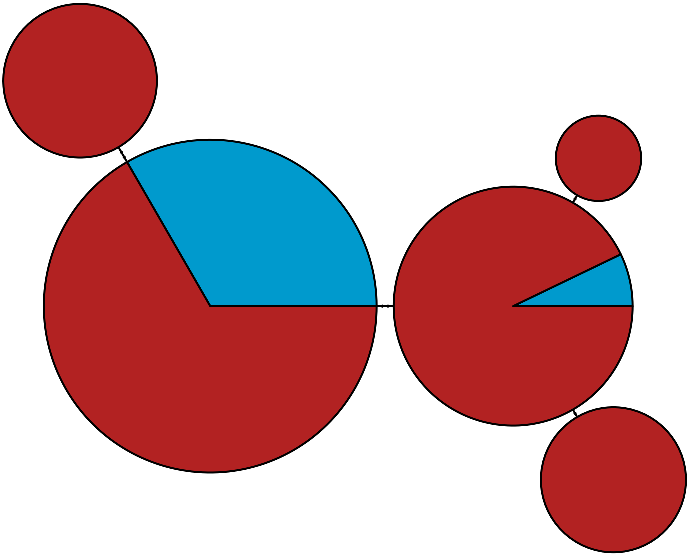

MBPA_000089929 Stramenopiles

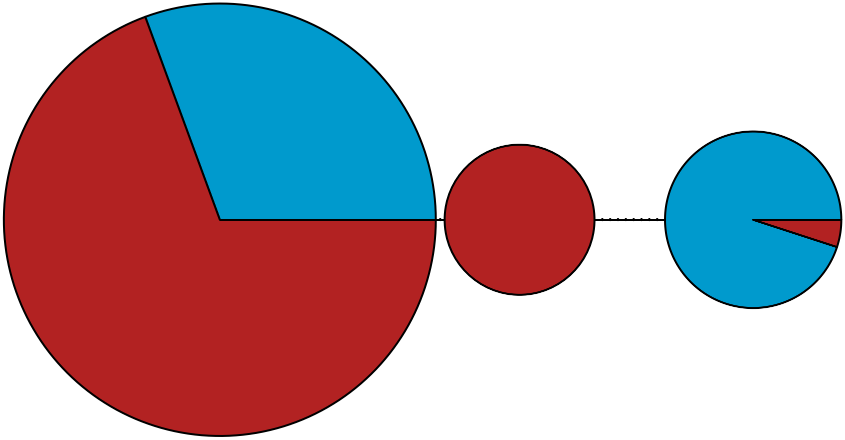

MBPA_000090227 Metazoa

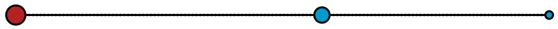

MBPA_000092583 Metazoa

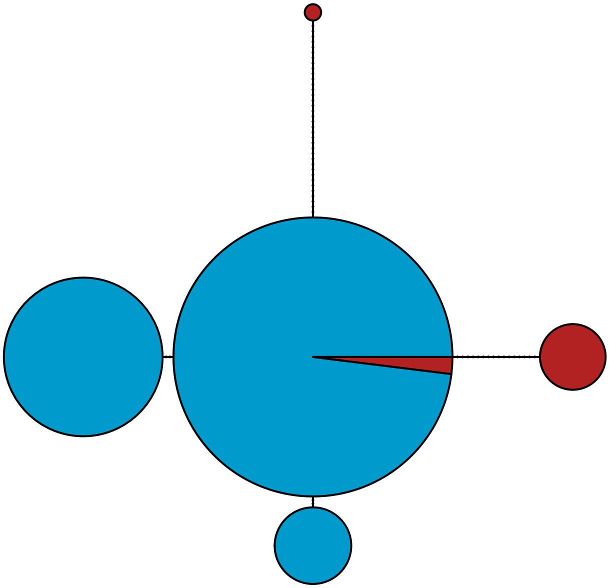

MBPA_000092639 Metazoa

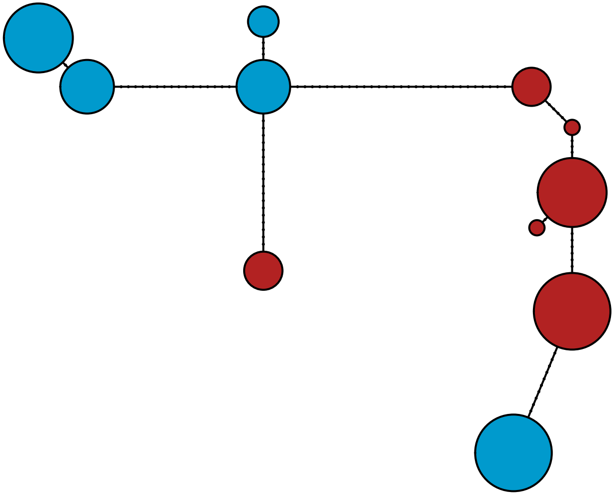

MBPA_000092849 Metazoa

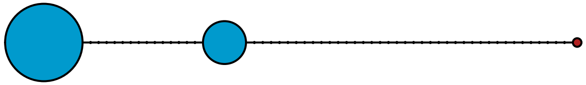

MBPA_000093092 Stramenopiles

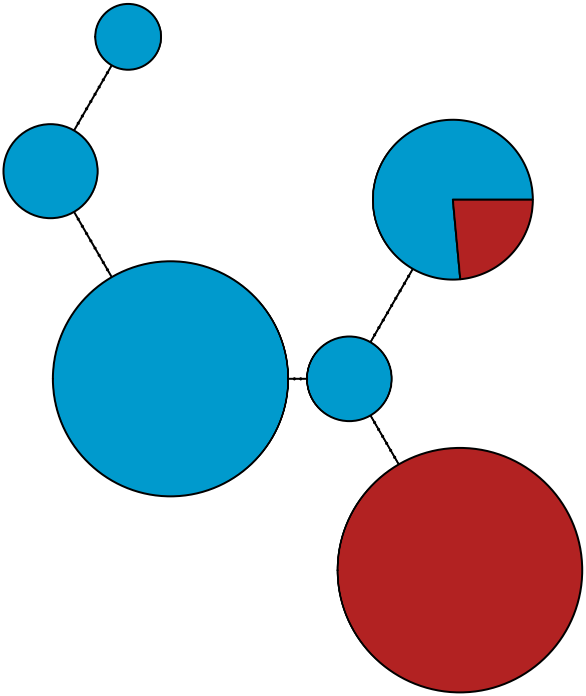

MBPA_000095064 Rhodophyta

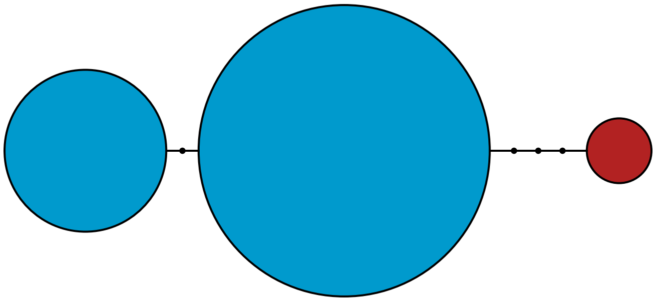

MBPA_000097088 Metazoa

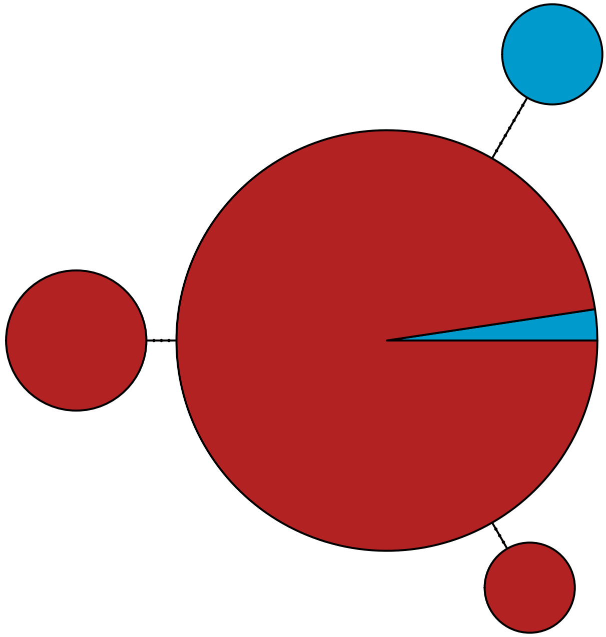

MBPA_000099451

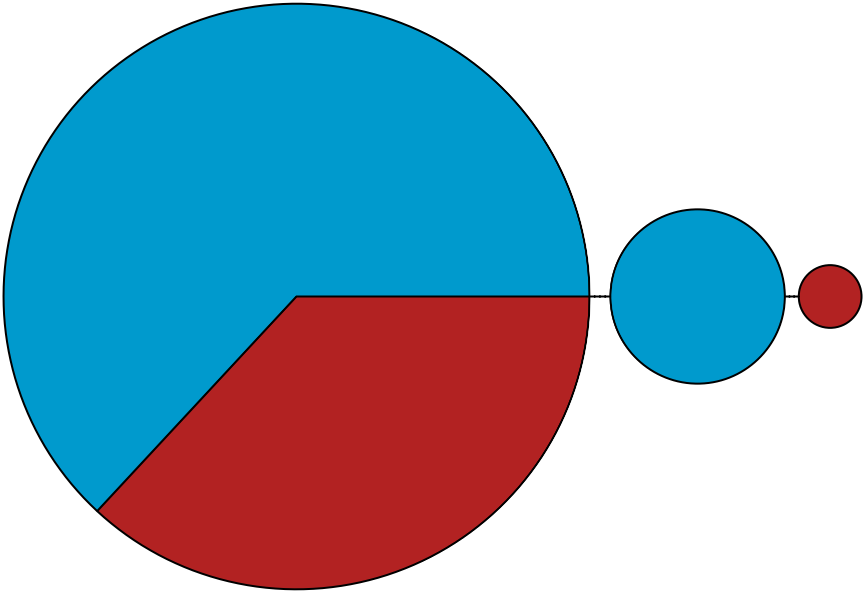

MBPA_000099769

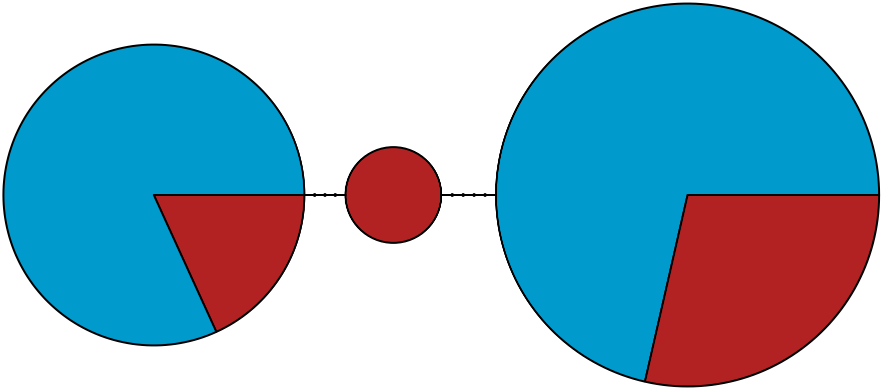

MBPA_000103432 Lobosa

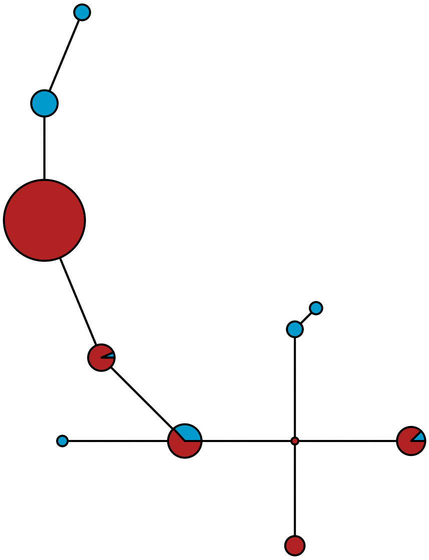

MBPA_000104492 Metazoa

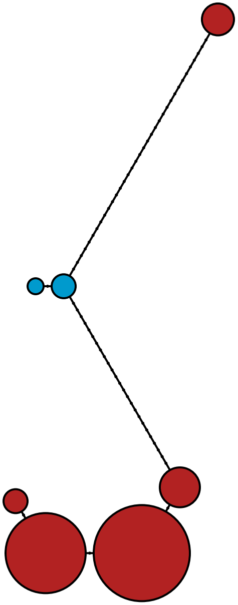

MBPA_000105646 Rhodophyta

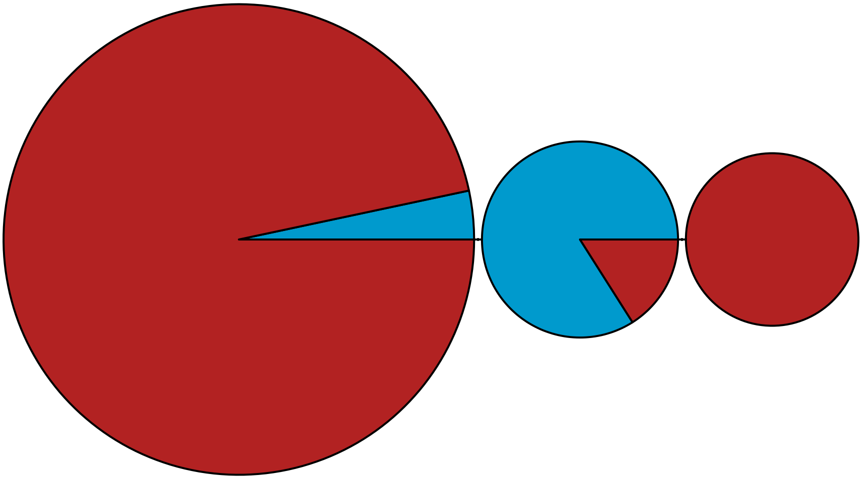

MBPA_000105665 Metazoa

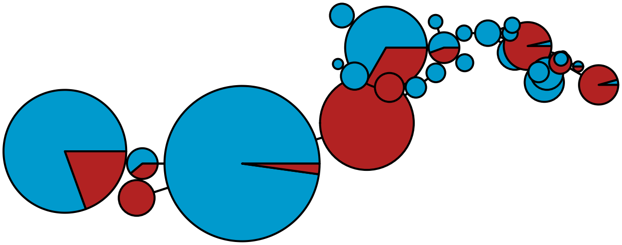

MBPA_000105765

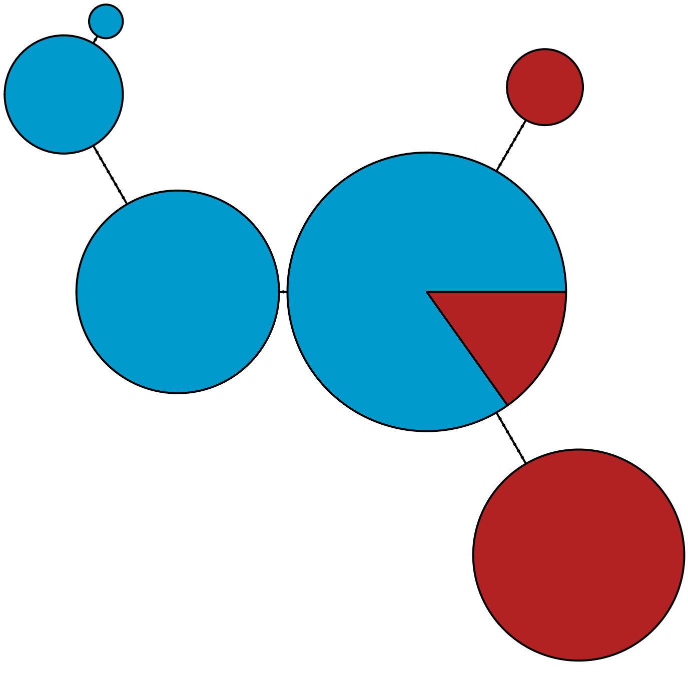

MBPA_000106995 Stramenopiles

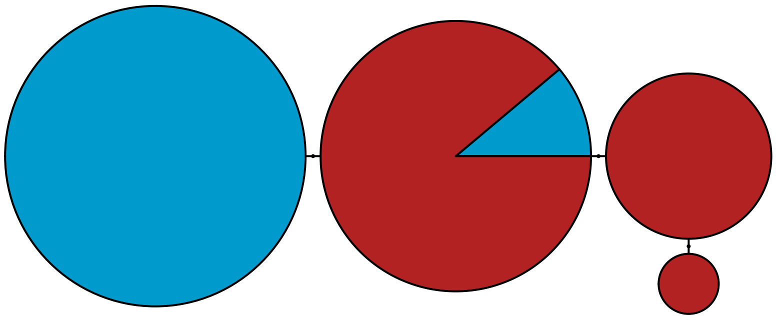

MBPA_000108530

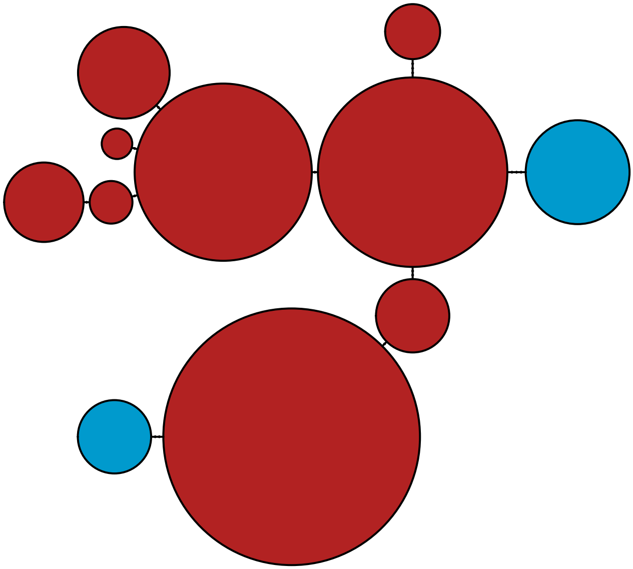

MBPA_000111858 Rhodophyta

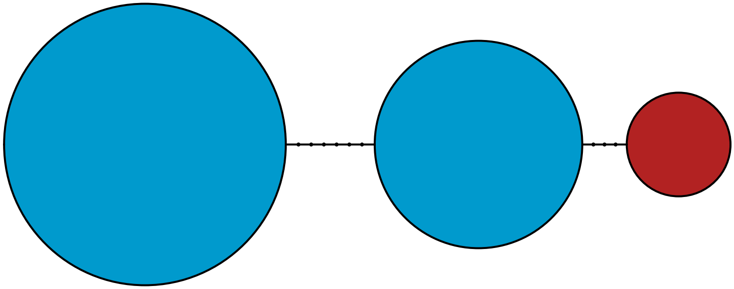

MBPA_000117637 Metazoa

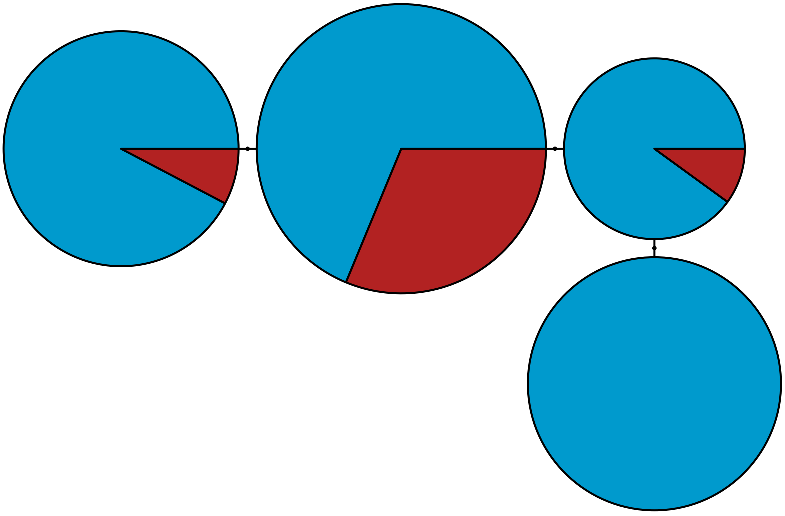

MBPA_000123345

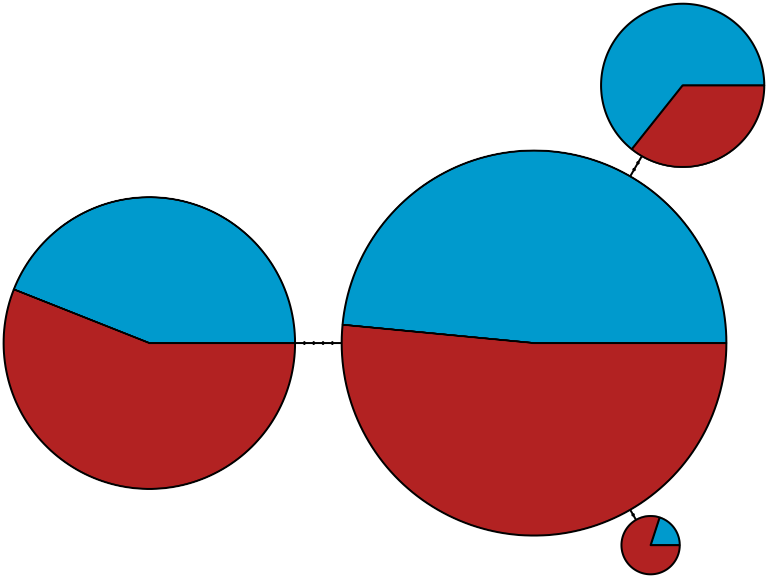

MBPA_000123662 Stramenopiles

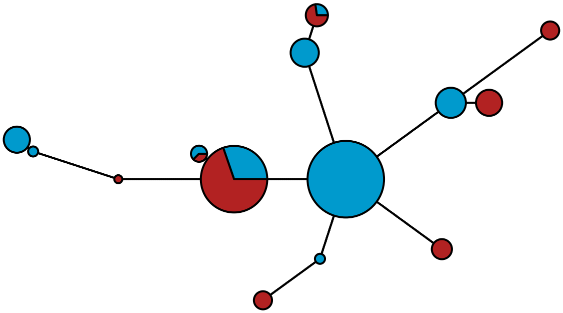

MBPA_000124145 Stramenopiles

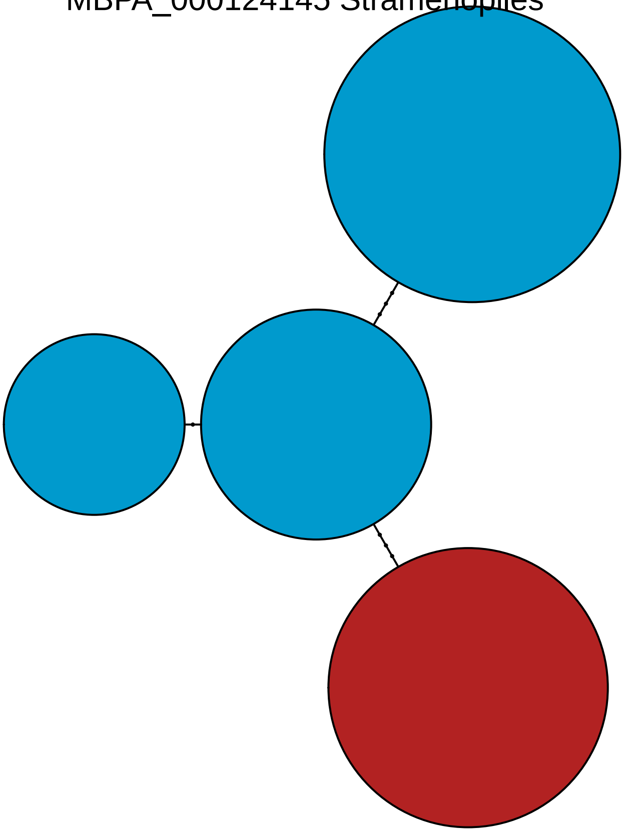

MBPA_000124425

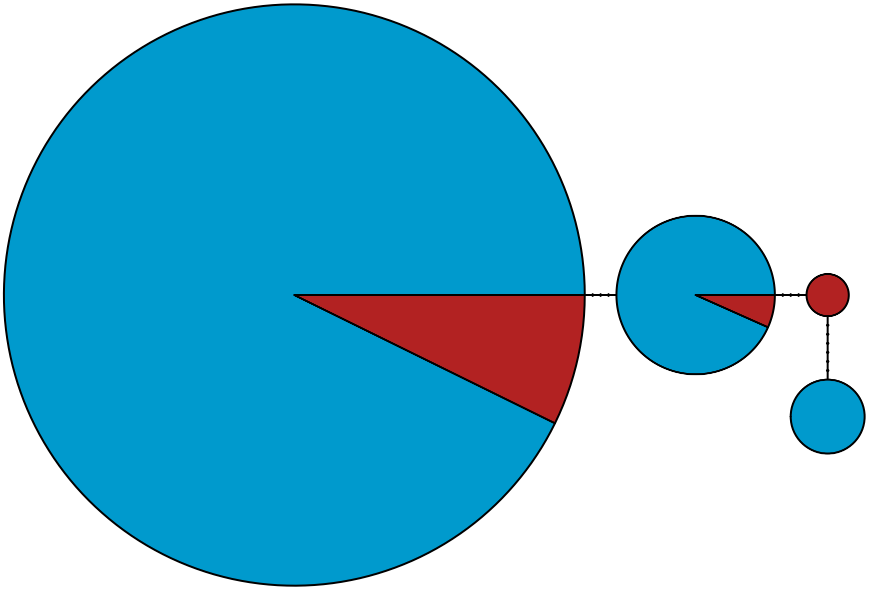

MBPA_000126019

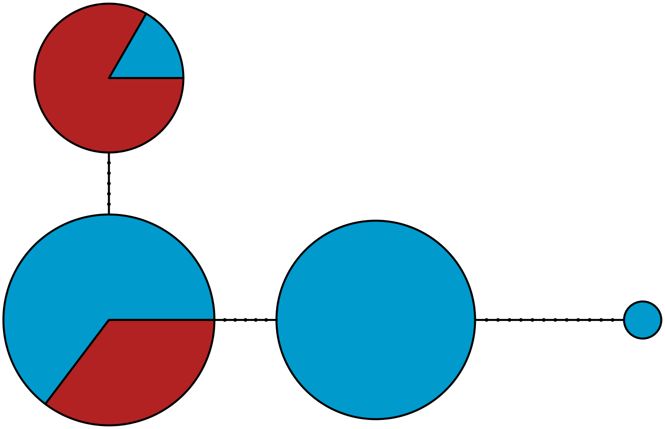

MBPA_000130732 Metazoa

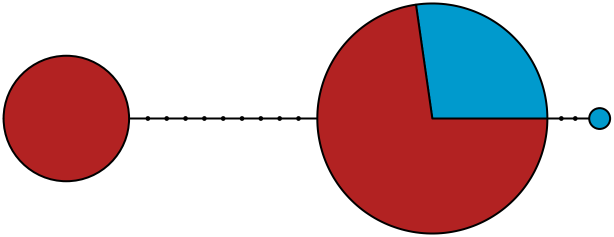

MBPA_000134877 Metazoa

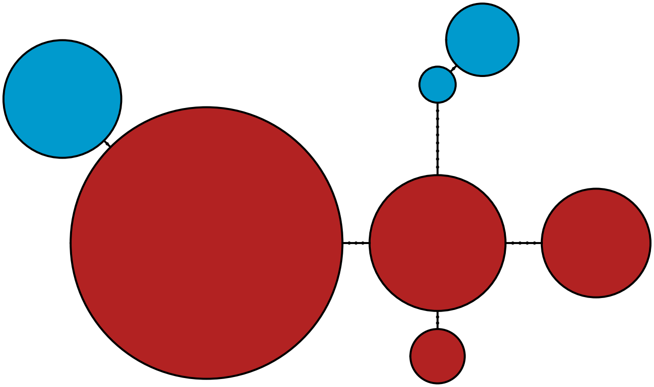

MBPA_000136012 Stramenopiles

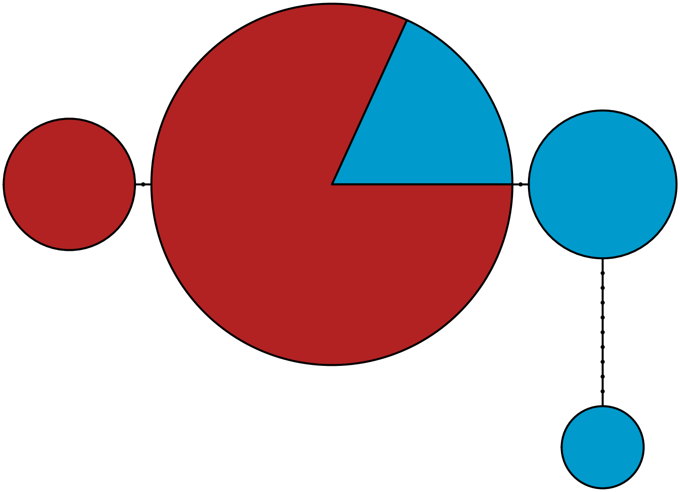

MBPA_000140352 Stramenopiles

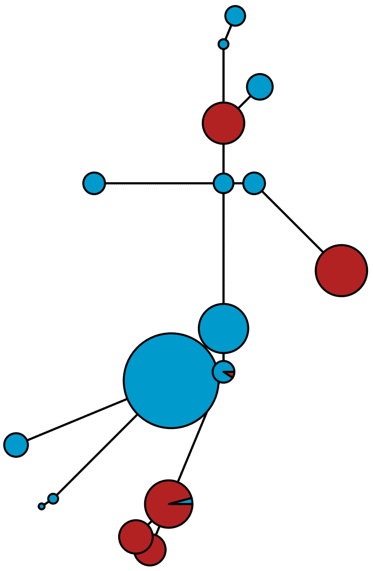

MBPA_000157139 Stramenopiles

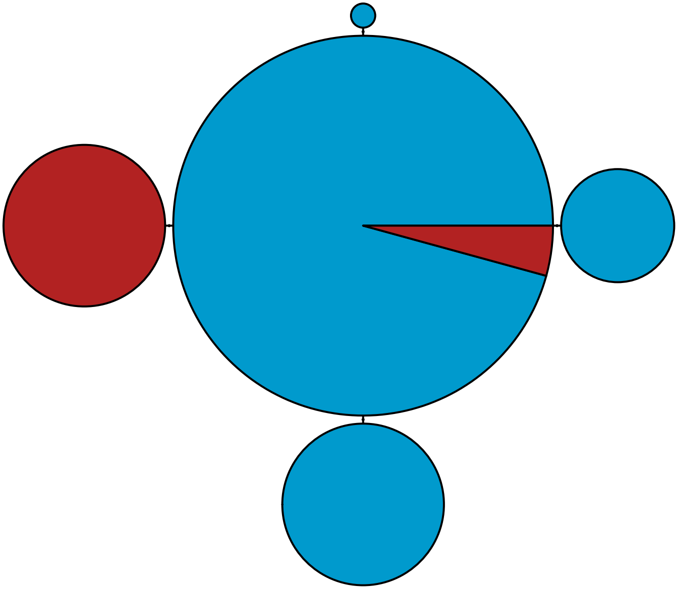

MBPA_000157502 Stramenopiles

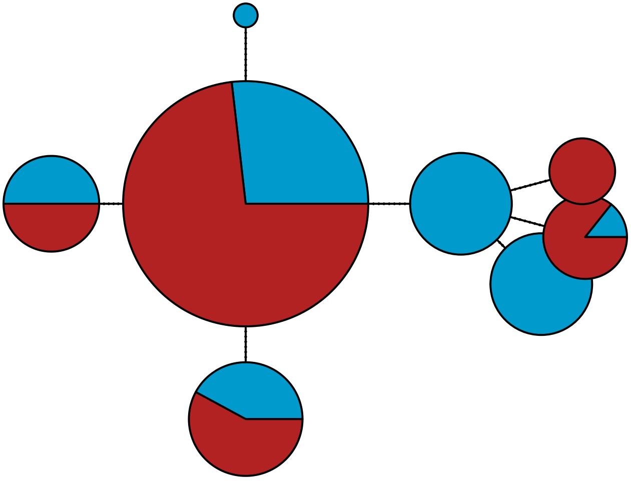

MBPA_000158774 Metazoa

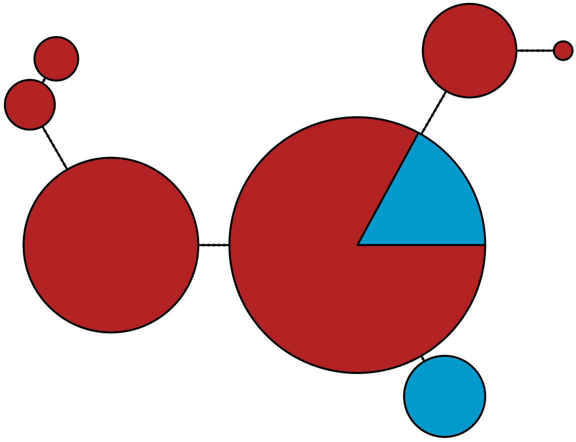

MBPA_000170237

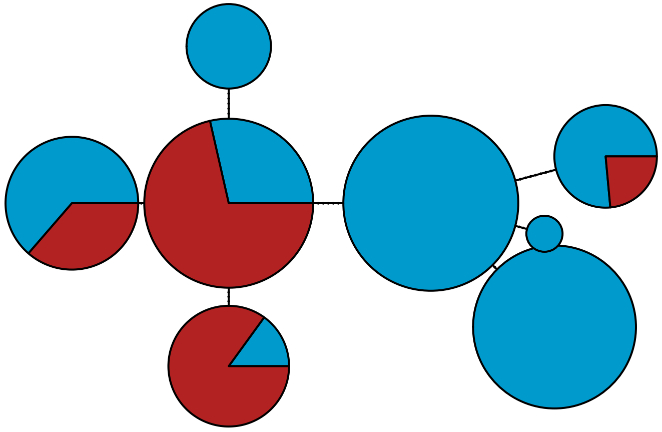

MBPA_000175827 Rhodophyta

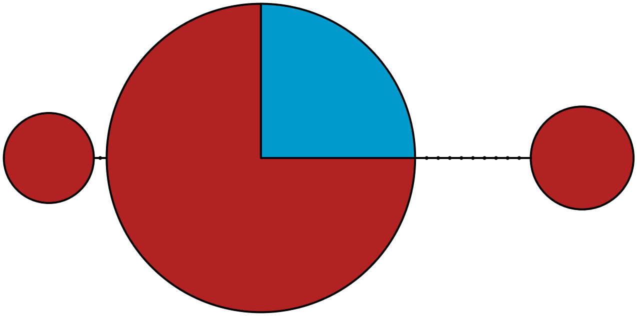

MBPA_000178984 Metazoa

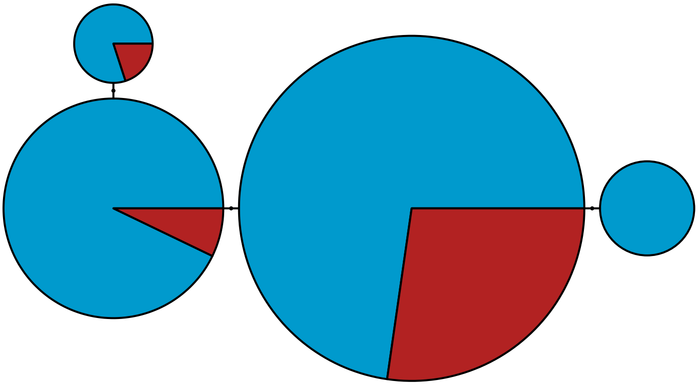

MBPA_000182011 Stramenopiles

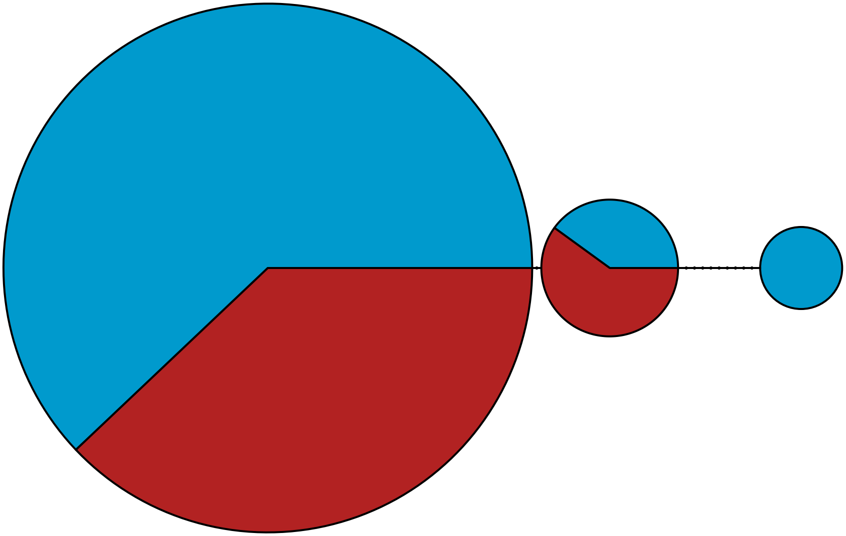

MBPA_000183538 Stramenopiles

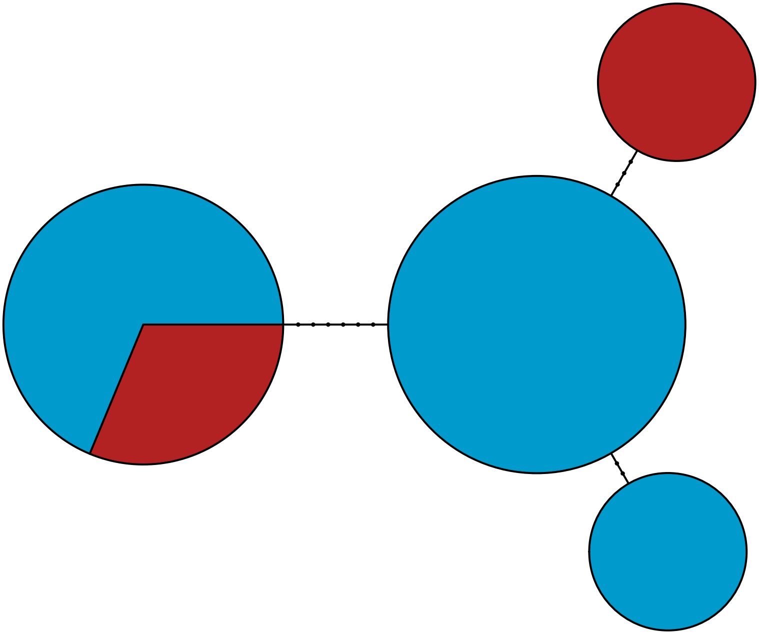

MBPA_000193298 Metazoa

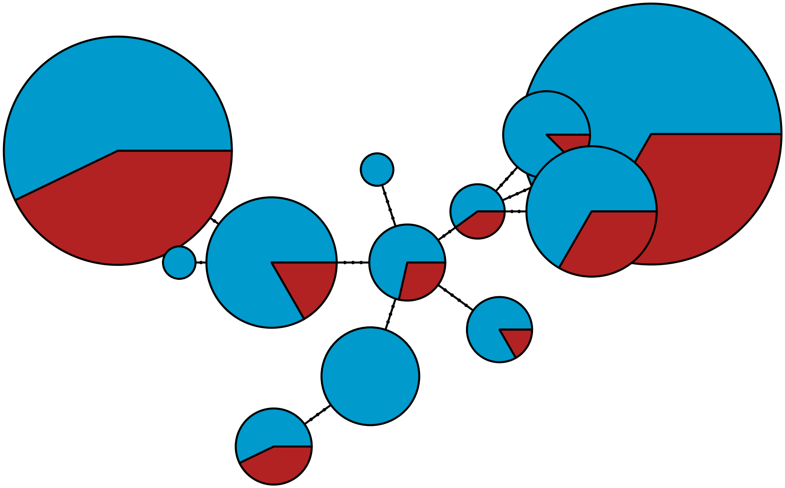

MBPA_000195626

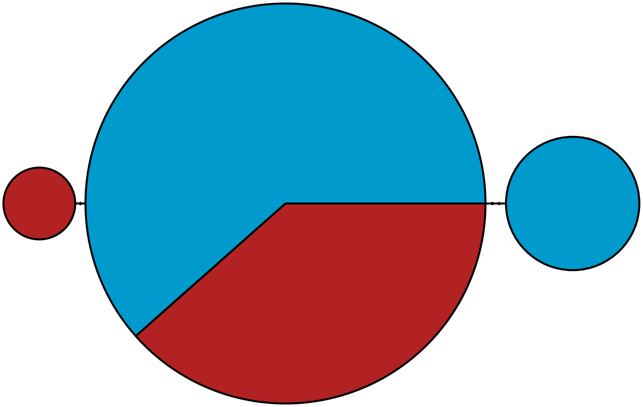

MBPA_000205830

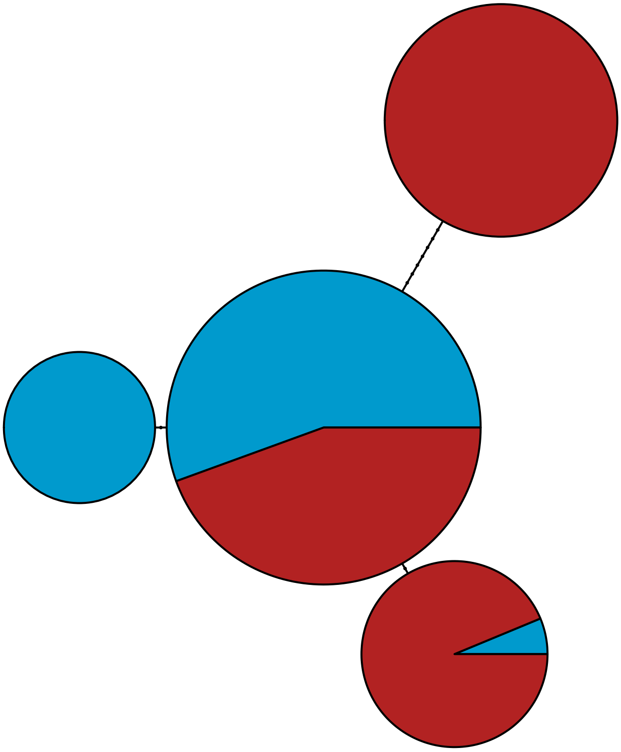

MBPA_000206007

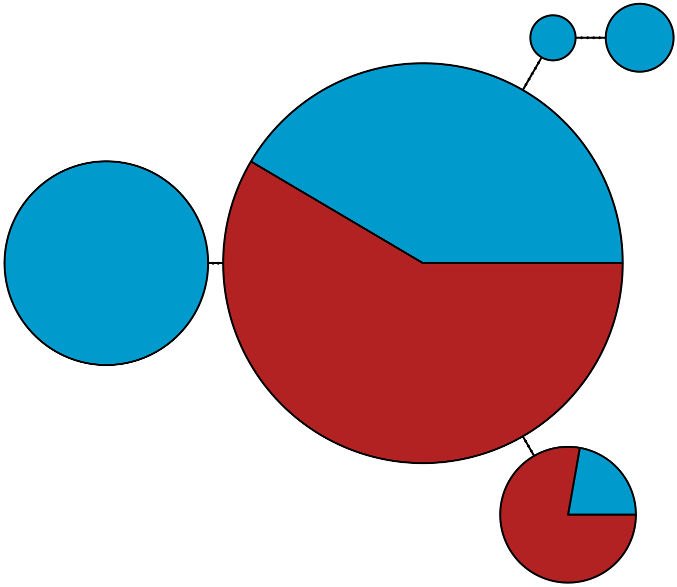

MBPA_000209094

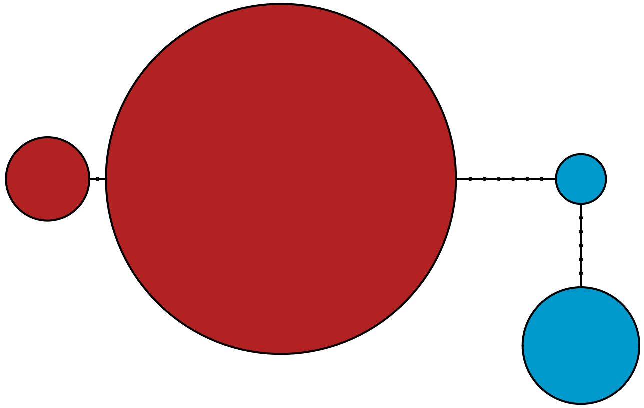

MBPA_000214230 Rhodophyta

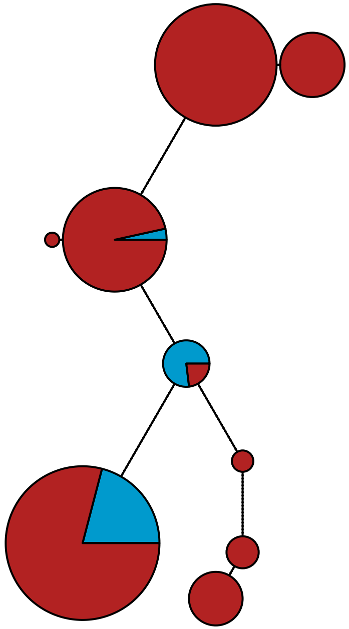

MBPA_000217727 Stramenopiles

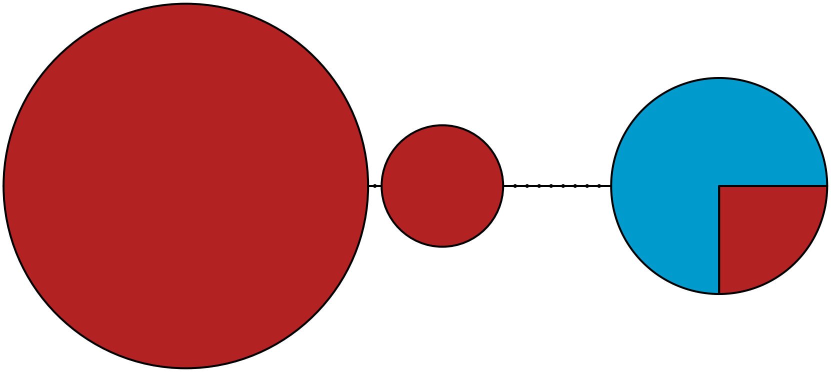

MBPA_000217887 Stramenopiles

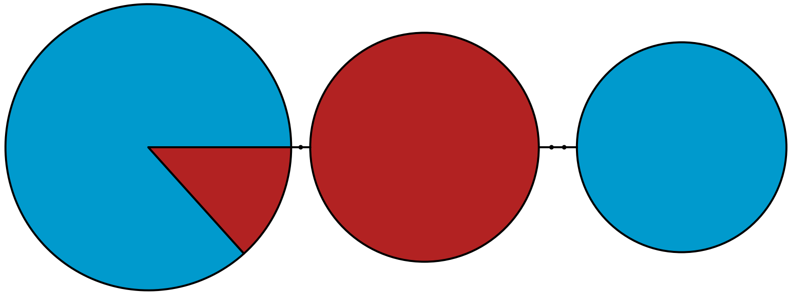

MBPA_000221050

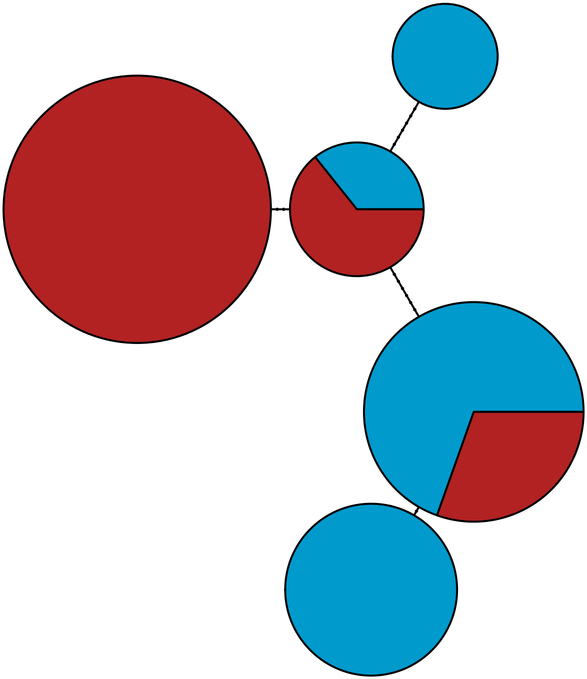

MBPA_000223760 Stramenopiles

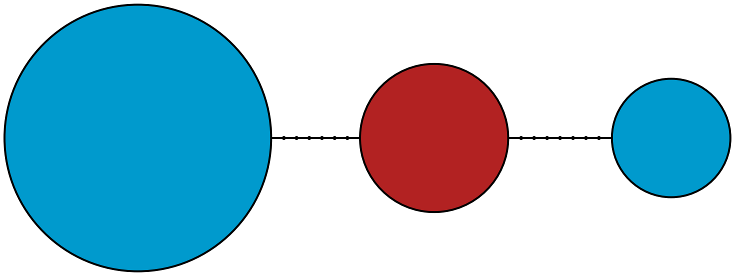

MBPA_000224754 Hacrobia

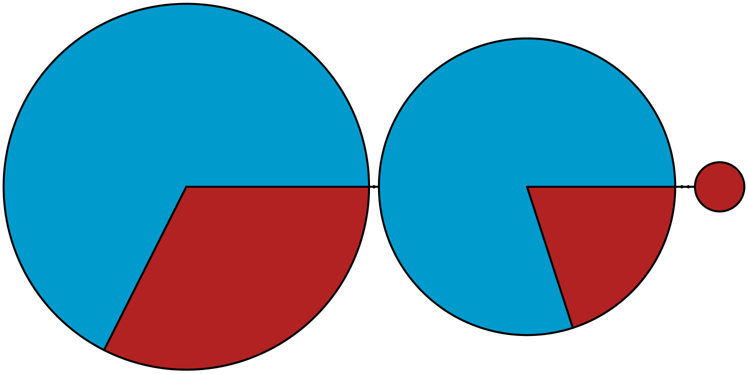

MBPA_000228037 Metazoa

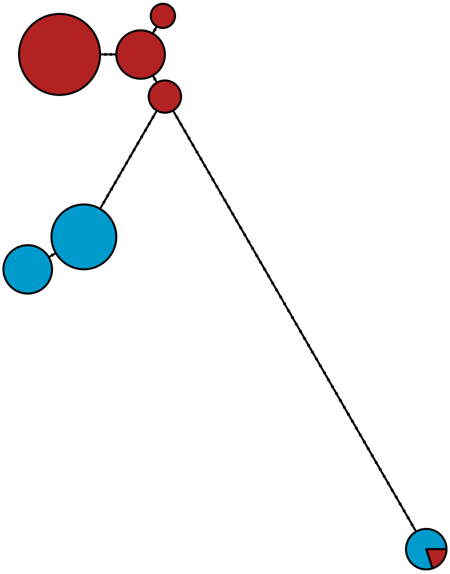

### MBPA_000228040 Metazoa

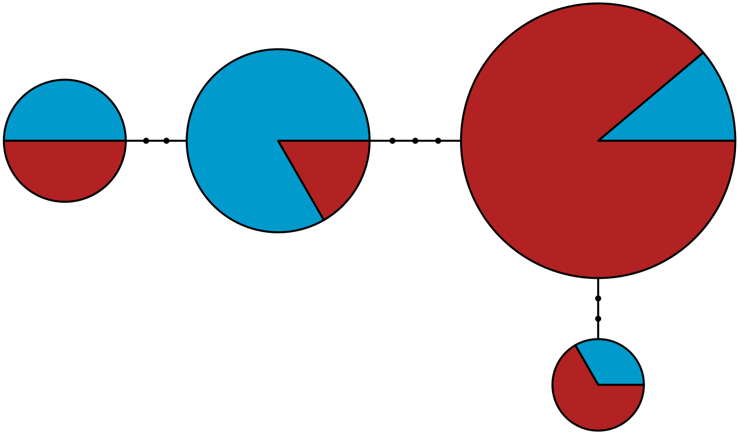

MBPA_000230114

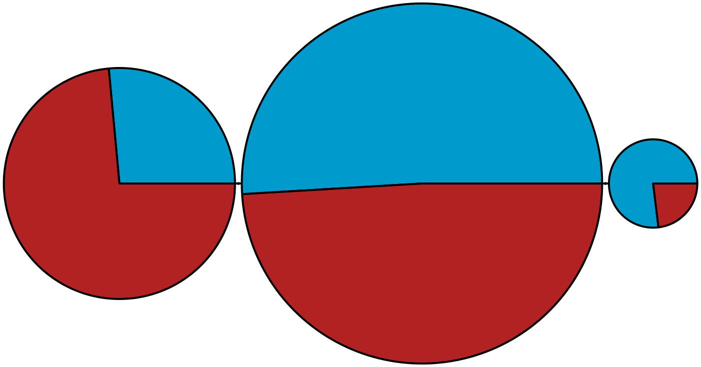

MBPA_000242347 Rhodophyta

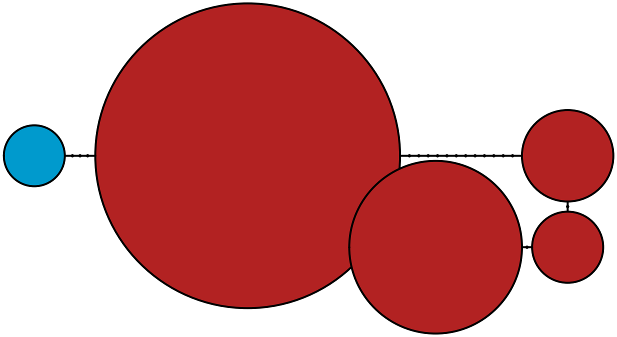

MBPA_000261919 Metazoa

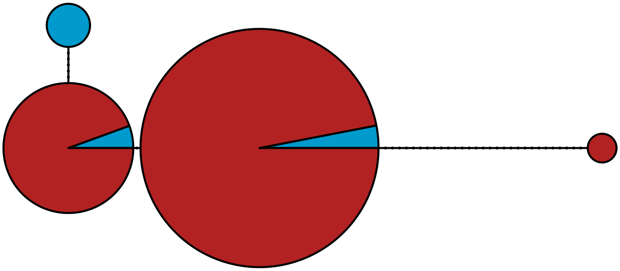

MBPA_000268858 Viridiplantae

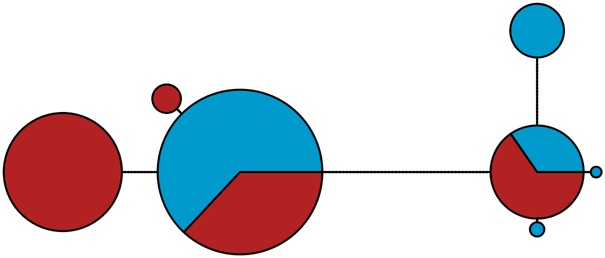

MBPA_000268936 Metazoa

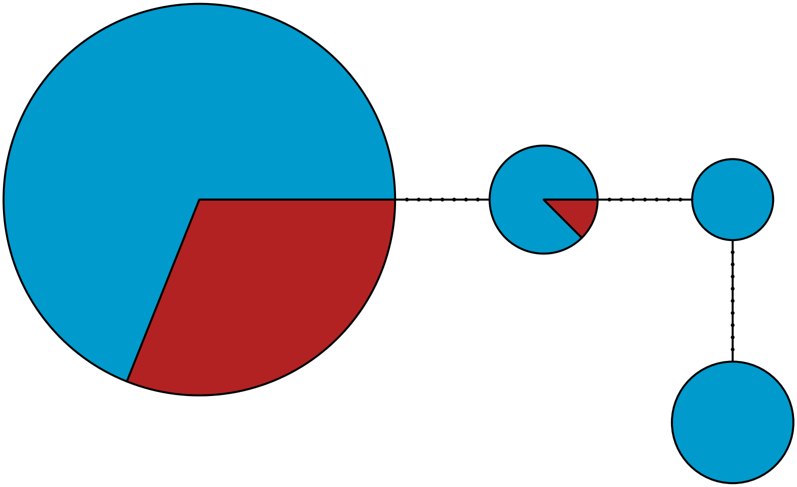

MBPA_000278260 Metazoa

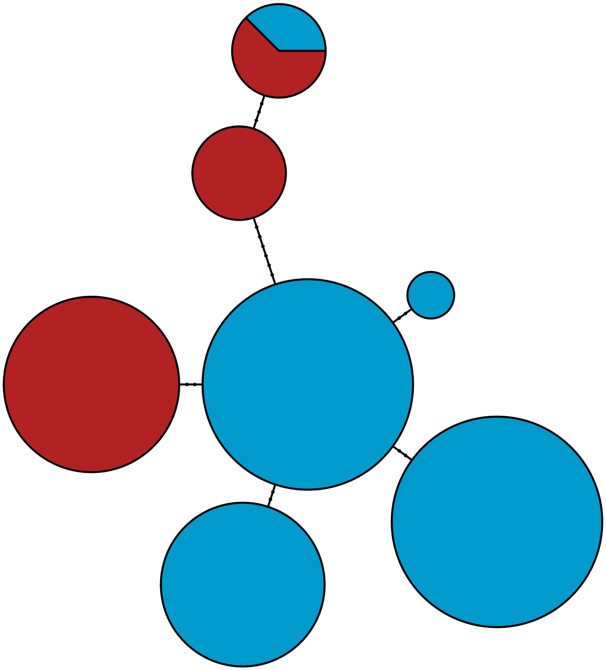

MBPA_000290334 Metazoa

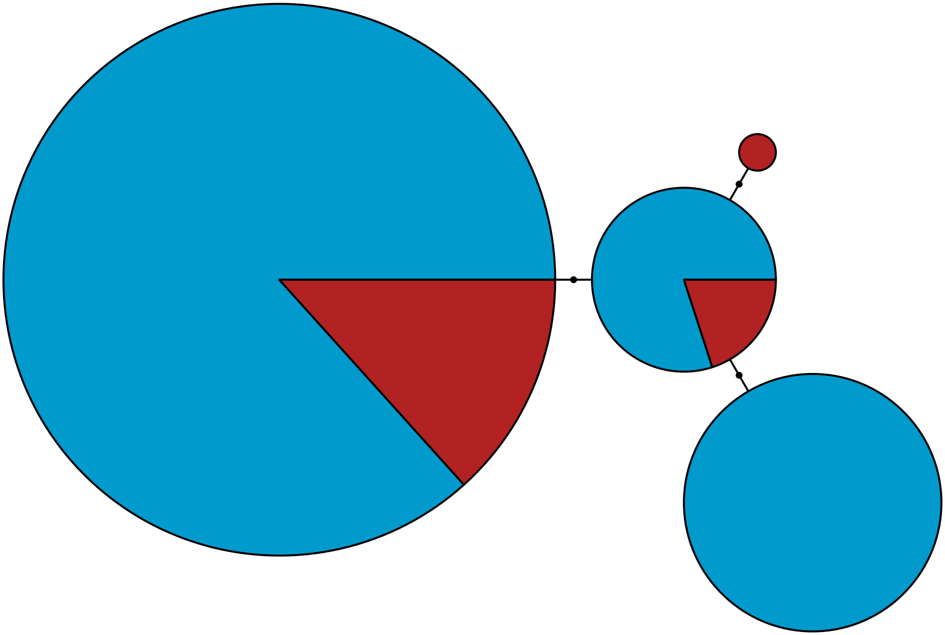

MBPA_000298014 Rhodophyta

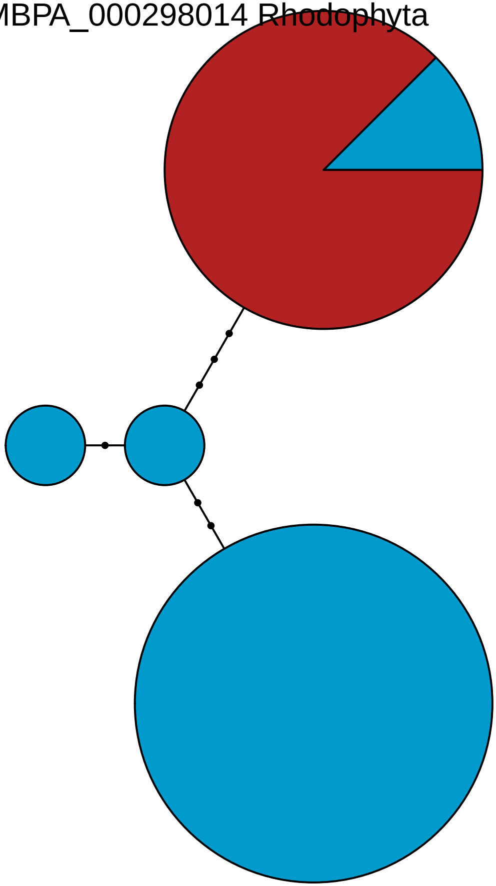

MBPA_000300574

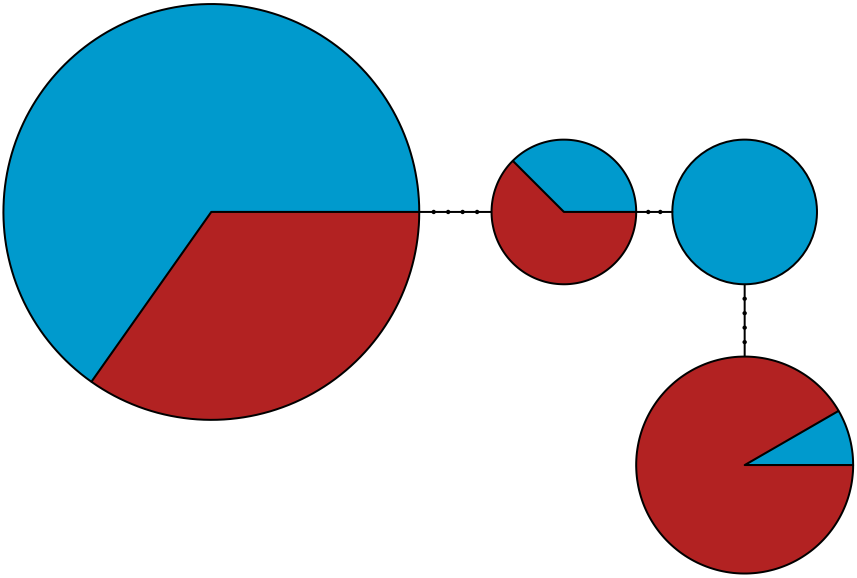

MBPA_000307595 Metazoa

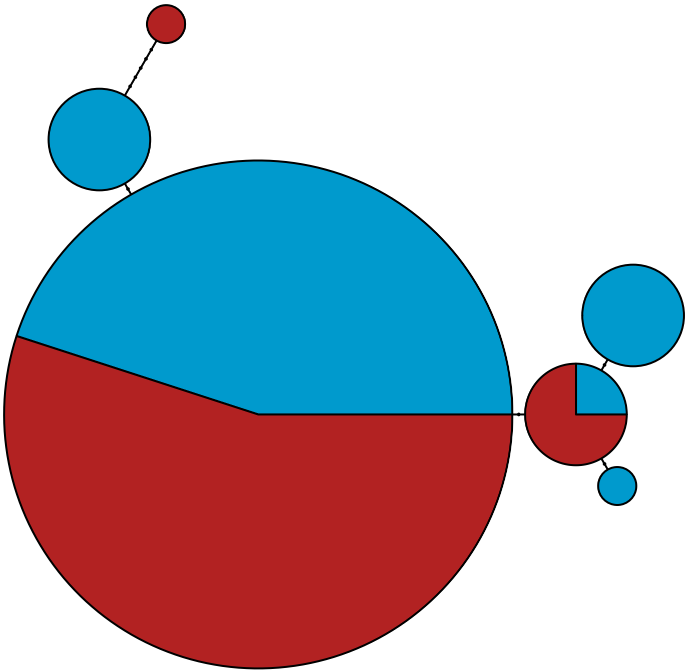

MBPA_000326515

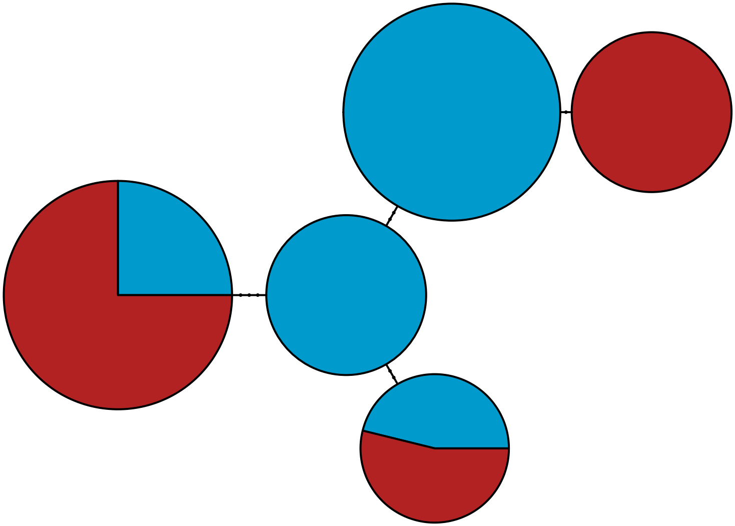

MBPA_000330949

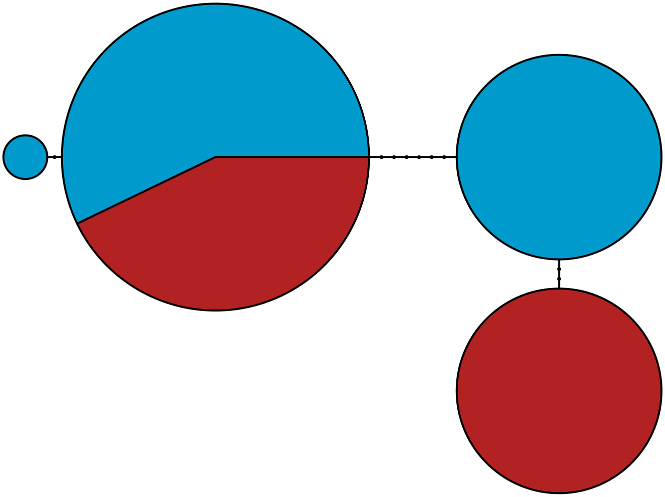

MBPA_000341561 Hacrobia

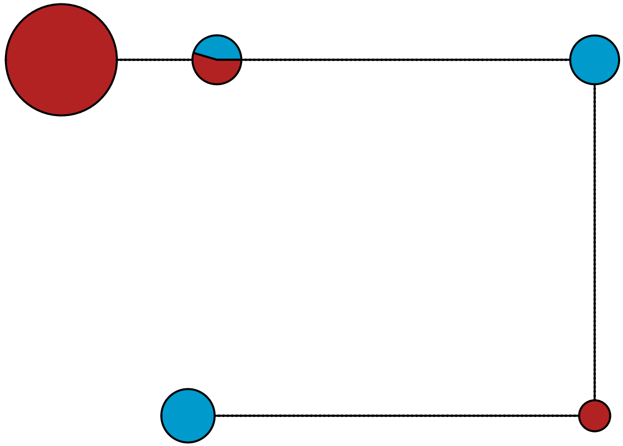

MBPA_000354144 Stramenopiles

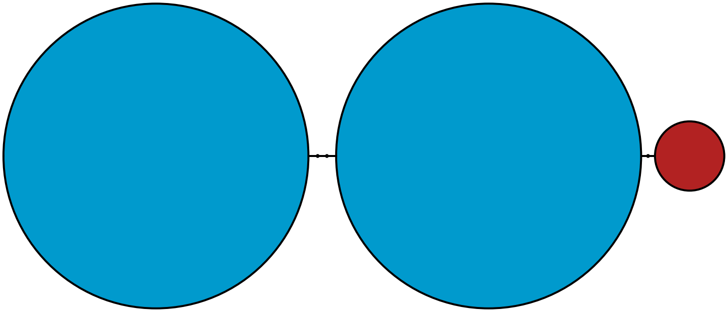

MBPA_000389095 Metazoa

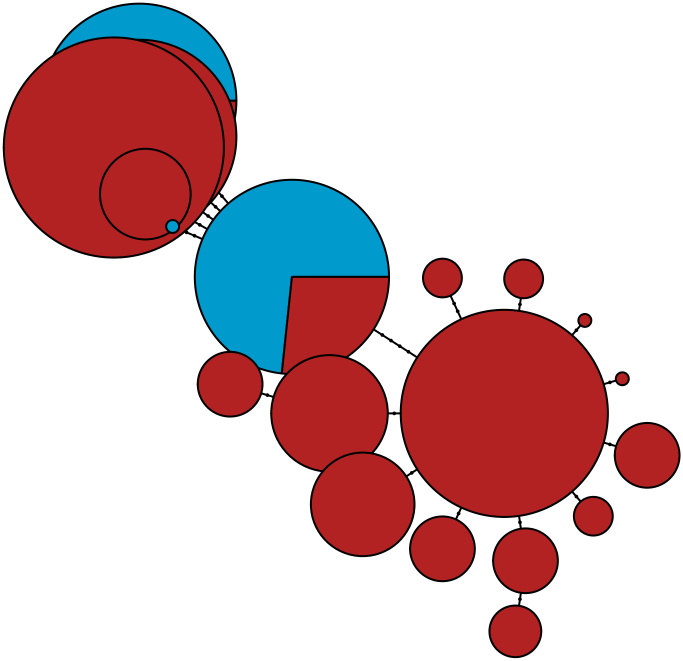

MBPA_000451838

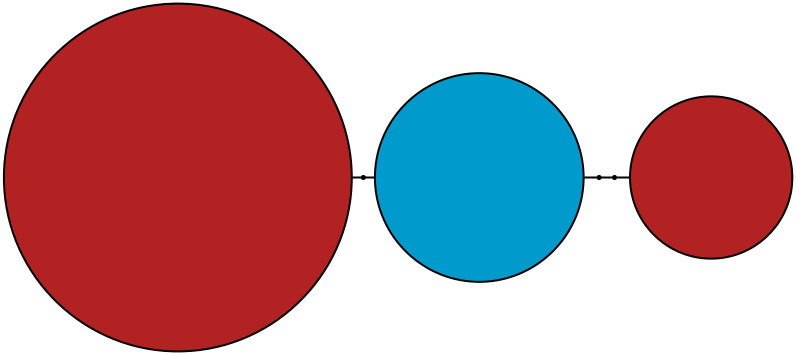

MBPA_000464129 Metazoa

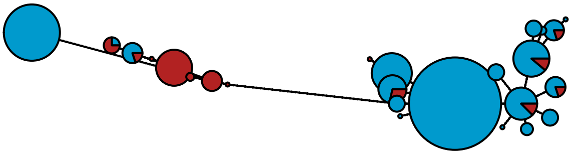

MBPA_000533475

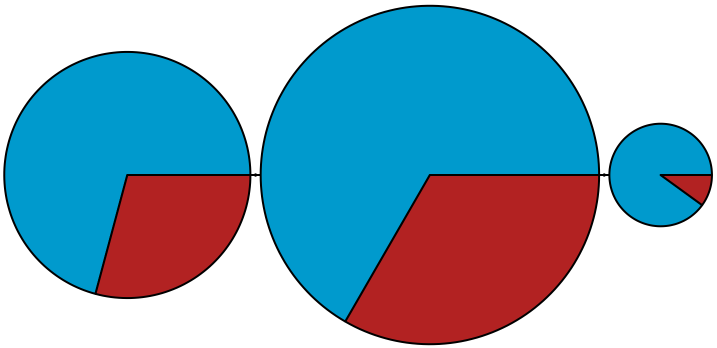

### MBPA_000556318 Stramenopiles

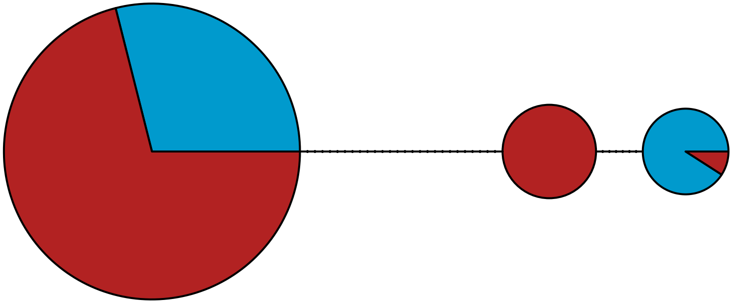

MBPA_000619797

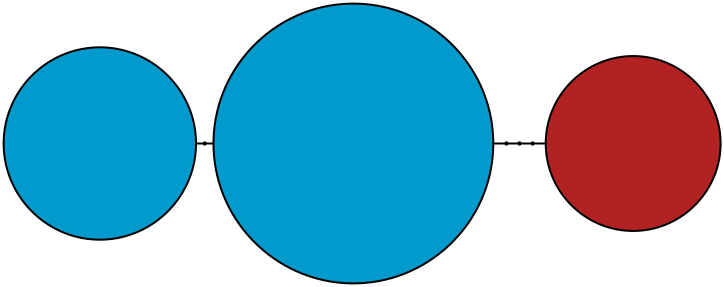

MBPA_000692213

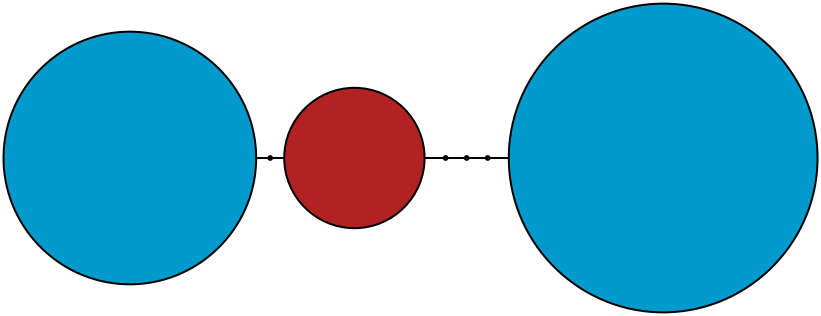

MBPA_000919505 Metazoa

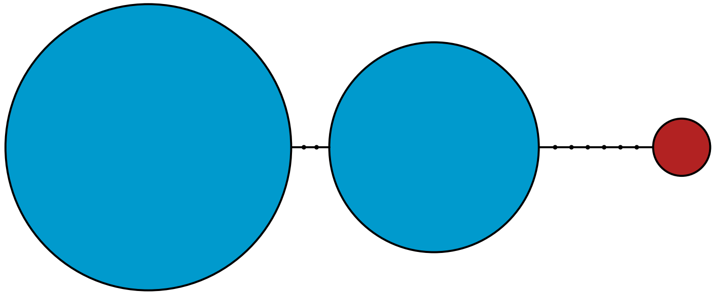

MBPA_001053177

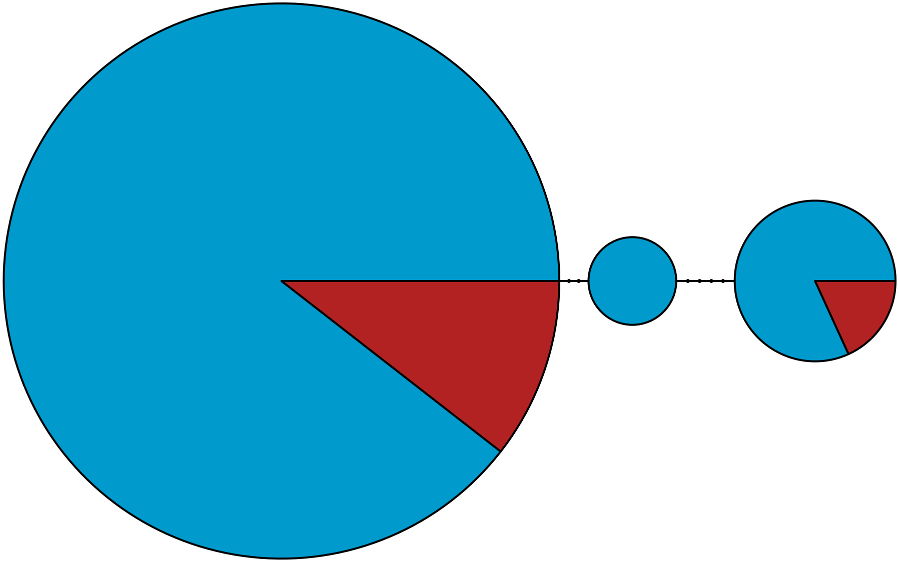

MBPA_001091579

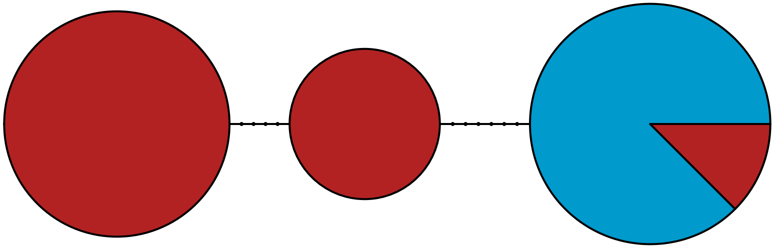

MBPA_001124203 Stramenopiles

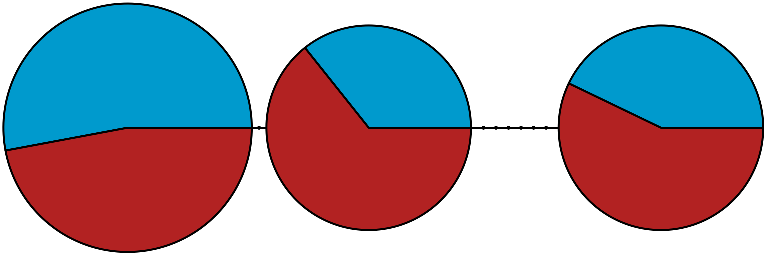

MBPA_001241510 Stramenopiles

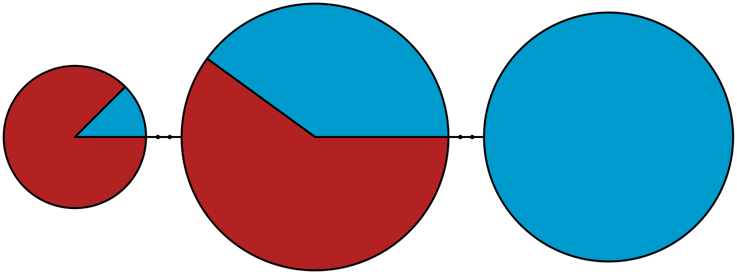

MBPA_003736101 Metazoa

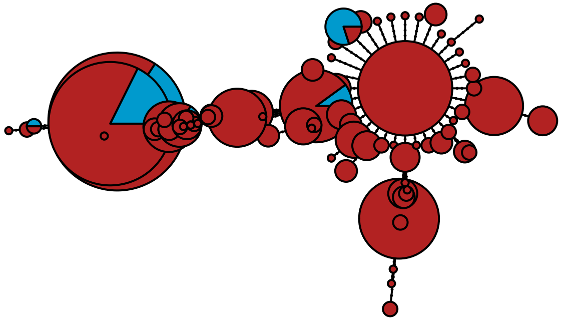

MBPA_003736115 Metazoa

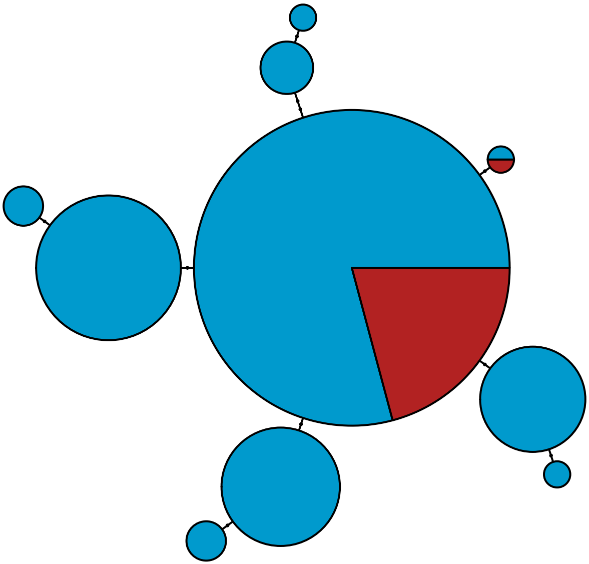

MBPA_003736743 Metazoa

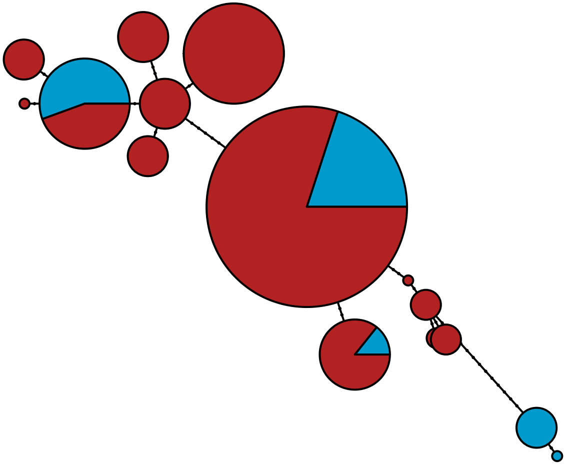

MBPA_003737888 Metazoa

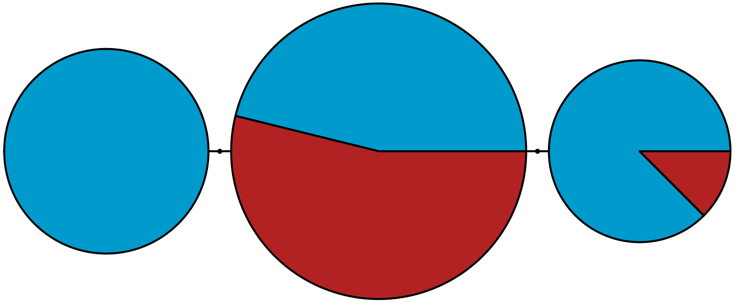

MBPA_003742170 Metazoa

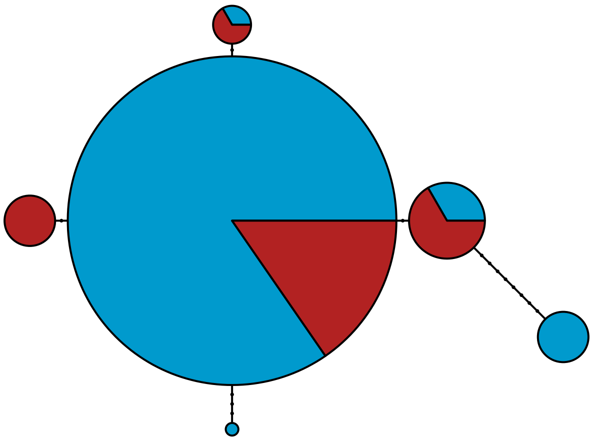

MBPA_003745411 Metazoa

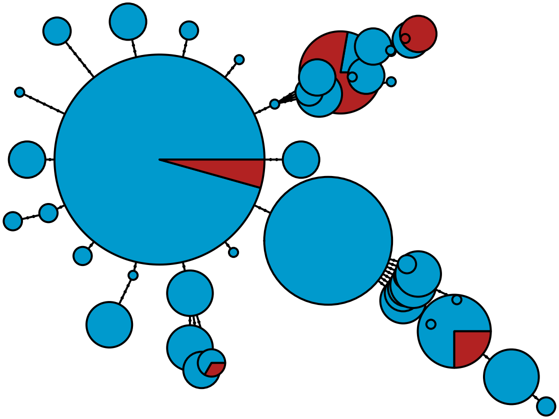

MBPA_003756087 Metazoa

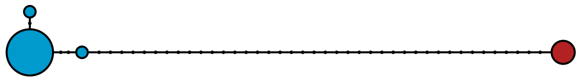

MBPA_003782205 Metazoa

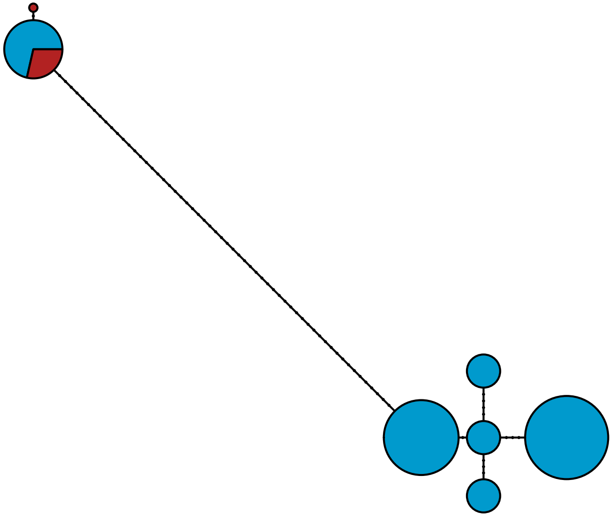

MBPA_003938994

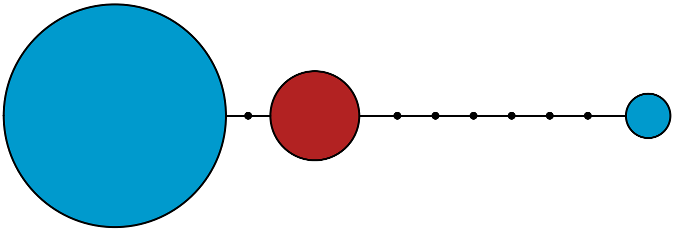
